# Hierarchical TPR:ZC3HC1 interactions drive stepwise assembly of serially arrayed nuclear basket appendages extending into vertebrate cell nuclei

**DOI:** 10.64898/2026.10.08.757499

**Authors:** Philip Gunkel, Mark Bates, Daniel Härtter, Volker C. Cordes

## Abstract

The nuclear basket (NB) is a fibrillar scaffold anchored to the nuclear pore complex (NPC), with protein TPR constituting its structural backbone. Recently, ZC3HC1 was identified as an additional structural NB component interconnecting TPR subpopulations. However, it had remained unclear how ZC3HC1 and TPR engage each other and how the resulting ensembles are arranged relative to the NB. Here, we uncover a hierarchy of sequential interactions between distinct ZC3HC1 and TPR domains, resulting in serially arrayed TPR:ZC3HC1 ensembles appended to the NB. Projecting into the nucleus, these ensembles extend beyond the NB dimensions traditionally considered prototypic. We also find that the copy numbers of NPC/NB-associated TPR are far higher than commonly anticipated and sufficient to support this modular organizational principle. Together with sufficient ZC3HC1 copy numbers, our data define a model in which alternating recruitment of ZC3HC1 and TPR extends the NB’s spatial and functional reach into the nuclear interior.

## Introduction

The nuclear basket (NB), an eightfold-rotationally-symmetric structure formerly described as fish trap-like (e.g., Maul, 1976; Jarnik and Aebi, 1991; Ris, 1991), is composed of delicate fibrils attached to the nuclear ring (NR) of the nuclear pore complex (NPC). NBs are likely to be present in the vast majority of phyla, as clear homologs of the NB scaffold-forming protein TPR, described below, can be identified across most of them (e.g., Kuznetsov *et al*., 2002; Holden *et al*., 2014; Gunkel *et al*., 2023). Furthermore, NPC-associated TPR has been detected in a wide range of tissues and cell types with only a few exceptions (e.g., Cordes *et al*., 1997; Kuznetsov *et al*., 2002; Gunkel *et al*., 2021), further indicating a widespread presence of NBs or NB-related structures.

The currently prevailing depictions of a prototypic NB, however, derive mainly from electron microscopy (EM) images of vitellogenic amphibian oocytes, with additional EM data from avian oocytes and salivary gland cells of insects such as *Chironomus tentans* (e.g., Ris, 1989, 1991; Jarnik and Aebi, 1991; Kiseleva *et al*., 1996; Goldberg *et al*., 1997). Such cells have virtually no nuclear envelope (NE)-associated chromatin and their nuclei are large enough for manual dissection, which allows their NBs to be readily exposed and then visualised as three-dimensional, unsectioned structures by conventional transmission EM (TEM) or scanning EM (SEM).

In such specimens, the NB fibrils project rectilinearly from the NPCs into the nuclear interior, and for preparations without additional heavy-metal coating, the fibrils were recurrently described as being ≤6 nm in apparent diameter (e.g., Franke, 1970, 1974a; Franke and Scheer, 1970a, 1970b; Scheer, 1972; Cordes *et al*., 1993; Ris, 1997; Gunkel *et al*., 2021). Towards their distal end, the fibrils in such EM specimens appear to branch and interlink laterally, forming a structure referred to as the terminal ring (TR; e.g., Jarnik and Aebi, 1991; Ris, 1991; Goldberg and Allen, 1996; Gunkel *et al*., 2021). A TR, however, may not occur in all cell types and perhaps not even at every NPC within the same nucleus, as suggested by early ultrastructural studies on cultured mammalian cells, which described the NPC-associated fibrils as either closed (“fish-trap-like”) or open-ended (e.g., Maul, 1971, 1976, 1977), with even some early models depicting the NR-attached fibrils as not interconnected (Franke, 1970, 1974a; Franke and Scheer, 1970a). More recently, cryo-electron tomography of murine 3T3 cells did not reveal distal lateral interconnections between NB fibrils either, modelling them instead as thick (∼10 nm) standalone struts (Singh *et al*., 2024). On the other hand, cryo- electron tomographs of *Dictyostelium* nuclei have shown NBs with a TR (Beck *et al*., 2004, 2007). Thus, it remains unclear whether the TR represents a genuine structural feature only in certain cell types, a structure only assembled on demand, or an artefact arising during specimen preparation.

In addition, one or several NB-like structures (NBLS) can be seen appended to the distal part of some of the prototypic NBs in amphibian and avian oocytes. When several NBLS occur stacked on top of each other, the resulting arrangement resembles a cylindrical extension projecting deeper into the nucleoplasm (e.g., Franke, 1974a, 1974b; Scheer *et al*., 1988; Ris and Malecki, 1993; Goldberg *et al*., 1997; Gunkel *et al*., 2021). While such NB-associated cylinders do not appear to exist in *Chironomus tentans* salivary gland cells (e.g., Goldberg and Allen, 1996; Soop *et al*., 2005), structures corresponding to the oocyte’s cylindrical NB:NBLS ensembles have been proposed to also occur in human somatic cells. There, they would correspond to the NPC-associated chromatin exclusion zones (CEZs) of extended cylindrical shape that can even be visualised in HeLa cells (Krull *et al*., 2010).

Among the first proteins described as being located at the NPC-attached nuclear fibrils were TPR in vertebrates (Cordes *et al*., 1997; Frosst *et al*., 2002) and its homologs Mlp1p and Mlp2p in budding yeast (Strambio-de-Castillia *et al*., 1999; Kosova *et al*., 2000). TPR homologs were also identified in insects, plants, and numerous other species across the eukaryotic realm (e.g., Zimowska *et al*., 1997; Kuznetsov *et al*., 2002; Jacob *et al*., 2007; Xu *et al*., 2007; Holden *et al*., 2014; Mitic *et al*., 2022; Gunkel *et al*., 2023). Models eventually proposing TPR as the fibril-forming scaffold protein of the NBs in vertebrates, insects and budding yeasts (Krull *et al*., 2004, 2010; Soop *et al*., 2005; Niepel *et al*., 2013; Gunkel *et al*., 2021) were supported by the loss of all NPC-associated CEZs in TPR-depleted HeLa cells (Krull *et al*., 2010) and by the absence of NB-like structures in Mlp1p/Mlp2p-deficient yeast (Niepel *et al*., 2013). Likewise, selective NB disassembly in *Xenopus* oocytes was accompanied by detachment of NB- associated TPR (Gunkel *et al*., 2021), but not of other NPC proteins that had previously been considered structural NB components.

Various functions have been attributed to TPR. In animal cells, TPR has been implicated in chromatin organisation, transcription regulation, replication, and senescence (Mendjan *et al*., 2006; Skaggs *et al*., 2007; Krull *et al*., 2010; David-Watine, 2011; Yap *et al*., 2012; Lelek *et al*., 2015; Zhang *et al*., 2016; Boumendil *et al*., 2019; Kosar *et al*., 2021; Uhlířová *et al*., 2021; Wu *et al*., 2021; Bartlett *et al*., 2024; Liu *et al*., 2026), as well as in autophagy (Funasaka *et al*., 2012; Dewi *et al*., 2021), DNA repair (Duheron *et al*., 2017), in meiotic recombination (Wu *et al*., 2025), and in promoting or controlling nuclear export of different classes of molecules (Bangs *et al*., 1998; Shibata *et al*., 2002; Soop *et al*., 2005; Skaggs *et al*., 2007; Coyle *et al*., 2011; Rajanala and Nandicoori, 2012; Fish *et al*., 2019; Aksenova *et al*., 2020; Lee *et al*., 2020; Zuckerman *et al*., 2020; Chen *et al*., 2021; Li *et al*., 2021; Tingey *et al*., 2022; Bhat *et al*., 2023; Cao *et al*., 2024). TPR may act in these processes directly or by serving as an operational platform for other NB-resident proteins. For example, TPR provides one of the NPC-positioned docking sites for GANP, the scaffold protein of the TREX-2 complex (Umlauf *et al*., 2013; Wickramasinghe *et al*., 2014; Aksenova *et al*., 2020; Gunkel and Cordes, 2022), which plays a major role in the export of naturally intronless transcripts (Wickramasinghe *et al*., 2010; Aksenova *et al*., 2020; Woldegebriel *et al*., 2020; Bhat *et al*., 2023). TPR also acts as a docking site for various other proteins, among which are the SUMO protease SENP1 (Schweizer *et al*., 2013; Duheron *et al*., 2017; Gunkel and Cordes, 2022), the ubiquitin E3 ligase COP1/RFWD2 (Yi *et al*., 2006; Ouyang *et al*., 2020), and the spindle assembly checkpoint proteins MAD1 and MAD2 during interphase (Lee *et al*., 2008; Schweizer *et al*., 2013; Cunha-Silva *et al*., 2020; Jackman *et al*., 2020; Gunkel and Cordes, 2022). However, these TPR-interacting proteins are commonly not considered needed for NB assembly and they are not required for keeping TPR or its yeast homologs positioned at the NPC (e.g., Lee *et al*., 2008; Mossaid *et al*., 2020; Bensidoun *et al*., 2022).

The predominant isoform of human TPR is a 267 kDa protein (e.g., Cordes *et al*., 1997; Kuznetsov *et al*., 2002) comprising two major domains: The carboxy-terminal domain (CTD; ∼77 kDa) was shown to be mostly unstructured and considered highly flexible (Hase *et al*., 2001). Among several evolutionarily conserved features, it contains a conspicuous, acidic residue cluster of so far unknown function (Hase *et al*., 2001; Kuznetsov *et al*., 2002). By contrast, the amino-terminal domain (NTD; ∼190 kDa), also termed the rod domain, forms coiled-coil (CC) homodimers of rod-like appearance (Hase *et al*., 2001), with similar CC structures also described for *Sc*Mlp1p (Kosova *et al*., 2000; Stankunas and Köhler, 2024). Immuno-EM data indicated that the N-terminal parts of TPR’s CC homodimers are folded back onto themselves when they are anchored to the NPC, positioning their N-termini near the TR region (Krull *et al*., 2004; Gunkel *et al*., 2021). A similar hairpin-like conformation was also considered likely for the NPC-anchored Mlps in budding yeast (Niepel *et al*., 2013; Stankunas and Köhler, 2024) and was recently incorporated into models of yeast and mammalian NBs based on electron-tomographic (ET) data (Singh *et al*., 2024; Obarska-Kosinska *et al*., 2026). Different segments of these folded N-terminal regions are thought to constitute both the NPC- binding domain and the rectilinear parts of the NB fibrils (Krull *et al*., 2004; Gunkel *et al*., 2021; Singh *et al*., 2024; Obarska-Kosinska *et al*., 2026). In contrast, the distal parts of TPR’s rod domain, which can also form CCs (Hase *et al*., 2001), were suggested to constitute the lateral interconnections referred to as the TR (Krull *et al*., 2004; Gunkel *et al*., 2021), which in these TR-containing NB models had been described as a delicate zigzag-like arrangement in *Xenopus* oocytes (Goldberg and Allen, 1996; Gunkel *et al*., 2021). In mammalian cells, however, in which no TR had been detectable in the cryo-ET data, also the distal parts of TPR’s rod domain were mostly assigned to the actual NB fibrils (Singh *et al*., 2024; Obarska-Kosinska *et al*., 2026).

Besides TPR, a protein named ZC3HC1 is the only other bona fide NB component currently known to have a structural role at the NB. Human ZC3HC1 is a ∼55 kDa protein comprising two similarly built and evolutionary conserved zinc ion-coordinating modules collectively termed the nuclear basket interaction domain (NuBaID), whose integrity is essential for its interaction with TPR and the NB (Gunkel *et al*., 2023). ZC3HC1 binds to TPR polypeptides already anchored to the NPC, which positions ZC3HC1 at the distal end of the NB fibrils (Gunkel *et al*., 2021). At this site, ZC3HC1 can then recruit additional TPR polypeptides and link them to those already part of the NB (Gunkel and Cordes, 2022).

The amount of TPR added to the NB in this way can be substantial. In several human cell lines of different germ layer origin, approximately 45–55% of all NPC/NB-associated TPR was found to be positioned there in a ZC3HC1-dependent manner (Gunkel *et al*., 2021). In *S. cerevisiae*, Pml39p (for pre-mRNA leakage, 39 kDa; Palancade *et al*., 2005; Hashimoto *et al*., 2022), noted to be the yeast ZC3HC1 homolog (Gunkel *et al*., 2021), likewise contains a NuBaID and can also interconnect subpopulations of Mlp1p (Gunkel *et al*., 2023).

In the current study, we addressed central questions regarding the ZC3HC1:TPR relationship at the NB. Specifically, we asked how ZC3HC1 engages apparently distinct TPR subpopulations, in what order these interactions occur, how many copies of TPR and ZC3HC1 are actually positioned at the NB and directly involved in these interactions, and, in particular, where the large subpopulations of ZC3HC1-dependent TPR are positioned relative to the NB. To this end, we mapped the ZC3HC1-binding domains (ZBDs) in TPR, identified additional ZC3HC1 elements required for recruiting TPR polypeptides, and unravelled the hierarchy of interactions among these domains. Modeling the identified interactions using AlphaFold3 (Abramson *et al*., 2024) corroborated our experimental findings. In addition, we determined the relative amounts and absolute copy numbers of NB-associated ZC3HC1 and TPR in HeLa cells, to assess which structural configurations are in principle numerically feasible, and to relate these values to the relative amounts of TPR and ZC3HC1 previously determined in HeLa and several other human cell types (Gunkel *et al*., 2021). Finally, we clarified how ZC3HC1 and distinct TPR subpopulations are positioned relative to one another at the NB, using conventional super-resolution microscopy and state-of-the-art single-molecule localisation microscopy.

Together, our data reveal an organisational principle by which sequential TPR–ZC3HC1 interactions give rise to regular NB-like appendices built onto the NB proper. The results further suggest that the prevailing view of the NB in vertebrate somatic cells as a structure restricted to the classically depicted NB proper is incomplete. Instead, our data allow us to propose a model in which the NB is an extendable structure, to which ZC3HC1:TPR ensembles can be appended as one or several serially arranged NBLSs that extend into the nuclear interior and thereby expand the spatial and functional reach of NPC-associated TPR.

## Results

### ZC3HC1 interacts with two distinct ZBDs of TPR’s rod domain

To determine the TPR regions capable of interacting with ZC3HC1, we performed yeast two- hybrid (Y2H) experiments. We deemed this approach suitable for domain mapping, because numerous segments of TPR that covered the entire NTD in a sequence-overlapping manner were known to homodimerise in the Y2H system, indicative that CC formation can also occur between shorter fragments of TPR’s NTD (Hase *et al*., 2001). Furthermore, Y2H screening had previously enabled the isolation of *Hs*TPR cDNAs from a human Y2H library using *Hs*ZC3HC1-encoding bait vectors (Supplemental Figure S1A).

We first paired full-length *Hs*ZC3HC1 with sequence-overlapping TPR segments covering the canonical isoform of *Hs*TPR, which comprises 2363 amino acids (aa; Figure 1A). Only two segments, aa 1–175 and aa 172–651, allowed colony growth upon co-expression with ZC3HC1 under selective conditions (up to 100 mM 3-AT), indicating robust interaction in the Y2H system (Figure 1B; Supplemental Figure S1B1). By contrast, no interaction was observed with segment aa 110–377, suggesting two non-overlapping ZBDs in TPR.

**Figure 1.**
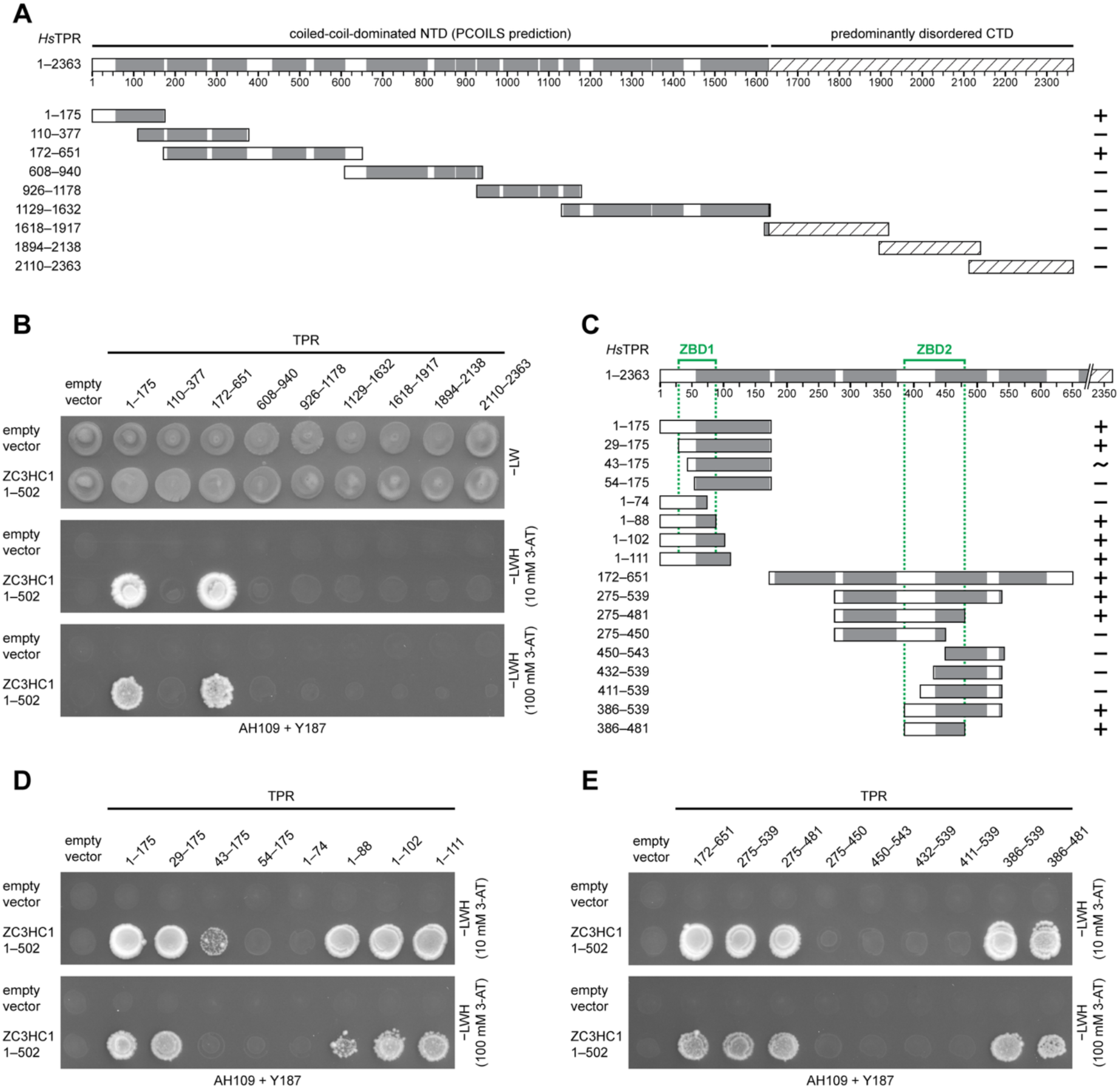
ZC3HC1 binds to two distinct segments of TPR’s NTD. **(A)** Schematics of human TPR segments encoded by Y2H vectors covering TPR’s canonical isoform of 2363 aa. Grey boxes indicate CCs as predicted by PCOILS (≥80% probability within windows of 28 aa; (Zimmermann *et al*., 2018). The hatched box represents the CTD. TPR segments had been fused at their NT to GAL4-AD. These prey plasmids had been transformed into MATα yeast cells, followed by mating with MATa yeast cells transformed with a bait vector expressing either GAL4-BD alone or a fusion of GAL4-BD to the NT of full-length *Hs*ZC3HC1. Y2H interactions with ZC3HC1 (shown in B) are indicated for each segment (+, colony growth; −, no colony growth). **(B)** Representative colony growth on medium lacking leucine and tryptophan (−LW), revealing successful mating between the MATα and MATa yeast strains, and colony growth indicating Y2H interactions between TPR segments and ZC3HC1, after replica-plating onto medium lacking leucine, tryptophan, and histidine (−LWH), here supplemented with 10 or 100 mM 3-AT. **(C)** Additional hsTPR segments encoded by Y2H vectors, used for fine-mapping ZBD1 and ZBD2. Y2H interactions with ZC3HC1 (shown in D and E) are indicated (+, colony growth; -, no colony growth; ∼, notably attenuated colony growth). Green brackets and green dashed lines mark the mapped boundaries of the essential parts of each ZBD. (**D, E**) Visualisation of Y2H interactions after replica-plating onto −LWH medium supplemented with 10 mM 3-AT (see Supplemental Figure S1 for corresponding −LW plates). For ZBD1 (D), colony growth was observed only with segments near the canonical NT that included at least aa 29–88. For ZBD2 (E), colony growth required at least aa 386–481, encompassing part of TPR’s formerly defined NBD.

Next, we mapped these domains in greater detail. The shortest TPR segments that still allowed ZC3HC1 interactions as robust as those with longer segments were aa 1–102, 29–175 and 386–481, while further truncations either attenuated or abolished colony growth (Figure 1C–1E; Supplemental Figure S1C1 and S1D1). Similar results were also obtained with a distinct pair of yeast strains (Supplemental Figure S1B2, S1C2, and S1D2). Furthermore, ZC3HC1 did not interact with the N-terminal extension of the non-canonical, second isoform of TPR, here termed TPR-L, which in humans comprises 76 additional aa, of which 24 are encoded by an exon upstream of the commonly used start codon (Supplemental Figure S1E and S1F).

Altogether, we concluded from these data that the essential residues of TPR’s first and second ZBD lie between aa 29–88 and 386–481, respectively, while some additional, flanking residues may only contribute, either directly or indirectly, to strengthening the interaction with ZC3HC1.

### The ZBD1 of NPC-anchored TPR provides the initial binding site for ZC3HC1 at the NB

Starting with ZBD1, we next investigated whether the two ZBDs identified by Y2H assays constitute physiological binding interfaces for ZC3HC1 at the NB. Previously, NB-resident ZC3HC1 had been localised at the TR (Gunkel *et al*., 2021), adjacent to the N-termini of NPC- anchored TPR (Supplemental Figure S2A–S2C; Krull *et al*., 2004; Gunkel *et al*., 2021). This co-localisation suggested that ZBD1 provides the initial docking site for ZC3HC1 at the NB.

To test this assumption, we employed CRISPR/Cas9 nickase (Ran et al. 2013a) to delete ZBD1-encoding sequences of the *TPR* gene in HeLa cells. According to the Y2H data, ZBD1 requires at least aa 43–88 (Figure 1D; Supplemental Figure S1C1), encoded across three exons corresponding to aa 1–50, 51–85, and 86–109 of the canonical *Hs*TPR isoform (Supplemental Figure S1E). Using the well-characterised HeLa P2 cell line (Gunkel *et al*., 2021; Gunkel and Cordes, 2022), we deleted three segments of exon 1 as well as the entire ORF of exon 1 and the 5’ portion of exon 2, thereby removing the ∼1.4 kb intron between exons 1 and 2. Each deletion was replaced in frame with the ORF for superfolder GFP (sfGFP; e.g., Pédelacq *et al*., 2006; Figure 2A1 and 2A2; Supplemental Figure S2D1-S2D3). Because TPR’s leucine zipper, possibly with a particular role in TPR homodimerisation, begins at aa 78 (Rodrigues and Park, 1993; Pal *et al*., 2017), exon 3 and most of exon 2 were left untouched.

**Figure 2.**
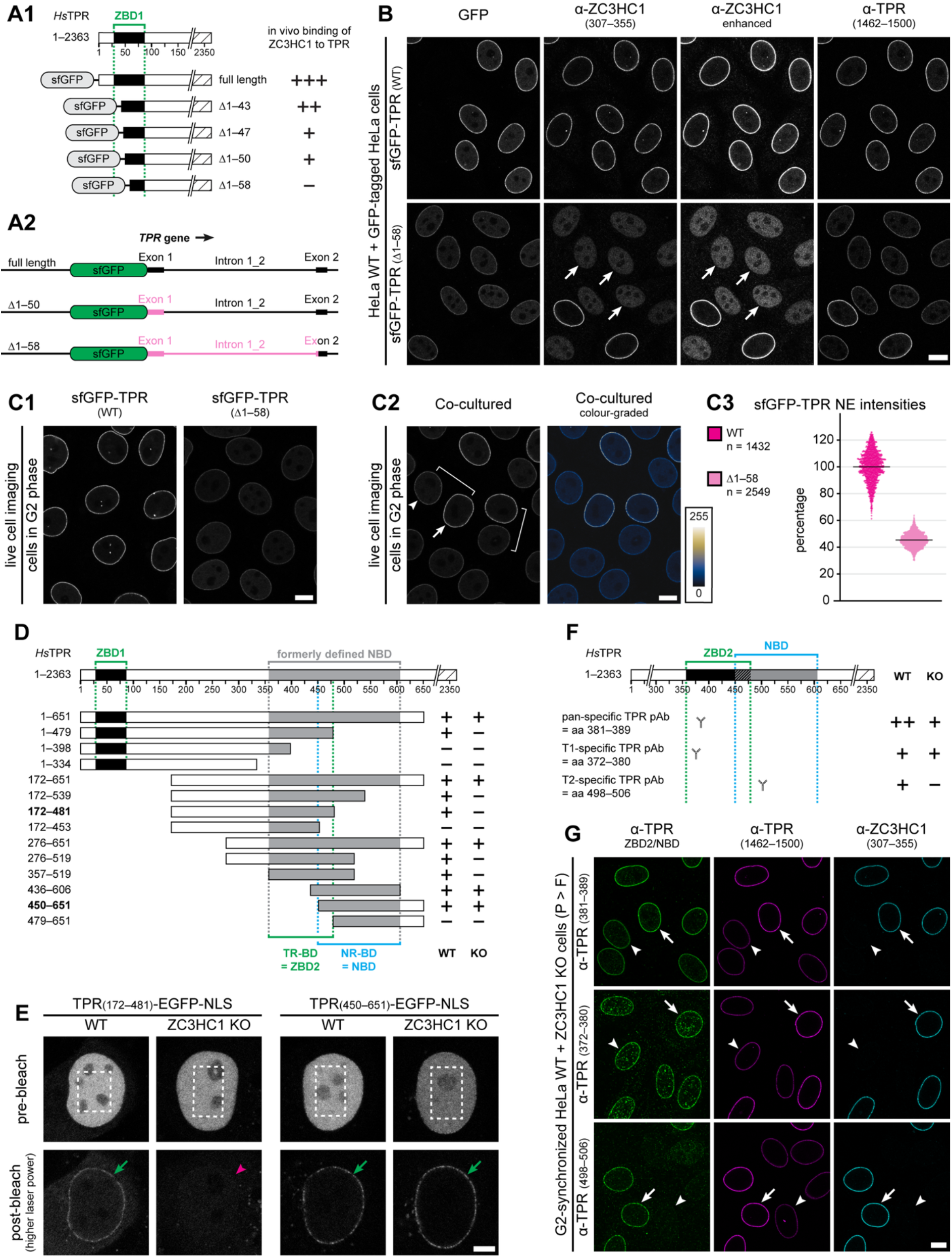
TPR’s ZBD1 provides the initial binding site for ZC3HC1 at the NB, NB-bound ZC3HC1 appends additional TPR via ZBD2, and TPR subpopulation-specific antibodies distinguish NPC-anchored from ZC3HC1-appended TPR. **(A)** Schematics of (A1) the sfGFP-tagged N-termini of WT TPR polypeptides and truncated variants expressed in HeLa cell lines, and (A2) corresponding regions of the CRISPR/Cas9-edited *TPR* gene, with deleted parts labelled in magenta and lengths drawn to scale. The capability of attracting ZC3HC1 to the NE is indicated (+++, bright ZC3HC1 signal intensities at the NE; ++, moderate; +, traces; -, not detectable). The horizontal bracket and dotted lines demarcate the ZBD1 region defined as essential in Y2H experiments. **(B)** Double-IFM of co-cultured populations of HeLa P2 WT cells with non-tagged TPR and HeLa P2 lines homozygously expressing either sfGFP-TPR or sfGFP-TPR Δ1–58. The co-cultured cells, cell-cycle-synchronised to G2, were fixed and immunolabelled with antibodies against ZC3HC1 and TPR, revealing that TPR amounts in TPR Δ1–58 cells are notably diminished and that ZC3HC1 is no longer NE-associated and instead appears as a nucleoplasmic pool (white arrows). Microscope settings were identical for both populations of co-cultured cell lines. ZC3HC1 staining is additionally shown enhanced. Corresponding pre-extracted cells are shown in Supplemental Figure S2D5. Bar, 10 µm. **(C)** Live-cell fluorescence microscopy of HeLa sfGFP-TPR and sfGFP-TPR Δ1–58 cells and quantification of sfGFP signal yields at NEs. Cells were grown either as single-type populations (C1) or as co-cultures (C2), all cell-cycle-synchronised to G2. The brackets demarcate two example pairs of a WT and a mutant cell within the co-cultured populations, with the nucleus of a WT cell additionally marked by an arrow, and an arrowhead marking the nucleus of a cell with sfGFP-TPR Δ1–58. The same micrograph is also shown colour-graded next to a colour look-up table (LUT). Bars, 10 µm. (C3) Automated image-based quantification of relative sfGFP signal yields at the NEs of neighbouring cells expressing either sfGFP-TPR or sfGFP-TPR Δ1–58, synchronised in G2 and imaged live as in C2. The arithmetic mean for the WT is set to 100%. **(D)** Schematics of expression vector-encoded deletion mutants of the canonical TPR isoform. NE-binding in WT and ZC3HC1 KO cells is indicated on the right. Grey boxes mark regions previously assigned to the NBD. Green and blue brackets with dotted lines demarcate the two overlapping domains now mapped as essential for ZBD2 (aa 357–479) and for the actual NBD (aa 450–606). Bold numbers represent segments shown in E. **(E)** Live- cell imaging and photobleaching of representative HeLa P2 WT and ZC3HC1 KO cells transiently transfected with expression vectors encoding *Hs*TPR aa 172–481 and aa 450–651. The nucleoplasmic pool of overexpressed GFP-tagged TPR was bleached (marked rectangles), followed by acquisition of a post-bleach image at higher laser power to visualise those TPR segments that had engaged with the NE. TPR aa 172–481 and aa 450–651 were detectable at the NE in WT cells, as was aa 450–651 at the NE in ZC3HC1 KO cells (green arrows), whereas aa 172–481 did not bind to the NE in ZC3HC1 KO cells (magenta arrowhead). Bar, 10 µm. **(F)** Schematic of the hsTPR segment aa 1–651 and the relative positions of three representative *Hs*TPR peptide antibodies (Y), which, in situ, bind either both the NPC-anchored and TR- attached subpopulations of TPR (pan-specific), only the NPC-anchored subpopulation (T1- specific), or only the TR-attached subpopulation (T2-specific). Relative NE signal intensities in co-cultured WT and ZC3HC1 KO cells, visualised by IFM and exemplified in G, are indicated on the right. **(G)** Triple-IFM of co-cultured HeLa WT and ZC3HC1 KO cells. The left row shows labelling with the TPR epitope-specific rb-antibodies presented in F, the middle row, as a reference, shows labelling with a TPR mAb (1462–1500) that recognises both subpopulations; and the right row shows labelling with ZC3HC1-specific gp-antibodies to distinguish WT and KO cells. Arrows mark NEs of representative WT cells while arrowheads mark NEs of ZC3HC1 KO cells. Antibody TPR 381–389 yields NE signal intensities about twofold higher in WT than in KO cells. By contrast, antibody TPR 372–380 labels WT and KO cell NEs at similar intensity, indicating labelling of only the T1 subpopulation, whereas the KO cell NEs remain unlabelled with antibody TPR 498–506, indicating labelling of only the T2 subpopulation. Bar, 10 µm.

Cell proliferation and viability of the resulting homozygous lines (TPR Δ1–43, Δ1–47, Δ1– 50, Δ1–58) were not noticeably affected under standard cell culture conditions, and the truncated TPR polypeptides still bound similarly stably to the NPCs as intact TPR in wild type (WT) and ZC3HC1 knockout (KO) cells, exemplified for TPR Δ1–58 (Supplemental Figure S2D4 and S2D5). In the TPR Δ1–47 and Δ1–50 cells, the NB-associated amounts of ZC3HC1 were strongly reduced but remained detectable (Figure 2A1; Supplemental Figure S2E). By contrast, in TPR Δ1–58, ZC3HC1 was no longer detectable at the NB but throughout the nuclear interior instead (Figure 2B), demonstrating that ZBD1 is the exclusive docking site for ZC3HC1 at the NB.

Consistent with the complete loss of ZC3HC1 from the NB, the amounts of NE-associated TPR in the TPR Δ1–58 cells were markedly reduced (Figure 2C1). This decrease was particularly evident when the mutant cells and WT cells expressing intact TPR were co- cultured as cell-cycle-synchronised mixed populations and imaged side by side (Figure 2C2). Quantification of the NE-associated sfGFP signals for the TPR Δ1–58 mutant revealed a reduction, on average, to about 45% of WT levels. Similar results were obtained both with our conventional approach for quantifying NE-associated signals (Gunkel *et al*., 2021; Supplemental Figure S2F) and with a newly developed image analysis pipeline (Figure 2C3; Supplemental Figure S2G). In both cases, the measured reduction matched essentially that seen in ZC3HC1 KO cells (Gunkel *et al*., 2021).

### The ZBD2 of additional TPR polypeptides enables their attachment to the NB through binding to ZC3HC1

Because ZC3HC1, once bound to the ZBD1 of NPC-anchored TPR, can attract additional TPR polypeptides and append them to the NB (Gunkel and Cordes, 2022), we asked whether TPR’s ZBD2 is required for this subsequent interaction. However, CRISPR/Cas9n deletions within the ZBD2 region were unsuitable to test this, given TPR’s molecular architecture (Hase *et al*., 2001; Krull *et al*., 2004) and the proximity of other binding domains of TPR (see further below, and Gunkel *et al*., MS in preparation). We therefore chose an alternative approach and compared NE binding of short TPR segments expressed in HeLa WT and ZC3HC1 KO cells.

Earlier work had shown that homodimerising TPR segments that contain some or all parts of TPR’s NPC binding domain (NBD), as it had been previously defined (aa 357–606), can bind to the NPC in HeLa WT cells (Hase *et al*., 2001), even in the absence of endogenous TPR (Hase and Cordes, 2003). Now knowing that the ZBD2, with its essential residues aa 386–481, lies within this region, we asked whether NE attachment of some of the previously tested TPR segments reflected a ZC3HC1-dependent interaction via ZBD2 rather than ZC3HC1- independent anchorage via the genuine NBD.

To distinguish between these possibilities, we used previous expression vectors encoding Myc-tagged TPR segments and novel vectors encoding EGFP-tagged TPR segments, with all segments additionally tagged with an SV40 NLS to ensure nuclear import. Together, these segments covered the ZBD2 and historical NBD in an overlapping manner (Figure 2D). Following transient transfection of HeLa WT and ZC3HC1 KO cells, the Myc-tagged TPR segments were visualised by immunofluorescence microscopy (IFM) and the EGFP-tagged segments by live cell imaging (LCI), the latter combined with nucleoplasmic photobleaching to allow post-bleach assessment of NE localisation, as exemplified for TPR aa 172–481 and aa 450–651 (Figure 2E). Altogether, these experiments revealed that all previously NE-positive segments (Hase *et al*., 2001) again localised at the NE when expressed in WT cells, as did several of the new ones. In ZC3HC1 KO cells, however, a subset of the segments failed to bind to the NE (Figure 2D and 2E).

These data revealed that residues required for ZC3HC1-independent NPC anchorage of TPR lie between aa 450 and 606, a region that lacks most residues essential for a functional ZBD2. Conversely, TPR polypeptides that bound to the NE only in the presence of ZC3HC1 lacked parts of aa 450–606 but contained at least aa 357–479 and thus overlapped the region containing the residues essential for a functional ZBD2 (Figure 2D).

Together, these findings show that the previously defined NBD actually comprises two domains involved in positioning TPR at the NE. One is the genuine NBD, required for ZC3HC1-independent anchorage at the NPC and now mapped more narrowly to aa 450–606. The other is ZBD2, with its essential residues aa 386–481, and now proven to be required for the ZC3HC1-dependent appendage of additional TPR polypeptides to the NB. Both domains, though, appear to depend on a shared region, aa 450–479, in order to be functional (Figure 2D).

#### NPC-anchored and TR-appended TPR can be distinguished in situ by TPR- subpopulation-specific antibodies targeting different NBD and ZBD2 epitopes

Identifying TPR’s NBD-overlapping ZBD2 as the domain by which ZC3HC1 tethers additional TPR to the NB posed an apparent conflict between previous studies: ZC3HC1 had been localised at the TR and not at the NR (Gunkel *et al*., 2021; Supplemental Figure S2A– S2C), implying that ZBD2-mediated appendage of additional TPR polypeptides would place their NBDs at the TR as well. However, in an earlier study using *Hs*TPR-specific peptide antibodies for double-blinded post-embedding immuno-TEM (iTEM), the NBD had been localised at the NR, not the TR (Krull *et al*., 2004).

Clues toward an explanation came from immuno-EM of *Xenopus* oocyte NEs (Gunkel *et al*., 2021), when using a panel of epitope-overlapping peptide antibodies against the *Xl*TPR region formerly defined as the NBD. These antibodies had all been proven to be epitope- specific and to not intramolecularly cross-react with other TPR segments. When immunolabelling subpopulations of oocyte NBs that appeared to have some fibrous material collapsed onto their distal ends, which possibly had occurred during the NE isolation process, some of these antibodies, nonetheless, only labelled the NB, close to its anchor points at the NR. Others, however, labelled both NR and TR, and still others only the TR region. This pattern suggested TPR subpopulation-specific epitope masking or inaccessibility, with some epitopes affected only in the NPC-anchored subpopulation and others only in ZC3HC1-appended subpopulation, perhaps represented by the collapsed fibrillar material, while a third set of epitopes was accessible in both. In the following, the NPC-anchored TPR polypeptides will be referred to as the T1 subpopulation, whereas all ZC3HC1-appended TPR polypeptides will collectively be referred to as the T2 subpopulation.

To test whether a corresponding scenario would also apply to proliferating human cells, we raised new peptide antibodies against the *Hs*TPR region spanning the genuine NBD and ZBD2. After identifying those antibodies suitable for IFM and confirming their specificity for TPR (Supplemental Figure S2H), we compared NE labelling in HeLa WT versus ZC3HC1 KO cells. Because the KO cells lack the NE-associated T2 subpopulation, we reasoned that this comparison should reveal antibodies that specifically recognise T1 or T2 in situ.

The results fell into three categories: First, one group of antibodies, here exemplified by anti-TPR aa 381–389, labelled the NE in both WT and ZC3HC1 KO cells, yet with about twofold higher intensity in the WT (Figure 2F and 2G). We therefore regarded the corresponding epitopes as equally well accessible in the T1 and T2 subpopulations. This difference in labelling intensity also matched that seen with standard TPR antibodies that target epitopes outside the ZBD2-NBD region (e.g., aa 1462–1500, Figure 2G) and label T1 and T2 proportionally in vitro and in situ (Gunkel *et al*., 2021; Gunkel and Cordes, 2022). The second group of antibodies, exemplified by anti-TPR aa 372–380, labelled the NEs of WT and ZC3HC1 KO cells at similar levels (Figure 2G), indicating that the corresponding epitopes were labelled in the T1 subpopulation but masked or inaccessible in T2. The third group of antibodies, exemplified by anti-TPR aa 498–506, only labelled the NEs of WT but not of ZC3HC1 KO cells (Figure 2G), indicating that these epitopes were labelled in the T2 subpopulation but not in T1.

Together, these data, showing selective in situ labelling of either the T1 or T2 subpopulation of TPR, allowed us to reconcile earlier observations and now establish that the ZBD2-adjacent NBD can occur at both the NR and the NB’s distal end.

#### The NuBaID of ZC3HC1, sufficient for docking to the ZBD1 of NPC-associated TPR, requires flanking elements to recruit additional TPR polypeptides via their ZBD2

Having delineated the roles of TPR’s ZBDs, we further examined which regions of ZC3HC1 are required for its assembly with distinct TPR subpopulations. Recently, we had shown that the two BLD (BIR-like domain) modules forming the NuBaID of ZC3HC1 are essential for interacting with TPR in HeLa cells. Furthermore, several single aa substitutions within the NuBaID abolished Y2H-binding to TPR regions that we here mapped as ZBD1 and ZBD2 (Figure 1; Gunkel *et al*., 2023). However, other questions remained, including which parts of ZC3HC1 bind to ZBD2.

In Y2H experiments, a minimal, NuBaID-only ZC3HC1 variant that lacked the loop and both NuBaID-flanking regions, and even tolerated the loss of four C-terminal NuBaID residues (mutant 72–290_398–467; Gunkel *et al*., 2023), bound to ZBD1 but not to ZBD2 (Figure 3A; Supplemental Figure S3A). On the other hand, loop deletion (aa 291–397) in an otherwise intact ZC3HC1 (1–290_398–502) impaired neither binding to ZBD1 nor to ZBD2. By contrast, deletion of N-terminal residues 1–71 (72–502) markedly diminished binding to ZBD2, and removal of C-terminal residues 468–502 (1–467) abolished it. C-terminal truncations that left the NuBaID intact revealed that deletion of aa 491–502 (1–490) was sufficient to abolish Y2H- binding to ZBD2 while preserving binding to ZBD1. Binding to ZBD1 was lost only upon C- terminal truncations further into the NuBaID (e.g., mutant 1–462; Figure 3A; Supplemental Figure S3A).

**Figure 3.**
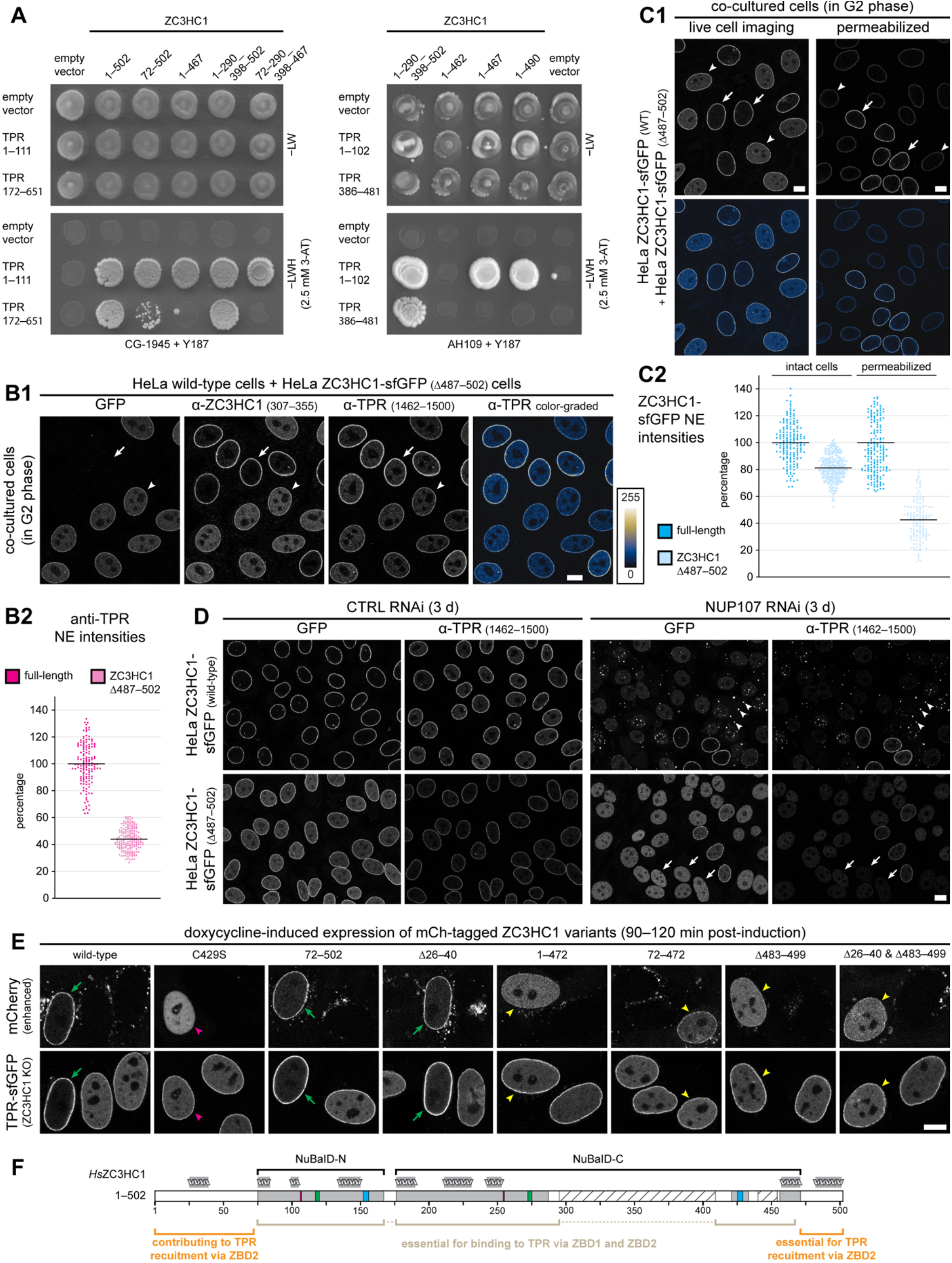
The intact NuBaID of ZC3HC1 is sufficient for binding ZBD1 but requires NuBaID-flanking elements to recruit additional TPR polypeptides via binding to ZBD2. **(A)** Y2H interaction experiments between full-length ZC3HC1 or deletion mutants thereof and TPR segments containing either ZBD1 or ZBD2. Colony growth on −LW medium and after replica-plating onto −LWH medium supplemented with 2.5 mM 3-AT is shown for a subset of pairing combinations of WT or deletion mutants of ZC3HC1 with TPR segments containing either ZBD1 (TPR 1–111, 1–102) or ZBD2 (TPR 172–651, 386–481). Equivalent results were obtained with both strain pairings (CG-1945 + Y187; AH109 + Y187). An almost complete NuBaID (72–290_398–467) suffices for Y2H binding to ZBD1 while failing to bind ZBD2; this also holds for ZC3HC1 mutants lacking C-terminal NuBaID-flanking sequences (1–467, 1–490). **(B)** Triple-FM of a co-cultured population of G2-synchronised HeLa P2 WT cells with non-tagged ZC3HC1 and HeLa P2 cells homozygously expressing ZC3HC1-sfGFP Δ487–502, and quantification of NE-associated IF-signal yields for TPR. (B1) Cells were fixed and immunolabelled for TPR and ZC3HC1. A WT cell nucleus is marked by an arrow, while an arrowhead marks the nucleus of a representative cell with ZC3HC1-sfGFP Δ487–502, in which the NE-associated amounts of TPR are markedly reduced. The same micrograph is shown colour-graded next to an LUT. Bar, 10 µm. (B2) Automated image-based quantification of NE- associated TPR immunofluorescence in G2-synchronised cells with either non-tagged WT ZC3HC1 or ZC3HC1-sfGFP Δ487–502. The arithmetic mean for the WT is set to 100%. **(C)** Fluorescence microscopy of HeLa ZC3HC1-sfGFP and ZC3HC1-sfGFP Δ487–502 cells, and quantification of sfGFP signal yields at NEs. (C1) Cells were grown as co-cultures cell-cycle- synchronised to G2 and imaged either as live cells or after permeabilisation, followed by fixation. The nuclei of two WT cells are marked by arrows, while arrowheads mark the nuclei of two cells with ZC3HC1-sfGFP Δ487–502. The same micrographs are also shown colour- graded, as in B1. Bars, 10 µm. (C2) Automated image-based quantification of sfGFP signal yields at the NEs of neighbouring cells with ZC3HC1-sfGFP or ZC3HC1-sfGFP Δ487–502, synchronised in G2 and imaged as in C1. **(D)** HeLa cells with either ZC3HC1-sfGFP or ZC3HC1-sfGFP Δ487–502 after treatment with control siRNAs and NUP107 siRNAs, and subsequent immunolabelling for TPR. A few cells that remained non-transfected with the NUP107 siRNAs are shown as reference and are marked by arrows in magenta. In the NUP107 siRNA-transfected cells, NE labelling for TPR and both ZC3HC1 variants was often no longer detectable, with some of the TPR-deficient NEs in the ZC3HC1-sfGFP cells marked by arrows in white. In such NUP107-deficient cells, numerous TPR- and ZC3HC1-positive foci were present instead, with some marked by white arrowheads. By contrast, in NUP107-deficient cells with ZC3HC1-sfGFP Δ487–502, such foci were not detectable; instead, both TPR and ZC3HC1-sfGFP Δ487–502 appeared uniformly distributed throughout the nucleus, excluding the nucleoli. Bar, 10 μm. **(E)** Live-cell fluorescence microscopy of cell-cycle-synchronised HeLa P2 ZC3HC1 KO cells homozygously expressing TPR-sfGFP and transiently transfected with a Tet-On-inducible expression vector encoding mCherry-tagged WT or mutant versions of ZC3HC1. Cells were imaged 120–150 min after induction with doxycycline. Nuclei in which the nucleoplasmic pool of TPR-sfGFP appeared largely re-appended to the NE upon ectopic expression of WT ZC3HC1 or certain mutants, and in which the ectopic ZC3HC1 is also bound to the NE, are marked with green arrows. A nucleus in which the nucleoplasmic TPR-sfGFP pool remained unchanged, as upon expression of mCherry-ZC3HC1 1–25_41–482_500–502, which cannot bind to the NE, is marked with an arrow in magenta. Arrowheads in yellow mark nuclei in which the ZC3HC1 mutants bind to the NE but fail to recruit TPR, which remains nucleoplasmic instead. Bar, 10 μm. **(F)** Schematic of ZC3HC1. Grey boxes denote the two BLDs, with their zinc-finger motifs C-X(2)-C (green) and H-X(3)-C (blue) highlighted. The hatched box indicates the largely disordered loop, dispensable for TPR interactions. Schematics of α-helices above the boxes outline their positions and relative lengths. Brackets mark segments essential for ZBD1 binding (grey) and those additionally required for, or contributing to, ZBD2 binding (orange).

Similar Y2H results were obtained with the *S. cerevisiae* homologs: deletion of either the N- or C-terminal region flanking the NuBaID of Pml39p did not noticeably affect binding to the Pml39-binding domain 1 (PBD1) of Mlp1p but abolished binding to PBD2 of Mlp1p (Supplemental Figure S3B).

Next, to examine whether the NuBaID-flanking sequences are required for tethering TPR to the NB, we used CRISPR/Cas9n to edit the *ZC3HC1* gene in HeLa cells. While no cell line with an N-terminal deletion was obtained, consistent with earlier difficulties editing *ZC3HC1* at its start codon (e.g., Gunkel and Cordes, 2022), we generated homozygous lines expressing all ZC3HC1 with a C-terminal sfGFP tag, both in its wild-type form and in a mutant form, the latter lacking aa 487–502 (Figure 3B1; Supplemental Figure S3C). While the mutant (Δ487– 502) co-localised with TPR at the NE, quantification of TPR’s NE-associated amounts revealed a reduction, on average, to about 44% of WT levels (Figure 3B), indicating that ZC3HC1 1– 486 cannot stably tether additional TPR polypeptides to the NB.

Of note, this reduction of NE-associated TPR was not accompanied by an obvious reciprocal increase in nucleoplasmic TPR. This resembles our observations in various ZC3HC1 KO substrains, in which NPC-associated TPR is consistently reduced by about half relative to WT cells, while most of these substrains nevertheless do not display an evident nucleoplasmic TPR pool (Gunkel *et al*., 2021; Gunkel and Cordes, 2022). Only a few ZC3HC1 KO substrains derived from the same progenitor line maintain a pronounced nucleoplasmic TPR pool, for reasons that remain unresolved, and these substrains can be used for TPR re-recruitment experiments (Gunkel and Cordes, 2022), as will be also shown below.

Furthermore, we compared the NE-associated signal intensities of sfGFP-tagged ZC3HC1 Δ487–502 with sfGFP-tagged full-length ZC3HC1 in co-cultured cells. The two lines were easily distinguishable because intact ZC3HC1 appeared located almost exclusively at the NEs, whereas the mutant in this particular cell line was also present throughout the nuclear interior (Figure 3C1). However, following image-based quantification of the NE-associated GFP signal intensities in live cells, we found ZC3HC1 Δ487–502 only reduced by ∼20% relative to WT ZC3HC1. Only after TX-100 extraction did the NE-associated amount of ZC3HC1 Δ487–502 drop, on average, to about 42% of WT levels (Figure 3C).

We next asked whether removal of the CTD of ZC3HC1 would affect the formation of cytoplasmic and nuclear TPR foci, since WT ZC3HC1 interconnects TPR polypeptides within such foci, as they disappear upon induced degradation of ZC3HC1 (Gunkel and Cordes, 2022). Such foci arise upon RNAi-mediated depletion of NUP153 (e.g., Hase and Cordes, 2003; Gunkel and Cordes, 2022), which is an NR protein binding to the Y-complex (e.g., Vasu *et al*., 2001; Vollmer *et al*., 2015), and even more so upon RNAi of NUP107, which is an actual component of the NR’s Y-complex (e.g., Belgareh *et al*., 2001; Harel *et al*., 2003; von Appen *et al*., 2015) and contributes to TPR docking sites at the NPC (e.g., Kim *et al*., 2014; Gunkel *et al*., 2021; Gunkel and Cordes, 2022).

Here, we performed NUP107 RNAi in the cell lines expressing ZC3HC1-sfGFP or ZC3HC1-sfGFP Δ487–502. In both cell lines, knockdown of NUP107 resulted in essentially complete detachment of TPR and both ZC3HC1 versions from the NPC. However, while numerous TPR- and ZC3HC1-containing foci were detectable in the presence of ZC3HC1- sfGFP, they were absent in the cells with ZC3HC1-sfGFP Δ487–502, indicating that ZC3HC1 cannot stably interconnect TPR polypeptides without its C-terminal segment (Figure 3D).

We next examined to which extent each NuBaID-flanking region contributes to re- appending soluble nucleoplasmic TPR back to the NB. For this purpose, we used a previously characterised HeLa ZC3HC1 KO cell line in which a substantial amount of those TPR polypeptides that can no longer be appended to the NB remain soluble in the nucleus (Gunkel and Cordes, 2022). Induced expression of WT ZC3HC1 in these cells had quantitatively restored NB attachment of essentially all soluble TPR within 30–60 min, whereas a NuBaID mutant (C429S) had not (Gunkel and Cordes, 2022). Here now, we accordingly studied mCherry-tagged ZC3HC1 mutants lacking either the entire N- or C-terminal region or parts thereof. These NuBaID-flanking regions are predicted to be mostly unstructured, except for two α-helices (aa 26–40 and aa 483–499; Gunkel *et al*., 2023), which we deleted internally in some mutants, to preserve the protein’s native termini and avoid potentially destabilising new ends (e.g., Varshavsky, 2019).

The outcome can be summarised as follows: All N-terminal mutants with an intact NuBaID (e.g., aa 72–502 and 1–25_41–502) re-appended TPR to the NB quantitatively, revealing that the entire N-terminal region is dispensable for TPR binding in human cells, unlike in the Y2H system (Figure 3E; Supplemental Figure S3D). By contrast, a NuBaID-intact mutant lacking both NuBaID-flanking regions (aa 72–472) failed to recruit TPR, as did a mutant that only lacked the entire C-terminal region (aa 1–472). Similarly, 1–486-mCherry failed to recruit TPR, while traces of recruitment were sometimes observed with mCherry-1–486, indicating that a C-terminal tag on 1–486 exacerbates the ZBD2-binding defect, possibly via steric hindrance. Finally, internal deletion of the C-terminal α-helix (aa 1–482_500–502) often abolished recruitment, and when both α-helices were deleted (aa 1–25_41–482_500–502), TPR recruitment was never detected (Figure 3E).

Together with previous results (Gunkel *et al*., 2023), these data indicated that docking of ZC3HC1 to NPC-anchored TPR requires only an intact NuBaID, whereas quantitative recruitment of additional TPR polypeptides via ZBD2 depends on NuBaID integrity plus the C-terminal flanking region, in particular its α-helix. The N-terminal α-helix, for its part, appears to provide further stabilisation of the ZC3HC1:ZBD2 interaction (Figure 3F).

#### The mean copy numbers of TPR and ZC3HC1 per NPC are numerically compatible with a widespread occurrence of NB-appended ZC3HC1:TPR ensembles in different cell types

Next, to assess which arrangements between both proteins are numerically feasible at the NPC, we set out to quantify the NB-associated copy numbers of ZC3HC1 and TPR. Earlier work, comparing WT and ZC3HC1 KO cell lines, had shown that ZC3HC1 is required for appending about half of the NE-associated TPR to the NBs of several human tumour and immortalised non-tumour cell lines (Gunkel *et al*., 2021; Gunkel and Cordes, 2022), and this likewise applied to IMR-90 fibroblasts with finite life span. In fact, quantification of TPR’s NE-associated amounts in IMR-90 ZC3HC1 KO cells revealed a reduction, on average, down to ∼43% of WT levels (Figure 4A), while ZC3HC1 RNAi in IMR-90 cells reduced the mean amounts of NE- associated TPR to about 60–61% (Supplemental Figure S4A).

**Figure 4.**
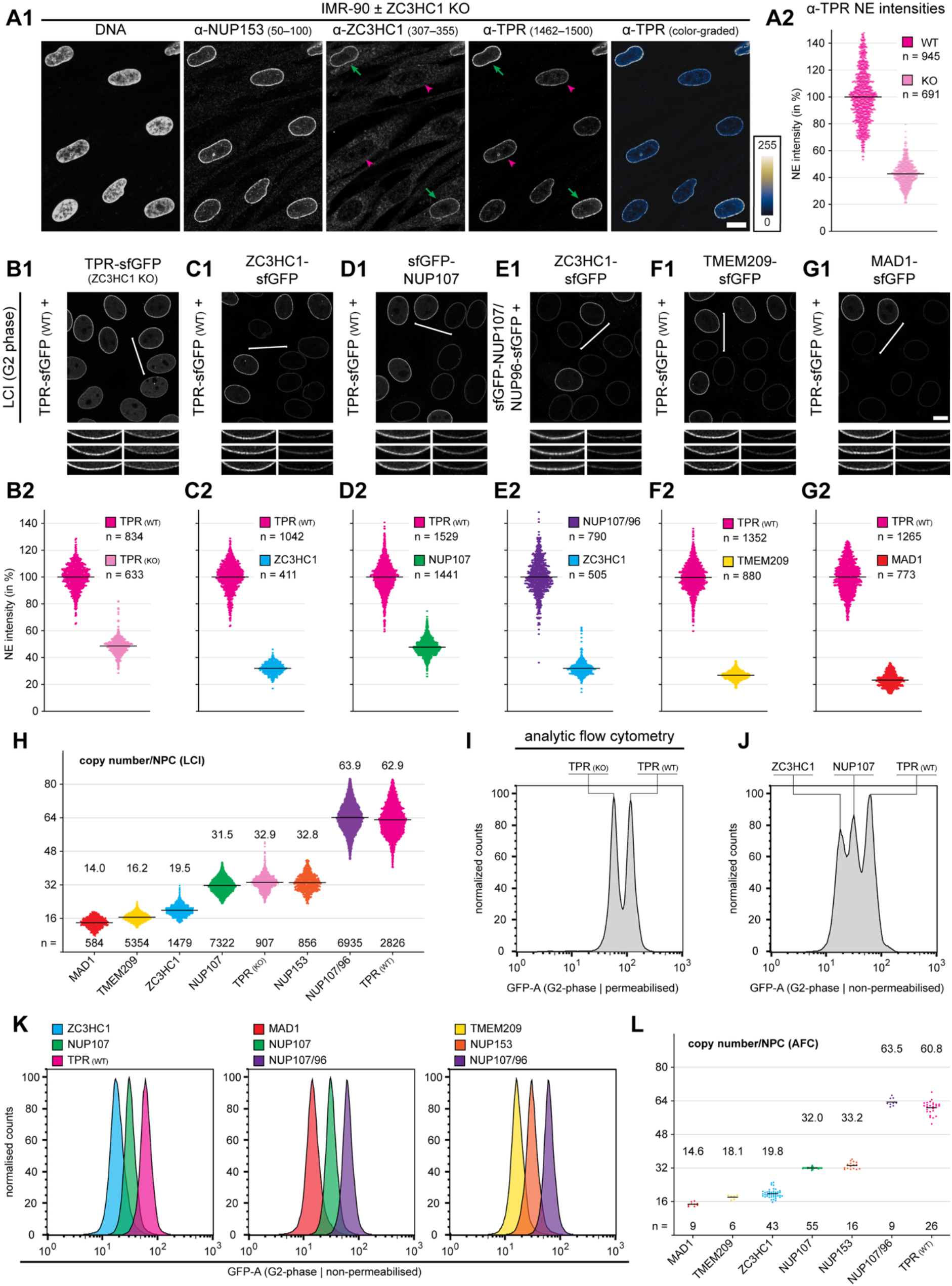
The mean TPR copy number per NPC in HeLa exceeds not only that of ZC3HC1 but also that of NUP107, revealing unexpected TPR abundance. **(A)** Triple-IFM of co- cultured IMR-90 WT and ZC3HC1 KO cells, and quantification of NE-associated IF-signal yields for TPR. (A1) Cells were fixed and immunolabelled for TPR, ZC3HC1, and NUP153, the latter as reference. The micrograph for TPR is also shown colour-graded next to an LUT. The nuclei of two WT cells are marked by green arrows, while arrowheads in magenta mark the nuclei of two representative ZC3HC1 KO cells with NE-associated TPR markedly reduced. Bar, 10 µm. (A2) Automated image-based quantification of NE-associated TPR immunofluorescence in IMR-90 WT and ZC3HC1 KO cells. **(B-G)** LCI of co-cultured, G2- synchronised populations of pairs of different HeLa cell lines (B1-G1), including representative NE segments from each pair at higher magnification, and automated image- based quantification of the NE-associated sfGFP signal intensities for each pair (B2-G2). In B1-G1, brackets mark exemplary pairs of the two different cell lines. Bars in B1-G1 (overview images), 10 µm. In B, WT and ZC3HC1 KO cells expressing TPR-sfGFP were compared, whereas in C–G, only WT cells were compared, each expressing different sfGFP-tagged proteins, including TPR-sfGFP with either ZC3HC1-sfGFP (C), sfGFP-NUP107 (D), or TMEM209-sfGFP (F), or MAD1-sfGFP (G), and ZC3HC1-sfGFP with sfGFP-NUP107 plus NUP96-sfGFP (E). **(H)** Summary scatter dot plot, compiled from numerous image-based quantifications, as exemplified in D–G and including corresponding analyses for NUP153 and MAD1 shown in Supplemental Figure S4M5 and S4N5. Relative NE-associated fluorescence ratios were converted into absolute copy numbers based on the respective copy-number references. The quantifications shown in this plot were restricted to datasets in which cell lines expressing GFP-tagged TPR, ZC3HC1, NUP153, or MAD1 had been co-cultured with one of the three copy-number reference cell lines expressing either sfGFP-NUP107 plus NUP96- sfGFP, sfGFP-NUP107, or TMEM209-sfGFP. **(I–L)** Quantification by AFC. Single-parameter histograms (I-K) display the relative fluorescence of ∼50,000 analysed HeLa cells per line, with the respective peak values set to 100%. Fluorescence intensities shown represent the total integral sums of the signal pulses produced by the cells passing through the laser beam (GFP- A). Cells were either analysed as co-cultured populations of two (I) or three cell lines (J), or as separate cultures analysed in parallel with identical AFC settings (K). To remove the soluble nucleoplasmic pool of sfGFP-TPR from ZC3HC1 KO cells, while retaining NB-associated TPR, the co-cultured and G2-synchronised WT and ZC3HC1 KO cells were briefly detergent- permeabilised before AFC (I; see also Supplemental Figure S4E3), unlike when the G2- synchronised WT cells expressing different sfGFP-tagged proteins were analysed as intact cells (J and K). Similar distribution plots obtained from G1 cells and from co-cultured cell populations are shown in Supplemental Figure S4E3 and S4P. The x-coordinate of the residual, NE-associated fluorescence peak in the permeabilised ZC3HC1 KO cells amounted to only about half the value of that of equally permeabilised WT cells (I). Relative to the x-coordinate of the sfGFP-NUP107 fluorescence peak in intact cells in G2, the values obtained for TMEM209-sfGFP, ZC3HC1-sfGFP, TPR- sfGFP, and sfGFP-NUP107/NUP96-sfGFP were 52.0%, 60.2%, 192.1% and 196.7%, respectively. Relative to sfGFP-NUP107/NUP96-sfGFP, the values for TMEM209-sfGFP and sfGFP-NUP153 were 26.4% and 49.8%, respectively. **(L)** Summary scatter dot plot based on the AFC data of multiple independent experimental repeats. Relative fluorescence peak positions obtained from AFC analyses of intact cells were converted into absolute copy numbers based on the respective copy-number references.

However, actual copy numbers per NPC were unavailable for ZC3HC1, and reported numbers for TPR in human cells varied widely, ranging from eight copies in DLD-1 cells (Li *et al*., 2021) to 16 (Otsuka *et al*., 2021) or ∼32 copies per NPC in the HeLa Kyoto line (Ori *et al*., 2013). Provided that the NB may indeed be a structure of eight-fold rotational symmetry, some of the reported copy numbers for TPR appeared irreconcilable with other findings, namely that a TPR homodimer represents the smallest NB assembly subunit (Hase *et al*., 2001), and that about half of all NPC-associated TPR in different cell lines requires ZC3HC1 for such a localisation (Gunkel *et al*., 2021; Gunkel and Cordes, 2022).

To address this issue, we started with determining the mean copy numbers for NPC/NB- associated TPR and ZC3HC1 in HeLa P2 cells. To this end we employed lines in which either all TPR or all ZC3HC1 were sfGFP-tagged (Gunkel and Cordes, 2022; Supplemental Figures S2D and S3C) and, as a reference, a HeLa P2 line in which all NUP107 carried an N-terminally appended sfGFP tag (Supplemental Figure S4B). These cells were to be cell-cycle- synchronised and analysed by LCI and analytical flow cytometry (AFC) to quantify the sfGFP signal intensities at the NEs and in whole cells. One would then determine the copy numbers for TPR and ZC3HC1 per NPC by comparing their signal intensities with those for NUP107, whose copy number is firmly established at 32 per NPC in HeLa Kyoto cells (Bui *et al*., 2013; Ori *et al*., 2013; von Appen *et al*., 2015; Politi *et al*., 2018).

We regarded determining the copy numbers of NPC-associated TPR and ZC3HC1 by LCI and AFC in HeLa P2 cells, using NUP107 as the main reference, as justified and diagnostically conclusive for the following reasons. First, in interphase HeLa P2 cells, NE-associated NUP107, TPR and ZC3HC1 reside almost exclusively at the NPCs, with TPR and ZC3HC1 appended to essentially all NPCs, and neither protein being part of any NE-aligned structures located between the NPCs (Gunkel *et al*., 2021, and see further below). In addition, TPR and ZC3HC1 remain associated with the NPC/NBs and are not degraded under any of the buffer and moderate temperature conditions that also apply to LCI or AFC (Gunkel *et al*., 2021; and our unpublished data; Gunkel and Cordes, 2022).

Second, HeLa P2 contains essentially no soluble NUP107 and only minor amounts of soluble TPR and ZC3HC1 (Gunkel *et al*., 2021). In addition, cytoplasmic annulate lamellae (AL) are rare in HeLa P2, with small AL only sporadically observed (e.g., Supplemental Figure S4B5). We therefore considered the amounts of NUP107 associated with AL pore complexes (e.g., Walther *et al*., 2003; Lin and Sumara, 2024) negligible in HeLa P2, while TPR and ZC3HC1 are absent from AL anyway (Cordes *et al*., 1997; Gunkel *et al*., 2021). Furthermore, unlike some other HeLa sublines (e.g., Griffis *et al*., 2002), HeLa P2 also lacks nuclear NUP98 bodies that recruit NUP107 (Morchoisne-Bolhy *et al*., 2015). Thus, when analysing cell-cycle- synchronised HeLa P2 cells harvested in G1 and G2, we assumed that the GFP signals even in intact cells would almost entirely originate from NPC- and NB-associated proteins when analysed by flow cytometry.

Third, we knew that the non-canonical TPR-L isoform would not interfere with our TPR copy-number determinations, even though TPR-L, expressed in different cell lines, is also detectable in small but significant amounts in HeLa P2 cells (Supplemental Figure S4C). Since TPR-L and canonical TPR appear to associate similarly well with NPCs and NBs, we initially considered it possible that the presence of untagged TPR-L may confound copy-number determinations based on N-terminally tagged canonical TPR. However, we found this concern not to be an issue because sfGFP tagging of TPR at the canonical start codon abolished detectable expression of TPR-L (Supplemental Figure S4D1). On the other hand, when TPR was sfGFP-tagged at its CT, this resulted, as expected, in co-cultured populations of TPR- sfGFP and TPR-L-sfGFP polypeptides. However, the ratio between TPR-sfGFP and TPR-L- sfGFP at the NPCs remained similar to that between untagged TPR and TPR-L (Supplemental Figure S4D2). Consequently, the later determined copy numbers for N-terminally tagged sfGFP-TPR were highly similar to those obtained for C-terminally tagged TPR (see further below).

Fourth, NPC density in HeLa P2 cells, and thus the amounts of NUP107, were known to remain essentially unchanged upon ZC3HC1 KO, as compared to the WT cells (Gunkel *et al*., 2021). This knowledge had allowed us to attribute changes in NPC-associated TPR copy numbers in ZC3HC1 KO cells exclusively to compositional changes at the NB, rather than in part to a reduction in NB numbers. In addition, we already knew, at least for TPR, that the relative amounts in WT and ZC3HC1 KO cells that one could determine by LCI and AFC corresponded to those relative amounts previously determined by IFM and immunoblotting (Gunkel *et al*., 2021). Specifically, using the already available HeLa P2 WT and ZC3HC1 KO cell lines expressing TPR C-terminally tagged with sfGFP (Gunkel and Cordes, 2022), quantification via LCI had already revealed that the total amount of NE-associated TPR-sfGFP was reduced by about half in the ZC3HC1 KO cells (Figure 4B; Supplemental Figure S4E1 and S4E2). Flow cytometry yielded very similar results after detergent extraction, which removed any residual soluble TPR that was no longer NE-associated in the ZC3HC1 KO cells (Figure 4I; Supplemental Figure S4E3). Together, these data underscored that quantification via sfGFP tags would yield TPR copy number relationships equivalent to those of the untagged proteins.

Fifth, prior studies had shown that potential non-steric optical effects, like GFP–GFP self- quenching at high local GFP concentrations, are negligible in live cell experiments (e.g., Wang *et al*., 2005; Gather and Yun, 2014). Accordingly, because GFP multimerisation does not produce detectable self-quenching, AFC-measured fluorescence had been shown to scale proportionally with cellular GFP content (Leonetti *et al*., 2016). Nonetheless, we also confirmed the linear detection range of our AFC settings using fluorescent calibration beads that carried different amounts of coupled fluorophore and found that the fluorescence intensities of sfGFP-NUP107 cells, as well as those of two additional HeLa P2 reference cell lines described further below, all fell within the linear range defined by these beads. Furthermore, the relative fluorescence intensities of these reference cell lines also scaled proportionally in LCI, revealing that they fell within the linear detection range of our microscope settings (see further below).

Sixth, several other prerequisites for rendering our approaches diagnostically conclusive had been fulfilled: all HeLa P2 cell lines had undergone extensive quality control, which confirmed that (i) all alleles of the *NUP107*, *TPR*, *ZC3HC1* and additional NUP and NB genes had been correctly tagged, (ii) the sfGFP-expressing cells were proliferating as well as their tag-free equivalents, (iii) NPC densities were indistinguishable between non-tagged cells and those expressing the sfGFP-tagged proteins, and (iv) immuno-labelling intensities at the NEs of cells expressing either tagged or tag-free proteins were likewise indistinguishable, indicating that the tagging did not considerably affect the number of copies incorporated into the NPC or NB (Supplemental Figure S4B4 and further below).

Finally, along with very similar results in other cell lines (Gunkel *et al*., 2021), we also knew that the relative amounts of NE-associated NUP107, ZC3HC1, and TPR in HeLa P2 are essentially indistinguishable from those in HeLa Kyoto (Supplemental Figure S4F1 and S4F2), the latter being a more commonly used HeLa subline in which copy numbers of NPC proteins had previously been determined (e.g., Ori *et al*., 2013; Otsuka *et al*., 2023). Furthermore, we had generated a HeLa Kyoto ZC3HC1 KO cell line and found, just as in HeLa P2 ZC3HC1 KO cells, that the amount of NE-associated TPR was reduced by about half compared to the WT, while NPC densities at the NEs of WT and ZC3HC1 KO cells remained unaffected (Supplemental Figure S4F3–S4F6). Taken together, these findings indicated that the absolute copy numbers we aimed to determine for TPR and ZC3HC1 in HeLa P2 could also be applied to HeLa Kyoto, thereby allowing direct comparison with previously reported data. The absolute copy numbers determined in HeLa P2 would furthermore allow us to relate these values to the relative amounts of TPR and ZC3HC1 previously determined for several other human cell types, in parallel to those for HeLa P2 (Gunkel *et al*., 2021).

As an aside, we regard it as noteworthy that the modal chromosome number and the range of chromosome numbers we had determined for HeLa P2 (Gunkel *et al*., 2021) are highly similar to those for HeLa Kyoto (Landry *et al*., 2013). Since HeLa P2 and HeLa Kyoto are neither progenitor nor progeny of each other, we consider it remarkable that independent screening projects yielded sublines with very similar, relatively narrow aneuploid karyotypes that, albeit unbalanced, exhibit stability over many generations (Gunkel *et al*., 2021). This degree of karyotypic convergence is consistent with both sublines having undergone similar processes of karyotypic selection and stabilisation from the much more heterogeneous HeLa progenitor population.

Having established the methodological prerequisites and validated our approach in HeLa P2 cells, we started with determining the copy number relationships between the NB and NPC components using LCI-based quantification of NE signal intensities. First, the comparison of co-cultured populations of WT cells with either TPR-sfGFP or ZC3HC1-sfGFP yielded a mean ratio of ∼3.1:1 (Figure 4C; Supplemental Figure S4G1). Next, we compared co-cultured populations of cells with TPR-sfGFP and sfGFP-NUP107 (Figure 4D; Supplemental Figure S4G2), or with sfGFP-NUP107 and ZC3HC1-sfGFP (Supplemental Figure S4G3). While some sfGFP-NUP107-expressing cells could occasionally also be identified by the presence of a few small GFP-positive AL, the sfGFP-NUP107 cells were, in any case, easily distinguishable from the other two lines by their markedly different NE fluorescence levels. Quantifications yielded mean fluorescence intensity ratios of ∼1.9–2.1:1 between TPR-sfGFP and sfGFP-NUP107 in both G1 and G2 (Figure 4D; Supplemental Figure S4G2), and a mean ratio of ∼1.6:1 between sfGFP-NUP107 and ZC3HC1-sfGFP (Supplemental Figure S4G3). Furthermore, highly similar ratios were also determined when using the HeLa P2 cell line expressing sfGFP-TPR instead of TPR-sfGFP, indicating that the copy numbers of NPC-associated TPR polypeptides were essentially unaffected by whether TPR was tagged at its N- or C-terminus (Supplemental Figure S4H).

In addition, we compared co-cultured cell populations comprising either TPR-sfGFP- or ZC3HC1-sfGFP-expressing cells together with additional reference cell lines that we had generated by CRISPR/Cas9-editing. Among these was a HeLa P2 cell line expressing all NUP107 and NUP96 as GFP-tagged polypeptides (Supplemental Figure S4I), corresponding to 64 GFPs per NPC. As a further reference with 16 GFP-tagged polypeptides per NPC, we had also created a HeLa P2 line homozygously expressing sfGFP-tagged TMEM209 (Supplemental Figure S4J). TMEM209 is a transmembrane NPC protein in metazoans (Fujitomo *et al*., 2012; Tang *et al*., 2020; Fang *et al*., 2024; Kohlhause *et al*., 2026) that had been proposed (Field *et al*., 2014) to be the homolog of yeast NPC protein Pom34p (Miao *et al*., 2006), the latter occurring in 16 copies per NPC (Kim *et al*., 2018; Akey *et al*., 2023). In isolated NEs of *Xenopus* oocytes analysed by mass spectrometry (Gunkel *et al*., 2021), we had found *Xl*TMEM209 to be a similarly abundant protein that remained stably NPC-associated, i.e. in a Triton X-100-resistant manner, and in HeLa P2 cells, we had eventually found TMEM209 to occur at ∼16 copies per NPC as well (see further below). In co-cultured populations of cells expressing either sfGFP-NUP107/NUP96-sfGFP, sfGFP-NUP107, or TMEM209-sfGFP, all three cell lines could thus be easily distinguished from one another.

Their NE-associated fluorescence intensities showed an approximately 1:2:4 relationship, consistent with the ∼16:32:64 copy-number relationship of TMEM209:NUP107:NUP107/NUP96 (Supplemental Figure S4K).

When we then analysed co-cultured populations of cells expressing either sfGFP- NUP107/NUP96-sfGFP or ZC3HC1-sfGFP, these two cell lines could, as expected, be readily distinguished, again allowing quantification of their NE-associated sfGFP intensities, which yielded a mean fluorescence intensity ratio of ∼3.1:1 between NUP107/NUP96 and ZC3HC1 (Figure 4E). Similarly, in co-cultured populations of cells with TPR-sfGFP and TMEM209- sfGFP, these two cell lines too could be easily distinguished (Figure 4F1), and the LCI-based ratio between TPR-sfGFP and TMEM209-sfGFP was ∼3.8:1 (Figure 4F2). These data thus underscored the results obtained via comparison with sfGFP-NUP107. Furthermore, we correspondingly studied co-cultured populations of TPR-sfGFP-expressing ZC3HC1 KO cells together with WT cells expressing either sfGFP-NUP107/NUP96-sfGFP or TMEM209-sfGFP, which yielded LCI-based ratios of ∼2.0:1 and ∼0.5:1, respectively (Supplemental Figure S4L). Altogether, such LCI-based signal intensity rations provided a mean copy number of ∼20 per NPC for ZC3HC1 in WT cells. In ZC3HC1 KO cells, the mean copy number for TPR was ∼33 while being ∼63 in WT cells (Figure 4H). The latter value thus already indicated a higher total amount of TPR per NPC than previously reported.

In addition, we also used a HeLa P2 cell line in which we had CRISPR/Cas9-edited all *MAD1L1* alleles to encode C-terminally sfGFP-tagged MAD1 (Supplemental Figure S4M). MAD1 is a CC-forming protein (Jin *et al*., 1998; Sironi *et al*., 2002; Kim *et al*., 2012; Fischer *et al*., 2022; Chen *et al*., 2023) that resides at the NB proper and engages TPR’s T1 subpopulation (Campbell *et al*., 2001; Lee *et al*., 2008; Krull, 2011). Since MAD1 plays a central role in assessing the NB fibril architecture in *Xenopus* oocytes and HeLa P2 cells (as discussed further below), we also determined its NPC-associated copy number. Comparison with sfGFP-NUP107 with MAD1-sfGFP yielded a ratio of 2.3:1, corresponding to a mean of ∼14 NPC-associated MAD1 copies per NPC (Supplemental Figure S4M5). The nuclei of MAD1-sfGFP cells could thus also be easily distinguished from those of TPR-sfGFP cells in a co-cultured population of both cell lines (Figure 4G1), with a mean fluorescence intensity ratio of ∼4.3:1 between TPR-sfGFP and MAD1-sfGFP (Figure 4G2).

As an aside, we had also determined the NPC-associated copy numbers of sfGFP-NUP153, which also engages with TPR’s T1 subpopulation at the NB. Using a HeLa P2 line expressing all NUP153 N-terminally tagged with sfGFP (Supplemental Figure S4N), we found this TPR binding partner to occur at ∼33 copies per NPC (Figure 4H; Supplemental Figure S4N5).

Next, we analysed the different cell lines by AFC, after having verified that the chosen AFC settings provided a linear detection range spanning both the fluorescent calibration beads and the fluorescence intensities of the different reference cell lines (Supplemental Figure S4O).

First, we compared WT and ZC3HC1 KO cells expressing TPR-sfGFP, after having permeabilised these cells to remove soluble TPR from the ZC3HC1 KO cells. As for the subsequent comparisons with the other cell lines, we analysed both co-cultured populations comprising two or three cell lines as well as each cell line on its own (Figure 4I; Supplemental Figure S4E3). The comparison of the TPR-sfGFP signal intensities in WT and ZC3HC1 KO cells yielded mean ratios ranging from ∼1.8 to 1.9:1.

When subsequently analysing intact WT cells expressing sfGFP-tagged TPR, NUP107 or ZC3HC1, a ratio of ∼1.9:1 was determined between TPR-sfGFP and sfGFP-NUP107, while the ratio between sfGFP-NUP107 and ZC3HC1-sfGFP was ∼1.6:1, in both G2 and G1 (Figure 4J and 4K; Supplemental Figure S4P). Of note, even in co-cultured cell populations comprising all three cell lines these could be readily distinguished from one another, as reflected by three clearly separated fluorescence-intensity peaks (Figure 4J).

Furthermore, comparative AFC analyses using the other reference cell lines expressing sfGFP-NUP107/NUP96-sfGFP and TMEM209-sfGFP yielded mean signal intensity ratios of ∼3.5:1 between sfGFP-NUP107/NUP96-sfGFP and ZC3HC1-sfGFP and ∼3.3:1 between TPR- sfGFP and TMEM209-sfGFP (Figure 4K; Supplemental Figure S4P).

In toto, AFC analyses yielded copy number relationships between TPR, ZC3HC1 and the other proteins very similar to those determined by LCI. Thus, the AFC data, too, revealed that the NPC-anchored T1 subpopulation in HeLa P2 cells comprises 32 TPR monomers in the absence of ZC3HC1, that the total amount of all NPC-associated TPR subpopulations in WT cells corresponds to an average of ∼61 TPR copies per NPC, and that the NB-wide mean copy number of ZC3HC1 is ∼20 (Figure 4L). In addition, the AFC data yielded a mean copy number of ∼15 MAD1 and ∼33 NUP153 polypeptides per NPC, matching the LCI-based quantification data.

### The ZC3HC1-appended TPR polypeptides project into the nuclear interior beyond the NB’s prototypic dimensions and thereby extend the NB scaffold that surrounds the CTDs

Addressing the question of how these additional, ZC3HC1-appended amounts of TPR are positioned relative to the NB and its NPC-anchored T1 subpopulation, one could consider four scenarios: If a distal TR exists, or if the NB’s distal end is treated as a corresponding reference plane, the T2 polypeptides could either run perpendicular or parallel to this plane. In the perpendicular arrangements, the T2 polypeptides would project either towards the NPC (scenario 1) or into the nuclear interior (scenario 2). When instead arranged parallel to the TR, they could be either laterally arranged in a ring, then perhaps even constituting the TR itself (scenario 3), or laterally protruding from the NB’s distal end, then possibly interconnecting neighbouring NBs (scenario 4).

We sought to distinguish between these scenarios by stimulated emission depletion (STED) microscopy (Klar *et al*., 2000), using our CRISPR/Cas9-edited cell lines expressing sfGFP- tagged TPR. Because this approach would visualise only TPR’s termini rather than tracing entire fibrils, we initially reasoned that two prerequisites had to be met to obtain unambiguous positional information. First, the tagged terminus to be detected had to be sufficiently distant from TPR’s ZBD2 and constrained to that distance by structural rigidity. The length and rigidity of the CC between the NT and ZBD2 ensured this spatial separation, as homodimers comprising aa 1–398 form rectilinear rods of ∼51 nm (Hase et al. 2001), which would even suffice to bridge the ∼45–50 nm spacing between the NR and the TR of the NB proper (Krull *et al*., 2004; Gunkel *et al*., 2021). Therefore, we would first visualise the sfGFP-tagged N- termini of TPR in the WT and ZC3HC1 KO lines and in the mutant line TPR Δ1–58, using fluorophore-coupled nanobodies against GFP to enhance signal intensities. By contrast, we considered the CT less suitable for resolving the arrangement of the different TPR subpopulations relative to one another, because we regarded the mostly unstructured CTD as too flexible (Hase *et al*., 2001).

Second, while the N-termini of the T1 subpopulation, positioned ∼45–50 nm from the NR (Krull *et al*., 2004; Gunkel *et al*., 2021), would indicate the TR endpoint of rectilinear fibrils between NPC and TR, an additional, differently coloured landmark was required at the NPC. We chose the NR and visualised it by labelling NUP153 with antibodies specific for a segment between its inner nuclear membrane-inserting amphipathic helix and its NBD, via which NUP153 docks to the NR (Enarson *et al*., 1998; Vollmer *et al*., 2015). Localising these fixed points, i.e. the NBD of NUP153 and the N-termini of TPR’s T1 subpopulation, and treating them as the endpoints of the eight straight NR-TR axes of the NPC-anchored NB would then allow us to determine the positions of TPR’s T2 N-termini relative to them, and thereby distinguish between scenarios 1–3. Scenario 4, by contrast, did not require a landmark at the NR because single-colour STED microscopy had already sufficed to discard this option: inspection of the NBs in top-on view at the nuclear poles of WT and ZC3HC1 KO cells expressing sfGFP-TPR had shown that the GFP signals were confined to clearly outlined, roundish, mostly donut-shaped structures in both cell lines, and did not reveal prominent lateral protrusions from the NBs in WT cells (see further below).

To visualise the positions of TPR’s N-termini and NUP153 relative to each other along the NPC–NB axis, we used two-colour 2D-STED microscopy (e.g., Göttfert *et al*., 2013). While this approach would turn out to be sufficient to distinguish between scenarios 1–3 (see further below), it imposed several constraints that we had to consider. First, because we needed to inspect the NEs in an equatorial focus plane, the distance between the objective lens and the NPCs at the nuclear equator would be relatively large. This in turn meant that we could not expect to reach the degree of lateral resolution achievable when NPCs are closer to the objective, as is the case at one of the nuclear poles. Second, because conventional 2D-STED microscopy generally provides low spatial resolution along the optical axis (Z), signals from NPCs and NBs located above and below the equatorial focal plane would contribute to the measured intensity. Consequently, when interpreting line profiles across the NE, we needed to consider the mean NPC density per NE surface area and NE curvature (Supplemental Figure S5A). Together, these limitations meant that it would generally not be possible to resolve two TPR subpopulations at the nuclear equator when they are only 40–50 nm apart and emit fluorescence of the same wavelength.

Nevertheless, when WT and mutant specimens were imaged within the same session using identical STED settings, the sfGFP fluorescence band at the NE of WT cells reproducibly appeared broader by visual inspection than in the ZC3HC1 KO and TPR Δ1–58 cells (Figure 5A; Supplemental Figure S5B). This difference in band width was obvious in all three separate sessions in which we directly compared WT and mutant cells by STED microscopy.

**Figure 5.**
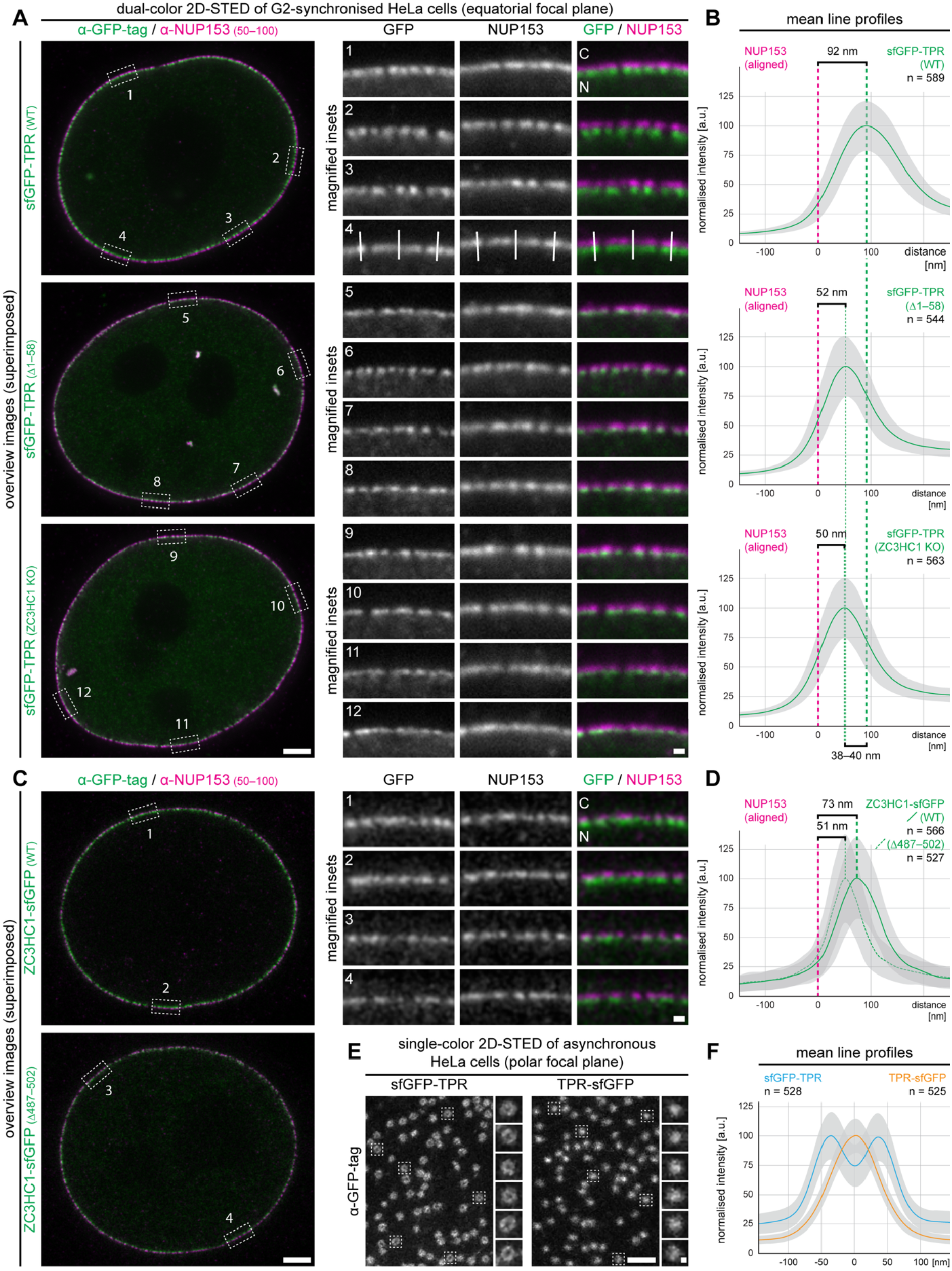
NB-associated TPR and ZC3HC1 in HeLa are components of a CTD-enclosing construction that extends into the nuclear interior. **(A)** STED microscopy of HeLa P2 cell lines expressing sfGFP-TPR (WT), sfGFP-TPR Δ1–58, and sfGFP-TPR in ZC3HC1 KO background. Cells were labelled with anti-GFP nanobodies and anti-NUP153 antibodies. Images of representative nuclei were acquired at the equatorial focal plane, revealing that the sfGFP signal distribution at the NEs of WT cells is notably broader than in the mutant lines that are no longer capable of recruiting additional TPR to the NB. Image acquisition settings were identical for all cell lines. Representative NE regions, marked as numbered rectangles in the overview images, are shown as magnified insets on the right. Lines in inset 4, drawn perpendicular to the NE, indicate examples for line profiles shown in B. Bars, 2 μm (overviews) and 200 nm (insets). Cytoplasmic (C) and nuclear (N) sides of the NEs are indicated. **(B)** Averaged line profiles for NE-associated sfGFP-TPR, based on >500 profiles across the NEs of five different nuclei per cell line. TPR profiles were aligned to the averaged signal intensity peaks of the corresponding NUP153 profiles. The resulting means for sfGFP relative to NUP153 are shown in green, with grey shading indicating the standard deviations. The mean sfGFP intensity peaks were set to 100%. Distances (brackets) between the peaks for NUP153 (dashed lines in magenta) and TPR (green dashed lines) are indicated. **(C)** STED microscopy of the HeLa P2 cell lines expressing ZC3HC1-sfGFP or ZC3HC1 Δ487–502-sfGFP. Bars, 2 μm (overviews) and 0.2 μm (insets). **(D)** Averaged line profiles for NE-associated ZC3HC1- sfGFP and ZC3HC1 Δ487–502-sfGFP, processed as in B and based on >500 NE profiles from >5 different nuclei per cell line. **(E)** Single colour STED microscopy of the WT HeLa P2 cell lines expressing sfGFP-TPR or TPR-sfGFP, labelled only with anti-GFP nanobodies. Images were acquired at the nuclear pole, revealing ring-like signal distributions for sfGFP-TPR and dot-like signal clusters for TPR-sfGFP. Bars, 500 nm (overviews) and 50 nm (insets). **(F)** Averaged line profiles for NE-associated sfGFP-TPR and TPR-sfGFP, based on >500 line profiles per cell line drawn across the centres of the signal accumulations in the top-on views of the NBs shown in E. The two mean profiles were aligned by matching the central trough of the sfGFP-TPR double peak to the single peak of TPR-sfGFP.

Nonetheless, we did not use these differences in band width or in absolute fluorescence intensities for quantitative interpretations, because in our STED setup the fluorescence intensities measured on different coverslips did not scale sufficiently reproducibly and proportionally with known copy-number differences. This finding was not wholly unexpected, because in STED microscopy the detected fluorescence is not determined by fluorophore number alone, but is additionally modulated by the depletion process, i.e. by how efficiently the STED beam suppresses fluorescence at the focal spot, which can differ between specimens even when microscope settings are kept constant.

We therefore used the two-colour data only to extract positional information, namely the distance between the NR landmark, provided by the NUP153 peak, and the sfGFP peaks for TPR and ZC3HC1 along line profiles across the equatorially focused NEs. To this end, we collected more than 500 line profiles per cell line from NE regions that were selected as being close to an ideal equatorial focus plane and that showed no NUP153–TPR boundary violations (Supplemental Figure S5A6–S5A8). Across these line-profile datasets, the sfGFP peak for intact TPR in WT cells was located markedly farther from the NPC than in ZC3HC1 KO and TPR Δ1–58 cells, with the latter two yielding essentially the same NUP153-to-sfGFP peak distance (Figure 5B). The larger NR-to-peak distance in WT cells ruled out scenario 1, which would predict a shorter NR-to-peak distance in WT cells, and scenario 3, which would predict very similar NR-to-peak distances in the WT and mutant cells. Instead, the larger NR-to-peak distance in WT cells supports scenario 2, in which NB-appended additional TPR polypeptides would project deeper into the nuclear interior.

In addition, we acquired STED images of NB-associated ZC3HC1-sfGFP and its mutant version ZC3HC1-sfGFP Δ487–502 and again noted that the sfGFP fluorescence band for the WT appeared notably broader than for the mutant (Figure 5C; Supplemental Figure S5C). The averaged line profiles revealed that the NR-to-peak distance for ZC3HC1-sfGFP was larger than for the ZC3HC1 mutant, yet smaller than the NR-to-peak distance for sfGFP-TPR in WT cells. By contrast, the NR-to-peak distance for ZC3HC1-sfGFP Δ487–502 was essentially identical to the NR-to-peak distances of TPR in both mutant lines (Figure 5B and 5D).

Finally, we also visualised the sfGFP-tagged C-termini of TPR in WT and ZC3HC1 KO cells. When focused on the NE equator, the fluorescence band in WT cells again appeared broader by visual inspection than in the KO cells, similar to what we observed for N-terminally tagged TPR (Supplemental Figure S5D). However, when visualising the NBs in top-on view, i.e. by focusing tangentially on the NE in the polar region of the nucleus closest to the objective, the C-terminal signals appeared as roundish dots, whereas the N-terminal signals mostly appeared as donut-shaped distributions (Figure 5E). Accordingly, XY line profiles drawn across the centres of these objects yielded a single central maximum for the CT signals and two lateral maxima for the NT signals, the latter separated by ∼90 nm (Figure 5F). Thus, these top- on views indicated that the N-termini of TPR surround its C-termini. Furthermore, inspection of top-on views of ZC3HC1-sfGFP likewise revealed a donut-like distribution, similar in appearance to that of sfGFP-TPR but smaller than that of sfGFP-NUP153 visualised for comparison (Supplemental Figure S5E).

Taken together, these STED data pointed to the following. First, the ZC3HC1-appended T2 polypeptides project toward the nuclear interior and thus extend beyond the dimensions commonly considered prototypic for the NPC-anchored NB. Second, because TPR’s N- terminal sfGFP signals surround its C-terminal signals in top-on views, we concluded that TPR’s CTDs are surrounded by the fibrillar scaffold of the NBs and their fibrillar appendices, in agreement with a previous model of the NB (Gunkel *et al*., 2021). Third, because the sfGFP fluorescence bands at the NEs of the mutant cell lines are expected to represent only those polypeptides that are part of the NPC-anchored NB proper, the notably broader band observed for WT ZC3HC1 is consistent with at least two spatially distinct ZC3HC1 subpopulations, of which the second is positioned farther away from the NB proper than the first. Finally, the reproducible finding that the NR-to-GFP peak distance for TPR’s N-termini in WT cells exceeds that of ZC3HC1 may indicate that the ZC3HC1-appended T2 population itself comprises at least two spatially distinct subpopulations, hereafter referred to as T2a and T2b, with T2b positioned even farther from the NPC than the second ZC3HC1 subpopulation.

Overall, although conventional STED microscopy could not directly resolve the individual subpopulations as separate layers, it revealed reproducible positional differences that point to a spatial ordering of TPR and ZC3HC1 subpopulations along the longitudinal NPC-NB axis. Together with the interaction hierarchy established above, these findings supported an NPC- associated serial arrangement comprising several TPR subpopulations and intervening ZC3HC1 subpopulations.

### The NPC-appended TPR:ZC3HC1 ensembles appear as stratified, often near-cylindrical structures stacked onto an NB

To visualize the three-dimensional (3D) organisation of the NPC-associated TPR:ZC3HC1 ensembles in greater detail, we next used recently introduced 4Pi-STORM (stochastic optical reconstruction microscopy; Bates *et al*., 2022). In 4Pi-STORM, fluorescence is collected through two opposing objectives and recombined in an interferometric detection path, sharpening the axial point-spread function and enabling single-molecule localisation with nanometer precision not only in the lateral (x/y) plane but also axially (z). As a result, this method can provide nearly isotropic 3D resolution.

We studied five of our HeLa P2 lines by 4Pi-STORM, comprising those expressing sfGFP- TPR, TPR-sfGFP, and ZC3HC1-sfGFP in the WT background, and sfGFP-TPR and TPR- sfGFP in the corresponding ZC3HC1 KO cells. Like 2D STED microscopy, 4Pi-STORM would not visualize the NB as a continuous object. However, TPR’s NTs and the NR would again outline the scaffold’s geometry.

To allow for a direct comparison with a previous study of the NPC based on 3D single- molecule localisation microscopy (Sabinina *et al*., 2021), we used protein ELYS as the NR landmark for STORM. ELYS is positioned at the outer perimeter of the NR through binding to the Y-complex (e.g., Rasala *et al*., 2006; Bilokapic and Schwartz, 2012; von Appen *et al*., 2015; Bley *et al*., 2022; Huang *et al*., 2022; Tai *et al*., 2022). For STORM, we detected ELYS with primary antibodies, while the GFP tags were labelled with anti-GFP nanobodies, all conjugated to photoswitchable dyes. Images of the labelled NR and NB components were then acquired by focusing tangentially on the nuclear pole facing the objective. This was expedient because 4Pi-STORM provides not only top-down images representing the x/y plane but, as pointed out above, also high-resolution information in the orthogonal x/z and y/z planes.

In the resulting x/z and y/z images, distinct layers of regularly arranged GFP localisations were apparent for sfGFP-TPR and ZC3HC1-sfGFP (Figure 6A). In the WT cells, often three “layers” of GFP localisations were visible for sfGFP-TPR and two layers for ZC3HC1-sfGFP, with seemingly rather similar inter-layer spacing for both proteins. In rare cases, sparse additional signals were even observed at positions suggestive of a partial fourth TPR layer (Figure 6A) and a partial third ZC3HC1 layer, while only one layer of sfGFP-TPR was visible in the ZC3HC1 KO cells (Figure 6A). The signals of the CTD-tagged TPR-sfGFP polypeptides, by contrast, did not appear regularly arranged but were rather distributed at random within an area in front of the NPCs, which in the WT cells was more extensive than in the ZC3HC1 KO cells (Figure 6A).

**Figure 6.**
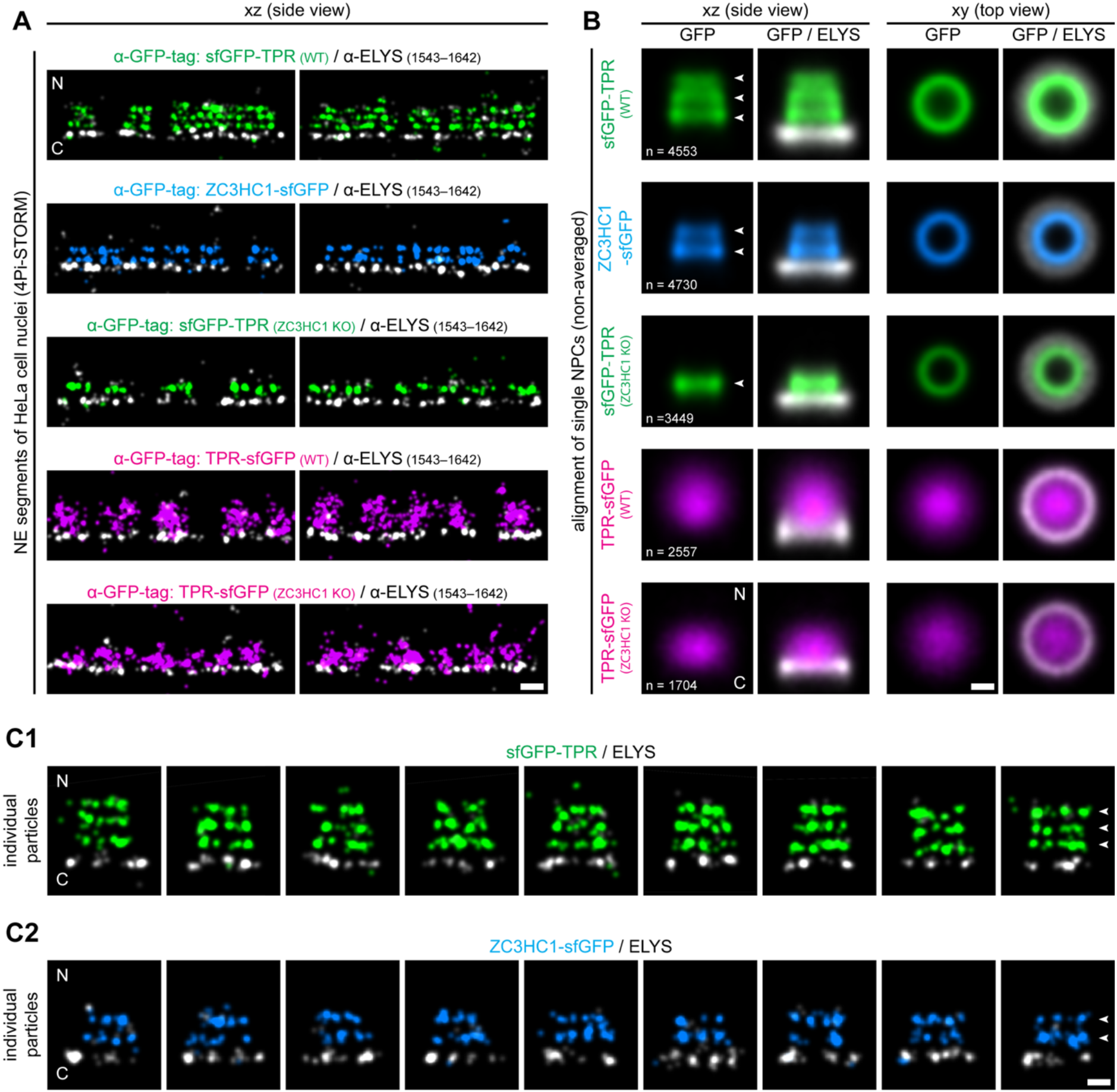
Visualization of stratified patterns of TPR and ZC3HC1 signals indicates CTD- enclosing scaffolds of cylindrical and truncated-cone-like appearance. **(A)** 4Pi-STORM of HeLa P2 cells expressing sfGFP-TPR, ZC3HC1-sfGFP, and TPR-sfGFP in the WT background, and sfGFP-TPR and TPR-sfGFP in ZC3HC1 KO cells. Images of representative NE regions are shown as x/z or y/z projections, revealing that the NE-adjacent signals for sfGFP-TPR and ZC3HC1-sfGFP appear arranged in stratified layers, whereas those for TPR- sfGFP appear more irregularly dispersed in front of the NPCs. Cytoplasmic (C) and nuclear (N) sides of the NEs are indicated. GFP fluorescence signals are shown in false colour, whereas immunolabelled ELYS is shown in white. Bar, 100 nm. **(B)** Composite views of COM-aligned stacks of NPC:NB- and NPC:NB:NBLS-containing particles. All automatically picked particles per cell line were pooled and computationally aligned and overlaid by COM registration in their original acquisition orientation. The x/z views reveal three layers of sfGFP- TPR and two layers of ZC3HC1-sfGFP in WT cells, only a single sfGFP-TPR layer in ZC3HC1 KO cells, marked by arrowheads, and a cloud-like distribution of TPR-sfGFP in both WT and ZC3HC1 KO cells. In x/y views, sfGFP-TPR and ZC3HC1-sfGFP appear as rings that overlap with the ELYS ring, while the TPR-sfGFP signals again appear as a cloud-like distribution around the NB’s longitudinal axis, surrounded by the sfGFP-TPR and ZC3HC1-sfGFP signals. Bar, 100 nm. **(C)** Selections of nine individual particles each, from the pools of automatically picked WT sfGFP-TPR- and ZC3HC1-sfGFP-containing particles, that exhibit a rather cylindrical appearance. Bar, 100 nm.

Next, to analyse the average appearance of the sfGFP distributions, we used an automated approach to identify and extract NE subvolumes containing individual NPCs and their NB:NBLS appendices, together referred to as “particles” in the following. Automated particle picking applied a cylindrical mask of 200 nm in diameter and 300 nm in height, under the constraint that no two cylinders overlapped. It was guided by annular ELYS signals, also accepting partial rings identifiable as ring segments. This approach commonly identified several hundred particles from the regions of interest (ROIs) surrounding the nuclear pole (Supplemental Figure S6A). Particle picking was performed on at least five nuclei per cell line. All particles of the same cell line were then pooled and computationally aligned by centre-of- mass (COM) registration in the original acquisition orientation, i.e. without correction of particle tilts relative to the z axis caused by the minor degree of NE curvature in the pole region, to generate an averaged representation (Figure 6B).

Even though this population average should not be read as the common appearance of all NBs, such stacking and alignment of many NB:NBLS entities corroborated the impression that ZC3HC1 and TPR’s N-termini are defining structural parts of the NB and its appendices. In the resulting x/z and y/z projections, sfGFP-TPR and ZC3HC1-sfGFP exhibited stratified arrangements, with three distinct layers for sfGFP-TPR and two for ZC3HC1-sfGFP in WT cells, and only a single sfGFP-TPR layer in ZC3HC1 KO cells. In the x/y projections, both sfGFP-TPR and ZC3HC1-sfGFP formed ring-like patterns that overlapped with the ELYS ring (Figure 6B). Displaying the GFP-tag and ELYS data separately indicated little crosstalk from ELYS into GFP, and virtually none from GFP into ELYS across cell lines (Figure 6B; Supplemental Figure S6B).

In the population averages for sfGFP-TPR and ZC3HC1-sfGFP in WT cells, signal intensities were highest in the NPC-proximal layer, corresponding to the NB proper, and progressively decreased in the more distal layers (Figure 6B). This suggests that the NPC- proximal layer contains, on average, more copies of each protein than any of the more distal layers. Although STORM signal intensities in individual particles are not generally proportional to protein copy numbers, stochastic variations in fluorophore blinking tend to average out across thousands of aligned particles, allowing more reliable estimates of average differences in protein abundance between the layers. Nevertheless, potential systematic differences in labelling efficiency between layers and limitations in particle alignment may still affect the layers’ relative signal intensities.

Unlike for sfGFP-TPR, the sfGFP signals of the C-terminally tagged TPR polypeptides appeared more evenly, i.e. cloud-like, dispersed along the NB’s longitudinal axis. The outline of this cloud in the z projection appeared spherical in the ZC3HC1 KO cells and ellipsoidal in the WT cells, in which it extended farther from the NPC than in the KO cells. In the x/y projections, the CTDs appeared encircled by a ring of ELYS signal (Figure 6B), and in neither cell type did the CTD distribution suggest participation in a regular structure.

While the population averages for sfGFP-TPR and ZC3HC1-sfGFP in WT cells exhibited a mean appearance reminiscent of a truncated cone (Figure 6B), the pool of automatically selected particles also included numerous examples with a more cylindrical appearance (Figure 6C). Although signal intensities often varied considerably between layers and, owing to the intrinsic variablity of STORM signals, could not be used to reliably infer relative copy numbers per layer in individual particles, these more cylindrical arrangements again frequently exhibited three layers of sfGFP-TPR and two layers of ZC3HC1-sfGFP.

Altogether, these 4Pi-STORM observations support the existence of stratified, scaffold- forming TPR:ZC3HC1 ensembles extending beyond the NB proper, with truncated cone-like or near-cylindrical appearances, while TPR’s CTDs exhibit a more diffuse distribution.

### Current evidence supports a model in which groove- and clamp-mediated interactions of ZC3HC1 with the segmentally folded rod domains of TPR homodimers generate serially arrayed, extended fibrillar scaffolds

Having shown that TPR:ZC3HC1 assemblies can comprise successive layers, we next used molecular structure predictions to visualise how TPR’s structural dimensions and the interaction domains of both proteins could give rise to such arrangements. Since the resulting scaffold would also constrain the spatial distribution of TPR’s CTDs, the predicted organisation and conserved features of this domain too were included in this analysis.

AlphaFold3 predicts TPR’s CTD for the greater part to be disordered (Figure 7A; Supplemental Figure S7A1), consistent with circular dichroism (CD) measurements of CTD polypeptides, which repeatedly indicated predominantly random coils, yet also a measurable proportion of β-sheet structure (Hase *et al*., 2001). The latter was likewise predicted by AlphaFold when several CTDs were analysed together (Supplemental Figure S7A2). AlphaFold additionally predicts a short alpha-helix within the CTD’s central part, comprising aa 2148–2154 (Supplemental Figure S7A1), as does PONDR (Predictor of Natural Disordered Regions; e.g., Garner *et al*., 1999; Supplemental Figure S7B). This region plays a role in TPR’s interaction with different types of RNA-binding proteins (our unpublished data to be published elsewhere) and is flanked by conspicuous phosphosites. In fact, the CTD is not only extensively phosphorylated during mitosis but also harbours multiple interphase phosphosites (Figure 7A; Supplemental Figure S7A1) that are regulated in response to extracellular stimuli (www.phosphosite.org), including sites linked to MAPK-dependent signalling (e.g., Eblen *et al*., 2003; Olsen *et al*., 2006; Vomastek *et al*., 2008; Pan *et al*., 2009; Rajanala *et al*., 2014). Furthermore, despite very low CTD sequence identity across phyla, other short motifs and compositional features are nonetheless conserved in metazoans and metaphyta (Kuznetsov *et al*., 2002). Among these features, a small C-terminal cluster rich in arginine and glycine residues and, in particular, a prominent acidic residue cluster in the central CTD region are particularly conspicuous (Figure 7A).

**Figure 7.**
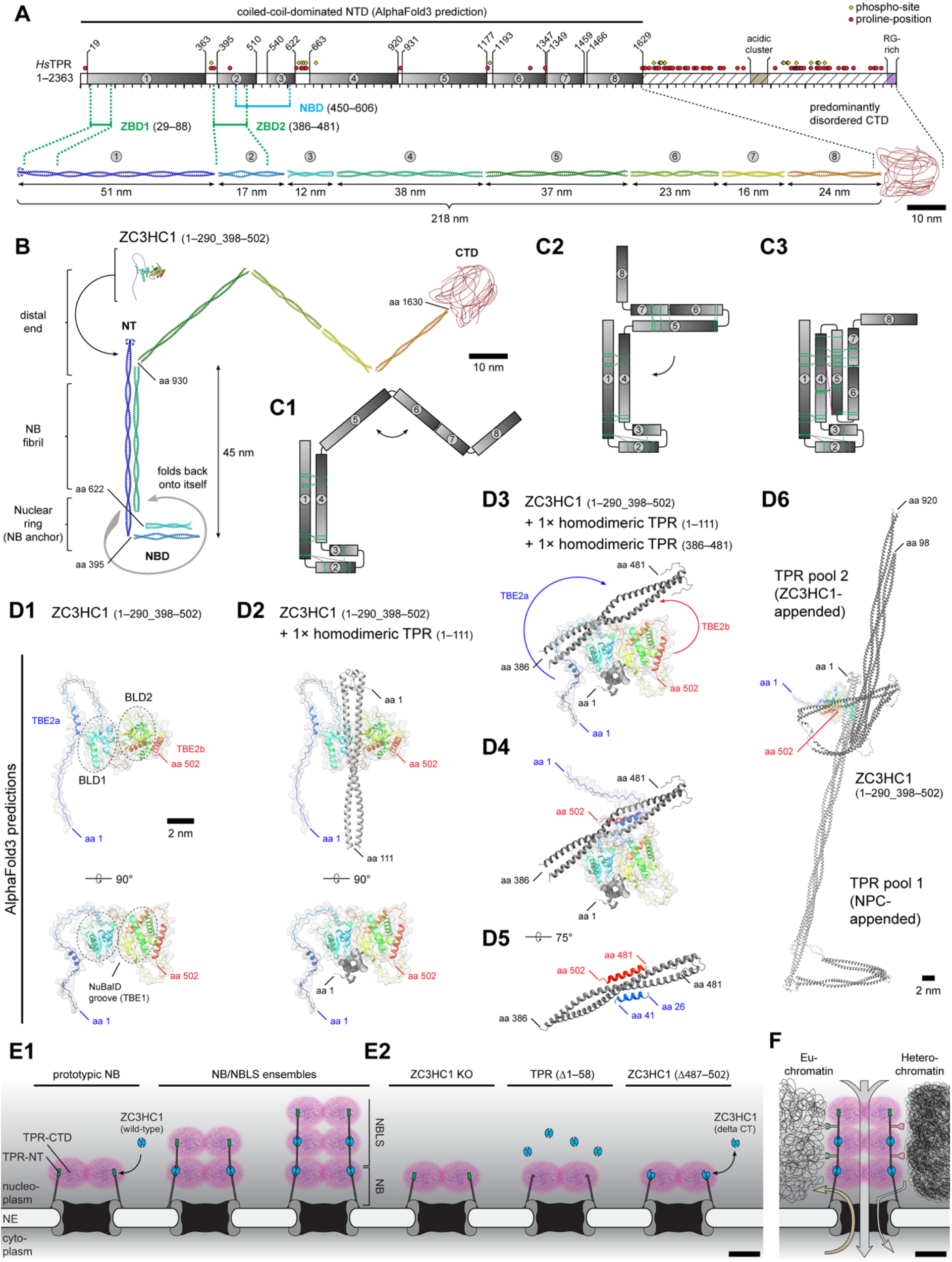
Towards the model of a stratified NB:NBLS scaffold of variable length, stepwise assembled by ZC3HC1-driven “groove-and-clamp” binding of folded TPR homodimers. **(A)** Schematic of TPR with AlphaFold3-predicted CC-forming regions (grey boxes). Corresponding AlphaFold3 models of TPR homodimers (rainbow colouring from blue NT to red CT) illustrate the eight CC segments (CC1–8) in the NTD and a mostly disordered CTD; hinge regions between segments are omitted. Approximate lengths are given for the NTD and each CC segment. Brackets and dashed lines mark the essential parts of ZBD1, ZBD2 and NBD. Proline residues and repeatedly detected phospho-Ser/Thr residues are indicated, as are an acidic residue cluster and an arginine/glycine-rich region within the CTD. **(B)** Model of a TPR homodimer of the T1 subpopulation in which CC4 folds back onto CC1 upon NR binding (Krull *et al*., 2004; Gunkel *et al*., 2021), yielding an antiparallel arrangement of the two CCs, thereby aligning four *a*-helices that form part of the rectilinear NB fibril. Additional folding of CC3 onto CC2 would contribute to rendering the ZBD2 inaccessible for ZC3HC1, while exposing the NBD for its interaction with the NR. CC segments are shown to scale relative to each other. Binding of ZC3HC1 to the ZBD1 of the TPR homodimer is indicated, with the ZC3HC1 structure also predicted by AlphaFold3. **(C)** Simplified schematics of the homodimeric NTD, with CC segments shown in three alternative conformations: (C1) back- folding of CC4 onto CC1 (as in B); (C2) additional back-folding of CC6/7 onto CC5; and (C3) further back-folding of CC5–7 onto CC1/4. Lengths of the CC segments are shown to scale relative to each other. Green lines indicate XL-MS cross-links, including CC1–4 links that support the initial hairpin-loop model. Additional cross-links suggest that the additional back- folding indicated in C2 and C3 may also occur. The single magenta line denotes the only cross- link incompatible with C3, because the linked residues in CC4 and CC5 would be too far apart. For clarity, cross-links between CTD residues are not shown. **(D)** AlphaFold3-predicted interfaces between ZC3HC1 and TPR homodimers at ZBD1 and ZBD2. For clarity, the large unstructured loop within BLD2 of ZC3HC1 is omitted. CC segments are shown in dark grey and the outline of ZC3HC1 in light grey, with selected regions additionally coloured. **(D1)** Structure prediction of ZC3HC1, with its two NuBaID-forming BLD modules (encircled) arranged such that the TBE1 groove is created between them. In addition, the *a*-helices of TBE2a and TBE2b within the otherwise unstructured regions flanking the NuBaID are highlighted in blue and red, respectively. **(D2)** Prediction of the ZBD1:ZC3HC1 interaction, illustrating that the TBE1 groove accommodates ZBD1 as part of the homodimeric CC1. **(D3- D5)** Prediction of the ZBD2:ZC3HC1 interaction, illustrating how the NuBaID-flanking *a*- helices of TBE2a and TBE2b change positions upon ZC3HC1 engagement with a second TPR homodimer (D3) and eventually clasp ZBD2 as part of the homodimeric CC2 (D4 and D5). **(D6)** In a composite created in Chimera, the TBE1 groove binds ZBD1 of one TPR homodimer of the T1 subpopulation (here only aa 1–920 are depicted), while TBE2a and TBE2b clasp ZBD2 of another homodimer belonging to the T2 subpopulation (here only aa 98–920 are depicted), thereby forming a fibrillar extension of the NB. **(E)** Model of the stepwise assembly of NB-like structures (NBLS) appended to the NPC-anchored NB. All fibrils of both NB and NBLS are here considered as consisting of folded TPR homodimers. **(E1)** Two NR-anchored TPR homodimers per asymmetric NB unit form two aligned bundles of altogether eight α- helices that together constitute each fibril of the NB proper in proliferating mammalian cells, with only one of these bundles here shown for simplicity. Once bound to the NB proper, a single ZC3HC1 can then recruit an additional TPR homodimer and append it to the NB. While it remains an open question whether the resulting NBLS is invariably composed of only eight instead of 16 TPR homodimers, the super-resolution microscopic data, interpreted in the context of the interaction hierarchy established here, indicate that the NB-appended T2 population of TPR can recruit additional ZC3HC1 polypeptides, which in turn recruit further TPR polypeptides. In principle, this alternating recruitment of ZC3HC1 and TPR may permit the stepwise addition of several NBLS layers until either ZC3HC1 or TPR becomes limiting. The resulting stratified scaffold would enclose a continuous CTD-containing interior (shown in magenta), which would contain an abundance of acidic residues due to the numerous CTDs’ acidic clusters. **(E2)** In the absence of ZC3HC1, or when TPR lacks its ZBD1 and thus cannot recruit ZC3HC1, or when ZC3HC1 lacks its C-terminal region required for ZBD2 binding, no NBLS can form, with TPR remaining confined to the fibrils of the NB proper. **(F)** Model of an NB with serially arrayed NBLS appendages. These NBLSs would extend the chromatin exclusion zone in front of the NPC, may provide additional sites of interaction with neighbouring chromatin and thereby play a role in local transcriptional regulation or in steps of RNP maturation, and may also contribute to defining different transport pathways (here depicted as arrows) to and from the NPC.

By contrast, AlphaFold predicts eight CC segments (CC1–CC8) for the two NTDs of a TPR homodimer. These CCs are arranged in parallel and in register (Figure 7A), in agreement with former results (Hase *et al*., 2001). The CCs are separated by short, flexible spacer sequences. Some of these spacers, too, contain phospho-serine or phospho-threonine residues, repeatedly detected in MS datasets (Figure 7A; e.g., Hornbeck *et al*., 2015; www.phosphosite.org) and possibly contributing to regulated interactions with other proteins and between the homodimeric parts of TPR. Furthermore, despite low overall sequence identity between TPR homologs from different phyla, several conserved sequence features also occur in the NTD, including distinct prolines in some of the spacers (Figure 7A; Kuznetsov *et al*., 2002; Gunkel *et al*., 2023) where they may promote CC disruption and flexibility.

The predicted length of CC1 is sufficient to span the 45–50 nm distance between TPR’s NT and the NR (Figure 7B), as determined by iTEM (Krull *et al*., 2004; Gunkel *et al*., 2021) and consistent with similar lengths of recombinant, homodimeric CC1 of TPR and Mlp1p (Hase *et al*., 2001; Stankunas and Köhler, 2024). AlphaFold further predicts that CC4 folds back onto CC1 (Figure 7B; Supplemental Figure S7C), consistent with iTEM evidence for backfolding of TPR and Mlp1p that positions the NT at the TR, and the NBD, which comprises parts of CC2 and CC3 (Figure 7A; Supplemental Figure S7C), at the NR (Krull *et al*., 2004; Niepel *et al*., 2013). Because AlphaFold generates this configuration with CC2 and CC3 slightly displaced from their more likely in vivo orientation, we repositioned these two CCs in Chimera via their flexible spacers to illustrate a more plausible geometry (Figure 7B). In this representation, CC2 and CC3, and thus ZBD2 and NBD, would lie ∼40–45 nm from the NT and ZBD1.

At this point, however, it remained uncertain how TPR’s other CCs would contribute to the NB scaffold. Previously, we had proposed that the CC5–8 segments of neighbouring NB fibrils might form a zig-zag-like arrangement that constitutes the TR (Gunkel *et al*., 2021), whereas CC1 and CC4 would fold onto each other and thereby form the rectilinear NB fibrils. This model provided an explanation for how the TRs seen in oocyte NBs could come about and it also harmonised with the oocyte’s NB fibril thickness of less than 6 nm, which could well accommodate two copies of such single-folded TPR homodimers, next to several other NB- associated proteins. This model had also considered immuno-localisations via epitope-specific antibodies by which we had located the distal parts of CC4, at ∼ aa 920, and of CC8, at ∼ aa 1630, in the TR region (Krull *et al*., 2004; Supplemental Figure S2A–S2C). Moreover, we had found a monoclonal TPR antibody, whose epitope we had mapped to aa 1462–1500 (Gunkel *et al*., 2021) and thus to the distal end of CC7, to label the TR region but not the NR in SR microscopy of ZC3HC1 KO cells (our unpublished data).

On the other hand, however, this model lacked experimental support regarding the positions of CC5 and CC6, since we have not yet succeeded in isolating an epitope-specific antibody that would have been suitable for immunolocalising the distal end of CC5 at ∼ aa 1180. Therefore, it remained unknown how CC5 and CC6, located between aa 920–1459, are actually positioned relative to the other CCs. Furthermore, our previous model in which CC5–8 form a zig-zag- like TR could not be reconciled with the recently characterised NBs in yeast and murine 3T3 cells, which had been found to lack a TR and whose tomographic electron density volumes for the NB fibrils were about 9–10 nm in diameter (Singh *et al*., 2024). Moreover, we had by then determined the inventory of NB-resident proteins in *Xenopus* oocytes and HeLa cells (our unpublished data), after having compared oocyte NEs with and without NBs (Gunkel *et al*., 2021) as well as lamina-NPC-NB fractions of auxin- and mock-treated HeLa TPR degron cells (Gunkel and Cordes, 2022) by mass spectrometry. However, when considering the copy numbers of all NB proteins and their arrangements relative to TPR at the NB fibrils, their total mass did not suffice to fill the tomographic electron density volumes of the 3T3 cells’ NB fibrils when only considering the CC1 and CC4 segments of two TPR homodimers as the scaffold-forming parts of the NB fibril.

We therefore wondered whether a TR may possibly be a speciality of NBs in cells without perinuclear chromatin, and whether TPR homodimers in other cell types, in which the NBs are surrounded by chromatin, might adopt a more highly folded and thereby more compact conformation, instead of forming a TR. In this context, we had found it remarkable that previous cross linking mass spectrometry (XL-MS) datasets for yeast NPCs had revealed not only intermolecular cross-links between Mlp1p and Mlp2p, but also numerous intramolecular cross-links which, in the case of Mlp1p, included not only those between CCs corresponding to TPR CC1 and CC4, but also many involving other Mlp1p CCs (Kim *et al*., 2018; Akey *et al*., 2022; Singh *et al*., 2024; Supplemental Figure S7D). This raised the question of whether these cross-links might indicate a multiply folded Mlp1p rod domain, and whether comparable cross-links may also have been detected for TPR. We therefore set out to data-mine the XL- MS datasets that were by then available from whole human cells or their isolated nuclei (Liu *et al*., 2017; Cardon *et al*., 2019; De Graaf *et al*., 2019; Bartolec *et al*., 2023; Bräuer *et al*., 2025).

While this inspection revealed hardly any cross-links between TPR and any of its genuine binding partners, and none with ZC3HC1, the datasets included numerous cross-links between TPR’s CC1 and CC4, indicating antiparallel alignment of these two CCs and thus supporting their backfolding as proposed in our previous models. The same held true for crosslinks between CC2 and CC3, which likewise indicated antiparallel alignment and backfolding of these two CCs (Figures 7C1; Supplemental Figure S7E1 and S7E2). More strikingly, though, numerous additional cross-links had been detected between CC5 and CC6/7 and between CC1/4 and CC5–7, suggestive of two additional intramolecular folding events involving the distal CC regions, namely between CC4 and CC5, CC5 and CC6/7, CC1 and CC6, and CC4 and CC6/7 (Figure 7C2 and 7C3; Supplemental Figure S7E3–S7E5). With the exception of a single cross-link between CC4 and CC5, detected only once in whole HEK293 cells (Bartolec *et al*., 2023; highlighted in magenta in Figure 7C3 and Supplemental Figure S7E4), all other cross-links were compatible with a plausible sequence of folding events leading to the formation of a bundle of four rectilinearly aligned α-helices per TPR homodimer (Figure 7C1– 7C3; Supplemental Figure S7E2–S7E5), and thus to a total of eight rectilinearly aligned α- helices per NB fibril.

On the other hand, the available cross-link data allowed some other hypothetical arrangements to be ruled out, at least for the immortal human cell lines that yielded these data, because the observed cross-links rendered such arrangements impossible. This included a parallel but staggered alignment of TPR homodimers, which could therefore finally be excluded as an explanation for how fibrillar NB extensions might be formed (Supplemental Figure S7E6), a scenario that had already been considered unlikely previously (Hase *et al*., 2001; Gunkel *et al*., 2021). Similarly, the available cross-linking data also excluded formation of a TR by a process in which the branched-off CC5–7 segments of neighbouring fibrils would align in a parallel but staggered manner (Supplemental Figure S7E7).

However, provided that some as yet unknown mechanism would prevent intra-homodimeric folding of CC5–7 onto CC1/4 in certain cell types, such as perhaps in full-grown amphibian oocytes, the cross-links between CC5 and CC6/7 would in principle allow antiparallel alignment of the branched-off CC5–7 segments of neighbouring fibrils, resulting in inter- homodimeric cross-links (Supplemental Figure S7E8). As proposed previously (Gunkel *et al*., 2021; see also Goldberg et al. 1996), such an alignment of TPR CCs would have to occur in a criss-crossed, interleaved manner, with the branched-off CC5–7 segments of each fibril aligning antiparallel with those of the next-but-one fibrils on either side. Since two TPR homodimers exist per NB fibril, this could allow formation of a TR of zig-zag-like appearance (Supplemental Figure S7E9).

By contrast, no cross-links have so far been detected that involve CC8. This CC may therefore be involved neither in formation of the NB fibril nor of the TR, and may instead act as a flexible jib for the CTD (Supplemental Figure S7E10).

Although we could not yet exclude that some of the cross-links derived from TPR conformations specifically occurring only for nuclear import, from TPR assembly intermediates, or from NBs that had partially collapsed during isolation, with the latter scenario potentially adding to the considered issue of false-positive cross-links in large-scale XL-MS (e.g., Bartolec *et al*., 2020), the reported TPR cross-links may instead indeed have captured genuine conformations at the NB. If so, two such multiply folded TPR homodimers, each corresponding to a compact bundle of four two-stranded coiled coils, could together form two laterally associated helical bundles, in principle sufficient to generate a fibril-like structure with an approximately ellipsoidal cross-section, with minor and major axes of about 4 and 8 nm, respectively. Such a structure would thus approach the dimensions of the tomographic electron- density volumes assigned to NB fibrils in 3T3 cells (Singh *et al*., 2024) and now more recently also in human cell types (Obarska-Kosinska *et al*., 2026).

We next returned to AlphaFold3 to inspect the interaction interfaces predicted between TPR and ZC3HC1 and to assess whether they were consistent with our experimental data. We had already concluded that ZBD1, comprising aa 29–88, represents a prototypic homodimeric CC, based on previous evidence that the smallest stable arrangement between TPR polypeptides in vivo is a soluble homodimer (Hase *et al*., 2001) and that aa 19–142 of TPR’s canonical isoform forms an essentially uninterrupted CC (Pal *et al*., 2017). AlphaFold3 confirmed the homodimeric nature of ZBD1 as part of CC1 and positioned ZBD1 in a groove formed by the two BLD modules constituting the NuBaID of ZC3HC1 (Figure 7D1 and 7D2; Gunkel *et al*., 2023). This groove will hereafter also be referred to as TPR-binding element 1 (TBE1) of ZC3HC1.

By contrast, ZBD2, as initially mapped to aa 386–481, would have included both a non-CC segment and the N-terminal part of CC2, because the next shorter construct we had tested, starting at aa 411, no longer interacted with ZC3HC1 (Figure 1C). However, AlphaFold3 predicts ZC3HC1 to interact only with a short segment of CC2, aa 415–444, which is clasped in a clamp-like manner by the two α-helices in the NuBaID-flanking extensions of ZC3HC1 (Figure 7D3–7D5). We now refer to these *a*-helices as the N-terminal TBE2a and the C- terminal TBE2b. The loss of ZBD2:TBE2a/TBE2b interaction upon truncating TPR to aa 411 suggests that the N-terminal part of CC2, starting at aa 395, is required for proper CC2 formation and thus for a functional ZBD2.

Next, we also asked how AlphaFold would handle two copies of ZC3HC1, the reason being that the BLDs of ZC3HC1 are closely related to the BIR (baculovirus inhibitor of apoptosis protein repeat) domains (e.g., Higashi *et al*., 2005; Kokoszynska *et al*., 2008; Gunkel *et al*., 2021, 2023), of which at least the BIR1 type is capable of homodimerisation (e.g., Lu *et al*., 2007; Mace *et al*., 2010; Hou *et al*., 2017; Yang *et al*., 2020). Although we found no evidence in our previous studies for stable homodimerisation of ZC3HC1 (Gunkel *et al*., 2021, 2023), and although most of our current data argue against ZC3HC1 homodimers being part of the TPR:ZC3HC1 arrangements once they are fully assembled, some of our data may nevertheless suggest that ZC3HC1 homodimers could play a transient role as an assembly intermediate (see Discussion). We therefore noted with interest that AlphaFold3 predicted a ZC3HC1 homodimer (Supplemental Figure S7F) which could embrace the ZBD1 of a single TPR homodimer and thereby involve the TBE1 of both ZC3HC1 polypeptides (Supplemental Figure S7G).

Furthermore, we also considered it noteworthy that the TBE1 appears promiscuous in silico, as AlphaFold can dock several TPR CC fragments or even other CC-forming proteins into it (Supplemental Figure S7H). Since such non-specific interactions are not observed in vivo (Gunkel *et al*., 2021), additional, yet unknown features must confer specificity for the TBE1:ZBD1 interaction. By contrast, the clamp-like TBE2a:ZBD2:TBE2b interactions appear specific, as AlphaFold generated this configuration only with ZBD2 but not with other TPR CCs or unrelated CC proteins tested.

Except for the uncertainties regarding ZC3HC1 homodimerisation and promiscuity of TBE1:TPR interactions, the AlphaFold3 predictions are largely consistent with the experimentally established interactions. They underscore our conclusion that a single ZC3HC1 polypeptide can link two TPR homodimers, by first binding via its TBE1 to the ZBD1 of one TPR homodimer and subsequently clasping the ZBD2 of another TPR homodimer with its NuBaID-flanking TBE2a and TBE2b (Figure 7D6). Of further note, AlphaFold predicts that Pml39p and Mlp1p engage in a similar manner. The two BLD modules of Pml39p form a groove, here called Mlp-binding element 1 (MBE1), which accommodates an Mlp1p homodimer via a ZBD1-homologous PBD1, while two α-helices in the NuBaID-flanking extensions of Pml39p, representing Mlp1p-binding element 2 (MBE2), would clasp another Mlp1p homodimer via its ZBD2-homologous domain PBD2 (Supplemental Figure S7I).

Together with the quantitative and spatial data, these observations support a stepwise NB– NBLS assembly model (Figure 7E1): ZC3HC1 first binds to one of the NB fibril’s two NPC- anchored TPR homodimers via TPR’s ZBD1, then recruits an additional TPR homodimer via ZBD2, followed by alternating recruitment of further TPR and ZC3HC1 polypeptides. By contrast, in the absence of ZC3HC1, or when TPR’s ZBD1 is missing, or when ZC3HC1 can bind ZBD1 but not ZBD2, such extended structures cannot form, leaving only the NPC- anchored T1 subpopulation of TPR (Figure 7E2).

However, in the WT cells, the NB-appended ZC3HC1:TPR ensembles spatially extend beyond the dimensions of the NB proper, and we propose that these ensembles thereby also expand the functional reach of NPC-associated TPR. In proliferating somatic cells, these fibrillar scaffolds may be involved, for example, in perinuclear chromatin organisation and local transcription regulation. Furthermore, by enclosing the CTDs of numerous TPR polypeptides, and thereby positioning numerous copies of the C-terminal RG-rich cluster, the phosphosite-flanked central α-helix, and the acidic cluster of the CTD within the scaffold interior, we additionally propose that these extensions also contribute to defining transport and cargo modification pathways to and from the NPC (Figure 7F).

## Discussion

### (1) A hierarchy of engagements between two ZBDs and three TBEs: attracting and being attracted

Previous work had shown that the initial attachment of ZC3HC1 to the NB and the subsequent recruitment of additional TPR by NB-bound ZC3HC1 can occur as temporally uncoupled binding events (Gunkel and Cordes, 2022). In the current study, we describe how these stepwise interactions involve two ZBDs of TPR and three TBEs of ZC3HC1. First, TBE1, predicted to be a groove formed by the two BLD modules of ZC3HC1, binds the homodimeric segment of CC1 that constitutes TPR’s ZBD1. Since NBs of ZC3HC1 KO cells can bind recombinant soluble ZC3HC1 in the absence of any soluble TPR (Gunkel and Cordes, 2022), we can now conclude that the ZBD1 of NB-associated TPR can act as the initial docking site for a soluble ZC3HC1 polypeptide without an attached TPR homodimer. In a second step, the NuBaID-flanking TBE2a and TBE2b, presumably acting together as a bipartite clamp, enable NB-bound ZC3HC1 to attract a soluble TPR homodimer by clasping the homodimeric CC2 segment that constitutes ZBD2. ZBD2 is thus the domain by which additional TPR polypeptides can be appended to an existing NB or NBLS.

Furthermore, since single amino acid substitutions within the NuBaID, likely disrupting the overall integrity of its bimodular architecture, abolish Y2H interactions not only with ZBD1 but also with ZBD2 on its own (Gunkel *et al*., 2023), one can conclude that structural integrity of the NuBaID is essential for enabling interactions with ZBD2. However, the ability of ZC3HC1 to bind T2 polypeptides via ZBD2 does not depend on prior binding to T1 polypeptides via ZBD1, as evidenced by Y2H interactions between ZC3HC1 and ZBD2- containing TPR segments that lack ZBD1.

Therefore, we cannot exclude that soluble ZC3HC1 in some in vivo situations, may already clasp the ZBD2 of a soluble TPR homodimer or accommodate its *Z*BD1 (see further below). However, such pre-assembled trimeric ZC3HC1:TPR subcomplexes would have to remain limited to only one of these two interaction modes. If formed via a ZBD1:TBE1 interaction in solution, such a subcomplex could in a next step be anchored to the NPC; if formed via a ZBD2:TBE2a/TBE2b interaction, it could be recruited to the free ZBD1 of an NPC-anchored TPR. However, simultaneous engagement of both ZBD:TBE pairs in solution is apparently prevented, since the formation of higher-order TPR:ZC3HC1:TPR ensembles is usually restricted to the NE.

In fact, while substantial amounts of soluble ZC3HC1 generated by ectopic expression can bind soluble TPR in interphase cell extracts, these ZC3HC1:TPR ensembles remain soluble after centrifugation at 250.000 × g (Gunkel *et al*., 2021). Furthermore, in *Xenopus* egg extracts that have been rendered competent for NPC/NB assembly, a certain proportion of the soluble ZC3HC1 and TPR also occurs as ZC3HC1:TPR subcomplexes, but these subcomplexes likewise remain soluble as long as the actual NPC/NB assembly process has not been initiated (Gunkel *et al*., 2021). Moreover, upon induced expression of ZC3HC1 in ZC3HC1 KO cells harbouring a large pool of nucleoplasmic TPR, this soluble TPR is quantitatively attached to pre-existing NBs and not forming any TPR-containing foci elsewhere (Gunkel and Cordes, 2022; Figure 3E). Together, these findings suggest that soluble ZC3HC1 and TPR can in principle interact through one of their TBE:ZBD pairs, but appear not to engage both pairs simultaneously, thereby avoiding large-scale premature cross-linking via concurrent ZBD1:TBE1 and ZBD2:TBE2a/TBE2b interactions.

However, when NPC anchorage of TPR is no longer possible, as upon RNAi of NUP107, ZC3HC1-dependent TPR foci readily form. Since formation of these foci requires integrity of both TBE1 and TBE2b (Gunkel and Cordes, 2022; Figure 3), and since they rapidly disassemble again upon induced degradation of ZC3HC1 (Gunkel and Cordes, 2022), one can conclude that they result from coupled TBE1:ZBD1 and TBE2:ZBD2 interactions with multiple TPR homodimers. Importantly, ZC3HC1 levels in these RNAi experiments are even far lower than in the recruitment experiments in ZC3HC1 KO cells, whereas TPR amounts are in the same order of magnitude. This argues against a simple concentration effect as the cause of foci formation. However, as also discussed further below, it currently remains unknown why the conditions for higher-order ZBD1:ZC3HC1:ZBD2 interactions are usually met only at the NB, and why disruption of the NPC/NB-anchoring route renders the soluble proteins permissive for ectopic foci assembly. It also remains unknown which of the TBEs or ZBDs are normally prevented from premature ectopic engagement.

### (2) Determinants of ZBD2 and NBD accessibility: exposing and masking

While the principle by which ZC3HC1 and TPR are licensed for higher-order interactions remains unclear, our data suggest that ZC3HC1 binding to ZBD2 and NUP binding to the adjacent NBD are mutually exclusive, at least in the NR and TR contexts examined here. In fact, binding of NUPs to the NBD of NPC-anchored TPR appears to exclude ZC3HC1 from binding the adjacent ZBD2, since ZC3HC1 is detectable neither at the NR in *Xenopus* oocytes nor in somatic cells (Gunkel *et al*., 2021; Figures 5 and 6; Supplemental Figure S2A–S2C). Conversely, neither NUP153 nor NUP107 or other Y complex proteins are detectable in the TR region or at NBLSs (e.g., Krull *et al*., 2004; Gunkel *et al*., 2021; Sabinina *et al*., 2021; Figure 5A; our unpublished data). Taking a closer look at the ZBD2 and NBD, several scenarios can be envisaged for how such mutual exclusivity at the NR and TR may come about.

The ZBD2 of TPR, with its essential residues mapped to aa 386–481, largely corresponds to CC2, which ranges from aa 395–510. Within this region, aa 415– 444 would align with the TBE2 clamp of ZC3HC1 according to AlphaFold3. By contrast, the residues essential for TPR’s NBD are located between aa 450–606. The NBD thus includes a large part of CC2 and almost the entire CC3, which comprises aa 540–622. While the precise docking sites for NUP153 and the Y complex still need to be determined, double aa substitution mutants such as L458D/M489D and L458P/M489P have been shown to impair binding of NUP153 to TPR (Hase and Cordes, 2003). Moreover, these mutations, in particular L458P/M489P, also abolish any type of TPR association with the NE, including appendage to existing NBs (Hase *et al*., 2001), and to NPCs that had been rendered TPR-deficient but remained capable of re- appending TPR (Hase and Cordes 2003). These findings are thus consistent with aa 450–481 having been identified in the current study as a region that provides features essential for functionality of both the ZBD2 and NBD, underscoring the functional overlap between both domains.

In the simplest scenario, binding of NUP proteins to the NBD may directly prevent binding of ZC3HC1 to the ZBD2, and vice versa, by steric hindrance. However, this would still leave open why NUPs prevent ZC3HC1 from binding to the ZBD2 of TPR’s T1 subpopulation at the NR, and, conversely, why NUPs do not detectably bind to the NBD of TPR’s T2 subpopulation at the TR. One possibility may be that NUPs can bind the NBD only when a distinct constellation between Y complex, NUP153, and TPR, existing only at the NR, is provided. In addition, such an NR-specific constellation could have a higher effective affinity for the NBD than ZC3HC1 has for the ZBD2. This constellation could involve an NR-specific arrangement between the Y complex and NUP153, or a specific conformation of TPR’s NBD that can be established only at the NR. These two possibilities are not mutually exclusive and may well complement each other.

In fact, we consider it possible that, within such an NR-specific constellation, an antiparallel alignment of TPR’s CC2 and CC3 may be a prerequisite for a functional NBD. Indeed, the recent finding of a crosslink between aa 477 in CC2 and aa 582 in CC3 (Bräuer *et al*., 2025) may point to such an alignment. Folding of CC3 onto CC2 could then mask the ZBD2 in a manner that prevents the TBE2 α-helices of ZC3HC1 from embracing CC2. Conversely, if such a CC2:CC3 alignment forming a functional NBD is established only at the NR, but not in the TPR T2 subpopulation, this would explain why NUPs are not found at the TR. There, ZC3HC1 would instead be able to bind the ZBD2 of TPR’s T2 subpopulation.

In this context, it should also be noted that the L458/M489 substitution mutants do not abolish homodimerisation of TPR via the neighbouring CCs (Hase and Cordes 2003). However, they probably prevent proper formation of CC2 and thereby directly disrupt the ZBD2. Furthermore, impaired CC2 formation would then no longer allow proper alignment with CC3 and may thus also prevent formation of a functional NBD.

Therefore, instead of a simple scenario in which ZC3HC1 or NUP153 and Y complex would prevent binding of the respective other to the ZBD2 or NBD merely by steric hindrance, one could assume that aligning or not aligning CC3 onto CC2 defines whether ZC3HC1 or NUPs can bind to ZBD2 or NBD. In this view, a “closed” aligned state would allow NUP binding to the NBD and prevent ZC3HC1 binding to ZBD2, whereas an “open”, not aligned state would result in the opposite.

NR-specific arrangements of Y complex and NUP153, or NR-specific CC2:CC3 folding, would also be consistent with the observation that neither native soluble NUP153 nor native soluble Y complex proteins, including NUP107, occur bound to native soluble TPR in substantial amounts in vivo (Shah *et al*., 1998; Harel *et al*., 2003). These findings also argue against a scenario in which premature interactions between soluble ZC3HC1 and the ZBD2 of soluble TPR would be prevented by soluble Y complex proteins or by soluble NUP153 masking the ZBD2 by binding the NBD of such soluble TPR. Moreover, even in a hypothetical setting in which NBD-binding NUPs would mask the ZBD2 of a soluble TPR homodimer, one would still need to explain why NB-bound ZC3HC1, but not soluble ZC3HC1, would then be able to unmask the ZBD2 again for appending TPR to the NB. Thus, this scenario also does not solve the above-mentioned problem of how the ZBD2 may become accessible specifically during the process of appending additional TPR to the NB.

By contrast, the proposed scenario of a constellation-dependent NUP:NBD interaction at the NR, together with a higher effective affinity of the NR-specific NUP constellation for the NBD than of ZC3HC1 for ZBD2, would provide a setting in which TPR anchorage to the NPC and subsequent, ZC3HC1-mediated TPR appendage to the NB would occur in a highly ordered manner. In this setting, the TPR homodimers would first occupy all TPR-binding sites at a newly assembled NPC, which would ensure that assembly of the NB proper is completed before any additional TPR homodimers are appended to it, and thereby prevent additional TPR from being appended to still incomplete NBs.

Differences in NBD and ZBD2 accessibility may also correlate with our finding that certain TPR antibodies specifically detect only one or other TPR subpopulation in situ. However, the inaccessibility of the corresponding epitopes may have different causes. For the TPR T2- specific antibody presented here, which recognises an epitope between aa 498–506, steric masking of this epitope, either by a bound protein or by intramolecular folding of TPR itself, appears conceivable. This epitope is part of CC2, which comprises aa 395–510, and is located close to ZBD2, whose essential residues were mapped to aa 386–481 and whose aa 418–441 segment is predicted to be clasped by the TBE2 α-helices of ZC3HC1. The aa 498–506 epitope is probably not a site of an omnipresent post-translational modification (www.phosphosite.org), and when part of a T1 homodimer, it is neither masked by ZC3HC1 nor by any of the other proteins identified as NB-resident components (our unpublished data). We therefore consider it possible that its inaccessibility may be caused by one of the NR proteins binding to TPR’s ZBD2-overlapping NBD, mapped to aa 450–622. Alternatively, this epitope may be masked by an NR-specific alignment of CC3 onto CC2.

For other TPR antibodies, different mechanisms may underlie their subpopulation specificity, such as subpopulation-specific post-translational modification. This could be the case for the T1-specificity of the TPR antibody recognising an epitope between aa 372–380, and thus between CC1 and CC2. Within this region, S373 has repeatedly been identified as phosphorylated in numerous phosphoproteomic studies addressing constitutive and signal- induced phosphorylation in interphase and mitosis (www.phosphosite.org). The positioning of S373 just before the ZBD2 may indicate that it could play a role in CC2:CC3 folding and unfolding. Thus, both the interaction of TPR’s T1 subpopulation with the NPC and the interaction of TPR’s T2 subpopulations with ZC3HC1 could be regulated by phosphorylation and dephosphorylation of S373.

In addition, other conspicuous TPR residues may also contribute to stimulus-induced assembly or disassembly of the NB proper and of NBLSs. Among these residues are two, T641 and T653, which are frequently found to be phosphorylated (www.phosphosite.org). Located just after the NBD and CC3, it is conceivable that T641 and T653 may also play a role in CC2:CC3 folding and unfolding.

Furthermore, phosphorylation of ZC3HC1 may also contribute to such assembly or disassembly processes. In fact, the N- and C-terminal TBE2 α-helices, comprising aa 29–37 and aa 482–499, are flanked by residues recurrently reported phosphorylated, including S24, T28, S58, S62 and S483 (www.phosphosite.org). Phosphorylation or dephosphorylation at these sites might affect how NB-immobilised ZC3HC1 may specifically interact with TPR subpopulations.

However, while it is tempting to speculate that TPR subpopulation-specific post- translational modifications, and possibly also modifications of ZC3HC1, contribute to the regulation of their interactions at the NB and NBLSs, this remains to be investigated. Ongoing work will need to clarify which of these residues are modified in each TPR subpopulation, whether such modifications alter the folding of TPR’s CC segments, or change ZBD2/NBD accessibility. Future work with phosphopeptide-specific antibodies and single aa substitutions targeting these sites may thereby help answer the question to what extent stimulus-induced phosphorylation affects assembly, maintenance, or disassembly of the NB and NBLSs.

### (3) ZC3HC1-appended TPR: extending NBs beyond the commonly expected

Apart from defining the sequential order by which first ZC3HC1 and then ZC3HC1-dependent subpopulations of TPR are attached to the NB, the two ZBDs also determine how these additional amounts of TPR are positioned relative to the NB. We find these TPR polypeptides appended to the distal end of the NB proper and oriented towards the nuclear interior, thus extending beyond the dimensions of what is commonly regarded as a prototypic NB.

NB-like structures regularly stacked onto each other and appended to some of the NBs in full-grown, i.e. stage VI oocytes of *Xenopus laevis* have been described in the past, either as “*cylindrical extensions … with regularly spaced transverse fibrils*”, with a length of about 250 nm (Scheer *et al*., 1988), or as “*hollow cables with … 50 nm periodic structure*” attached to the NB (Ris, 1997). Furthermore, fibrillar material projecting 250–300 nm deep into the nuclear interior of *Xenopus* stage VI oocytes was found by immuno-EM to be positive for TPR along its entire length, as was NPC-associated fibrillar material reaching up to about 150–200 nm into the nuclear interior in human tissue cells (Cordes *et al*., 1997). Moreover, long CEZs at the nuclear side of NPCs in ultrathin sections of HeLa cells were later also proposed to correspond to these cylindrical NB extensions (Krull *et al*., 2010). Although variable in length, most of these CEZs measured 100–150 nm, while some reached 150–250 nm, reminiscent of channel-like CEZs described in other cell types (e.g., Visser *et al*., 2000; Rego *et al*., 2008), including rat pancreatic cells (Watson 1959). In fact, the original definition of the term “pore complex” did not refer only to elements located directly at the pore through the NE, but also included such cylindrical channels extending deeper into the nuclear interior (Watson, 1959).

Furthermore, analysis of consecutive ultrathin sections through HeLa cells had indicated that these CEZs are often longitudinally surrounded by chromatin, suggesting that they could indeed have a cylindrical shape. Since TPR was essential for their formation (Krull *et al*., 2010), and since TPR had been detected in fibrillar material at corresponding positions in other mammalian cell types (Cordes *et al*., 1997), it was thus already conceivable that TPR was not only a structural component of the NB proper, but also contributed to structures forming the long cylindrical CEZs. Finally, more recent iTEM of NEs from *Xenopus* late-stage II oocytes revealed a seemingly periodic labelling pattern at the nuclear side of NPCs not only for TPR, but also for ZC3HC1 (Gunkel *et al*., 2021). This oocyte stage is characterised by high transcriptional activity and precedes the onset of perinuclear chromatin retraction from the NE in stage IV (e.g., Dumont, 1972; Hill and Macgregor, 1980; Rasar and Hammes, 2006).

Based on these observations, we proposed a model in which both the NB and NBLSs are built from TPR, with ZC3HC1 acting as linker between them (Gunkel *et al*., 2021; Gunkel and Cordes, 2022). Extension of the prototypic NB by a single NBLS would generate a cylindrical structure protruding about 40–45 nm further into the nucleoplasm. Such an NB-appended NBLS could then recruit further ZC3HC1, which in turn could recruit additional TPR, resulting in several NBLSs being stacked on top of each other. This could also explain how the long CEZs noted in numerous cell types can come about.

Thus, several earlier data and models had already indicated that fibrillar extensions of the NB proper, with regular spacing suggesting a repetitive arrangement, may exist not only in the special case of the full-grown amphibian oocyte lacking perinuclear chromatin, but also in early stages of amphibian oogenesis as well as in tumour and non-tumourigenic human somatic cells. However, the prevailing view has long remained that of an NB without additional superstructures. This may partly reflect the fact that most EM images of NBs stemmed from *Xenopus* oocytes and showed NBs without such extensions, which, as an aside, can also rapidly disassemble upon incubation in many standard buffers (e.g., Gunkel *et al*., 2021; our unpublished data). In addition, NBs of *Chironomus tentans* salivary gland cells, another cell type in which NBs can be visualised comparatively well due to the lack of perinuclear chromatin, do not contain such NBLSs, most likely because most insect families, including Diptera, lack a ZC3HC1 homolog (Gunkel *et al*., 2023). Together, these circumstances may have reinforced the view that the NB typically lacks further appended structures. NBLSs occasionally shown on top of the NB may then have been regarded as a special feature of *Xenopus* oocytes.

Against this background, and in light of the data presented here, we now conclude that such NB extensions are not restricted to vertebrate oocytes, but are likely to be widespread also in somatic cells. This conclusion is also supported by the copy numbers of TPR and ZC3HC1 determined here for HeLa cells, which are sufficient to build such structures, as will be discussed further below. It is also supported by the relative amount of ZC3HC1-associated TPR detected in different cell lines. We previously showed that in proliferating cell lines of different germ layer origin, including tumour-derived and immortalised non-tumour cell lines, about half of the total NPC-associated TPR is linked to NBs via ZC3HC1 (Gunkel *et al*., 2021). The current study now extends this observation to the fibroblast line IMR-90, in which about 50% of the total NPC-associated TPR is likewise connected to NBs through ZC3HC1 (Figure 4A), and similar proportions of T1 and T2 subpopulations of TPR were also determined for a wide range of other cell types, including tumour and non-tumour cell lines, primary cells and embryonic stem cells (our unpublished data). These findings indicate that in all these different cell types, the amounts of NB-associated TPR are sufficient to build structures similar to those here presented for HeLa cells.

Our data therefore support a model in which the NB proper represents only the NPC- proximal part of a larger cylindrical scaffold built of TPR and ZC3HC1. More specifically, we propose that ZC3HC1-appended TPR homodimers can form regular structures that maintain the octagonal symmetry of the NB while extending for variable distances into the nuclear interior. This model of octagonally symmetric NB:NBLS superstructures differs from a recent model of NB appendices, based on electron-tomography of human cells, in which only a few individual TPR homodimers were graphically depicted as appendages to some of the NB fibrils (Obarska-Kosinska *et al*., 2026).

Our concept of ZC3HC1-dependent NBLSs as common structural features of human cells also brings our current model closer to Watson’s original, broader concept of the pore complex, which included cylindrical channels extending for variable distances into the nuclear interior (Watson, 1959). As discussed further below, the length of these NBLSs might even be adjustable on demand.

Interestingly, a recent study in yeast (Schmidt *et al*., 2026) also proposed NB appendices as a regular, octagonally organised structure rather than as sporadically appended individual fibrils. Referring to the arrangement of NPC-associated TPR and ZC3HC1 in *Xenopus* oocytes (Gunkel *et al*., 2021), Pml39p-associated Mlp1p homodimers were considered to be part of an octagonally symmetric arrangement extending further into the nuclear interior (Schmidt *et al*., 2026). However, the actual spatial arrangement of these Pml39p-associated Mlp1p homodimers relative to the NB proper was not directly visualised, and the positioning of the NB-appended Mlp1p homodimers was thus also discussed as being uncertain. In fact, while an early iTEM study had suggested that Mlp1p can be associated with fibrils extending from the vicinity of NPCs towards the nuclear interior (Strambio-de-Castillia *et al*., 1999), another study proposed that Mlp1p is not only part of the NB, but may also be arranged horizontally along the plane of the NE and bridge neighbouring NPCs (Niepel *et al*., 2013). Thus, how Pml39p- associated Mlp1p homodimers are arranged relative to the NB proper in yeast remains to be clarified.

In animals, lateral connections between NBs, or NBLSs, have so far been observed only sporadically in oocytes of different amphibian species. In stage VI *Xenopus* oocytes, local regions of the highly folded NE can contain numerous NBs with long NBLS cylinders which sometimes appear to be laterally interconnected (Franke and Scheer, 1970a; Ris and Malecki, 1993; Cordes *et al*., 1997; Arlucea *et al*., 1998). These superstructures were also found to be immunogold-positive for TPR (Cordes *et al*., 1997). In addition, TR-appended lattice-like structures that laterally interconnect NBs have been described in oocytes of the newt *Triturus cristatus* (Goldberg and Allen, 1992). These lattices were considered equivalent to the laterally interconnected NBLS cylinders described for *Xenopus* oocytes (Ris, 1997; Arlucea *et al*., 1998), and were likewise proposed to be built from TPR polypeptides (Goldberg *et al*., 1997; Arlucea *et al*., 1998). Whether these lattices in *Triturus* oocytes also contain ZC3HC1 remains to be shown. Clearly, however, for somatic mammalian cells there is currently no evidence for TPR or TPR:ZC3HC1 ensembles being arranged horizontally along the NE and connecting neighbouring NBs. Instead, available localisation data indicate that all NPC-associated TPR, including its ZC3HC1-appended subpopulations, extends from individual NPCs towards the nuclear interior.

### (4) More TPR than commonly expected: considering numerically possible constellations with NB-associated ZC3HC1

To assess which supramolecular arrangements between TPR and ZC3HC1 at the NB and beyond are numerically possible, we determined their mean copy numbers per NPC and found TPR to be more abundant at the NEs of proliferating cultured cells than formerly anticipated. In fact, the total amount of TPR per NPC had previously been reported to range between eight copies in human DLD-1 cells (Li *et al*., 2021) and 16 copies in HeLa Kyoto cells (Otsuka *et al*., 2023), when determined by light microscopy of FP-tagged versions of TPR, or to be 32 copies in HeLa Kyoto and possibly other proliferating cell lines (Ori *et al*., 2013; Obarska- Kosinska *et al*., 2026), when determining the amounts of NE-associated, non-tagged TPR by mass spectrometric means.

Given that a homodimer is the smallest stable TPR ensemble within the cell (Hase *et al*., 2001), we cannot explain how a copy number of only eight TPR polypeptides would be sufficient for the formation of an eight-fold rotationally symmetric NB. On the other hand, a number of 16 NPC-associated TPR monomers is in principle conceivable. This scenario would mean that only one of the two Y-complex rings forming the NR would be decorated with TPR. However, since we found the amounts of NPC-associated TPR to be highly similar in HeLa P2 and HeLa Kyoto, our data strongly argue against an average of only 16 TPR polypeptides being associated with each NPC in HeLa Kyoto cells.

By contrast, a total of 32 TPR copies, corresponding to 16 homodimers, as determined by mass spectrometry, would indicate that these homodimers would indeed be able to form an NPC-anchored NB in which both Y-complex rings are loaded with TPR. Such numbers, however, would hardly leave any TPR for being additionally appended to the NB, a point also reflected in the schematic depiction of ZC3HC1-appended TPR homodimers, which are shown as attached only to individual but not all NB fibrils (Obarska-Kosinska *et al*., 2026). Furthermore, since only a total of eight ZC3HC1 polypeptides were determined to exist per NPC (Obarska-Kosinska *et al*., 2026), this would exclude any more regular appendage of additional TPR homodimers beyond a first NB-appended NBLS, unlike in a situation in which an average of about 20 stably NB-associated ZC3HC1 polypeptides is available.

However, we regard it as an important aside to also point to a recent study showing that TPR protein levels in TPR^+/−^ mice are drastically reduced, at least in spleen and tail tissues (Liu *et al*., 2026). If the immunoblot signals shown in that study reflected a linear detection range, this raises the question of whether the amount of TPR in some TPR^+/−^ cell types might only allow formation of a normal NB on some NPCs or of a minimal NB scaffold on all NPCs, then with only one of the two nuclear Y-complex rings decorated with TPR homodimers. If the latter were the case, it would mean not only that a total of only 16 TPR copies could occur at the NPC in steady state, but also that such amounts would already suffice for these mice to be viable. On the other hand, though, an alternative explanation might be that the murine WT tissues examined harbour large amounts of nucleoplasmic TPR, which would then be absent from TPR^+/−^ cells, without necessarily affecting the copy numbers of NPC-associated TPR. Clarifying which of these scenarios, or possibly even others, may hold, could be of fundamental importance for how the functional significance of the NB and NBLSs, as will be discussed further below, can be interpreted.

Furthermore, comparison of the relative GFP signal intensities in yeast cells expressing either GFP-tagged Mlp1p, Mlp2p or Pml39p suggested that the Mlp1p:Mlp2p:Pml39p amount relationship is 4:2:1 (Zsok *et al*., 2024). While absolute numbers had not been determined, mean copy numbers per NPC for Mlp1p, Mlp2p, and Pml39p were suggested to be 16, 8 and 4, respectively (Zsok *et al*., 2024), whereas mean per-NPC copy numbers determined for Mlp1p and Mlp2p in other studies were 15 for Mlp1p and 13 for Mlp2p (Mi *et al*., 2015), about eight each for Mlp1p and Mlp2p (Kim *et al*., 2018), and 5.2 for Mlp1p and 3.9 for Mlp2p (Rajoo *et al*., 2018). However, these overall low numbers were considered to be due to not all yeast NPCs being equipped with an NB (Rajoo *et al*., 2018), and, in fact, even among those NPCs that are positive for Mlp1p, only about 28% display a complete NB (Singh *et al*., 2024). Together, these data thus indicate that NPC loading with Mlp homodimers in yeast is more heterogeneous than in mammalian cells. Taking this into account, it was recently proposed that the actual composition of a complete yeast NB on its own, i.e. without possible NB extensions, consists of 16 copies each of Mlp1p and Mlp2p. To this NB proper, eight copies of Pml39p could then be bound, which in turn could recruit 16 copies of a second Mlp1p subpopulation (Schmidt *et al*., 2026). This again would be in line with the previously observed loss of an NPC-associated subpopulation of Mlp1p in *pml39*Δ cells (Gunkel *et al*., 2023), and explain the Mlp1p:Mlp2p:Pml39p ratio of 4:2:1 (Zsok *et al*., 2024).

Furthermore, in this recently proposed assembly model, one NPC-anchored Mlp2p homodimer and one Mlp1p homodimer per NB fibril would be required for cooperatively recruiting one Pml39p, which could then recruit yet another Mlp1p homodimer. However, this second Mlp1p homodimer, lacking a second Mlp2p homodimer aligned with it, would be unable to recruit yet another copy of Pml39p (Schmidt *et al*., 2026).

By contrast, in HeLa, HCT116 and DLD-1 cells, and presumably in numerous other human cell lines, we are dealing with much higher copy numbers and with multilayered NB extensions that are much more omnipresent than previously assumed. In fact, we now have to consider mean values of about 61–63 TPR copies and 20 ZC3HC1 copies per NPC in HeLa cells, and likely similar copy numbers in many other human cell lines. We thereby have to consider at least three NPC-associated subpopulations of TPR and two of ZC3HC1 that are positioned at different distances from the NPC proper, as can be deduced from our STED micrographs and seen in the STORM images. This multilayered arrangement already indicates that not every aspect of the self-limiting assembly model proposed for the yeast NB (Schmidt *et al*., 2026) can be directly applied to the NBs and their NBLSs in vertebrates.

Anyhow, one would not necessarily expect that the assembly processes leading to NB extensions in yeast and mammalian somatic cells would have to be identical: The approximately spherical nucleus of budding yeast has a diameter of about 1.8–2.0 µm (e.g., Wang *et al*., 2016) while the nuclei of somatic mammalian cell types, ranging from spherical to ovoid in shape, are generally much larger, with diameters commonly ranging between 5 and 20 µm (e.g., Lammerding, 2011; Gillooly *et al*., 2015; Skinner and Johnson, 2017; Sebastian *et al*., 2021 and references therein). In addition, such mammalian nuclei often harbour varying amounts of heterochromatin unevenly distributed along the NE. Therefore, NBLS assembly in mammalian nuclei may need to occur without an inherent self-limitation, allowing stacks of NBLSs to span larger distances when required. On the other hand, ZC3HC1 and the long NB extensions enabled by this protein may also have disadvantages, resulting in an evolutionary balancing act between the pros and cons of possessing ZC3HC1, as previously suggested (Gunkel *et al*., 2023). In mammals, the advantages of being able to form long NB extensions have apparently outweighed their possible disadvantages. In budding yeast, by contrast, where a comparable need for cylindrical stacks of NBLSs reaching into the interior of a much smaller nucleus may not exist, the potential disadvantages of such long NB extensions may have contributed to the emergence of the length-restricting mechanism outlined above.

Nevertheless, we should not exclude that cooperativity between the two NPC-anchored TPR homodimers, which would correspond to the NPC-anchored pair of an Mlp1p and an Mlp2p homodimer, may play some role also in mammals in the initial recruitment of ZC3HC1 to the NB. Further along this line, if NBs composed of only 16 copies of TPR existed in TPR +/- mice, it would also enhance our understanding of the NB assembly process in vertebrates to know whether such minimalistic NBs would still be capable of binding ZC3HC1, i.e. without cooperativity between two aligned TPR homodimers.

Here, however, we first need to address the question of how ∼62 copies of TPR, distributed essentially across three NPC-associated layers, and ∼20 copies of ZC3HC1, mainly present in two layers, can be arranged relative to one another within the NPC- and NB-associated TPR:ZC3HC1 ensembles of a proliferating human cell line. Since the numbers determined for both proteins are not exact multiples of eight, one also has to assume a certain degree of heterogeneity between the NB extensions at different NPCs, which may also be reflected in the STORM image compilations (Figure 6B). In this regard, it appears conceivable, but not mandatory, as discussed below, that such differences between individual NPC:NB:NBLS ensembles may affect primarily those layers of TPR and ZC3HC1 most distal from the NPC. Heterogeneity regarding these layers may be caused by the available TPR and ZC3HC1 amounts being insufficient for building extensions of identical TPR and ZC3HC1 copy number on all NBs.

For TPR, one can make an additional assumption. If we assume that the NB proper in WT cells is composed of the same number of 32 NPC-anchored TPR T1 polypeptides that we determined as the absolute TPR copy number per NB in ZC3HC1 KO cells, this would leave about 30 TPR copies in the WT cells as the T2 subpopulations. These remaining copies would then have to be distributed across the two clearly discernible additional layers of TPR, if the rarely observed fourth TPR layer is regarded as numerically irrelevant in this context. This T2 subpopulation could thus be subdivided into two further subpopulations, T2a and T2b, with T2a corresponding to a first NBLS, here referred to as NBLS1, and T2b corresponding to a further NBLS, here referred to as NBLS2, which would be appended to NBLS1. If we further assume eight-fold rotational symmetry also for the NBLSs, this would mean that the first NBLS would consist of 16 TPR copies of the T2a subpopulation. The remaining 14 TPR copies of the T2b subpopulation could then either form consistently incomplete NBLS2 structures, or be arranged such that about 75% of all NB:NBLS1 ensembles would carry a second, complete NBLS2, whereas the remaining 25% of NB:NBLS1 ensembles would lack such an NBLS2. A gradual transition between both scenarios is of course also conceivable, with different proportions of the NB:NBLS1 ensembles carrying complete or incomplete NBLS2s, or lacking any additional TPR T2b. Averaged across all NPCs, however, this distribution could in any case be simplified as a 32xT1:16xT2a:16xT2b-partial arrangement.

One could argue, however, that the appearance of the NBs and NBLSs in *Xenopus* oocytes does not seem to fit well with such a 32xT1:16xT2a:16xT2b distribution. When comparing the NB and NBLS fibrils in conventional TEM and SEM micrographs of stage VI oocytes, no obvious difference in fibril thickness has so far been apparent (e.g., Scheer *et al*., 1988; Ris and Malecki, 1993; Gunkel *et al*., 2021). If the NB proper in *Xenopus* stage VI oocytes is indeed composed of 32 TPR polypeptides, as will be discussed further below, one could thus assume that oocyte NBLSs likewise contain 32 TPR copies if NB and NBLS fibril diameters are indeed the same.

On the other hand, we cannot exclude that thickness differences are no longer recognisable because of preparation artefacts introduced during conventional EM sample processing, which will be discussed further below. In addition, we cannot exclude that the TPR copy number per NBLS differs between oocytes and somatic cells, and that oocytes can indeed assemble NBLSs whose fibrils contain twice as many TPR polypeptides as those in somatic cells. More informative for the situation in human cells would be comparisons between the thickness of NB and NBLS fibrils as they appear in electron-tomographic datasets. Based on the primary ET data shown so far, which also include possible NBLS fibrils (Obarska-Kosinska *et al*., 2026), we could not, however, draw firm conclusions regarding possible thickness differences between NB and NBLS fibrils after measuring these structures in the images presented. In this context, we therefore await further insights from future ET studies.

The 32xT1:16xT2a:16xT2b-partial scenario, however, is only one of several that one can reflect upon. For example, in one of these other scenarios, 32 TPR copies would again be assigned to the NB proper and thus to the T1 subpopulation. In this case, however, each NBLS too would be built from 32 TPR polypeptides. The discrepancy between the resulting 96 TPR polypeptides that would be required to equip all NPCs with NBs plus two NBLSs, and the actually available number of only about 61–63 TPR polypeptides for such arrangements on average, could then be resolved only by assuming considerable heterogeneity in the number and completeness of NBLSs associated with individual NBs. Such heterogeneity could manifest as significant numbers of NBs without an NBLS, next to others with a single, complete NBLS composed of 32 TPR copies, and yet others with NBs that carry either two complete NBLSs composed of altogether 64 TPR copies, or a complete NBLS1 plus incomplete NBLS2. This scenario would at least harmonise with the abovementioned fact that no clear difference between NB and NBLS fibril thickness has so far been observed in *Xenopus* oocytes. Apart from that, however, we cannot readily reconcile this scenario with our STORM data, as the frequent occurrence of three TPR layers would, under this model, have to be accompanied by a large proportion of NBs lacking ZC3HC1-dependent TPR extensions, for which we find little evidence.

In another scenario, all NPCs in WT cells would have an NB, which, however, would not necessarily need to be composed of 32 T1 copies. Indeed, the initial assumption could be wrong, namely that the 32 TPR copies per NPC determined in ZC3HC1 KO cells would also have to represent the T1 copy number in WT cells. Instead, one could imagine that a latent excess of TPR in the KO cells leads to quantitative saturation of all TPR-binding sites at the NPC. In WT cells, by contrast, competition between TPR recruitment to the NPC and TPR recruitment to the NB might result in situations in which not both Y-complex rings are quantitatively occupied by TPR, because part of the TPR pool has already been captured by ZC3HC1 and appended to the NB. If the overall number of T1 polypeptides were lower, more TPR copies would be available for the T2a and T2b subpopulations. However, in order to then reach the average of ∼62 TPR copies per NPC, one would then need to further postulate that among the subpopulation of NBs comprising 32 TPR copies some would also need to have 32 TPR copies of the T2a subpopulation appended to them. Altogether, this scenario would thus come along with a particularly high degree of heterogeneity between the NB:NBLS ensembles, which one could perhaps describe as 16–32xT1:16–32xT2a:16–32xT2b-partial. However, this scenario would invalidate some of the other considerations discussed above, according to which it would appear more plausible that all TPR-binding sites at the NPC would need to be occupied first before further assembly of NBLSs can occur via binding to ZC3HC1. Furthermore, the current ET data from murine and human cells do not appear to point to pronounced heterogeneity in the thickness of the presented NB and NBLS fibrils (Singh *et al*., 2024; Obarska-Kosinska *et al*., 2026). At this point, we thus cannot regard any of these and yet other scenarios regarding the TPR constellations within the NB:NBLS ensembles as proven or categorically excluded, because the actual TPR copy-number relationships between the individual layers remain unknown. The COM-aligned STORM data suggest that the NPC- proximal T1 layer contains, on average, more TPR than T2a, and T2a more than T2b, but do not establish numerical ratios. Importantly, any model must also accommodate the independent finding that only about 45–55% of the total NPC-associated TPR is bound via ZC3HC1 (Gunkel et al., 2021; Gunkel and Cordes, 2022; current study).

Furthermore, we also still need to assess which scenario may be compatible with the mean copy number of about 20 ZC3HC1 polypeptides per NPC. If we again assume that the binding sites for ZC3HC1 at the NB or at an NBLS occur in sets of eight, or in multiples of eight, the number of 20 would at first glance allow only for two basic distribution patterns. In the one scenario, 16 copies of ZC3HC1 would all belong to subpopulation Z1, forming a complete layer of ZC3HC1 polypeptides sandwiched between TPR’s NB-forming subpopulation T1 and its NBLS1-forming subpopulation T2a. The remaining four copies of ZC3HC1, belonging to subpopulation Z2a appended to NBLS1, would then again recruit some TPR polypeptides, but not in numbers sufficient to form complete NBLS2 structures on all NB:NBLS1 ensembles. In the other scenario, the 20 copies would be allocated to two complete layers with eight ZC3HC1 copies each, Z1 and Z2a, plus another, again incomplete layer, Z2b, with four copies.

A 16:4 distribution, in which 80% of the ZC3HC1 copies would belong to the proximal Z1 population, would appear difficult, though, to reconcile with the ZC3HC1 signal distribution observed by STED microscopy, with the notably broader ZC3HC1 fluorescence band in WT cells and the markedly greater distance of its signal peak from the NR than in the ZC3HC1 mutant cells. Furthermore, the relative signal intensities of the two ZC3HC1-sfGFP layers in the COM-aligned STORM datasets do not appear to reflect the fourfold difference expected from such a distribution either.

By contrast, an 8:8:4 distribution might appear more readily compatible with the STED data, since two equally sized but spatially separated Z1 and Z2a subpopulations, plus a smaller Z2b subpopulation positioned even farther away from the NPC, would be consistent with the broader ZC3HC1 signal distribution in WT than in ZC3HC1 mutant cells and with the resulting signal peak being positioned farther from the NR. However, while the STED data do not exclude such a distribution, the actual position of the remaining four ZC3HC1 polypeptides would remain elusive, as the averaged line profile (Figure 5D) does not allow to unambiguously infer a smaller Z2b subpopulation positioned even farther from the NB than the Z2a polypeptides. More importantly, the COM-aligned STORM data (Figure 6B) provide no indication of a numerically substantial third Z2b subpopulation, as no prominent third layer of ZC3HC1-sfGFP has been identified in this compiled dataset.

Nonetheless, there might appear to be a possible way out of this dilemma if one again assumed that the NBs and their NBLS extensions are much more heterogeneous in their composition than we have assumed so far. One could then assume that some NBs would have 8 copies of the Z1 subpopulation bound to them while others would have 16 copies of Z1, depending on whether their NB and NBLS fibrils contain one or two aligned TPR homodimers. Such heterogeneity could yield on average a two-layered ZC3HC1 distribution with a more signal-intense proximal than distal ZC3HC1 layer, without requiring a uniform 8:8:4 distribution across three layers. Furthermore, such heterogeneity between individual NB:NBLS ensembles would nonetheless allow us to describe the relative relationships between TPR and ZC3HC1 in the different layers in a simplified manner as 2xT1:1xZ1:2xT2a:1xZ2:2xT2b, meaning that one ZC3HC1 polypeptide would in principle link an already anchored TPR homodimer to another TPR homodimer of the next assembly stage.

However, while this basic copy number relationship is in line with many of the currently available data, it does not fully reconcile all our results so far. Thus, further work will be required to clarify how NBs and NBLSs are composed in HeLa and other cell types, which range of structural conformations exists not only between different cell types but also within a given nucleus, and how such intranuclear variability may then come about.

### (5) Putative ZC3HC1 homodimerisation: considering a transient assembly intermediate

After considering possible T1:ZC3HC1:T2 arrangements, one also needs to consider two further observations that raise the possibility that ZC3HC1 may transiently homodimerise during the assembly process. First, in previous work, we had noted, upon adding TPR-free recombinant ZC3HC1 in NB assembly buffer to NE scaffolds prepared from ZC3HC1 KO cells, that two equal-sized subpopulations of ZC3HC1 bound to these NBs. However, only one of them remained persistently bound, whereas the other one detached rapidly and quantitatively when the NBs were washed and then incubated in buffer alone (see Supplemental Figure S6C in Gunkel and Cordes, 2022). Second, in the current work, we noted that apparently two subpopulations of ZC3HC1 Δ487–502-sfGFP bind to TPR’s NPC-anchored T1 subpopulation. Again, only one of them, which we now know comprises eight copies of ZC3HC1, remained robustly bound to the NBs during cell permeabilisation, whereas the other, also comprising eight copies of ZC3HC1, was quantitatively detached from the NBs by the same procedure (Figure 3C2).

These findings need to be considered together with the facts that the BLDs of ZC3HC1 are closely related to the BIR domains of other proteins (e.g., Higashi *et al*., 2005; Kokoszynska *et al*., 2008; Gunkel *et al*., 2021, 2023), that at least BIR1 is capable of homodimerisation (e.g., Lu *et al*., 2007; Mace *et al*., 2010; Hou *et al*., 2017; Yang *et al*., 2020), and that AlphaFold3 can predict a ZC3HC1 homodimer in an arrangement with TPR (Supplemental Figure S7G). Thus, we wonder whether transient homodimerisation of ZC3HC1 may represent one step in an ordered assembly sequence leading to a higher-order TPR:ZC3HC1:TPR ensemble. In such a scenario, the first ZC3HC1 copy, here referred to as Z1a, would bind robustly to an NPC- anchored TPR homodimer of the T1 subpopulation. However, for recruitment of a TPR homodimer of the T2 subpopulation, a second ZC3HC1 copy, here referred to as Z1b, might be needed, with Z1b only weakly homodimerising with Z1a. Once the T2 homodimer has been recruited and stably engaged, Z1b would no longer be required and could detach, leaving only the robustly bound Z1a as the actual link between the two TPR homodimers.

In principle, one could also imagine that Z1a and Z1b bind to the two NPC-anchored TPR homodimers that together form one of the eight NB fibrils (see further below). In such a scenario, again only Z1a would be capable of binding robustly on its own to one of these TPR homodimers, whereas Z1b would become robustly anchored at the NB only after recruitment of a T2 homodimer, and would engage only weakly with Z1a in the absence of a T2 homodimer. However, this scenario would demand that 16 copies of ZC3HC1 were sandwiched between the T1 and the first T2 subpopulation of TPR, and since the average number of ZC3HC1 polypeptides per NPC in HeLa is about 20, this would only leave 4 copies for appending the second to the first T2 subpopulation. As discussed above, this setting would appear difficult to reconcile with the ZC3HC1 signal distributions observed by STED microscopy, in particular with the markedly greater distance of its signal peak from the NR in the WT than in the ZC3HC1 mutant cells.

More generally, it remains unclear whether the second, weaklier NB-associated ZC3HC1 interacts directly with the stably bound ZC3HC1. Neither Y2H experiments using ZC3HC1 as both bait and prey nor in vitro interaction experiments using soluble, differently FP-tagged versions of ZC3HC1 provided evidence for interactions between soluble ZC3HC1 polypeptides (our unpublished data). Therefore, one would have to postulate that any homodimerisation between Z1a and Z1b could occur only when Z1a adopts a distinct NB- bound conformation that enables binding with Z1b. However, since this assumption remains speculative, it remains an open question whether physiological ZC3HC1 homodimers exist at all - and, if so, whether they function as transient intermediates during the TPR:ZC3HC1:TPR assembly process.

### (6) The NB proper: a fishtrap, basket, or cylindrical cage?

After considering how additional TPR can be appended to the NB through ZC3HC1, one also needs to reassess the structure of the NB proper itself, in light of the recent ET studies of the NBs in yeast and mammalian cells (Singh *et al*., 2024; Obarska-Kosinska *et al*., 2026).

What is now referred to as the NB proper may have been first described more than five decades ago in TEM images of human melanoma cells as an NPC-associated structure composed of eight fibrils (Maul, 1971). A similar structure observed in SV40-infected monkey kidney CV1 cells was then described as fishtrap-like (Maul, 1976). The corresponding structure was again referred to as a fishtrap, but also as a nuclear cage or basket, when visualised by SEM of *Xenopus* oocyte NEs (Ris 1989, 1991; Jarnik and Aebi 1991).

Over the following decades, numerous SEM studies of NBs in different cell types, including avian oocytes as well as *Chironomus tentans* salivary gland cells (e.g., Kiseleva *et al*., 1996; Goldberg *et al*., 1997), consolidated the view of the NB as a basket-like structure. In this view, the distal ends of the eight NB fibrils are inclined towards the longitudinal axis, giving the NB the overall appearance of a truncated cone. A compatible conclusion was also drawn from stereometric measurements of the smallest versions of the NPC-associated CEZs in poliovirus- infected HeLa cells. Although individual fibrils could not be resolved in these thin sections, the dimensions of these CEZs were similar to the space enclosed by NBs in amphibian oocytes, i.e. a truncated pyramid with an octagonal base (Krull *et al*., 2010).

However, recent ET studies now challenge the general validity of this conventional EM- based view, first in yeast and mouse 3T3 cells (Singh *et al*., 2024), and more recently also in human cell types (Obarska-Kosinska *et al*., 2026). In these studies, the mammalian NB was modelled as a circular arrangement of freestanding fibrils, presented as either nearly parallel to the NB’s central axis (Obarska-Kosinska *et al*., 2026) or as tilted inward by only 6° (Singh *et al*., 2024).

Therefore, one first has to ask whether the largely perpendicular arrangement of the NB fibrils on the NR in mammalian cells, as suggested by the recent ET studies, rules out that the truncated-cone-like appearance of the NB seen in many conventional EM studies reflects a physiological NB conformation. Alternatively, one would have to conclude that the appearance of the NB as a fishtrap- or basket-like structure merely reflects an artefact of chemical fixation and dehydration.

There are indeed several reasons why inward inclination of NB fibrils could result from conventional EM sample preparation. One conceivable source of such an artefact may be the numerous, largely unstructured CTDs of TPR, which emanate from the distal parts of the NB- fibril-forming rod domains of TPR and extend into the space enclosed by these fibrils. If these CTDs, possibly together with CTD-interacting proteins, form a dense phase of intrinsically disordered regions (IDRs) within this space, aldehyde fixation could alter the physical state of this phase. In fact, FA fixation has been shown to substantially change the apparent morphology of phase-separated IDR-driven condensates, and low concentrations of GA can even solidify such condensates under defined ionic conditions (Irgen-Gioro *et al*., 2022; Beck *et al*., 2024). Since the CTDs of vertebrate and insect TPR contain about 10–13 lysine residues, and since FA can additionally react with several other aa residues, FA- and GA-mediated cross- linking could in principle convert the initially liquid phase into a collapsed and thus more densely packed state, with such shrinkage and compaction possibly further pronounced during subsequent dehydration. While the NPC-proximal ends of the NB fibrils are anchored at the NR, shrinkage of the CTDs connected to the distal ends could pull these ends towards the longitudinal axis and thereby cause the fishtrap- or basket-like, aka truncated-cone-like appearance of the NBs. In principle, we have to consider that FA fixation of our specimens for STORM microscopy may also have caused a certain degree of CTD cross-linking and compaction and may thereby have contributed to the truncated-cone-like appearance in the composite views of the COM-aligned NB:NBLS ensembles. In this particular case, however, we also cannot categorically exclude that this appearance in the stacked STORM images may, to a minor extent, have been influenced by the fact that the 200 nm-diameter, 300 nm-high cylindrical volumes used for particle extraction were not computationally corrected for potential tilts of individual NB:NBLS ensembles relative to the z-axis. Such tilts could occur when particles are selected from NE regions that are not perfectly planar but slightly curved, as at nuclear poles. However, given the 200 nm diameter of the extraction volume, an NBLS diameter of only about 90 nm, and the distance of about 120 nm between ELYS, representing the NR, and the NPC-distal part of an NB:NBLS:NBLS ensemble, loss of signal from the most distal parts of an NBLS would, if at all, be expected only at tilt angles of about 25° or more. Even for NPCs selected from slightly curved NE regions at nuclear poles, such strong tilts should, if they occur at all, represent rare events.

Assuming that the often-discussed conformational variability of the NB could reflect different degrees of fixation- and dehydration-induced CTD compaction, this may then also apply to the pronounced inclination of the NB fibrils described to occur during the translocation of a very large mRNP, namely the Balbiani ring (BR) particle, in the salivary gland cells of *Chironomus tentans*. There, striking differences in NB appearance during BR particle translocation were interpreted as translocation-driven structural rearrangements of a remarkably dynamic NB (Kiseleva *et al*., 1996, 1998). While we do not exclude that physiological plasticity of the NB could play a role in translocating the BR particle or other cargo complexes through the NB, we propose that conventional EM alone may not suffice to unambiguously distinguish native conformational variability from fixation- and dehydration- induced artefacts.

While the recent ET depictions of the NB scaffold in human cells convey the impression of a static and invariant structure composed of evenly arranged rigid rods, this view also contrasts with an atomic force microscopy (AFM)/STORM study that attributed pronounced conformational flexibility and variability to NBs in U-2 OS cells. In that study, NBs were described as “soft” structures capable of regularly adopting a broad range of conformations, including collapsed and largely disassembled states (Vial *et al*., 2023). However, we believe that these AFM/STORM-derived conformations also need to be interpreted with caution, because sample preparation conditions similar to those used in this study have been shown to promote NB disassembly and collapse of NB fibrils when applied to manually isolated *Xenopus* oocyte NEs with initially intact NBs (Krull, 2011; Gunkel *et al*., 2021; see also Jarnik and Aebi 1991). Furthermore, such conditions also destabilise NBs in cultured mammalian cells and have therefore been avoided in cell fractionation experiments (Gunkel *et al*., 2021; Gunkel and Cordes, 2022). Thus, we wonder whether the various NB conformations described by AFM may at least in part reflect preparation-induced NB destabilisation, rather than conformations commonly adopted by intact NBs in vivo.

At the same time, though, also the recent ET data might not describe all possible mammalian NB conformations. Since these NB reconstructions are based on subtomogram averaging, some conformational heterogeneity of individual NBs may, in principle, have been lost during such averaging. In this respect, these ET data may face a limitation similar to that of our COM- aligned STORM data, in which potential heterogeneity in the appearance of NBs and their appendices, whether physiological or already induced by FA fixation, would likewise not be visible in these composite views. For the HEK293 dataset (Obarska-Kosinska *et al*., 2026), the NPC particles used for averaging were manually picked from filtered tomograms, and, with NB fibrils reportedly also visible in the primary data, we currently cannot assess whether fibril appearance may have played a role in particle selection. In fact, we do not know whether NPCs with particularly conspicuous NB fibrils may have been preferentially selected, either because being already apparent in the primary data or because they were especially well visible after filtering, potentially underrepresenting other NB conformations. However, since NB fibril appearance was not stated to be a selection criterion, we consider this point merely an uncertainty of method description whose clarification would further strengthen the interpretation of these ET data.

Nonetheless, although we still refrain from definitively excluding the possibility that, in some cell types or under certain conditions in vivo, the NB may adopt a physiological conformation resembling a fishtrap- or basket-like structure, the recent ET data suggest that the most plausible explanation for this appearance in conventional EM specimens is a fixation- and dehydration-induced deformation artefact.

### (7) The NB proper: delicate fibrils or thick struts?

A second point requiring a reassessment of the conventional EM-based data relates to NB fibril thickness. In conventional EM of amphibian oocytes, the NB fibril width appears to range between about 3–5 nm, when both coating with heavy metals and treatment with tannic acid are omitted (e.g., Franke, 1970; Franke and Scheer, 1970a; Scheer, 1972; Jarnik and Aebi, 1991; Cordes *et al*., 1993; Krull, 2011; Gunkel *et al*., 2021). By contrast, the subtomogram averages of the recent ET studies indicate much thicker fibrils, at least with regard to the longer axis of their cross-sections. In 3T3 cells, NB fibril width was reported to be about 10 nm (Singh *et al*., 2024), consistent with their designation as struts. Furthermore, although no numerical diameter was provided for the NB fibrils in human cells (Obarska-Kosinska *et al*., 2026), the longest axes of the cross-sections shown for the subtomogram-averaged, rod-like portions of these NB fibrils appear to range between 9 and 10 nm (see also further below). Therefore, one again has to ask whether the thin fibrils seen in conventional EM may represent either a physiological, perhaps oocyte-specific state of the NB, or whether chemical fixation, osmication, and dehydration may have caused an artefactual reduction in apparent fibril diameter.

Before considering this possibility, however, it should be noted that we regard it as unlikely that only one of the two Y-complex rings forming the NR is equipped with TPR, such that only 16 instead of 32 TPR copies would form the NB proper in oocytes. However, before outlining two observations that argue against this scenario, one needs to consider the following: The NBs previously studied by conventional TEM or SEM with regard to NB fibril thickness were certainly, or at least most likely, from stage VI oocytes of well-fed frogs. In these oocytes, NE areas with extensive NBLS superstructures on top of the NBs are frequent. By contrast, in stage V oocytes from young, very slim frogs, which are rarely used for such investigations, stacks of NBLSs are seen only sporadically, rendering such oocytes more suitable for estimating the relative amounts of TPR associated with the NB proper alone. In fact, when the relative amounts of NE-associated TPR, ZC3HC1 and NUP107 in such stage V oocyte NEs were compared with those in *Xenopus* cultured cells, the ratios were rather similar (Gunkel *et al*., 2021). Since these data did not indicate a lower amount of TPR per NB in the stage V oocytes, and since we now have no reason to assume that the NB proper in *Xenopus* cultured cells contains only 16 instead of 32 TPR copies, this finding supports the notion that the NB proper in stage V oocytes is also built from 32 TPR copies. However, to extend this argument to stage VI oocytes of well-fed frogs, side-by-side preparation and inspection by SEM of stage V and stage VI oocytes from differently aged and differently fed frogs had to be conducted. This comparison did not reveal any obvious difference in the appearance of the NB fibrils themselves (our unpublished data), now supporting the conclusion that the NB proper in stage VI oocytes too is composed of 32 TPR copies.

A second observation also argues against a scenario in which the thin appearance of oocyte NB fibrils would simply reflect the presence of only one TPR homodimer per fibril. Although observed only rarely, one can occasionally find, also in stage VI oocytes, that one of the eight fibrils constituting an NB appears as a fibril that is even far thinner than those commonly found attached to the NR. These extremely thin fibrils are then always seen attached to the inner half of the NR and thereby appear lacking the other half of the NB fibril, which usually appears to be attached to the outer half of the NR. Together, these observations indicated early on that an NB fibril in amphibian oocytes represents an alignment of two fibrillar rods, of which the outer one very rarely can be missing. In SEM specimens prepared without tannic acid and without additional heavy-metal coating, these extremely thin inner fibrils appear to have a circular profile and diameters of only about 2–2.5 nm. By contrast, the presumably complete fibrils sometimes appear as flattened rods, when tilted NBs are inspected in side view, instead of their more common inspection in top view. When we measured these complete fibrils at their thinnest part in top views of the NBs in SEM micrographs, the width of what we regarded as the short axis of an elliptic or disco-rectangular cross-section was around 3 nm, whereas the long axis, visible in side view, could reach values between 4–5 nm (our unpublished data).

Thus, if the NB proper in *Xenopus* oocytes also contains 32 TPR copies, a complete NB fibril must accommodate two folded TPR homodimers. In addition, the fibril width would have to be large enough to also accommodate one MAD1 homodimer, because MAD1 is a CC protein that forms homodimers (Jin *et al*., 1998; Sironi *et al*., 2002; Kim *et al*., 2012) and because it is a common, TPR-binding component of the NB in human cell lines and *Xenopus* oocytes (e.g., Campbell *et al*., 2001; Hase *et al*., 2001; Lee *et al*., 2008; Krull, 2011; Gunkel and Cordes, 2022). MAD1 homodimers appear to engage TPR’s T1 population, but an NB:NBLS ensemble may play a role in keeping them stably NB-associated (our unpublished data). The MAD1 homodimers themselves are dispensable though for NB formation and maintenance, as also holds for several other NB-associated proteins like GANP and SENP1 (Lee *et al*., 2008; Mossaid *et al*., 2020; Bensidoun *et al*., 2022; Supplemental Figure S8A).

Furthermore, the C-terminal region of its extended CC folds back towards its N-terminal part (Sironi *et al*., 2002; Fischer *et al*., 2022; Chen *et al*., 2023). In fact, the two arms of a folded *Xl*MAD1 homodimer would be long enough to align with the entire length of the CC1:CC4 region of *Xl*TPR (Supplemental Figure S8B1), and this holds also for the human homologs (Supplemental Figure S8B2), as recently also predicted for the human NB fibril (Obarska-Kosinska *et al*., 2026). An extended alignment between TPR and MAD1 homodimers would also conform with MAD1 interacting with several CC segments of TPR in the Y2H system (our unpublished data).

Based on these assumptions, one can model the following arrangement. If the two TPR homodimers are folded hairpin-like only once, such that their CC4 and CC1 segments align and form two homodimeric CC1:CC4 bundles, but without additional alignment of CC5–7 with CC1:CC4, two principally conceivable arrangements could be formed together with one folded MAD1 homodimer (Supplemental Figure S8C4 and S8C5). In one arrangement, MAD1 would interact with both TPR bundles; in the other, it would align with only one of the two TPR bundles, whereas the MAD1-binding sites of the other TPR bundle would no longer be accessible because of TPR:TPR interactions. In the first case, the resulting bundle of six CCs could be simplified as TPR:MAD1:TPR, and in the second as TPR:TPR:MAD1, both of which, though, could have similar cross-section dimensions. In fact, given that the diameter of a prototypical dimeric CC can be about 1.6–1.8 nm (e.g., Morgan *et al*., 1993; Whitehead *et al*., 2007; Kennaway *et al*., 2009; Kawakita *et al*., 2016; Sapra *et al*., 2020), each once-folded homodimer would form one bundle of two aligned CCs, with cross-sectional axis lengths of about 3.2 × 1.6 nm. If two such TPR bundles lie side by side and one once-folded MAD1 homodimer aligns with either one or both of them, the resulting TPR:MAD1:TPR or TPR:TPR:MAD1 bundle would be expected to have cross-sectional axis lengths of about 3.0 × 5.6 nm (Supplemental Figure S8C4).

The fact that NB fibrils in conventional EM specimens were often measured as even thinner could reflect that the narrow side of the fibrils was preferentially measured. More importantly, however, artificial compaction of the filament during EM sample preparation, in particular through the use of OsO_4_, also has to be considered. Indeed, OsO_4_ in combination with GA can drastically alter and disrupt protein secondary structure and conformation, including a major loss of α-helical components (e.g., Lenard and Singer, 1968; Puchtler *et al*., 1985). More generally, aldehyde fixation, OsO_4_ fixation, and dehydration can cause protein aggregation and compaction, as well as collapse of hydrated structures (Studer *et al*., 2008; Akhtar, 2012; Xu *et al*., 2024), and could thereby also reduce the apparent diameter of NB fibrils to some extent. Therefore, the measured thickness of NB fibrils after conventional fixation, osmication, and dehydration should be generally interpreted with caution.

However, while measured oocyte NB fibril widths of about 2.5–5.0 nm still appear compatible with some degree of fixation- and dehydration-induced compaction of a bundle comprising one MAD1 homodimer and two TPR homodimers - when each homodimer is folded only once in a hairpin-like manner - it is much more difficult to imagine how these thin NB fibrils could accommodate four additional coiled-coil bundles. Such additional bundles would arise if the TPR homodimers folded severalfold, aligning not only CC4 onto CC1 but also CC1:CC4 with CC5:CC6/7. In this case, several arrangements would be conceivable; however, even the most compact versions of the more plausible arrangements would be expected to have cross-sectional axis lengths of about 7.2 × 4.8 nm or 8.8 × 3.2 nm (Supplemental Figure S8C5). Thus, all these arrangements would be substantially larger than the measured width range of the oocyte NB fibrils seen in conventional EM specimens.

At the same time, however, crosslinking data from different human cell types suggest that such multiple folding of TPR may indeed occur, at least in proliferating cell types (Liu *et al*., 2017; Cardon *et al*., 2019; De Graaf *et al*., 2019; Bartolec *et al*., 2023; Bräuer *et al*., 2025; Obarska-Kosinska *et al*., 2026). It is therefore plausible that TPR is indeed folded severalfold in somatic mammalian cell types, and a TPR:TPR:MAD1 bundle with cross-sectional axis lengths of 8.8 × 3.2 nm would also fit well with the recent cross-sectional views of subtomogram averages of NB fibrils from HEK293 cells and macrophages (Obarska-Kosinska *et al*., 2026). Although no numerical diameters for the human NB fibrils were provided, the authors described the averaged filament density as being divided into two thinner individual filaments, resulting in an overall two-pronged cross-section. Indeed, when measuring the axis lengths of the presented cross-sections through the subtomogram averages, the values obtained from the shown images are about 8.9–10 × 2.7–3.6 nm. The theoretical arrangement with cross- sectional axis lengths of 8.8 × 3.2 nm would therefore be consistent with both the existing crosslinks between different TPR CC segments, and with the thickness of NB fibrils as inferred from the subtomogram averages. Such a TPR:TPR:MAD1 bundle could likewise be accommodated within the NB struts of 3T3 cells, whose diameter was reported to be about 10 nm (Singh *et al*., 2024).

However, we consider it noteworthy that, when we measured the individual NB fibrils shown as primary ET data for HEK293 cells, macrophages, and lymphocytes (Obarska- Kosinska *et al*., 2026), as well as for 3T3 cells (Singh *et al*., 2024), all of them appeared rather thin, with apparent widths ranging between about 2.4 and 3.7 nm. Taking into account the tomographic planes in which the nuclear pores are shown in these micrographs, it is evident that the fibrils are viewed from different angles. In fact, one can conclude that the images do not show only the narrow side of the fibrils, but should also include fibrils whose broader side is being presented. This raises the question why no broad fibril is apparent in the primary data, and why the greater width becomes apparent only in the subtomogram averages.

However, even though not all questions concerning the folding and arrangement of the TPR homodimers as part of the NB proper in human cells may yet be answered, the current data require us to further consider how the thin NB fibrils in *Xenopus* oocytes can be explained. If TPR homodimers in *Xenopus* oocytes were also folded severalfold to generate a CC1:CC4:CC5:CC6/7 bundle, whose diameter would be difficult to reconcile with the measured fibril widths, one would again have to consider a preparation artefact. This, however, would require an extreme compaction of the fibrils. Alternatively, oocyte NBs may indeed differ from mammalian somatic cell NBs with regard to the folding of their TPR homodimers. At present, we cannot exclude that in oocytes only CC4 is folded onto CC1, whereas further folding, resulting in a CC1:CC4:CC5:CC6/7 bundle, occurs only in other cell types.

### (8) The NB proper: with or without a terminal ring?

The question of where the CC5–8 segments of the TPR homodimers are positioned in different cell types leads directly to the third issue regarding the NB proper, namely whether the TR is a structure physiologically existing in certain cell types or merely another imaginable artefact of sample preparation in conventional EM.

The distal lateral connections between the eight NB fibrils were initially described as a distal ring (Jarnik and Aebi, 1991), whereas the term TR was introduced shortly thereafter (Panté and Aebi 1993). In the early studies in which a distal ring, aka a TR, was mentioned, this structure indeed appeared as a ring-like element. Later, higher-resolution studies indicated that this ring may instead correspond to a zig-zag-like lateral arrangement between neighbouring NB fibrils. In this context, it was proposed that the NB fibrils would bifurcate at their distal ends and that these branches would then interweave with neighbouring branches and thereby form the basket ring (Goldberg and Allen, 1996). More recently, the zig-zag-like arrangement seemingly interconnecting the NB fibrils laterally was proposed to be formed by the second half of TPR’s homodimeric rod domain (Gunkel *et al*., 2021).

However, while the existence of a TR long appeared almost unquestioned, lateral connections between the distal ends of the NB fibrils in murine and human somatic cells clearly were not detectable in the recent ET studies (Singh *et al*., 2024; Obarska-Kosinska *et al*., 2026), even though such connections could in principle be compatible with the XL-MS data for human TPR (see also Supplemental Figure S7E8 and S7E9): Provided that the homodimeric rod domain of TPR indeed folds severalfold, such that the CC1 and CC4–7 segments together form the rectilinear NB fibril, the two 24 nm-long CC8 segments of the two TPR homodimers within each fibril would be long enough to interact with the CC8 segments of neighbouring fibrils, if all CC8 segments branched laterally away from each fibril on both sides at an angle of 90°. Since the circumference of an octagon-shaped TR with a circumradius of 45 nm would be 275 nm, and even that of an octagon with a circumradius of 50 nm would only be 306 nm, the corresponding side lengths of about 34 and 38 nm could be bridged by one of the CC8 pairs between neighbouring fibrils, in principle even allowing for an anti-parallel alignment over 14 or 10 nm, respectively.

However, if a regular TR composed of CC8 existed in human cells, the question remains why it would not have been detected in the recent high-quality ET datasets. In addition, in none of the XL-MS studies were crosslinks detected between CC8 segments. Furthermore, CC8 is certainly not long enough to form the zig-zag-like arrangement between neighbouring fibrils seen in amphibian (Goldberg and Allen, 1996; Gunkel *et al*., 2021). Thus, if the NB fibril in *Xenopus* oocytes would also represent an aligned bundle of CC1:CC4:CC5:CC6/7, what then would we see in SEM and TEM as a structure with a rather regular, zig-zag-like appearance? A simple explanation would be that essentially all NBs in *Xenopus* oocytes would have NBLSs appended to them, of which the vast majority would simply collapse onto the NB proper during sample preparation. This potential problem, namely that NPC-associated fibrils could shrink, tear, and then collapse during EM sample processing, had already been mentioned in early NPC studies (Franke, 1970). The regular appearance could then be explained by assuming that the angle at which the NBLS fibrils collapse onto the NB is structurally predetermined.

However, at least two findings are difficult to reconcile with this scenario. First, the fibrils of the NB proper in *Xenopus* oocytes are very frequently seen to not only bifurcate at their distal ends but also to be interconnected with the bifurcating distal ends of the next-but-one NB fibrils in an interwoven manner (Goldberg and Allen, 1996; Gunkel *et al*., 2021), making it difficult to imagine how the fibrils could interweave so regularly during collapse. Second, lateral interconnections between fibrils of the NB and the NBLSs can be seen at regular intervals of about 45–50 nm in apparently intact structures comprising the NB proper and several NBLSs stacked onto it in *Xenopus* oocytes (Franke, 1974a; Scheer *et al*., 1988; Ris and Malecki, 1993), further arguing against such a collapse scenario as an explanation for the TR. Therefore, we currently do not rule out the possibility that a TR in vertebrate oocytes, and possibly also in other cell types, could be formed in a cell-type-specific manner by the CC5–7 parts of the TPR homodimers, when laterally branching off from the NB fibril scaffold formed by the aligned CC1:CC4 parts. Formation of the TR would then occur either as proposed previously, involving interactions between one fibril and the next-but-one fibril (Gunkel *et al*., 2021), or in a version involving directly neighbouring fibrils, which would also be compatible with the crosslinks detected between CC5 and CC6/7 of human TPR (Supplemental Figure S7E8 and S7E9). Although we do not exclude the existence of a yet unknown adaptor protein, the TPR polypeptides in the outlined scenarios could on their own suffice for forming a TR. Furthermore, since lateral interconnections also appear to exist between the fibrils forming the NBLS stacks in *Xenopus* oocytes (Scheer *et al*., 1988; Ris and Malecki, 1993), we can imagine that the TPR homodimers of the NBLS too may be capable of laterally interconnecting themselves via the distal parts of their NTDs, thereby also forming TR-like structures (Supplemental Figure S7E8 and S7E9).

The reason why the NBs, and likely also the NBLSs, in oocytes might form a TR may relate to the fact that, in the later stages of oogenesis, chromatin is no longer associated with the NE, in contrast to most somatic cell types. In the latter, a more compact version of the NB fibril may be required for specific interactions with neighbouring chromatin and, in proliferating cells, for replication-associated chromatin homeostasis at or near NPC-associated chromatin, a process linked to TPR (Zhang *et al*., 2016; Kosar *et al*., 2021) and its homolog Mlp1p (Bermejo *et al*., 2011; García-Benítez *et al*., 2017). In such cells, the formation of a thick NB fibril composed of bundled TPR CCs might provide the required stability for keeping the fibril lastingly appended to the NPC, possibly also by generating additional binding sites for lateral interactions with proteins that connect the NB and NBLSs to flanking chromatin, as discussed further below. By contrast, in cells lacking perinuclear chromatin, the thick version of an NB fibril as potential interaction platform for chromatin-binding proteins may not be required. Nonetheless, the NB fibrils in the oocyte might need to be stabilised too, to enable the NB to fulfil its tasks. In this scenario, such stabilisation would be provided by the cell-type-specific formation of a TR.

In other words, apart from a set of NB functions possibly common to all cell types, an NB in oocytes, or in other cells devoid of perinuclear chromatin, may not need to fulfil the same full range of tasks as in somatic cells, and thus may not require the particular NB structure that meets the demands in other cell types. Indeed, since oocytes store proteins for later use during embryogenesis, one could even imagine that many NBs, and their NBLS appendices, may even simply represent stockpiled material. Oocyte NBs may even engage in oocyte-specific interactions that involve lateral interactions between the CC5–8 segments of neighbouring NBs. Such interactions could possibly allow for the NB-associated lattices sometimes seen in the oocytes of different amphibian species (e.g., Goldberg and Allen, 1992; Cordes *et al*., 1997; Arlucea *et al*., 1998). It is even tempting to speculate that a CC5–8 segment that does not interact with any part of the same or another TPR homodimer, but instead remains pivotally tethered to the NB or NBLS fibril as a rod-like arm, may allow the two CTDs to extend away from the NB or NBLS fibril. In insect families that lack a ZC3HC1 homolog, and thus possibly also lack NBLSs, such hinge-mediated mobility of the second half of the NTD may even provide an alternative means of extending the spatial and functional reach of TPR.

However, before pursuing these speculations further, it is necessary to first clarify whether a TR may exist as a physiological arrangement in oocytes. One can then ask whether a TR occurs only in a cell-type-specific manner, or, in principle, even in all cell types, as a manifestation of different conformations, varying between NBs without a TR as the default situation and some with TRs, perhaps assembled in a regulated manner only on demand. In this context, we regard it noteworthy that, in the early descriptions of the fishtrap-like structure in mammalian cells, it was mentioned to exist both in an open and closed state, with the fishtrap fibrils described as appearing interconnected and forming a ring (Maul, 1976, 1977), which may represent NBs with and without a TR.

Either way, since all NBs in later-stage *Xenopus* oocytes appear to have a TR in EM specimens, either as an artefact or as a regular structure, ET of such oocytes, even though their vitrification is technically highly demanding due to their large size, may eventually contribute to distinguishing between these two possibilities. In addition, XL-MS data from *Xenopus* oocyte NEs might provide further insight. Neither strategy, however, makes it obsolete to raise an antibody that specifically recognises the CC5:CC6 transition, in order to pinpoint where this proxy for a severalfolded TPR homodimer is located, both in oocytes and mammalian cells. Crosslinks and structure prediction tools can provide valuable information, but since we cannot always tell for certain why and where a TPR folding event has occurred in cell samples exposed to conditions that may cause NB rearrangements, and since AlphaFold is not yet infallible, the classical approach of mapping a site of interest by old-school immunolocalisation might still turn out to provide the missing insight.

### (9) The NB, the NBLS and nucleoplasmic pools: Introducing the fields of employment

Even though several structural details of the NB proper and the NBLSs remain unresolved, our current data suggest that NB:NBLS ensembles are a common feature of HeLa cells and various other proliferating human cell types. Representing extensions of the NB proper into the nuclear interior, the ZC3HC1-dependent NBLSs would increase both the number and the spatial reach of NB-associated TPR polypeptides. This directly raises the question of which functions are fulfilled by such extended NB:NBLS ensembles. Do the NB proper and the NBLSs mainly act together as one supramolecular TPR:ZC3HC1 scaffold, or do TPR’s NPC-anchored T1 population and the ZC3HC1-dependent T2 population also have distinct, subpopulation- specific functional repertoires?

The functional rationale for ZC3HC1-dependent fibrillar TPR extensions of the NB proper that project into the nuclear interior of somatic cells appears straightforward. TPR is not merely an anchoring site for the established NB-resident proteins (see Introduction), but also appears to interact with additional proteins that associate with the NB only transiently and, in some cases, in a cell-type-specific manner. Beyond proteins involved in nucleocytoplasmic transport, mammalian TPR in interphase binds proteins involved in transcription regulation, RNP maturation, chromatin remodelling, DNA repair and diverse signal transduction pathways (e.g., Vomastek *et al*., 2008; Kosar *et al*., 2021; Chowdhury *et al*., 2026; our unpublished data), as will be discussed further below. Furthermore, in meiocytes, TPR appears to recruit S100PBP to the NE during early meiotic prophase, and because loss of S100PBP impairs XY crossover formation and increases autosomal crossover formation, there may be a link between NPC- or NB-anchored TPR, S100PBP and the regulation of meiotic recombination (Wu *et al*., 2025).

In view of these findings, NBLSs could expand the number of TPR binding sites available for such different processes, allowing the same proteins to engage TPR within both the NB proper and NBLS fibrils. Alternatively, PTMs or conformational features that distinguish NB from NBLS fibrils may permit interactions specific to distinct TPR subpopulations. More generally, we consider it possible that some proteins bind TPR only when it has adopted one particular conformation, but not when the same region of TPR is present in another structural context. Indeed, current evidence suggests that some proteins and protein complexes may preferentially use either the NB or NBLS as operational platforms, whereas other TPR-binding partners may engage both. However, it remains to be determined why some of these interactions preferentially involve the NPC-anchored T1 population, the ZC3HC1-dependent TPR subpopulations, or both, and whether others may cell-type-specifically involve TPR subpopulations even uncoupled from the NB, a possibility that will also be addressed further below.

Before considering the potential functions of the NBs and NBLSs in further detail, one first needs to once again point out though that TPR is essential in mice (https://www.mousephenotype.org/data/genes/MGI:1922066; our unpublished data) and *Drosophila* (Qi *et al*., 2004). While these genetic data do not formally prove that the NB as a structure is essential, they suggest that at least some NB-associated functions are indispensable in these animals. TPR even appears essential in at least some tumour cell lines like HeLa, as we could not produce a TPR KO cell line using CRISPR/Cas9n technology, in line with results for several other tumour cell lines (Hart *et al*., 2015; Aksenova *et al*., 2020). In other organisms, though, the TPR homologs are dispensable, with *Arabidopsis* tolerating KO of the TPR homolog NUA (Jacob *et al*., 2007; e.g., Xu *et al*., 2007) and *S. cerevisiae* being viable without Mlp1p and Mlp2p (Strambio-de-Castillia *et al*., 1999; Kosova *et al*., 2000).

ZC3HC1, on the other hand, is not strictly essential for viability even in mice (e.g., Illert *et al*., 2012; Gunkel *et al*., 2023; Aherrahrou *et al*., 2026; https://www.mousephenotype.org/data/genes/MGI:1916023; our unpublished data), and one can assume that NBLSs appended to the NB via ZC3HC1 are thus dispensable too for basic house-keeping functions. Pml39p in budding yeast is also non-essential (Palancade *et al*., 2005), and in some species, including most insect orders, ZC3HC1 homologs are naturally absent (Gunkel *et al*., 2023, see also further below). Consistent with this, NBLSs have not been detected in *Chironomus tentans* (Kiseleva *et al*., 1996, 1998; Soop *et al*., 2005), which belongs to the order Diptera, just like *Drosophila*.

Nevertheless, because a ZC3HC1 homolog exists in most animal, plant, fungal and amoebozoan species, it potentially provides some general advantage. Furthermore, since similar amounts of TPR occur appended to the NB via ZC3HC1 in various proliferating human cell types, including tumour and non-tumourigenic cell lines, primary cells, and human embryogenic stem cells (Gunkel *et al*., 2021; this study; our unpublished data), one can predict that NB-appended NBLSs are present in all these cell types. Assuming that this appendage of additional TPR to the NB extends the NB’s spatial reach, the question remains which repertoire of tasks would need to be fulfilled by the NB proper and the NBLSs.

### (10) The NB and NBLS: contemplating roles in perinuclear chromatin organisation

One morphologically evident function in which the NB and its NBLS extensions may act together is the formation of an NPC-associated chromatin exclusion zone (CEZ), and thus a chromatin-free “courtyard” in front of the NPC, as described previously for mammalian somatic cells (Krull *et al*., 2010). However, electron-microscopic images of such CEZs have to be interpreted with caution. As discussed in detail previously (Krull *et al*., 2010), one first needs to point out that an NPC-associated CEZ extending deep into the nuclear interior does not necessarily reflect the presence of a protein scaffold along the CEZ’s entire length: Chromosomes in the interphase nuclei of somatic cells are non-randomly arranged, with each chromosome occupying its own spatially restricted chromosome territory (ChT; e.g., Cremer *et al*., 1993). Although these ChTs are not strictly separated from each other by rigid boundaries, but can partly intermingle with neighbouring chromatin (Branco and Pombo 2006) through interchromosomal contacts in a transcription-, compartment- and cell-type-dependent manner (Mokhtaridoost *et al*., 2024), chromatin-poor clefts can nonetheless occur between neighbouring ChTs. Such clefts would primarily arise from the spatial arrangement and self- organisation of the ChTs themselves and could be maintained by the dynamic events at their peripheries, without requiring a dedicated, chromatin-excluding protein scaffold along the entire ChT perimeter (Krull *et al*., 2010). Accordingly, not every part of a chromatin-free zone seen in conventional TEM images of ultrathin sections of somatic cells would need to reflect the presence of TPR and ZC3HC1. Thus, in cases in which neighbouring ChTs are positioned at the NE, those parts of an NPC-associated chromatin-free zone that, in a TEM micrograph, appear to extend particularly deep into the nuclear interior could in principle - if such possibility is not excluded by serial sections - depict a sectioned interterritorial cleft instead of a scaffold- defined tubular channel (Krull *et al*., 2010).

This explanation, however, does not apply to the tubular, NPC-associated proximal parts of those CEZs that extend into chromatin-rich regions of a ChT, where they are surrounded by chromatin on all sides, and whose existence depends on the presence of TPR (Krull *et al*., 2010). For these CEZs, the more plausible interpretation remains that NPC-associated scaffolds exclude chromatin from the NPC entrance. In mammalian cells, in which ZC3HC1 appends additional TPR to the NB, such scaffold-defined CEZs would then comprise not only the NB proper but also the ZC3HC1-dependent NBLSs. In this view, the NB-attached NBLSs would thus extend the NPC-associated CEZs deeper into the nuclear interior, either directly into a ChT or into the cleft between neighbouring ChTs. Stacks of NBLSs would then correspond to the up to ∼200 nm long NPC-associated and TPR-containing filament bundles found in human tissues (Cordes *et al*., 1997) and to the similarly long NPC-associated CEZs observed in various somatic cell types (Krull *et al*., 2010 and references therein). The NBLSs would thus be the structural correlate of extended CEZ cylinders that emanate from NPCs into the nucleus.

However, we suspect that the flanking chromatin usually does not directly contact the fibrillar TPR-containing scaffold of the NB and NBLSs. Previously, it had been concluded that a very narrow, additional chromatin-free zone between the actual NB scaffold and the flanking chromatin keeps them separated from each other (Krull *et al*., 2010). In addition, in high- pressure frozen previtellogenic *Xenopus* oocytes, in which chromatin is still located near the NE, we noted an NE-aligned zone filled with fibrillar material between the NE and chromatin (Gunkel *et al*., 2021), which appears continuous with the narrow chromatin-free zone surrounding the NB scaffold (Figure 7F). This zone may correspond to a zone that aligns the NE and the NBs in the salivary gland cells of *Chironomus tentans* and excludes rRNPs upon shutdown of nuclear export (Kylberg *et al*., 2010). While the NE-aligning zone may be established by unstructured parts of transmembrane NE proteins, we consider unstructured material associated with the NB, and in vertebrates also with the NBLS, to contribute to the thin layer between the NB:NBLS scaffold and the neighbouring chromatin (Figure 7F). Candidates include the large unstructured loop of ZC3HC1 (Gunkel *et al*., 2023), parts of TPR’s CTDs that may not be entirely confined to the scaffold’s interior, and natively unfolded regions of nucleic acid-binding proteins that we found interacting with TPR. In this view, the region surrounding the NB would not only contribute to maintaining the NPC courtyard as chromatin-free but also to forming an interface with neighbouring chromatin that could be either transcriptionally inactive or active (see below), and to nuclear export of certain types of cargos (see below).

### (11) The NB and NBLS: pondering roles in transcription regulation

Although the Mlps are non-essential in budding yeast, one role assigned to the NB in this organism is transcription regulation, as subsets of transcriptionally active genes can be recruited to or be associated with NPC-anchored Mlp proteins (e.g., Casolari *et al*., 2004, 2005; Dieppois *et al*., 2006; Luthra *et al*., 2007; Tan-Wong *et al*., 2009; Bermejo *et al*., 2011; Texari *et al*., 2013). Furthermore, TPR and its homologs also across other species participate in transcription-regulatory interactions with chromatin or transcription factors (e.g., Mendjan *et al*., 2006; Skaggs *et al*., 2007; Vaquerizas *et al*., 2010; Su *et al*., 2018; Boumendil *et al*., 2019; Aleman *et al*., 2021; Uhlířová *et al*., 2021; Tyagi *et al*., 2023; Bartlett *et al*., 2024; Bednor *et al*., 2026; Chowdhury *et al*., 2026; Liu *et al*., 2026). In addition, elimination of TPR or its homologs results in altered transcriptomes (e.g., Aksenova *et al*., 2020; Lee *et al*., 2020; Zuckerman *et al*., 2020; Aleman *et al*., 2021; Kamel *et al*., 2024; Liu *et al*., 2026), which also holds for NPCs with and without NBs (Bensidoun *et al*., 2022), suggesting that the NB may regulate the expression of subsets of genes by enabling or maintaining their silenced or active states.

We consider it possible that the NBLSs in species with ZC3HC1 may execute similar processes. In this scenario, NBLSs extending deeper into the nuclear interior would allow for more lateral interactions with surrounding chromatin and engagement with genes beyond the reach of the NBs alone (Figure 7F). In line with this notion, a recent study in fission yeast has shown that both the CTD of Nup211, the homolog of TPR/Mlp1p, and Rsm1, the homolog of ZC3HC1/Pml39p, contribute to 3′-processing and transcription termination of gene-regulatory non-coding RNAs (Bednor *et al*., 2026). Strikingly, a rsm1 mutation that abolishes the second BLD module caused gene-regulatory phenotypes similar to those of Nup211 mutants with truncated versions of the CTD (Bednor *et al*., 2026). Moreover, Nup211 and Rsm1 mutations affected overlapping transcript classes, and their combination did not produce a pronounced synergistic phenotype, suggesting that both proteins contribute to the same gene-regulatory pathway (Bednor *et al*., 2026). Since the BLD-truncating Rsm1 mutation corresponds to ZC3HC1/Pml39p truncations that abolish NB association and recruitment of additional TPR/Mlp1 polypeptides (Gunkel *et al*., 2023), the Rsm1 phenotype could provide an example of a Rsm1/ZC3HC1/Pml39p-dependent subpopulation of Nup211/TPR/Mlp1p that selectively contributes to the regulation of specific gene subsets.

Further along this line, also mammalian ZC3HC1, and thus the TPR subpopulations forming the NBLS, appear required for the regulated expression of genes that are distinct from those affected by the NB: Comparing the transcriptomes of tumour and non-tumour WT and ZC3HC1 KO cells, we noted altered expression of distinct classes of genes in the absence of ZC3HC1, yet with the affected gene classes differing between tumour and non-tumour cells, indicating cell-type-specific up- and downregulation.

Further support for cell-type-specific functions of the ZC3HC1-dependent TPR subpopulations comes from their differential physicochemical stability. In certain non-tumour cell lines, in which the pool of ZC3HC1-appended TPR polypeptides is of a similar size as in HeLa cells, the T2 subpopulations of TPR cannot be released from the NBs under conditions that readily release and solubilise the T2 subpopulations in other cell lines, including most but not all tumour cell lines tested (our unpublished data). This suggests that cell-type-specific PTMs or additional binding partners, including chromatin-associated proteins, may stabilise the T2 population in some cell lines and thereby contribute to cell-type-specific NBLS functions.

Furthermore, upon RNAi of ZC3HC1, TPR, and the TREX2 protein GANP, we observed similarities between the transcriptomes of TPR KD and GANP KD cells, in line with previous findings (Aksenova *et al*., 2020), and these transcriptomes differed markedly from those following ZC3HC1 KD, and thus from cells without NBLSs (our unpublished data). Similarly, a recent study revealed striking differences between the transcriptomes of TPR- and ZC3HC1- deficient IMR-90 cells (Bartlett *et al*., 2025). While this study described ZC3HC1 as not appending additional TPR to the NBs, we found that NE-appendage of about half of all NE- associated TPR in IMR-90 cells depends on ZC3HC1 (Figure 4A; Supplemental Figure S4A). Together, we interpret these findings as indicative that ZC3HC1-appended TPR populations can engage with genes out of reach of TPR’s NPC-anchored T1 subpopulation. Furthermore, when interacting with transcriptionally active genes, the NB:NBLS ensembles may define different types of translocation pathways, allowing nascent mRNPs to be funneled directly into these fibrillar arrangements or routed along their outside towards the NPC (Figure 7F).

### (12) The NB and NBLS: questioning roles in quality control of intron-containing RNAs

Mlp1p/Mlp2p and Pml39p have been proposed to contribute to a quality-control step that prevents incompletely spliced mRNAs from reaching the cytoplasm. This conclusion was initially drawn from reporter-based assays indicating that loss of Mlp1p allows intron- containing transcripts to escape nuclear retention, without detectably impairing splicing itself (Galy *et al*., 2004). This concept was later extended to Pml39p, whereas the NPC-associated endoribonuclease Swt1p was implicated more broadly in nuclear mRNP quality control (Palancade *et al*., 2005; Skruzný *et al*., 2009; Bonnet *et al*., 2015). More recently, age- dependent displacement of the NB in yeast was also connected to leakage of unspliced pre- mRNAs, again suggesting that this pathway may become relevant when the NB structure is compromised (Mirkovic *et al*., 2025).

In the context of the current study, we regard the findings reported for Pml39p of particular relevance. In *mlp1*Δ and *pml39*Δ single-deletion strains, as well as in *mlp1*Δ *pml39*Δ double- deletion strain, leakage of an intron-containing reporter transcript was found to occur to a comparable extent, with no evidence for a relevant additive effect (Palancade *et al*., 2005; Bonnet *et al*., 2015). Since loss of Pml39p does not affect NPC anchorage of Mlp1p but prevents the recruitment of additional Mlp1 polypeptides to the NB (Bensidoun *et al*., 2022; Gunkel *et al*., 2023; Zsok *et al*., 2024; Schmidt *et al*., 2026), this would, within the framework of our current model, point to the Pml39p-recruited Mlp1 polypeptides as the Mlp1p subpopulation involved in preventing export of incompletely spliced transcripts.

In human cells, TPR has likewise been implicated in the control of intron-containing RNAs. Two studies using HIV-based reporter systems in primate cell lines concluded that depletion of TPR increases export of intron-containing transcripts (Coyle *et al*., 2011; Rajanala and Nandicoori, 2012). In addition, studies on endogenous human transcripts linked TPR to nuclear retention and/or exosome-dependent decay of incompletely spliced RNAs (Yap *et al*., 2012; Fish *et al*., 2019).

However, it remains necessary to critically assess whether mammalian TPR has a general central role in the surveillance of intron-containing transcripts, since TPR was found not to be required for nuclear retention in reporter systems that were both similar to and distinct from those used earlier (Iino, 2017; Lee *et al*., 2020). Furthermore, genome-wide transcriptome analyses after TPR depletion in U2-OS and MCF7 cells did not reveal a significant increase in the cytoplasmic fraction of intron-containing transcripts, suggesting that TPR is not required for nuclear retention of the bulk of unspliced mRNAs (Lee *et al*., 2020; Zuckerman *et al*., 2020). Consistent with this, genome-wide transcriptome analyses in HeLa cells revealed that neither loss of TPR nor loss of ZC3HC1 caused an increase in the total amount of intron- containing RNAs in the cytoplasm (our unpublished data). This suggests that the ZC3HC1-dependent TPR subpopulations, i.e. the NBLSs, are also not required for retaining the bulk of incompletely spliced mRNAs. However, since these findings were all obtained with tumour cell lines, ongoing research still needs to clarify to what extent this also applies to different types of non-tumour cells.

At present, however, it seems difficult to reconcile the phenotypes of ZC3HC1 knockout mice with a scenario in which only the ZC3HC1-dependent TPR subpopulations, i.e. the NBLSs, would play a fundamental role in mRNA quality control. Apart from moderate impairments already present in young animals, old ZC3HC1 knockout mice eventually appear to be, on average, in markedly better general health than their wild-type and heterozygous littermates (our unpublished data). Why this is the case remains to be clarified, but impairment of any crucial mRNA quality-control mechanism does not appear to be the most plausible explanation.

### (13) The NB and NBLS: sketching roles in cargo processing and translocation

Numerous studies have linked the TPR in mammalian cells or its homologs in other species to nuclear export of RNAs, including bulk or selected poly(A)+ mRNAs, stress-induced cellular mRNAs, viral mRNAs, lncRNAs, circRNAs, and tRNAs (e.g., Bangs *et al*., 1998; Shibata *et al*., 2002; Galy *et al*., 2004; Vinciguerra *et al*., 2005; Jacob *et al*., 2007; Skaggs *et al*., 2007; Xu *et al*., 2007; Bae *et al*., 2009; Aksenova *et al*., 2020; Lee *et al*., 2020; Zuckerman *et al*., 2020; Chen *et al*., 2021; Li *et al*., 2021; Bensidoun *et al*., 2022; Bhat *et al*., 2023; Cao *et al*., 2024; Vdovina *et al*., 2025; Zhou *et al*., 2026). In addition, NPCs with and without NBs in budding yeast were found to associate with different classes of mRNAs, indicating NB-specific RNA interactions (Bensidoun *et al*., 2022). Furthermore, various proteins involved in RNA processing, RNP maturation, and nuclear export have been identified as binding partners of TPR, its homologs, or the NB in experiments determining the protein interactome of TPR or its homologs by different means (e.g., Green *et al*., 2003; Niepel *et al*., 2005; Bretes *et al*., 2014; Dimitrova *et al*., 2015; Saroufim *et al*., 2015; Hakhverdyan *et al*., 2021; Kosar *et al*., 2021; Malik *et al*., 2023; De Castro *et al*., 2024; our unpublished data), or by comparing the protein inventory of NPCs with and without NBs in yeast, human tumour cells, and *Xenopus* oocytes (Aksenova *et al*., 2020; Gunkel *et al*., 2021; Bensidoun *et al*., 2022; our unpublished data).

However, apart from TPR being probably indirectly involved in the export of naturally intron-less or intron-poor mRNAs, by acting as a binding platform for the TREX-2 complex (e.g., Umlauf *et al*., 2013; Wickramasinghe *et al*., 2014; Aksenova *et al*., 2020; Gunkel and Cordes, 2022; Bhat *et al*., 2023; Vdovina *et al*., 2025), it remains largely unknown how TPR contributes to the export of other cargoes.

In this context, it is also important to distinguish between NB-associated proteins whose stable NE association may require the ZC3HC1-dependent TPR subpopulations and those that interact preferentially with the NB proper. For example, in mammalian cells, loss of ZC3HC1, and thus of the ZC3HC1-dependent TPR subpopulations, does not detectably affect NE association of GANP, SENP1 or MAD1 (Gunkel and Cordes, 2022). Likewise, NE localisation of the corresponding yeast proteins Sac3p, Ulp1p and Mad1p does not appear to be affected by the absence of Pml39p (Gunkel *et al*., 2023). This suggests that the NB proper can provide a specific interaction platform for some proteins that only bind to TPR within the T1 subpopulation. Such specific interactions may even help prevent ZC3HC1 from engaging the ZBD2s of T1 polypeptides, as exemplified for MAD1, for which AlphaFold predicts that its N-terminal α-helix masks the ZBD2 of a T1 homodimer (Supplemental Figure S8B3). Conversely, the NBLSs may provide additional or distinct interaction sites for other binding partners, including proteins whose association with TPR may even be more transient or cell- type-specific.

Furthermore, little is known about which transport routes towards the NPC exist for the different types of nuclear export cargos (e.g., Williams *et al*., 2018; Kehlenbach and Chook, 2025). Some models propose translocations exclusively through the NB’s TR while other findings point towards additional pathways that bypass the TR and instead enter or exit the NB between its fibrils (e.g., Iborra *et al*., 2000; Li *et al*., 2021).

We can envision a scenario in which the NB and NBLSs enable passage of export cargos along at least two distinct paths. In one, the bulk of cargos would be granted access to a central route through the scaffold-surrounded lumen filled with most parts of TPR’s CTDs. Cargos permitted access to this region would enter it either through the NB’s or NBLSs’ distal end or laterally between the NBLS-forming fibrils. Other export cargos not granted access to the central route would make use of a path along the outside of the NB:NBLS scaffold and only enter the scaffold-surrounded area, then laterally through the NB fibrils, close to the NPC, where we regard TPR’s CTDs to be largely absent (Figure 7F).

Furthermore, we consider it likely that most nuclear import cargos, once released from their nuclear import complexes, will not traverse the CTD meshwork but will instead exit the NB laterally near the NPC (Figure 7F). This view is consistent with earlier findings showing that TPR can interact with one of the nuclear import factors, Importin-β, itself, but not with cargo- loaded Importin-β (Shah *et al*., 1998).

Several CTD features are relevant in this context of selective cargo translocations. First, the CTDs of TPR homologs in vertebrates and other multicellular organisms have several FG repeats, even though fewer than in some distant species. Unlike Mlp1 of budding yeast, which lacks an FG repeat in its CTD, the CTDs of TPR homologs in other unicellular organisms, such as, e.g., the microalga *Nannochloropsis gaditana* (EWM27281), share similarities, including many FG repeats, with the unstructured FG-rich domains of nucleoporins. Such similarities could indicate that the TPR CTDs may have originated from a prototypic FG-repeat domain. Furthermore, some parts of the CTDs in vertebrates and other species have an aa composition similar to that of FG-repeat domains (Kuznetsov *et al*., 2002), and CTD polypeptides lacking the central acidic cluster (Figure 7A) have CD spectra (Hase *et al*., 2001) that resemble those of FG-repeat domains (Denning *et al*., 2002, 2003). Since the TPR CTDs amount, on average, to about 4.74 MDa per NPC in HeLa, one might envision that a meshwork composed of CTDs could exhibit sieve-like properties and act as a selective permeability barrier, similar to the barrier formed by FG-repeat domains within the NPC (Schmidt and Görlich, 2015). However, if NB:NBLS scaffold dimensions are about 70–80 nm in diameter and 80–120 nm in height (Krull *et al*., 2010; Gunkel *et al*., 2021; Figure 5) and if the CTDs were evenly distributed within this volume, their mean density would only be ∼13–26 mg/ml. These values are far below the 100–400 mg/ml reported for condensates of other intrinsically disordered proteins, including the NPC’s FG domains (e.g., Schmidt and Görlich, 2015; Abyzov *et al*., 2022; Ng *et al*., 2023). They would, though, be in the range of some disordered proteins forming low density, liquid-like condensates (e.g., Wei *et al*., 2017). Considering a mean of 62 CTD chains of 733 aa uniformly distributed within such cylindrical lumens, and assuming that inter-chain contacts (π–π, cation–π, or related) can occur on average between every 10 th, 15th, or 20th residue along each chain, this could generate a statistically isotropic 3D network with ∼1100– 2,100 nodes. For such node densities and cylinder volumes, the resulting mean spacing between the contact points, corresponding to the average mesh size of a CTD meshwork, would be ∼5– 8 nm.

However, the CTD may not be entirely disordered: CD spectra of CTD polypeptides indicate some β-sheet fractions in solution (Hase *et al*., 2001), and while AlphaFold3 predicts single CTD to be largely unstructured, it suggests β-sheet formation when several CTDs are considered together (Supplemental Figure S7A2). Therefore, since four CTDs, from two TPR homodimers, may cluster in each asymmetric NB unit, we wonder whether this could lead to stretches of β-sheets and modulate the presumed CTD network properties.

In addition, we had noted a short ordered segment within the CTD, predicted by PONDR (Predictor of Natural Disordered Regions; Li *et al*., 1999 Supplemental Figure S7B). This structured segment is now predicted by AlphaFold3 as an α-helix (Supplemental Figure S7A1), and it appears to be involved in interactions with nucleic acid-binding proteins and regulated by phosphorylation (our unpublished data). Furthermore, also numerous other CTD residues appear to be phosphorylated in response to extracellular stimuli (Figure 7A; www.phosphosite.org; e.g., Eblen *et al*., 2003; Olsen *et al*., 2006; Vomastek *et al*., 2008; Pan *et al*., 2009; Rajanala *et al*., 2014), indicating that the CTD’s functions can be modulated by signaling pathways during interphase.

Another conspicuous feature is a short cluster of RG and RGG motifs close to the CT of the CTD in vertebrates and other multicellular organisms (Kuznetsov *et al*., 2002), and such an RGG-rich region close to the CT also exists in Mlp1 of *Chaetonium thermophilum* (Dimitrova *et al*., 2015). These motifs could be involved in a wide range of processes (e.g., Thandapani *et al*., 2013; Chong *et al*., 2018; Chowdhury and Jin, 2023), including remodeling and maturation of mRNPs or phase separation between the NB:NBLS and the surrounding nucleoplasm. Several of these RG/RGG motifs also serve as targets for arginine methylation (Guo *et al*., 2014; Larsen *et al*., 2016).

The perhaps most striking CTD feature in animals is the acidic cluster in the middle of the CTD (Figure 7A; Kuznetsov *et al*., 2002). Such clustered acidic residues could contribute to unpacking and charge-based remodelling of mRNPs within the CTD lumen. A process of mRNP unpacking is exemplified by the Balbiani ring particles in the salivary gland cells of *Chironomus tentans*. These globular mRNPs of ∼45 nm in diameter dock to the TR, appear to undergo rearrangement within the NB lumen, and traverse the NPC as elongated, baton-like particles (Stevens and Swift, 1966; Kiseleva *et al*., 1996; Daneholt, 1997; Soop *et al*., 2005). Given that globular pre-mRNPs in avian erythroblasts measure 35–40 nm in diameter (Maxwell *et al*., 1981), that TREX-coated pre-mRNPs in human nuclei are similarly roundish and span 30–70 nm (Pacheco-Fiallos *et al*., 2023), and that human cytoplasmic mRNPs are ellipsoidal and distinctly smaller, with 35–50 nm in length but only 9–10 nm in width (Skabkin, 2004), it is reasonable to propose that similar remodelling principles apply to vertebrate mRNPs above a certain size and that such mRNPs may need to engage with the CTDs to become export-competent. These mRNPs would enter the CTD meshwork from the distal end of the NB:NBLS scaffold and possibly also laterally, to be matured and directed towards the NPC (Figure 7F). Extremely long transcripts, however, like the one for titin, which exceeds the length of the BR transcript by far, and whose estimated mRNP size may reach or exceed 70 nm in diameter, need to exit the nucleus via the NE budding pathway instead (Zaganelli *et al*., 2026).

In yeast, by contrast, isolated nuclear mRNPs are far smaller and already ellipsoidal, measuring 15–35 nm in length and 5–7 nm in width (Batisse *et al*., 2009). Such dimensions suggest that most yeast mRNPs may not require remodelling as extensive as in vertebrates to acquire export competence, a possibility that may correlate with the fact that the CTDs of Mlp1p and Mlp2p lack several of the sequence features characteristic of CTDs in other species (Kuznetsov *et al*., 2002). Nevertheless, the CTD of Mlp1p in *S. cerevisiae* has been shown to serve as a docking site for hnRNPs and other proteins involved in mRNA export (e.g., Green *et al*., 2003; Fasken *et al*., 2008; Dimitrova *et al*., 2015).

Apart from this, we can envisage a “fast track” to the NPC for those RNPs in animals that do not require extensive remodelling, including mRNPs from naturally intronless genes and others from immediate early response or heat-shock genes that must exit the nucleus rapidly under stress situations. Such RNPs might even be denied access to the CTD-filled area and instead move along the outer perimeter of the NB:NBLS scaffold (Figure 7F). There they could use the previously suggested 10–20 nm chromatin-free corridor between the NB fibrils and surrounding chromatin (Krull *et al*., 2010), potentially maintained by space holders between NB and chromatin proposed decades ago (Maul, 1971). After bypassing the CTD-filled region, these mRNPs would enter the NB lumen laterally between the NB fibrils, close to the NPC (Figure 7F), consistent with some earlier models (Iborra *et al*., 2000; Li *et al*., 2021).

Furthermore, confinement of a large acidic mass within the NB:NBLS-enclosed volume should also disfavor passage of basic proteins that have been imported into the nucleus, such as the histones. We therefore predict that most nuclear import complexes are disassembled directly after having passed the NPC. In this scenario, the released import cargoes, which in principle would mostly be small enough to enter the CTD-filled channel and would therefore need to be prevented from interacting with the CTDs’ acidic clusters, would enter the nucleoplasm by passing sideways between the NB fibrils close to the NR (Figure 7F).

Altogether, we can envisage a scenario in which the NB and NBLS define distinct transport pathways for different classes of cargo, either through the CTD meshwork or bypassing it (Figure 7F). In this model, the CTD-filled lumen would function as a selective translocation channel for the bulk of RNPs where they are matured for export and directed towards the NPC. While the NB is capable of handling such RNPs on its own, as in most insects and in ZC3HC1- deficient mice, the NBLSs would reduce the workload of the NB proper by providing access to this pathway over a longer distance. To what extent the CTD’s RGG motifs, FG repeats, short α-helix, acidic cluster and perhaps locally formed beta-strands contribute to such RNP maturation and other processes remains a topic of ongoing investigation.

### (14) NB and NBLS: distinguishing default assembly from local demand

Having discussed possible roles of mammalian NBLSs in chromatin organisation, transcription regulation, RNA quality control and cargo translocation, we finally consider whether such extensions are assembled by default or in response to local demand. This question is prompted by *S. cerevisiae*, where NB assembly is not a constitutive feature of all NPCs but has been linked to local transcription- and mRNP-dependent processes.

In budding yeast, only a subset of NPCs is equipped with Mlps, and thus NBs (e.g., Galy *et al*., 2004; Niepel *et al*., 2005, 2013; Bensidoun *et al*., 2022; Singh *et al*., 2024; Zsok *et al*., 2024; Schmidt *et al*., 2026). Cryo-ET resolved a stable, morphologically recognisable NB only on about 27% of all NPCs investigated, corresponding to a minority population of NPCs with a double nuclear Y-complex ring (Singh *et al*., 2024). Consistently, live-cell data indicated that Mlp1p, Mlp2p and Pml39p are present only on a subset of NPCs, whereas Nup60p, the NPC- anchored recruitment factor for Mlp1p (e.g., Stankunas and Köhler, 2024), is described as being found on all NPCs (Zsok *et al*., 2024). In fact, NB-less NPCs have been proposed to represent the default state, with appendage of Mlp1p, Mlp2p and Pml39p to NPCs, and thus NB formation or maintenance, depending on ongoing RNA Pol II transcription and mRNP maturation (Bensidoun *et al*., 2022). Consistently, inhibition of RNA Pol II or depletion of certain 3′-end-processing/polyadenylation factors, such as Pab1p, caused loss of Mlp1p from NPCs and its redistribution into the nucleoplasm or into nuclear granules (Bensidoun *et al*., 2022). Furthermore, FRAP (fluorescence recovery after photobleaching) experiments had shown a near-complete recovery of Mlp1p/Mlp2p fluorescence at the NE within a few seconds, and thus much faster than the recovery of GFP-tagged proteins of the NPC proper (Niepel *et al*., 2013). On the other hand, however, doRITE (dilution of RITE-labelled complexes) analyses revealed that an NPC-associated population of Mlp1p, Mlp2p and Pml39p can remain stably associated with individual NPCs over multiple cell divisions (Zsok *et al*., 2024). Thus, rapid exchange or redistribution of NE-associated Mlp fluorescence may coexist with a stably appended pool of the Mlp polypeptides at individual NPCs. Furthermore, upon certain natural stimuli, such as heat shock, Mlp1p, Mlp2p and Pml39p lose their steady-state presence at the NE and accumulate within intranuclear foci instead (Carmody *et al*., 2010; Bensidoun *et al*., 2022; Mohajan *et al*., 2025), and at least the NPC-uncoupled Mlp1p then appears capable of executing functions also at sites distant from the NE (Mohajan *et al*., 2025).

By contrast, in the vertebrate cell types examined so far, essentially all NPCs appear to be equipped with an NB proper. This conclusion can be drawn from, for example, SEM overview images of *Xenopus* oocyte NEs (e.g., Jarnik and Aebi, 1991; Kiseleva *et al*., 1996), TEM images of poliovirus-infected HeLa cells, showing NPC-associated HEZs with NB-like dimensions at essentially every cross-sectioned NPC (Krull *et al*., 2010), and FM images of various cell types showing co-localisations of TPR and ZC3HC1 with NPC proteins at essentially all NPCs (Gunkel *et al*., 2021). Furthermore, since about half of the total cellular amount of TPR is attached to the NE via ZC3HC1 in a wide range of human tumour cell lines and proliferating non-tumour cells (Gunkel *et al*., 2021; Gunkel and Cordes, 2022; current study and our unpublished data), we also consider it likely that the NBs in these cells at least are commonly equipped with NBLSs too. The few sporadically detectable NPCs in HeLa and other proliferating cell types that appear to lack TPR may be explained by incomplete labelling or by NB assembly not yet having been initiated. In fact, only very few NPC-containing vertebrate cell types appear to lack NBs, as appears to hold for mature spermatids. While TPR still appears NPC-associated across the entire NE in pachytene spermatocytes (Kuznetsov *et al*., 2002; Wu *et al*., 2025), TPR staining at the NE becomes concentrated in a single cluster during the early haploid stages of spermatogenesis (Kuznetsov *et al*., 2002), which correlates with NPC clustering in only one region of the NE in these stages (e.g., Friend and Fawcett, 1974; Fawcett and Chemes, 1979; Sutovsky *et al*., 2000; Ho, 2010). Then, in mature spermatids, NPC-associated TPR is no longer detectable (Kuznetsov *et al*., 2002; Santos *et al*., 2024), which correlates with the remaining spermatid NPCs apparently lacking a recognisable CEZ and/or NB (e.g., Franklin, 1968; Santos *et al*., 2024).

Several observations further indicate that assembly or maintenance of the NB proper in vertebrates neither requires the presence of genes in the immediate neighbourhood of an NPC nor ongoing transcriptional activity. First, in the late stages of *Xenopus* oogenesis, NPC and NB assembly continues in the absence of neighbouring chromatin. From stage IV onward, the chromosomes detach from the NE and become located deep within the nuclear interior, while growth of the giant oocyte nucleus and its NE continues, along with a further increase in NPC numbers. Since essentially all NPCs of stage VI oocytes are equipped with NBs, one can conclude that assembly of the new NPCs after chromosome withdrawal is accompanied by assembly of new NBs. Thus, at least in this cell type, NB assembly can occur even though any chromosomal genes that are still transcriptionally active are already located micrometers away from these NPCs, deep within a nucleus of approximately 400 µm in diameter. Second, even in amphibian erythrocytes, characterised by highly condensed chromatin and hardly any transcriptional activity (e.g., Grasso and Woodard, 1966; Small and Davies, 1970; Maclean *et al*., 1973; Krohne *et al*., 1981), almost all NPCs are equipped with TPR and ZC3HC1 (Gunkel *et al*., 2021). This indicates that, once assembled, vertebrate NBs can also be maintained in cell types with very low transcriptional activity.

This conclusion is further supported by perturbation experiments. After a four-hour transcriptional shutdown in HeLa cells, NPC association of both TPR and ZC3HC1 remains largely unaffected, with no notable detachment of either protein from the NE into the nucleoplasm (Gunkel *et al*., 2021), despite ZC3HC1 then being hyperphosphorylated. As an aside, the latter finding also demonstrates that certain ZC3HC1 residues can be phosphorylated without detectably disrupting the NB-associated ZC3HC1:TPR ensembles (Gunkel *et al*., 2021). Furthermore, neither knockdown of PABPN1 nor PABPC1, the latter being a human homolog of yeast Pab1p, causes any notable detachment of TPR or ZC3HC1 from the NPCs (Supplemental Figure S8D), unlike what has been found in yeast (Bensidoun *et al*., 2022).

In addition, FRAP experiments in HeLa cells provide no evidence for any highly dynamic exchange of NE-associated TPR, but rather support the notion that the TPR subpopulations remain durably associated with the NPCs and NBs (Supplemental Figure S2D4; our unpublished data). Moreover, neither TPR nor ZC3HC1 is notably detached from the NPCs in HeLa cells upon heat shock (Supplemental Figure S8E). Finally, when isolated NBs of ZC3HC1 KO cells are first incubated with cell extracts containing recombinant ZC3HC1, subsequently added recombinant TPR is stably appended to such NBs (Gunkel and Cordes, 2022), indicating that the formation of higher-order TPR:ZC3HC1:TPR ensembles can readily occur even in vitro.

In view of these findings, we conclude that NPCs constitutively equipped with NBs represent the default state in mammalian cells, and that transcriptional activity in general appears neither required for NB assembly nor for maintaining its integrity. This, in turn, suggests that the mammalian NB proper should be regarded as a constitutive structural component of the NPC, rather than an optional appendage whose assembly depends on ongoing transcriptional activity in the neighbourhood of an NPC. Furthermore, since neither TPR nor ZC3HC1 has so far shown the kind of large-scale stress-dependent detachment from the NPCs reported for Mlp1p and Pml39p (Carmody *et al*., 2010; Bensidoun *et al*., 2022; Mohajan *et al*., 2025), it is, in principle, conceivable that even the ensemble of NPC, NB and NBLS in proliferating mammalian cells may represent a de facto constitutive entity.

At present, we do not know which features distinguish the yeast NB, as a structure assembled or maintained only on demand, from an NPC-anchored mammalian NB assembled by default as part of a cognate NPC-NB entity. However, while the mammalian NB may be an essential structure that has to remain anchored to essentially every NPC, in order to act as an operational platform for other NB-resident proteins or to fulfil its own specific tasks, the NBLS may not need to be appended to every NB in every mammalian cell type, also because ZC3HC1 is not an essential protein and the NBLS may thus not be an essential structure either. In fact, we do not exclude that some mammalian cell types exist that commonly lack NB-associated NBLSs, and that other cell types may generally have several NBLSs appended to their NBs, resulting in longer NB extensions.

On the one hand, one could imagine that such variability between cell types merely reflects a matter of supply. In this simple scenario, TPR and ZC3HC1 would be alternatingly appended to all NBs in a largely stochastic manner as long as sufficient amounts of both proteins were available. Indications for such a scenario come, for example, from in vivo recruitment experiments in which endogenous soluble nucleoplasmic TPR in ZC3HC1 KO cells appears to become rather uniformly appended to pre-existing NBs upon induced expression of recombinant ZC3HC1 (Gunkel and Cordes, 2022). In addition, the rather uniform distribution patterns of NPC-associated TPR and ZC3HC1 in HeLa WT cells (Figures 5A and 5C) would also be compatible with stochastic recruitment of both proteins to the NBs until one of them becomes limiting.

Therefore, in this simple scenario, the amounts of both proteins in some cell types would suffice to form NB-appended cylinders, whereas in other cell types, a low amount of one protein would prevent the other, even if sufficiently abundant, from being further appended to the NB. One could thus speculate that an excess of TPR over ZC3HC1 may be the reason for why TPR in certain differentiating cell types also occurs in a nucleoplasmic state not physically linked to NPCs (Uhlířová *et al*., 2021; Malik *et al*., 2023; Chowdhury *et al*., 2026). Similarly, analyses of tissues and organs from newborn mice may point in the same direction. In these animals, the rather balanced relationship between TPR and ZC3HC1 seen in numerous cell lines appears not to exist in all tissues at P0 and P1 (our unpublished data). In fact, certain tissues in newborn mice appear to have a soluble nucleoplasmic surplus of either TPR or ZC3HC1. By contrast, hardly any soluble TPR and ZC3HC1 appear to exist in the corresponding tissues of healthy adult animals, with the relative amounts of both proteins instead appearing rebalanced again (our unpublished data). In principle, it is therefore conceivable that the number of NBLSs appended to an NB may be determined simply by transcriptional up- or downregulation of either *TPR*, *ZC3HC1* or both genes. In this context, one can also imagine that disease-associated dysregulation of either gene’s transcription could disturb cell-type- or differentiation-stage-specific TPR:ZC3HC1 copy-number relationships during postnatal development and in adult tissues, possibly contributing to tumourigenesis or other pathological processes.

However, a scenario in which the amounts of TPR:ZC3HC1 ensembles would only correlate with transcriptional activities at the *TPR* and *ZC3HC1* genes, followed by stochastic NBLS assembly as long as supplies are available, would be too simple, for several reasons. First, it would not consider that the nucleoplasmic TPR pool in NSPCs differs from the NPC-associated TPR pool by at least one phosphorylation event (Malik *et al*., 2023). This finding suggests that regulatory processes at the protein level may also play a role in whether TPR is bound to pre- existing NBs or NBLSs, or retained in a soluble nucleoplasmic state, in order to fulfil some functions not linked to the NBs. Second, a merely stochastic assembly of NBLSs would not explain how differently long NB extensions at neighbouring NPCs can come about in the nuclei of adult tissues. In fact, in numerous TEM micrographs of ultrathin sections from vertebrate tissues with non-dividing, terminally differentiated cells, NPC-associated HEZs can be seen whose lengths differ notably within the same nucleus, with short and long cylinders sometimes located close to each other within the same nuclear region. This in turn suggests a mechanism of local length regulation, indicating that NBLS extensions may also be assembled because of a local demand.

In fact, while assembly and maintenance of the vertebrate NB do not require neighbouring chromatin or ongoing transcription, and while this may also apply to at least the first NBLS appended to an NB (Gunkel and Cordes, 2022), we do not exclude that distinct chromatin loci may provide yet unknown stimuli that initiate assembly of additional NBLSs onto pre-existing NB/NBLS ensembles. In budding yeast, such a local demand may be transient and may affect different individual NPCs at different times, without requiring long-term commitment of a particular NPC to an extended NB ensemble. In a non-dividing heterochromatin-rich somatic cell, however, one may expect that such a local demand for a “length-adjusted” version of the NB extensions would have to be specified early during terminal differentiation, before the heterochromatin-rich regions at the nuclear periphery are fully established. This would also require both early formation of the NB extensions and early selection of those NPCs from which they would need to extend into CTs.

Indeed, it is difficult to imagine that the long NBLS cylinders seen in heterochromatic CTs are assembled only after an external signal, such as heat shock, has triggered the need for such structures. In the case of stress-induced transcriptional activation of distinct loci, which would require an immediate response, one would not expect the cell to wait until ZC3HC1 and TPR had been upregulated or at least redistributed within the nucleus in order to only then assemble long NBLSs and locally displace chromatin for connecting such loci with the nuclear periphery. If such long NB extensions and the chromatin regions that they link to the nuclear periphery might play roles in stress or immune responses, it appears more plausible that such NBLS stacks are formed in advance, perhaps soon after the last cell division, in order to be already available when a rapid cellular response to an external event has to be implemented.

Assuming that transcriptionally active genes located deeper within the nucleus may contribute to creating a local demand for long NB extensions, it may be less difficult to imagine how those HEZs that extend into clefts between neighbouring CTs come about. In this scenario, formation of the NBLS stacks could be directly related to their engagement with subsets of still transcriptionally active genes exposed at CT surfaces (e.g., Kurz *et al*., 1996; Verschure *et al*., 1999; Mahy *et al*., 2002; Chaumeil *et al*., 2006; Clemson *et al*., 2006). The more challenging questions concern those long, tubular HEZs that extend deep into a CT that, by staining, appears to be almost entirely heterochromatic. While we consider it possible that the basic construction principle for such particularly long NB extensions does not differ from that leading to the NB:NBLS ensembles in proliferating cells, their local formation, positioning and maintenance inside a CT may require additional determinants. In addition to TPR and ZC3HC1, other TPR-interacting proteins may contribute to keeping TPR positioned along the perimeter of such CEZs. These could include proteins that laterally connect the TPR scaffold to neighbouring chromatin, but also yet another architectural element, such as an additional cross-linker of TPR polypeptides, involved in assembling or stabilising certain extended TPR arrangements. We also cannot rule out that TPR homodimers located deeper within the nuclear interior acquire features that allow them to engage in sufficiently stable oligomeric interactions, perhaps in staggered arrangements, thereby permitting thin but extended TPR ensembles without requiring an additional cross-linking protein at every interface.

Whatever the precise structural implementation may be, the central issue remains what, in such an intraterritorial position, would need to be connected to an NPC-associated NBLS scaffold. Could such intraterritorial sites indeed represent loci that must remain accessible even in terminally differentiated cell types, including different types of leukocytes, in which the density of NPCs per unit NE surface area can be far lower and the heterochromatin content far higher than in proliferating cells? Would such loci need to be connected to NPCs in many or all cell types, or could individual NPCs establish cell-type-specific connections to distinct loci that must remain accessible only in particular terminally differentiated cells? Could such loci include transcriptionally competent islands within an otherwise compact and transcriptionally repressive CT? Could the NBLS stacks linking such loci to individual NPCs provide cargo translocation routes? Could the NBLS scaffold components at such loci also act as an operational platform required by distinct proteins executing their tasks at such loci? In other words, could these NB extensions help keep a discrete intraterritorial site transcriptionally competent and functionally connected to an NPC?

Although it remains elusive how the specific formation and stabilisation of a particularly long NBLS extension at one NB but not another within the same nucleus is achieved, we can imagine that the stimuli initiating the formation of such extensions, not only those between CTs but also those reaching into CTs, may originate from gene-regulatory processes at specific gene loci.

One conceivable class of intraterritorial sites that might induce or require the formation of NB extensions is represented by pericentromeric satellite loci. These loci are embedded in constitutive heterochromatin, but are not transcriptionally inert under all conditions. In mammals, pericentromeric satellite transcripts have been linked to the formation and organisation of chromocentres, to the establishment or maintenance of pericentromeric heterochromatin, and to defined stages in development and differentiation in which chromatin architecture is reorganised. This suggests that a pervasive demand for such pericentromeric satellite transcripts may already exist shortly after the last cell division. In addition, though, pericentromeric satellite transcription can also be strongly induced by cellular stress, including heat shock, and emerging data may link pericentromeric satellite/nuclear-stress-body pathways to inflammatory or antiviral responses (e.g., Saksouk *et al*., 2015; Brändle *et al*., 2022; Ninomiya *et al*., 2025).

For example, upon heat shock, human HSATIII RNAs are produced from pericentromeric satellite III regions and act as architectural lncRNAs in nuclear stress bodies (e.g., Jolly *et al*., 2004; Metz *et al*., 2004; Rizzi *et al*., 2004). Recent work further suggests that, during recovery from thermal stress, a fraction of these RNAs is exported, in an ALYREF-promoted manner, to the cytoplasm, where they contribute to recovery-associated processes (Ninomiya *et al*., 2025).

However, whether NB extensions reach out to these or other perichromatic regions, or to any other distinct gene loci, remains a topic for future research. At present, we can only assume that the complexity of interactions among TPR homodimers, ZC3HC1 and other proteins that may induce the formation of HEZs extending deep into the nuclear interior is far greater than we can currently infer.

Moreover, even though we regard the basic ZC3HC1-mediated NBLS assembly principle to hold for all NBLS-containing cell types, including not only proliferating and non- proliferating somatic cells but also the full-grown oocytes of *Xenopus* and other vertebrates, such oocytes raise yet further questions about the mechanisms underlying site-specific NBLS assembly. In fact, in the late stages of oogenesis, the lampbrush chromosomes undergo condensation and become clustered deep within the nuclear interior, leaving stage V and VI ooyctes devoid of perinuclear chromatin and thus eliminating neighbouring chromatin as potential source of stimuli for local NBLS assembly. Nonetheless, particularly long NBLS stacks, which, as mentioned above, are more frequently observed in stage VI than in stage V oocytes, appear to be distributed rather non-uniformly across the NE. Some NE regions feature numerous NBs with long stacks of NBLSs appended to them, whereas other regions feature NBs that appear to be devoid of NBLSs. Without any neighboring chromatin, the question remains as to which stimuli might induce the site-restricted assembly of particularly long NBLSs or prevent their assembly at other sites within this huge nucleus of more than 400 µm in diameter.

### (15) Nucleoplasmic TPR and ZC3HC1: functional pools uncoupled from the NPC?

Having so far focused on TPR and ZC3HC1 as scaffold components, we consider it important to also point at other findings of particular note, which indicate that subpopulations of TPR in different species may execute certain functions also when not physically linked to the NPC. First, in *Drosophila melanogaster*, which lacks an obvious ZC3HC1 homolog (Gunkel *et al*., 2023), notable amounts of the TPR homolog Megator/Mtor in some cell types (e.g., Zimowska *et al*., 1997; Zimowska and Paddy, 2002; Qi *et al*., 2004), but not necessarily in others (e.g., Lye *et al*., 2014), appear to exist not only at the NE but also at sites deep within the nuclear interior. Such subpopulations were detected, for example, in cells during different stages of larval development, where the nuclear Megator distribution was described as a discontinuous granular or particulate pattern without any obvious connectivity to NPCs (Zimowska and Paddy, 2002). Of particular note, upon heat shock of *Drosophila* larvae, most of this nuclear Megator was found to rapidly accumulate at the 93D heat shock locus (Zimowska and Paddy, 2002), which does not encode heat shock proteins but is transcribed into non-coding heat-shock RNAs and acts as a nuclear repository for several RNA-binding and pre-mRNA-processing factors during heat shock (e.g., Dangli *et al*., 1983; Prasanth *et al*., 2000). By contrast, positioning of the NPC-associated Megator subpopulation appeared to remain unaffected upon heat shock (Zimowska and Paddy, 2002). Provided that the particulate staining patterns within the nucleus, consistently observed with different Megator antibodies (Zimowska and Paddy, 2002; Qi *et al*., 2004), do not reflect unrelated antibody cross-reactions, these findings indicate that *Drosophila* TPR can exist in different structural states and may behave differently in response to different cellular stimuli, as previously proposed (Zimowska and Paddy, 2002). Furthermore, chromatin-IP experiments with *Drosophila* SL-2 cells revealed an association of Megator with actively transcribed chromosomal regions that, according to FISH analyses, were not restricted to the NE-proximal nuclear periphery but also occupied positions deeper within the nuclear interior, suggesting that both peripheral and intranuclear pools of Megator may engage in chromatin binding (Vaquerizas *et al*., 2010).

Furthermore, and as already mentioned above, Mlp1p and Mlp2p in budding yeast have been found to redistribute from the NE to nucleoplasmic foci upon heat shock (Carmody *et al*., 2010), and Mlp1p no longer tethered to the NPCs is required for organising heat-shock- response genes into chromatin clusters located deeper in the nucleoplasm (Mohajan *et al*., 2025). In addition, a nucleoplasmic pool of TPR has been reported in murine C2C12 myoblasts, where TPR binds chromatin at specific genes and contributes to transcriptional regulation during myogenesis (Uhlířová *et al*., 2021). Furthermore, a large nucleoplasmic pool of phosphorylated TPR (p-TPR), often associated with heterochromatin located deeper within the nucleus, was described for murine neural stem/precursor cells (NSPCs). This nucleoplasmic pool of p-TPR was no longer present in mature neurons, in which TPR was located primarily at the NE (Malik *et al*., 2023). MS analysis of TPR immunoprecipitates from NSPC lysates revealed that several transcription factors, chromatin modifiers and other chromatin-associated proteins bound specifically to either TPR or p-TPR, suggesting different roles in chromatin organisation and transcription regulation (Malik *et al*., 2023). Interestingly, some of these chromatin-associated proteins were also detected as specific TPR interactors in systematic Y2H screens of human cDNA libraries using segments of TPR’s NTD and CTD as baits (our unpublished data). Moreover, a nucleoplasmic pool of TPR was recently described for murine E16.5 fetal gonocytes (Chowdhury *et al*., 2026). In these cells, TPR was found to specifically interact with SPOCD1, a chromatin-associated piRNA-pathway scaffold factor, thereby helping to keep LINE1 loci accessible to the piRNA-directed methylation machinery (Chowdhury *et al*., 2026). This function was proposed to operate throughout the nucleoplasm, again raising the possibility that some TPR-dependent chromatin-regulatory functions need not be restricted to NPC-associated TPR (Chowdhury *et al*., 2026).

On the one hand, we wonder whether some of the murine TPR described as nucleoplasmic, based on microscopic images, may nonetheless be associated with NBs via ZC3HC1. We regard it as noteworthy that disruption of the SPOCD1–TPR interaction in gonocytes causes male sterility, and that male infertility is also one of the main phenotypes of ZC3HC1/NIPA- deficient mice (Illert *et al*., 2012; our unpublished data; https://www.mousephenotype.org/data/genes/MGI:1916023). Even though such a shared phenotype does not prove a common mechanism, it is tempting to speculate that the loss of ZC3HC1-dependent TPR extensions may also affect normal SPOCD1–TPR interactions to some extent.

On the other hand, though, we do not doubt the existence of notable amounts of nucleoplasmic TPR in certain developmental stages, tissues and species, including mammals, even though the soluble nucleoplasmic pools of *Hs*TPR, in more than 20 different human tumour and non-tumour cell lines, primary cells and stem cells tested so far are small, generally below 5% of TPR’s total cellular content (Hase *et al*., 2001; Gunkel *et al*., 2021; Gunkel and Cordes, 2022; our unpublished data). However, comparing corresponding tissues from newborn and adult mice, we noted, as mentioned above, that some P0 and P1 tissues appear to have a significant pool of soluble nucleoplasmic TPR, unlike in the adult tissues. This finding suggests that TPR may primarily perform NPC/NB-associated functions in the adult, yet may also play roles as an NB-uncoupled nucleoplasmic protein during differentiation processes (our unpublished data).

However, it remains to be clarified how NPC-associated TPR and TPR not physically linked to an NB contribute to the transcriptional regulation of different sets of genes, and how TPR- interacting factors involved in such regulatory processes distinguish between a mobile, nucleoplasmic version of TPR and TPR that is part of either an NB or NBLS.

In view of the different interactomes reported for p-TPR and TPR in NSPCs (Malik *et al*., 2023), it is conceivable that the NPC-anchored and nucleoplasmic versions of TPR might differ in their post-translational modification profiles. In addition, however, we also consider it possible that differences in the conformational states of NB-associated TPR and of a soluble, nucleoplasmic version could play a role. In this context, we regard it as notable that the SPOCD1-binding site on TPR has been mapped to TPR residues 775–877 (Chowdhury *et al*., 2026). This region lies within CC4, and thus in a CC segment that, together with CC1, forms the main scaffold of the NB fibril. On the one hand, this could exemplify lateral interactions between TPR fibrils of NBs or NBLSs with neighbouring chromatin. On the other, it raises the question of whether SPOCD1 recognises CC4 only in a specific conformational state. One can speculate whether the SPOCD1-binding site would be accessible in a multiply folded TPR homodimer as part of an NB fibril or in a perhaps less folded TPR homodimer as part of an NBLS fibril, or whether the SPOCD1-binding site is only exposed in an unfolded version of a nucleoplasmic TPR homodimer. In principle, similar considerations may also apply to a stress- regulated interaction between TPR and the transcription factor HSF1 identified in vitro (Skaggs *et al*., 2007). In this case, HSF1 was found to bind TPR close to its NT and to a TPR segment comprising aa 1218–1320 (Skaggs *et al*., 2007), and thus part of TPR’s CC6. If such an interaction were confirmed in vivo, one would again need to distinguish whether it reflects a lateral, HSF1-mediated interaction of NB or NBLS fibrils with neighbouring chromatin, or an interaction of HSF1 with nucleoplasmic TPR homodimers that are not linked to the NPC.

Further along this line, however, we also regard it as noteworthy that nucleoplasmic TPR in ZC3HC1 KO cells remains soluble, without engaging in any noticeable interactions with chromatin (Gunkel and Cordes, 2022). By contrast, upon detachment of ZC3HC1-appended TPR from the NBs in WT cells by degron-mediated rapid degradation of ZC3HC1, such TPR distributes throughout the nuclear interior, where it then is no longer extractable under conditions that readily extract nucleoplasmic TPR from ZC3HC1 KO cells (Gunkel and Cordes, 2022). One can now speculate whether TPR homodimers that had initially been part of NBLS fibrils may have acquired features, such as PTMs or a distinct conformation, that allowed for certain interactions with chromatin. Upon degron-mediated release of such TPR homodimers from the NBs, these features might then enable them to interact with chromatin sites that are normally out of the reach of an NBLS extension. By contrast, without such features, the nucleoplasmic TPR homodimers in ZC3HC1 KO cells would be kept from binding to inappropriate chromatin regions. However, it is also conceivable that in other, non-tumour cell types, and in particular during differentiation processes, the nucleoplasmic subpopulations of TPR may exhibit pool-specific features that allow them to be specifically attracted, when required, to distinct chromosome sites far out of the reach of the NBs and their NBLSs.

In summary, TPR and ZC3HC1 appear to occur in distinct subpopulations across cell types, with both overlapping and subpopulation-specific functions. The abundance of these subpopulations, as well as the copy number relationships between TPR and ZC3HC1, likely require tight control, particularly during differentiation, to ensure appropriate regulation of gene expression and other processes in different cell types. We therefore consider it likely that dysregulation of these copy number relationships, for example in disease or during aging, could contribute to a range of pathologies.

### (16) ZC3HC1: outlining a proposal for a new name

Finally, we wish to end by proposing that ZC3HC1, with its aliases HSPC216, NIPA and ILP1, be renamed. The original naming of ZC3HC1 (zinc finger C3HC-type protein 1; https://www.genenames.org/data/gene-symbol-report/#!/hgnc_id/HGNC:29913) was based on its annotated Pfam zf-C3HC motif (Finn *et al*., 2006). However, ZC3HC1 and its homologs actually do not possess an evolutionarily conserved C3HC motif but a C2HC zinc finger instead, exemplified by Pml39p, which lacks one of the cysteines eponymic for the C3 in the motif name (Gunkel *et al*., 2023).

The alias HSPC216 stemmed from its cDNA naming, with this cDNA being one among numerous ones isolated from hematopoietic stem/progenitor cells and with HSPC216 not having been further characterised (Zhang *et al*., 2000). The alias NIPA (nuclear interacting partner of ALK) was chosen after its cDNA had been isolated in a Y2H screen with a chimeric bait that included the receptor tyrosine kinase ALK (Ouyang *et al*., 2003). However, this interaction did not reflect an interaction between ZC3HC1 and ALK but instead between ZC3HC1 and an N-terminal segment of TPR that was part of the chimeric bait (Bischof *et al*., 1997; Gunkel *et al*., 2021) and that included TPR’s ZBD1 (Gunkel *et al*., 2021 and current study). ILP1 (inhibitor of apoptosis-like protein [IAP]-like protein 1), on the other hand, was chosen because parts of its sequence are similar to that of IAP proteins (Higashi *et al*., 2005), with such similarity indeed evident (Gunkel *et al*., 2023). However, while ILP1 (Higashi *et al*., 2005), respectively NIPA (Ouyang *et al*., 2003; Illert *et al*., 2012) had been portrayed as a protein with an anti-apoptotic function, other lines of experimental evidence have opposed this notion (Gunkel *et al*., 2021, and references therein). Finally, while *S. cerevisiae* Pml39p may play some role in the nuclear retention of unspliced mRNAs (Palancade *et al*., 2005; Bonnet *et al*., 2015), current evidence does not indicate that ZC3HC1 is required for a similar surveillance mechanism in mammals.

The proposal for a renaming of ZC3HC1 is instead grounded on evidence pointing (i) at a protein that is primarily located at the NB in different cell types across vertebrates (Gunkel *et al*., 2021) (ii) at a protein that functions as a structural element and linker between the TPR polypeptides forming the scaffold of the NBs and the NBLSs (Gunkel *et al*., 2021; Gunkel and Cordes, 2022 and current study), (iii) at a protein whose NuBaID can be described by a signature that also encompasses ZC3HC1 homologues, like Pml39p, that have not been detectable with the zf-C3HC signature (Gunkel *et al*., 2023), and (iv) at a protein whose function, in terms of interconnecting TPR subpopulations, appears to be evolutionarily conserved, at least in mammals and yeasts (Gunkel *et al*., 2023). In addition, “ZC3HC1” is cumbersome to pronounce and tends to slow down scientific discussion when referring to this protein, so we suggest that any new name should be easier to use in spoken communication.

We therefore propose to call the vertebrate ZC3HC1 protein from now on **FIBE**, as an abbreviation for a **f**ibril-**i**nterconnecting **b**asket **e**xtender that, together with its corresponding TPR homologue, allows for a unique type of protein ensemble at the NB.

## Materials and Methods Y2H experiments

Y2H analyses were performed essentially as described previously (Gunkel *et al*., 2023). In brief, haploid MATa and MATα *S. cerevisiae* strains (Supplemental Table S1) were transformed separately with bait and prey constructs (Supplemental Table S2), respectively. Empty bait and prey vectors served as controls for self-activation. After selection on −Leu or −Trp plates, single colonies were expanded in liquid culture, mated in 96-well plates, and plated on adenine-containing YPDA agar. After two days at 25°C, cells were replica-plated onto −Leu/−Trp (−LW) plates to verify mating success and, after a further three days, onto−Leu/−Trp/−His (−LWH) plates without or with increasing concentrations of 3-AT. Plates were then incubated at 25°C for up to 10 days to monitor growth indicative of bait–prey interaction.

### Cell culture and cell cycle synchronisation

HeLa cells and IMR-90 cells were routinely cultured in high-glucose DMEM (D6429; Sigma- Aldrich) supplemented with 10% fetal bovine serum (FBS; P40-37500; PAN-Biotech) and 1% Antibiotic Antimycotic Solution (AAS; A5955; Sigma-Aldrich). All cell lines used in this study are listed in Supplemental Table S3. Most cell lines were commonly passaged twice weekly and maintained at 37°C under humidified air with 5% CO2. During generation of new cell lines and after cell sorting, the medium was occasionally supplemented with 100 µg/ml Normocin (InvivoGen). All cell lines were routinely tested for mycoplasma and other microbial contaminations and confirmed to be free of these. Where required, cell cycle synchronisation was performed by double-thymidine block, essentially as described previously (Gunkel *et al*., 2021). For increased adherence, for example in experiments requiring permeabilisation before fixation, cells were occasionally seeded on surfaces coated with Laminin/Poly-D-Lysine (LPDL001; Sigma-Aldrich) or bovine collagen (#125-50; Cell Applications).

### Genomic editing in human cell lines and cell line characterisation

Gene tagging in HeLa cells with sfGFP was achieved by CRISPR/Cas9 nickase technology, essentially as described previously (Gunkel and Cordes, 2022). In brief, sgRNA sequences (Supplemental Table S4) were cloned into the bicistronic Cas9n expression vector pSpCas9n(BB)-2A-Puro (PX462 V2.0; Ran *et al*., 2013), kindly provided by Feng Zhang (Broad Institute, Cambridge, MA). In the specific case of the sfGFP-tagged TPR Δ1–58 cell line, four sgRNAs had to be used, of which two targeted exon 1 and the other two exon 2, while the donor plasmid (see Supplemental Table S2) carried homology sequences of exon 1 (left arm) and 2 (right arm). The cells were then transfected with the corresponding sgRNA/Cas9 vectors together with the donor plasmid and sorted for GFP signals using a Bio-Rad S3e cell sorter (Bio-Rad Laboratories, Hercules, CA). In most cases, such fluorescence activated cell sorting (FACS) was repeated to obtain cell populations enriched for cells with particularly high fluorescence, which then served as the starting material for the selection of monoclonal cell lines. Following examination by LCI to assess proper localisation of the tagged proteins at the NE/NPC, the enriched populations were seeded at low density in 10-cm dishes to allow growth of monoclonal colonies that could be manually picked and transferred into 18-well μ-slides with ibiTreat bottom (ibidi). Clones were further screened by LCI to again assess subcellular localisation and compare NE-associated brightness under identical microscope settings. The brightest NE-positive clones were selected for further expansion and characterised by comparative IFM and immunoblotting versus the corresponding progenitor cell line, as well as by genotyping via PCR and DNA sequencing (see Supplemental Table S5), with all these procedures performed as described previously (Gunkel *et al*., 2021; Gunkel and Cordes, 2022).

ZC3HC1 KO cell clones (HeLa Kyoto and IMR-90) were enriched by treating the transfected population with puromycin at a final concentration of 5 µg/mL for 12 h. While the finite lifespan of IMR-90 did not allow for further selection of monoclonal cell lines, the HeLa Kyoto populations of puromycin-surviving cells were subjected to serial dilutions to obtain monoclonal cell lines that were further screened via IFM, as described in detail previously (Gunkel *et al*., 2021).

#### Transfections and RNAi

Transfections for transient expression in HeLa cells were performed essentially as described previously (Gunkel *et al*., 2021, 2023; Gunkel and Cordes, 2022). In brief, cells were seeded either on coverslips for IFM or on four- or eight-well glass-bottom μ-slides (ibidi) for LCI. The next day, they were transfected with PolyJet (SignaGen) or FuGene HD (Promega) according to the manufacturer’s instructions. Transfected cells were either fixed for IFM or live-imaged 24 hours post-transfection. Transfection and induced expression of the all-in-one vectors encoding a constitutively expressed Tet-On transactivator and either a doxycycline-inducible WT or mutant version of mCherry-tagged *Hs*ZC3HC1, as presented in Figure 3E, were performed as described previously (Gunkel and Cordes, 2022). For gene-editing transfections in HeLa cells, cells were grown to about 70% confluency in 10-cm dishes and then transfected with donor plasmids and the corresponding sgRNA/Cas9n vectors using PolyJet or FuGene HD, followed by enrichment of edited cells by antibiotic selection or FACS. For *ZC3HC1* gene disruption in IMR-90 cells, we performed nucleofection using a 4D-Nucleofector (Lonza, Basel, Switzerland) according to the manufacturer’s instructions, including the IMR-90 protocol provided by Lonza. RNAi of HeLa cells was performed essentially as described previously (Gunkel *et al*., 2021; Gunkel and Cordes, 2022). In brief, cells were seeded at low density on coverslips and transfected the next day in Opti-MEM (GIBCO) with 12 nM of either negative control siRNAs or target-specific siRNAs (see Supplemental Table S6), using HiPerFect transfection reagent (QIAGEN) according to the manufacturer’s instructions. Cells were fixed 72 h after transfection. For RNAi in IMR-90 cells, cells were seeded on coverslips and transfected with either 25 nM (DharmaFECT protocol) or 12 nM (HiPerfect protocol) of either negative control siRNAs or siRNAs targeting *Hs*ZC3HC1 (see Supplemental Table S6), using either DharmacFECT (Dharmacon, Lafayette, CO, USA) or HiPerfect according to the manufacturer’s instructions. The Dharmacon siRNAs against *Hs*ZC3HC1 (L-016879-02) represented a pre-designed pool of four different siRNAs. For transfections with Ambion siRNAs against *Hs*ZC3HC1 performed in parallel, two siRNAs (s28276 and s28278) had been pooled. The transfections with siRNAs were repeated on day three after exchanging the culture medium. Cells were fixed six days after the first transfection and immediately subjected to immunolabelling for IFM.

### LCI and photobleaching

Live-cell imaging was performed essentially as described previously (Gunkel *et al*., 2021, 2023; Gunkel and Cordes, 2022). In brief, cells were grown in FluoroBrite DMEM (GIBCO) supplemented with FBS and AAS on multi-well ibidi glass-bottom μ-slides (#1.5H). DNA staining of living cells was achieved with 5-SiR-Hoechst (Bucevičius *et al*., 2019). Images were acquired with a Leica TCS SP5 or SP8 confocal laser-scanning microscope (Leica Microsystems). In cells with endogenously sfGFP-tagged polypeptides, the DNA stain was used to position the focal plane at the nuclear equator before image acquisition, thereby avoiding bleaching of the GFP signal prior to the actual acquisition, which was particularly important for quantifications. For all NE quantification datasets, image acquisitions were performed with the SP8 microscope. Nucleoplasmic photobleaching of living cells transiently expressing GFP-tagged polypeptides was carried out with the Leica TCS SP5 as described previously (Gunkel *et al*., 2023), with selected areas being pulsed with full laser power for 2 min, followed by acquisition of post-bleach images.

### Antibodies

All antibodies used in this study and their specifications are listed in Supplemental Table S7. The novel *Hs*TPR antibodies presented in Figure 2F and 2G were raised in rabbits. Following immunisation with a recombinant *Hs*TPR fragment (aa 354–452) synthesised in bacteria and purified therefrom, subpopulations of peptide-specific antibodies were isolated by affinity chromatography on series of sequence-overlapping peptides (from Peptide Specialties GmbH, Heidelberg, Germany), as recently described for collections of *Xl*TPR peptide antibodies (Gunkel *et al*., 2021). Synthesis of sequence-overlapping peptides corresponding to the N- terminal extension of the non-canonical version of hsTPR (TPR-L, Figure S1E), and the coupling of these peptides via either an N- or C-terminal cysteine residue to maleimide- activated keyhole limpet hemocyanin (KLH) was done at Peptide Specialty Laboratories (Heidelberg, Germany). Immunisation of rabbits and subsequent affinity chromatography of the peptide-specific antibodies used for Figure S1G was performed essentially as described previously (e.g., Cordes *et al*., 1997; Krull *et al*., 2004).

### IFM

Specimens for standard IFM were prepared essentially as described previously (Gunkel *et al*., 2021). In brief, cells grown on #1.5 coverslips (Gerhard Menzel B.V. & Co. KG, Braunschweig, Germany) were fixed with 2.4% formaldehyde (FA), freshly prepared in PBS, and then quenched with 50 mM NH_4_Cl in PBS. Permeabilisation with 0.25–0.5% Triton X-100 in PBS was carried out either for 5 min after fixation or, where indicated, for 2.5 min before fixation, in the latter case in PBS supplemented with freshly added 10 mM MgCl_2_ to avoid destabilisation of the NB. Blocking and incubations with primary and secondary antibodies (Supplemental Table S7) were performed in PBS with 1% BSA. DNA staining with PureBlu Hoechst 33342 (Bio-Rad Laboratories) was carried out during incubation with fluorophore- coupled secondary antibodies. The immunolabelled specimens were mounted in SlowFade Gold or Diamond Antifade Mountant (Invitrogen), the latter to better preserve FA-fixed GFP signals. Specimens were inspected with a Leica TCS SP5 or SP8 confocal laser-scanning microscope (CLSM; Leica Microsystems) equipped with a HC PL APO 63×/1.40 OIL CS2 objective (Leica Microsystems). The colour lookup table (LUT) C3_linear-protanopic- deuteranopic was obtained from https://sites.imagej.net/NeuroCyto-LUTs.

### Immuno-SEM

Immuno-SEM of manually isolated NEs from *Xenopus laevis* oocytes was performed as described in detail previously (Gunkel *et al*., 2021). All *Xenopus* husbandry and handling complied with German law and were approved by the Veterinary Institute of LAVES (Niedersächsisches Landesamt für Verbraucherschutz und Lebensmittelsicherheit; 33.42502- 05/A-005/07). In brief, oocytes were obtained by ovariectomy from anaesthetised animals and transferred into modified Barth’s saline. Nuclei were manually isolated in 5:1-H buffer supplemented with 5 mM MgCl_2_, transferred onto 5 mm² silicon chips, opened with tweezers and thereby spread onto the silicon chips, and rinsed to remove the nuclear contents. NEs were fixed with 2%FA in 5:1-H containing 5 mM MgCl_2_ for 10 min, quenched with 50 mM NH_4_Cl in 5:1-H for 5 min, and incubated in blocking solution containing 1% cold-water fish skin gelatine (#G7041; Sigma-Aldrich) in 5:1-H for at least 30 min. Incubation with primary antibodies (Supplemental Table 4) was carried out in blocking solution for 60 min, followed by several washes and subsequent incubation in blocking solution containing goat anti-rabbit F(ab’)2 fragments coupled to 10 nm colloidal gold (#810.166; AURION, Wageningen, Netherlands) for 60 min. The immunogold-labelled NEs were rinsed in 5:1-H, followed by post-fixation first with 2% FA in 5:1-H for 20 min and then with 0.5% glutaraldehyde (GA) and 0.1% tannic acid in 5:1-H overnight. The NEs were rinsed with water, then further fixed with 0.5% OsO_4_ in water for 20 min, followed by rinsing, subsequent staining with 0.1% uranyl acetate for 10 min, dehydration in a series of increasing ethanol concentrations, and critical point-drying via CO_2_. Mounted specimens were examined without further metal coating at 10– 15 kV in a Hitachi S-5500 field-emission in-lens scanning electron microscope (Hitachi High- Technologies), via concurrent secondary electron (SE) and backscattered electron (BSE) detection. The superimposition of the BSE signals onto SE micrographs, the plotting of the positions of the immunogold particles (IGPs) onto the scheme of an idealistic NPC and its NB in a face-on-view drawn to scale (Supplemental Figure S2B), and IGP distance measurements (Supplemental Figure S2A–S2C) were performed as described in detail previously (Gunkel *et al*., 2021).

### Immunoblotting

SDS-PAGE and immunoblotting were performed essentially as described previously (Gunkel *et al*., 2021). Following incubation with primary and HRP-conjugated secondary antibodies (Supplemental Table S7), the immunolabelled proteins were detected by enhanced chemiluminescence (ECL) using ECL Prime (Amersham/GE Healthcare), followed either by exposure to fluorographic X-ray films (Super RX-N UV, Fujifilm, Tokyo, Japan) or by detection with an Azure 280 chemiluminescent imaging system (Azure Biosystems, Sierra Court, CA, USA).

### FACS and AFC

FACS was performed on a Bio-Rad S3e cell sorter (Bio-Rad Laboratories) and used, in sorting mode, for enrichment of sfGFP-positive cells in the course of establishing cell lines with endogenously tagged proteins and, in scanning mode, for quantification of the established cell lines’ sfGFP fluorescence intensities by analytical flow cytometry. Adherent cells were detached with Trypsin-EDTA (GIBCO), and trypsinisation was stopped after 3–5 min by addition of Defined Trypsin Inhibitor (GIBCO). After centrifugation at 300× g for 1 min, the fluffy cell pellet was resuspended in a suitable buffer (PBS or NB-stabilising buffer; Gunkel *et al*., 2021), filtered through a 35 µm cell strainer, and transferred into 5 mL tubes with cell- strainer caps (Falcon, Corning, NY, USA) which were then loaded into the sorter. For AFC corresponding to Figure 4I and Supplemental Figure S4E3, the cells had been permeabilised with 0.25 % TX-100 in NB-stabilising buffer for 3 min, followed by sedimentation of the cells by centrifugation at 300 × g, quantitative removal of the TX-100-containing buffer and its replacement with TX-100-free NB-stabilising buffer. The permeabilised cells were then immediately subjected to AFC. ProLine Rainbow Beads (#1451085; Bio-Rad Laboratories) were used for calibration to ensure that GFP signals measured during quantitative analyses were within the linear range of the PMT detectors. Data acquisition was performed with the ProSort software (Bio-Rad Laboratories), and FlowJo (https://www.flowjo.com; Ashland, OR, USA) was used for further data analysis and Figure preparation.

### STED microscopy and image analysis

Cells were seeded on high-precision glass coverslips (#1.5H; Paul Marienfeld GmbH & Co. KG, Lauda-Königshofen, Germany) and synchronised to G2 phase. The cells were briefly rinsed with PBS, followed by fixation with 2.4% FA for 30 min and subsequent permeabilisation with 0.33% TX-100 in PBS for 5 min. Subsequent labelling of sfGFP-tagged polypeptides with an anti-GFP-tag nanobody (NanoTag Biotechnologies GmbH, Göttingen, Germany), as shown for all specimens in Figure 5, and additionally with an antibody against NUP153, as in Figure 5A and 5C, was performed according to the standard IFM procedure outlined above, yet without DNA staining. Specimens were mounted in SlowFade Gold (Invitrogen). STED images were acquired either with a Nikon Eclipse Ti2 microscope body (Nikon, Tokyo, Japan) equipped with an Abberior STEDYCON (Abberior Instruments GmbH, Göttingen, Germany) and a Plan Apo 100×/1.45 Oil objective (Nikon), or with a STED Expert Line microscope (Abberior Instruments GmbH) equipped with a UPlanSApo 100×/1.40 Oil objective (Olympus, Tokyo, Japan). Both STED microscopes were equipped with a pulsed 775 nm STED laser for depletion. Laser powers were optimised for each sample.

To determine the mean distribution profiles of sfGFP fluorescence across the NEs of WT and mutant cell lines in STED micrographs of NEs focused in the nuclei’s equatorial plane, the signal intensity profiles were captured with Fiji/ImageJ (https://imagej.net/Fiji) by drawing lines perpendicular to the NEs. More than 500 such lines were selected from five different nuclei, while omitting only those regions in which NUP153 labelling pattern indicated that the imaged NE was not in an equatorial focal plane. Importantly, line profiles were selected solely on the basis of the NUP153 distribution pattern, without viewing the sfGFP images. The corresponding sfGFP line profiles were only revealed after having completed collection of the entire NUP153 dataset. The signal intensity values for each of the NUP153 and sfGFP line profiles were then aligned using RStudio (R Core Team, 2026; R Foundation for Statistical Computing, Vienna, Austria; https://www.R-project.org/) and further processed in Excel (Office package; Microsoft, Redmond, WA, USA).

Unlike the sfGFP signals of the double-labelled NEs in equatorial focal plane, the line profiles of NBs labelled only with GFP nanobodies and captured as face-on images, shown in Figure 5E, had to be selected directly on the basis of their sfGFP signal distribution pattern, which introduced some bias. For NBs built of C-terminally sfGFP-tagged TPR, which appeared only as roundish dots, the criterion for choosing a line profile across the central signal maximum was a clearly outlined roundish signal cluster of near-spherical shape, while omitting dots of more ellipsoidal shape, which might reflect local curvature of the NE. Similarly, NBs built of N-terminally sfGFP-tagged TPR were selected when they were spherical and ring-like in appearance, again omitting those of ellipsoidal shape, but also those that did not reveal a clear signal trough between the two signal peaks of a line-crossed ring. The signal intensity values for these sfGFP line profiles were again processed in Excel.

### STORM imaging and image analysis

Cells were seeded on high-precision glass coverslips (18 mm diameter, 170 ± 5 μm thickness; #1.5H, Paul Marienfeld GmbH). Before use, one quarter of each coverslip had been sputter- coated on one side with a thin aluminum layer by the in-house workshop of our institute, thereby forming a mirror surface required for initial alignment of the sample and microscope objectives, as described previously (Bates *et al*., 2022). The asynchronous cells, grown on the coated side of the coverslips, were harvested at a maximum cell density of 70%, fixed for 30 min in freshly prepared 2.4% FA in PBS, and quenched with 50 mM NH4Cl in PBS for 5 min. The coverslips were then either stored at 6–8°C overnight or directly permeabilized with 0.2% TX-100 in PBS for 10 min, briefly rinsed with PBS, and incubated with several drops of Image- iT FX Signal Enhancer (# I36933; Thermo Fisher Scientific) as a first blocking reagent for 30 min. Following a second blocking step in PBS with 2% BSA for 60 min, the cells were incubated for 60 min in PBS containing 2% BSA together with an ELYS antibody (# HPA031658; Sigma-Aldrich), which had been coupled to amine-reactive AF647 (#A20006; Thermo Fisher Scientific) according to the distributor’s instructions, and a custom-made Cy5.5-coupled anti-GFP Q nanobody (NanoTag GmbH, Göttingen). Following three brief rinses in PBS and three subsequent washes in PBS for 5 min each, each coverslip was mounted onto a second coverslip of 30 mm diameter, which in turn had been glued onto a custom-made support consisting of a stainless-steel disc with an open aperture (Bates *et al*., 2022). All mounting steps, including application of the STORM imaging buffer (50 mM Tris pH 8.0, 10 mM NaCl, 10% glucose [w/v], 143 mM β-mercaptoethanol), were performed as described in detail previously, as was preparation of the enzymatic oxygen-scavenging system added to the imaging buffer (Bates *et al*., 2022). The resulting sandwich of the two coverslips and the stainless-steel disc was then magnetically coupled to the sample holder connected to the sample translation stage of the custom-built 4Pi-STORM microscope described previously in detail (Bates *et al*., 2022).

Image data was acquired using two-colour 4Pi-STORM and STORM data analysis was performed using a custom-written IDL application, as described previously (Bates *et al*., 2022). Specifically, the STORM image stacks and the PSF calibration stack were transformed to map the four image channels to a common coordinate system, resulting in co-registration of the image data within a root-mean-square deviation of less than 2 nm. Single-fluorophore localisation, phase correction, and drift correction were then performed, yielding a two-colour 3D localisation dataset for each sample ROI (Bates *et al*., 2022).

### Protein structure predictions and molecular graphics tools

Similar to previous structure predictions of ZC3HC1 and Pml39p (Gunkel *et al*., 2023), the predictions in the current study were initially obtained from AlphaFold2 (Jumper *et al*., 2021; Tunyasuvunakool *et al*., 2021), yet here using the computing resources provided by the COSMIC2 platform (https://cosmic2.sdsc.edu; Cianfrocco *et al*., 2017). The final versions for ZC3HC1 and Pml39p, as presented in Figures 7D and S7, were then generated with AlphaFold3 (Abramson *et al*., 2024) via the AlphaFold web-interface (https://alphafoldserver.com), and essentially confirmed the AlphaFold2 predictions.

Similarly, the predictions for the homodimeric segments of TPR and Mlp1p were initially obtained with AlphaFold Multimer, as an extension of AlphaFold2, likewise using COSMIC2. The predictions shown in Figure 7A and 7B represent such AlphaFold2/COSMIC2 predictions. By contrast, the interactions between ZC3HC1 and homodimeric TPR segments shown in Figure 7D2–7D5, as well as between Pml39p and the corresponding Mlp1p segments, shown in Supplemental Figure S7I2–S7I5, again represent AlphaFold3 predictions, as do the predicted interactions between ZC3HC1 and other human CC-forming proteins shown in Supplemental Figure S7H. The TPR:ZC3HC1:TPR ensemble shown in Figure 7D6 was created in ChimeraX (https://www.cgl.ucsf.edu/chimerax; Pettersen *et al*., 2021), using the CIF files provided by AlphaFold3. Chimera (https://www.rbvi.ucsf.edu/chimera; Pettersen *et al*., 2004) and ChimeraX were also used for in silico modifications, including certain spatial rearrangements of CCs and their positioning (Figure 7B, Supplemental Figure S8B), and for the colouring of AlphaFold3 structures in this study.

### Image analysis for signal quantification at NEs

ImageJ/FiJi software (versions 1.50i–1.54k, National Institutes of Health, USA; Schneider *et al*., 2012) was used for image-based quantification of NE-associated signal intensities presented in Supplemental Figures S2F, S4E, and S4G, with NE-segment selection and subtraction of cytoplasmic and nuclear background performed as described previously (Gunkel *et al*., 2021). To also quantify NE-associated signal intensities automatically, we established an image analysis pipeline. First, a U-Net convolutional neural network (Falk *et al*., 2019) was trained on 30 representative image crops extracted from the CLSM micrographs, with each crop containing 4–6 nuclei whose NEs had been manually annotated. Blurred NE segments, corresponding to nuclei considered not to be in the equatorial plane and therefore out of focus, were deliberately left unannotated, so that the network learned to preferentially predict sharply resolved, in-focus NE rings. The training dataset was expanded fourfold by on-the-fly data augmentation, including random rotations, random rescaling, and addition of artificial Gaussian noise. The network was trained for 50 epochs using a BCEDice loss until convergence. This compound objective combines binary cross-entropy with a soft Dice term at equal weights (0.5/0.5). It was proposed for class-imbalanced segmentation (Taghanaki *et al*., 2019) and was implemented using the bio-image-unet Python library (https://github.com/danihae/bio-image-unet). Once trained, the same network was used to generate NE masks from micrographs of all cell lines quantified in the current study. The NE probability maps returned by the network, which assign every pixel a value between 0 and 1, were binarised using a fixed probability threshold of 0.01, applied identically to all images and subjected to area closing with a 2-pixel threshold in order to close small gaps. Next, Euler characteristics for 2D binary images were used for shape-based filtering and calculation of the genus of each NE as a ring-like object. Only NEs presenting as a single closed ring with genus 1, i.e. as a continuous loop enclosing a single hole, were retained. By contrast, incomplete rings (genus 0), either with out-of-focus parts or representing NE segments at the image borders, and multi-hole structures (genus ≥ 2), representing clustered nuclei appearing in contact with each other, were excluded. In addition, nuclei with perimeters outside the range of 300–1500 pixels were filtered out. For each retained nucleus, the mean signal intensity within its ring-shaped NE mask was calculated. In addition, each NE mask was expanded by 5 pixels on both sides, and the original NE mask was then subtracted from this expanded mask, yielding two rim masks: an inner, nucleoplasmic mask and an outer, cytoplasmic mask, which were used for local background correction. The mean intensity values in each of these two rim masks were determined, weighted by 0.33, and subtracted as background from the mean intensity value of the actual NE mask. For co-cultured populations of two cell lines with distinct NE signal intensities, the same segmentation pipeline was applied to all nuclei. To assign the already quantified nuclei to one of the two cell lines, a two-component Gaussian mixture model was fitted to the background-corrected mean NE intensities. In cases in which only one of the two cell lines additionally showed a pronounced nucleoplasmic signal, as was the case for the non- permeabilised ZC3HC1 Δ487–502 cells shown in Figure 3C, the intensity measured in the nucleoplasmic rim mask was included as a second feature for assignment. Nuclei were then hard-assigned to one of the two Gaussian components, i.e. assigned to the component for which the model calculated the higher posterior probability. This assignment was used only for images in which the fitted model supported a robust separation of two cell populations: (i) the Bayesian information criterion (BIC) had to favor a two-component over a one-component Gaussian mixture model by more than 10 (Kass and Raftery, 1995), (ii) each component had to account for at least 10% of the analysed nuclei, and (iii) the standardised distance between the component means had to be at least 2. This distance was calculated as Cohen’s d when only NE intensity was used, or as Mahalanobis distance when NE intensity and nucleoplasmic rim intensity were used together. In a two-component Gaussian model, a distance of 2 corresponds to an expected Bayes-optimal misclassification rate of approximately 16%. Images failing any of these criteria were excluded from cell-line-specific statistics. For data representation, we applied a cut-off of the upper and lower 1% of the total counts to remove outliers. The seeming overlap between two cell lines in pooled dot plots of NE signal intensities, as seen in Figures 2C3, 3B2, 3C2, 4A2, 4B2, 4D2 and 4F2, did not reflect an overlap at the level of individual images. Rather, it arose when nuclei from multiple independently normalised images or datasets were combined, such that data points originating from different images occupied overlapping intensity ranges in the pooled representation. When the same data were resolved at the level of individual images, the two populations remained separated, as exemplified in Supplemental Figure S2G.

## Author contributions

Conceptualization, V.C.C. and P.G.; data curation, P.G., V.C.C., M.B and D.H.; formal analysis, P.G., V.C.C., M.B and D.H.; investigation, P.G., V.C.C., M.B.; methodology, P.G., V.C.C., D.H. and M.B.; project administration, V.C.C. and P.G.; resources, V.C.C.; supervision, V.C.C.; validation, P.G., V.C.C. and M.B.; visualisation, P.G., V.C.C. and M.B.; writing–original draft preparation, V.C.C.; writing–review and editing, V.C.C., P.G., D.H and M.B. Artifical intelligence (AI) was used for refining text and help with code writing. All AI- generated contents were thoroughly checked by the authors. All authors have read and agreed to the published version of the manuscript.

Conflict of interest

The authors declare no conflict of interest.

## Acknowledgements

We gratefully acknowledge Frank Schwarz and Kerstin Mohr for contributions to Y2H experiments, Gabriele Hawlitscheck and Susanne Brandfass for contributions to cloning, and Steffen J. Sahl for helpful discussions. Furthermore, we thank Patrick Cramer, Jan Ellenberg, Gražvydas Lukinavičius, Melina Schuh, Feng Zhang, and in particular Dirk Görlich for kindly providing research materials and for sharing equipment. We are particularly grateful to Dirk Görlich and his coworkers Bianca Mußil, Katrin Gregor, and Lareen Gräser for kindly providing an as-yet unpublished TMEM209-specific nanobody, which they had isolated, purified, and labelled, and which we used for the characterisation of our TMEM209-sfGFP cell line.

## Abbreviations

aa, amino acid; AFC, analytical flow cytometry; BLD, BIR-like domain; CC, coiled-coil; CEZ, chromatin exclusion zone; ChT, chromosome territory; CT, carboxy-terminus; CTD, carboxy-terminal domain; FA, formaldehyde; IFM, immunofluorescence microscopy; KO, knockout; LCI, live cell imaging; NB, nuclear basket; NBLS, nuclear basket-like structure; NE, nuclear envelope; NBD, NPC binding domain; NPC, nuclear pore complex; NR, nuclear ring; NTD, amino-terminal domain; NT, amino-terminus; NuBaID, nuclear basket interaction domain; NUP, nucleoporin; PBD, Pml39 binding domain; ROI, region of interest; SEM, scanning electron microscopy; STED, stimulated emission depletion; STORM, stochastic optical reconstruction microscopy; TBE, TPR-binding element; TEM, transmission electron microscopy; TR, terminal ring; XL-MS, cross linking mass spectrometry; WT, wild type; Y2H, yeast two-hybrid; ZBD, ZC3HC1 binding domain

## Notes

### Competing Interest Statement

The authors have declared no competing interest.

### Summary of Updates

We have added 4Pi-STORM data visualizing the three-dimensional organization of NPC-associated TPR:ZC3HC1 ensembles, presented in the new Figure 6 and Supplemental Figure S6. The Discussion has been expanded to address the additional insights provided by these data. Furthermore, Mark Bates has been added to the author list in connection with the included STORM data. We have also made minor revisions to the Title, Abstract, and several other parts of the manuscript to incorporate and refer to the additional STORM data. Finally, we have corrected several incomplete references, including missing page numbers.

