## Supplemental Figures for "Hierarchical TPR:ZC3HC1 interactions drive stepwise assembly of serially arrayed nuclear basket appendages extending into vertebrate cell nuclei"

#### Supplemental Figure S1 (1/2)

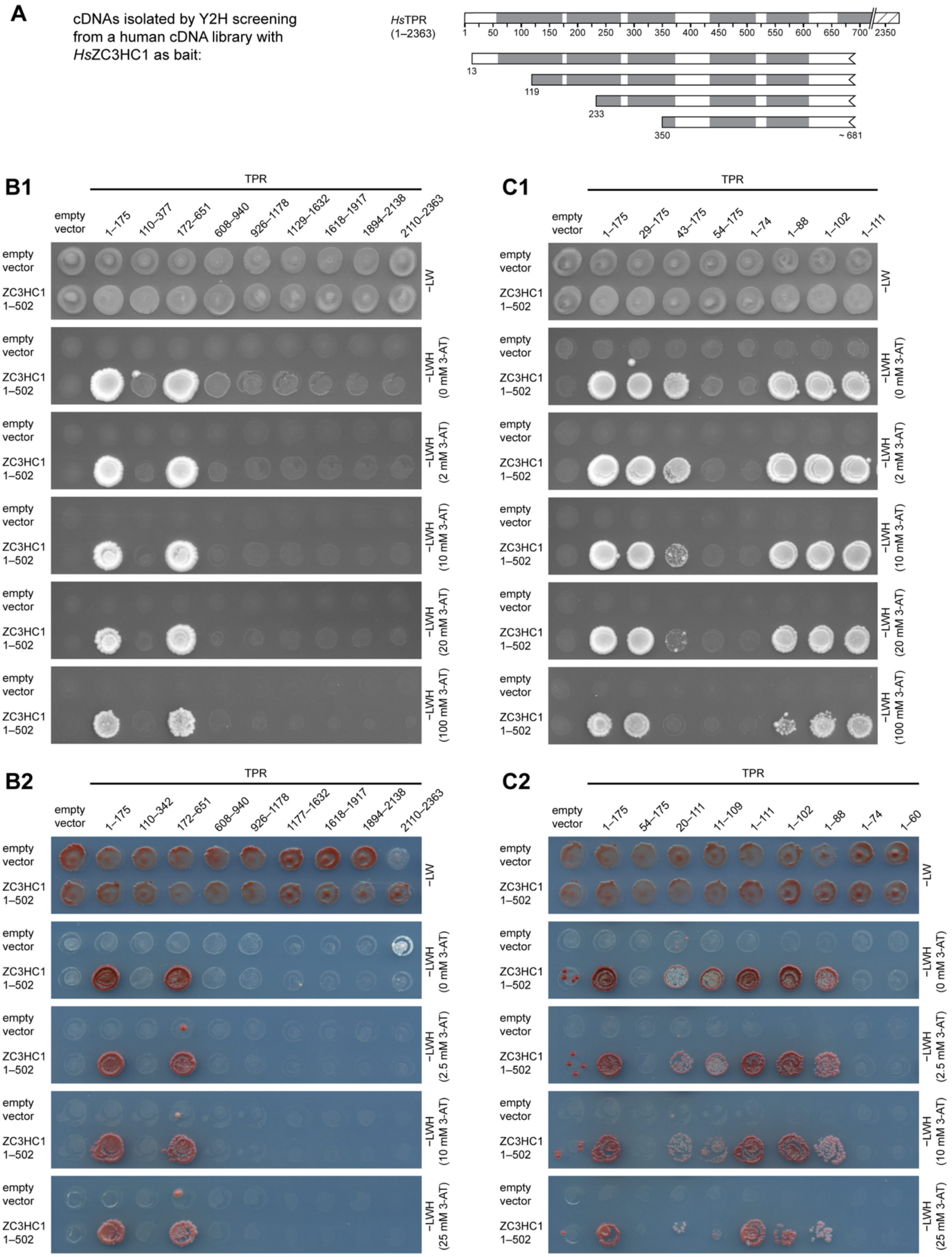

Supplemental Figure S1 (2/2)

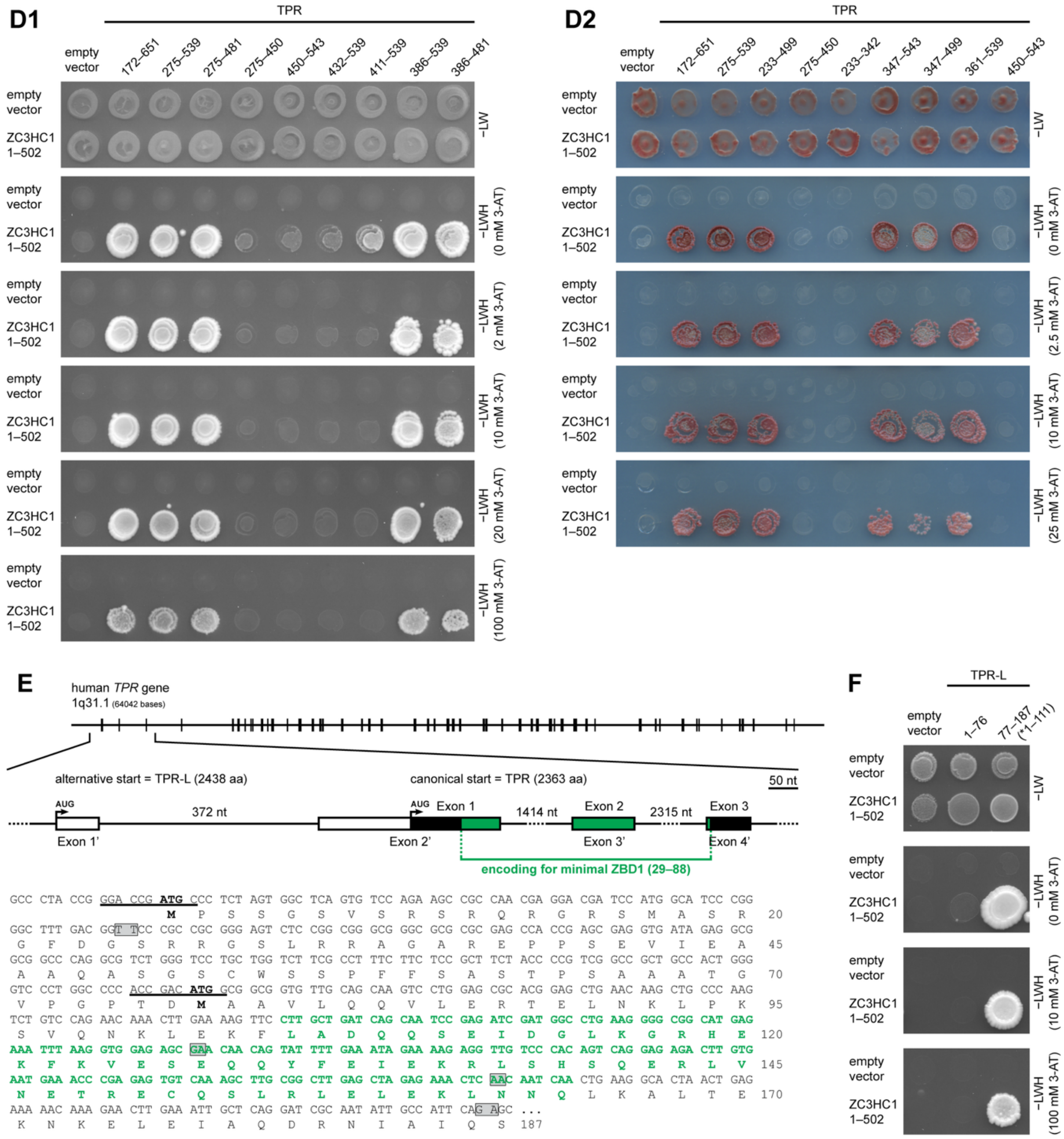


**Supplemental Figure S1. ZC3HC1 interacts with two distinct segments of TPR's NTD. (A)** Screening of a human Y2H library with a bait vector encoding full-length *Hs*ZC3HC1 yielded four different, partially overlapping cDNAs encoding N-terminal *Hs*TPR segments, here depicted schematically to scale relative to full-length TPR. The oligo(dT)-primed cDNA library was derived from poly(A)+ RNAs pooled from various adult human tissues (Normalized Matchmaker Human Universal cDNA Library, Clontech 638874). Screening of

$1.1 \times 10^7$ clones was performed at the Proteomics Core Facility for protein interaction at the German Cancer Research Center in Heidelberg, Germany, using the low-level expression vector (pGBT9) encoding GAL4-BD-ZC3HC1, without the addition of 3-AT to the selection medium. The only cDNAs isolated that represented the same protein more than once encoded the four overlapping TPR segments; a few additionally isolated solitary clones each encoded a different protein, of which none was SKP1 or another SCF-complex component formerly reported as ZC3HC1 binding partners (e.g., Bassermann *et al.*, 2005). Relative to the mRNA sequence of *Hs*TPR (NM_003292), the TPR ORFs began at nt 333, 652, 933 or 1344, corresponding to aa 13, 119, 233 and 350, respectively. Their 3'-ends included an adenine-rich region downstream of nt 2343 that is prone to hybridise with oligo(dT) primers during cDNA synthesis, thereby tolerating some mismatches, and thus leading to EST sequences that do not represent splice variants of TPR (e.g., Kuznetsov *et al.*, 2002). Despite some uncertainty regarding the assignment of the TPR nucleotides around position 2343, all of the isolated cDNAs encoded TPR segments that included aa residue 681. Therefore, all these TPR segments included the formerly defined NBD region of TPR, located between aa 350–610 (Hase *et al.*, 2001). **(B)** Y2H experiments analysing interactions of full-length *Hs*ZC3HC1 with segments of *Hs*TPR. TPR segments were tagged at their NT with GAL4-AD. These prey plasmids and an "empty" GAL4-AD vector had been transformed into MATα yeast cells, followed by mating with MATa yeast cells transformed with a bait vector encoding GAL4-BD alone or a fusion of GAL4-BD to the NT of ZC3HC1. Representative colony growth is shown on −LW medium, revealing successful mating between the MATα and MATa yeast strains, while Y2H interactions are visualised by colony growth after replica-plating onto −LWH medium supplemented with different 3-AT concentrations. **(C, D)** Additional Y2H experiments using further TPR segments for fine-mapping of ZBD1 (C) and of ZBD2 (D). For ZBD1, colony growth (up to 100 mM 3-AT) was observed only with segments near the canonical NT that included at least 29–88. For ZBD2, colony growth required at least 386–481. **(E)** Schematics of the 5’ end of the human TPR gene, illustrating the relative position of an alternative start codon upstream of the ORF encoding the canonical isoform of TPR. The non-canonical, second isoform of human TPR, here termed TPR-L, comprises 76 additional aa, of which 24 are encoded by an exon (exon 1’) upstream of the more commonly used start codon. The Kozak sequences (Kozak, 1984, 1987) surrounding both start codons are underlined. Boxed nucleotides in the ORF of TPR indicate exon-exon junctions, including the last nucleotide of an upstream exon and the first nucleotide of the downstream exon. Sequences encoding the mapped minimal region of ZBD1 (aa 29–88), including the corresponding nucleotide and protein sequences, are highlighted in green. **(F)** Y2H experiments analysing interactions of full-length *Hs*ZC3HC1 with N-terminal segments of TPR and TPR-L, with interactions only observed with the ZBD1-containing N-terminal segment of the canonical TPR, but not with aa 1–76 of TPR-L.

#### Supplemental Figure S2 (1/5)

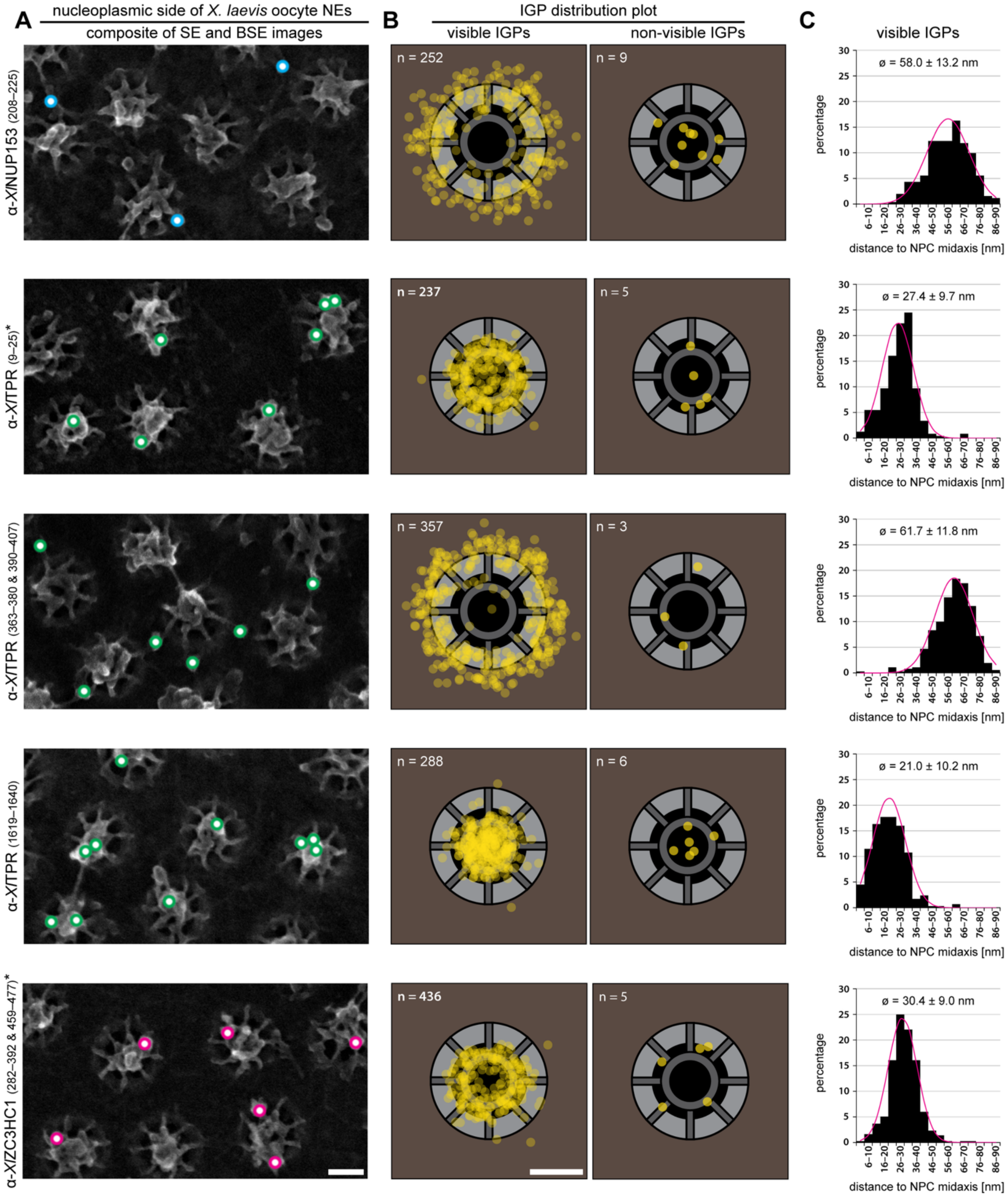

### Supplemental Figure S2 (2/5)

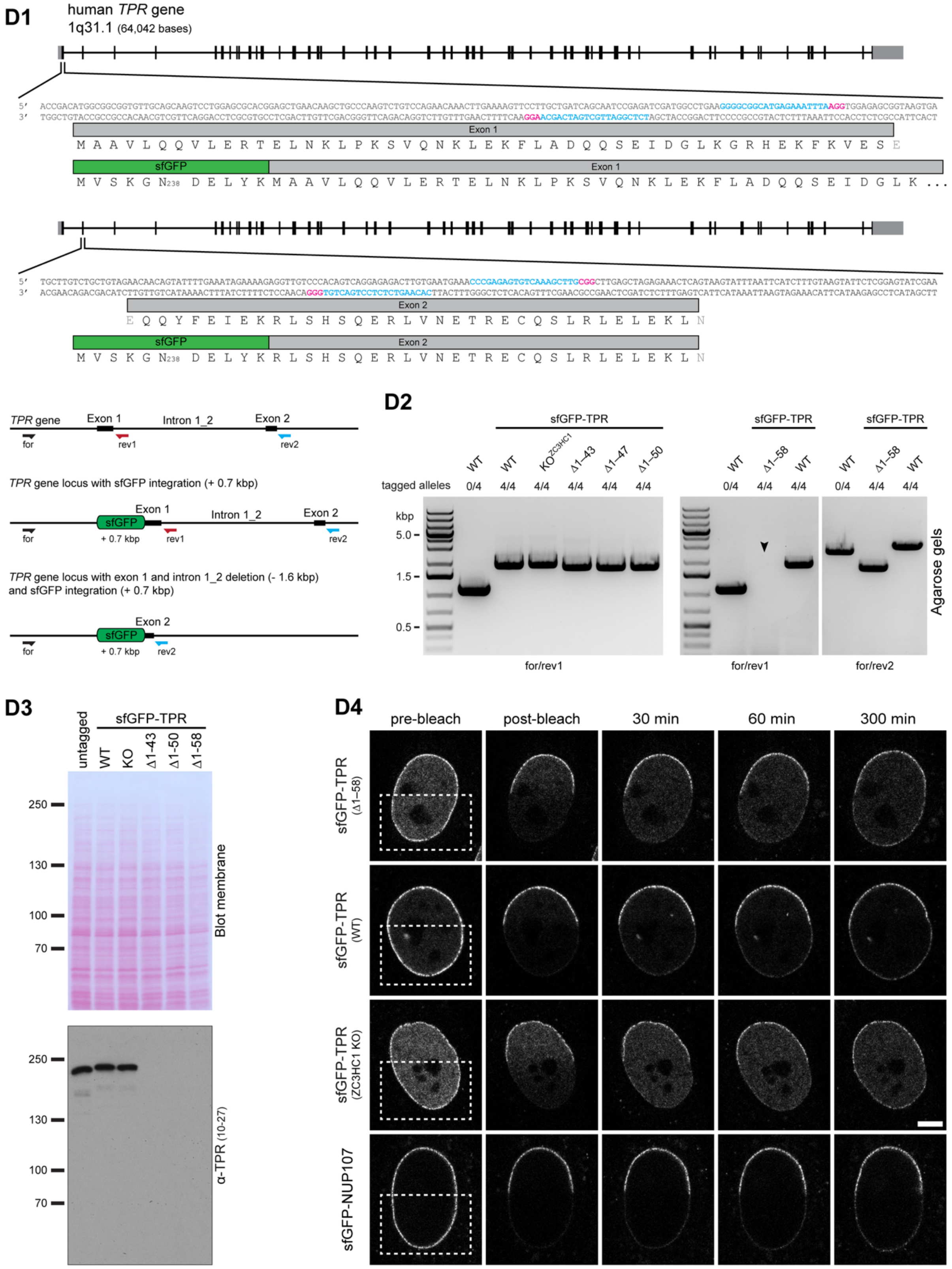

Supplemental Figure S2 (3/5)

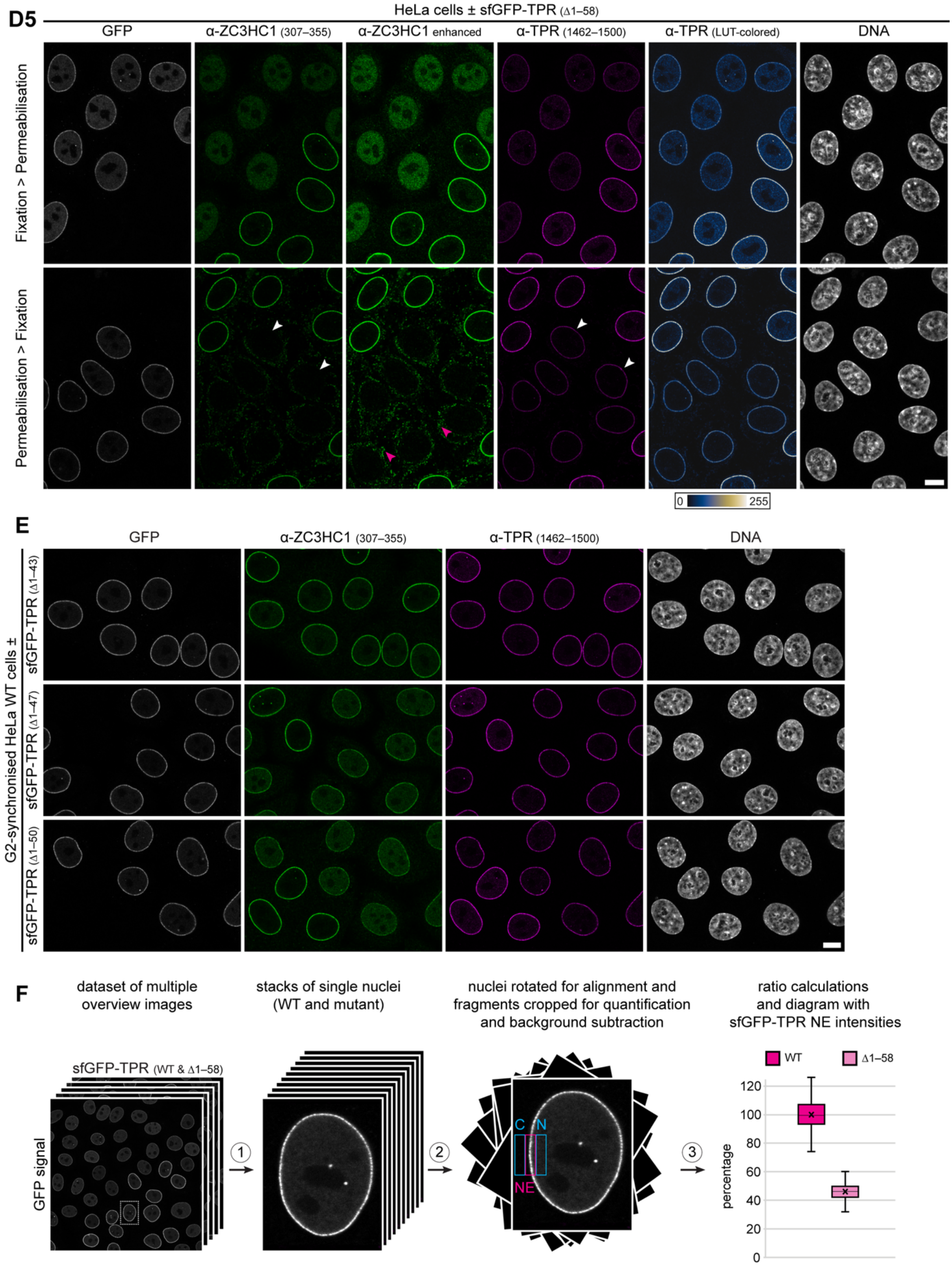

### Supplemental Figure S2 (4/5)

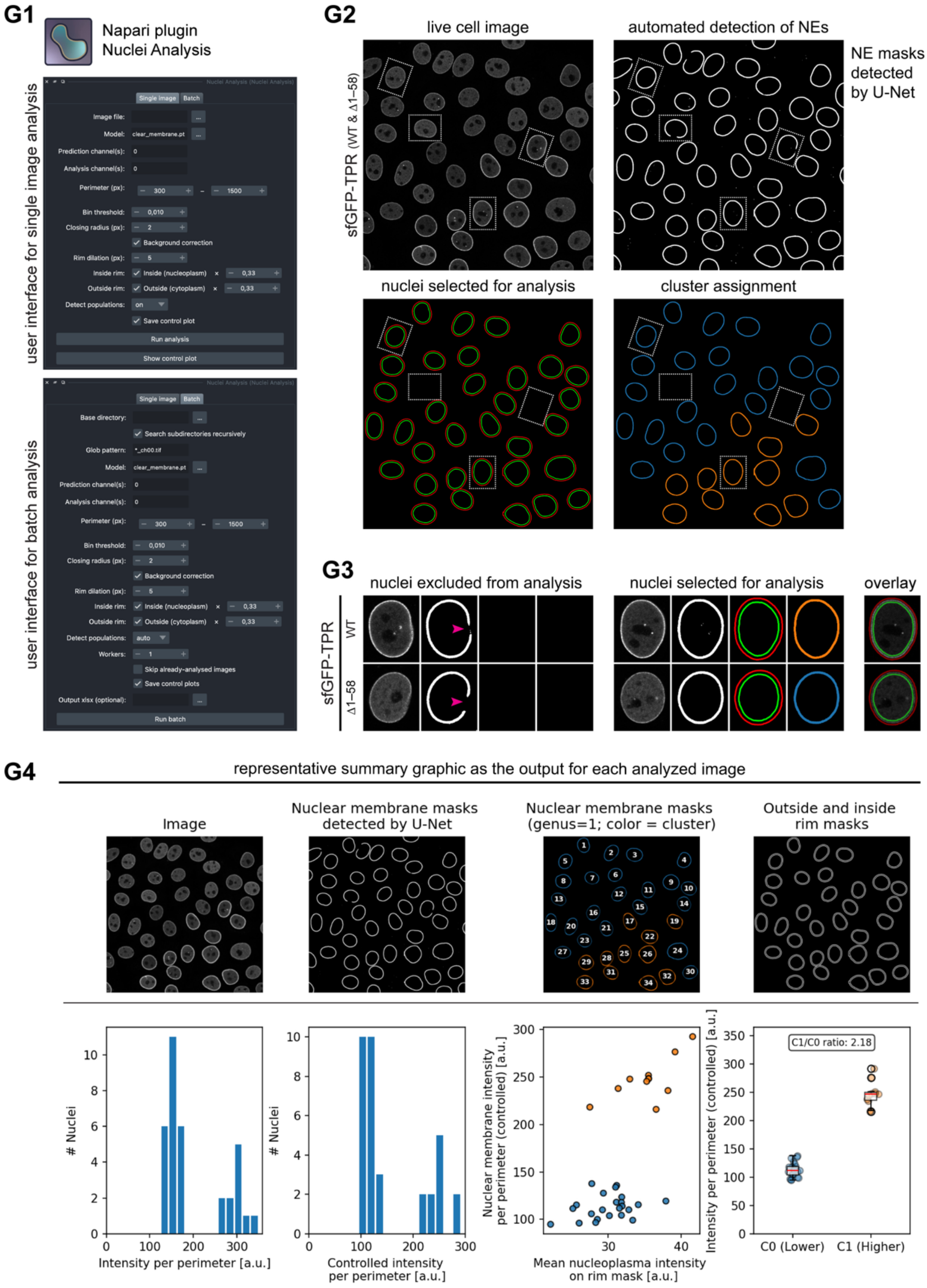

Supplemental Figure S2 (5/5)

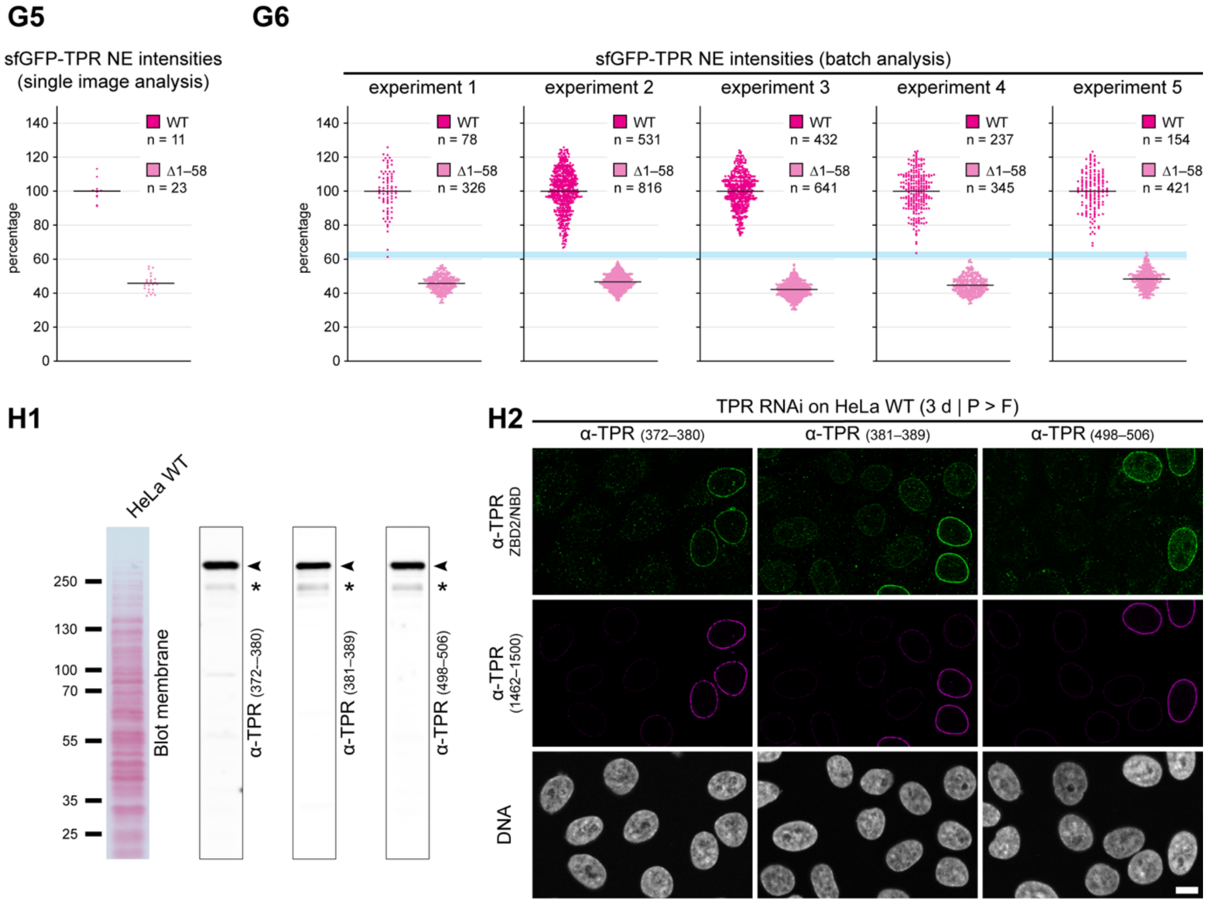


**Supplemental Figure S2. TPR's ZBD1 positions ZC3HC1 at the NB's distal end, quantitative analyses reveal the extent of subsequent recruitment of the TPR T2 subpopulation, and subpopulation-specific antibodies distinguish TPR T1 from T2 polypeptides. (A–C)** ZC3HC1 is located at the distal end of the NB proper, close to TPR's NT and the C-terminal end of TPR's rod domain, while TPR's NBD is located at the NR. **(A)** Immuno-SEM of manually isolated NEs from *Xenopus laevis* stage V oocytes, having selected NE regions in which NBs lacked apparent fibrillar material appended to their distal ends. Immunolabelling was performed with epitope-specific antibodies targeting different regions of ZC3HC1, TPR, and NUP153, the latter used as reference. TPR's NT was labelled with α-*Xl*TPR aa 9–25, TPR's NBD, as formerly defined (Hase *et al.*, 2001), with α-*Xl*TPR aa 363–380 and aa 390–407, and the C-terminal end of TPR's rod domain with α-*Xl*TPR aa 1619–1640. Image acquisition and analysis were performed as described previously (Gunkel *et al.*,

2021). Representative composite images show SE micrographs of the NE's nuclear face overlaid with BSE images revealing the positions of immunogold particles (IGPs), encircled in blue for NUP153, magenta for TPR, and green for ZC3HC1. Bar, 50 nm. **(B)** Distributions of IGPs relative to an idealised NPC and its NB in face-on view. Schematics are drawn to scale, with outer and inner NR diameters of 110 and 70 nm, respectively, and an outer TR diameter of 55 nm. Bar, 50 nm. IGPs visible in both BSE and SE mode, indicating positions on top of the labelled material, and IGPs invisible in SE mode, indicating positions buried beneath proteinaceous material, are shown separately. The datasets obtained with α-*Xl*TPR aa 9–25 and α-*Xl*ZC3HC1 aa 282–392 and aa 459–477 were already presented previously (Gunkel *et al.*, 2021), whereas the other TPR and NUP153 datasets had not yet been published at that time. **(C)** Bar diagrams corresponding to the datasets shown in B, representing the percentages of IGPs, grouped into 5-nm-wide windows, at different radial distances from the NB's longitudinal axis, here represented by the y-axis. Mean radial distances and standard deviations (SDs) were derived from all measured distances of each dataset. **(D)** Characterisation of HeLa P2 cell lines homozygously expressing N-terminally sfGFP-tagged full-length truncated versions of TPR. **(D1)** Schematic depiction of part of the human *TPR* gene comprising exons 1 and 2, which encode the N-terminal part of the canonical TPR isoform, and of the site of CRISPR/Cas9n-mediated insertion of the sfGFP ORF. As one example of the generation of the four N-terminal TPR deletion mutants, deletion of exon 1, intron 1 and part of exon 2, resulting in the mutant encoding sfGFP-TPR Δ1–58, is depicted. The relative positions of forward (for) and reverse (rev) primers used for the PCR analyses shown in D2 are indicated. **(D2)** Genotyping by PCR amplification, as described in further detail previously (Gunkel and Cordes, 2022), after on-target integration of the sfGFP ORF and deletion of *TPR* gene segments. Note that the HeLa P2 cell line likely has four *TPR* alleles, as in HeLa Kyoto (Landry *et al.*, 2013) and in the HeLa progenitor cell line CCL-2 (Adey *et al.*, 2013), while HeLa S3 likely has five *TPR* alleles (Adey *et al.*, 2013). PCR products, shown after agarose gel electrophoresis, derive from the original untagged (0/4) progenitor HeLa P2 line, the homozygous line in which all four *TPR* alleles encoded the sfGFP-tagged full-length canonical isoform of TPR (4/4), and the mutant lines in which all four *TPR* alleles encoded sfGFP-tagged and truncated TPR polypeptides (4/4). For comparison, the PCR product of the ZC3HC1 KO cell line expressing sfGFP-tagged full-length canonical TPR is also included. Both full-length sfGFP-TPR lines had been described previously (Gunkel and Cordes, 2022). The arrowhead indicates successful deletion of intron 1 in the sfGFP-TPR Δ1–58 line, which is no longer detectable when using the reverse primer rev1. **(D3)** IB of total cell extracts from untagged HeLa P2 cells, the homozygous cell line expressing sfGFP-tagged full-length TPR, and the mutant lines in which all *TPR* alleles had been truncated, next to total cell extracts from ZC3HC1 KO cells expressing sfGFP-tagged full-length TPR. Immunolabelling was performed with an epitope-specific *Hs*TPR antibody targeting aa 10–27 of the canonical TPR, with this epitope absent from the N-terminal deletion mutants. **(D4)** Fluorescence recovery after photobleaching (FRAP) experiments with the different cell lines expressing sfGFP-tagged full-length and truncated versions of TPR in WT and ZC3HC1 KO backgrounds. Photobleaching was performed to assess the stability of NPC association of TPR Δ1–58 relative to full-length TPR. A HeLa P2 line expressing all NUP107 as N-terminally sfGFP-tagged polypeptides, characterised in detail further below (Supplemental Figure S4B), was included for comparison. The rectangular areas marked in the pre-bleach images were subjected to pulses of full laser power for 2 min, followed by acquisition of post-bleach images at the indicated time points. GFP fluorescence recovered with similarly slow kinetics at the bleached NE segments in all cell lines expressing sfGFP-tagged TPR, indicating similarly stable NPC association. Bar, 5 µm**.** **(D5)** Double-IFM of co-cultured populations of HeLa P2 WT cells with non-tagged TPR and HeLa P2 cells homozygously expressing sfGFP-TPR Δ1–58, using the same cell-line combination and antibodies as in Figure 2B. Here, the G2-synchronised cells were either permeabilised before or after fixation, the latter procedure being identical to the one used for the experiment presented in Figure 2B, to assess the stability of NPC association of the TPR mutant. White arrowheads mark two TPR Δ1–58 cells in which the soluble ZC3HC1 pool, no longer capable of NB association, had been removed by permeabilisation and is therefore no longer visible. As an aside, magenta arrowheads mark cytoplasmic structures with which this particular ZC3HC1 antibody cross-reacts, yet only when cells were permeabilised with TX-100 prior to fixation. As in Figure 2B, the NE-associated amounts of TPR in TPR Δ1–58 cells are markedly diminished, but show no apparent further reduction upon permeabilisation before fixation. The TPR micrograph is also shown colour-graded next to a colour look-up table (LUT), illustrating the approximately twofold reduction in TPR immunolabelling intensity at the NEs of TPR Δ1–58 cells. Bar, 10 μm. **(E)** Double-IFM of co-cultured populations of HeLa P2 WT cells with non-tagged TPR and HeLa P2 lines homozygously expressing either sfGFP-TPR Δ1–43, sfGFP-TPR Δ1–47 or sfGFP-TPR Δ1–50. Only a minor reduction in NE-associated ZC3HC1 was observed in TPR Δ1–43 cells, with a somewhat stronger reduction in TPR Δ1–47 and TPR Δ1–50 cells, accompanied by corresponding decreases in the NE-associated amounts of TPR. None of these mutants, however, showed the approximately twofold reduction in total NE-associated TPR or the complete loss of NE-associated ZC3HC1 observed with TPR Δ1–58 (Figure 2B; Supplemental Figure S2D5). Bar, 10 μm. **(F)** Image-based manual quantification of relative NE-associated IF or sfGFP signal yields in co-cultured populations of two cell types expressing different amounts of the same or different versions of a given protein. Cells were co-cultured on the same coverslip and imaged using identical microscope settings. The flow chart exemplifies the procedure of NE-associated signal quantification, as described in detail previously (Gunkel *et al.*, 2021), for a co-cultured population of cells expressing sfGFP-TPR or sfGFP-TPR Δ1–58. In brief, randomly acquired overview images were used to compile (1) essentially all clearly focused equatorial nuclear sections without prior assignment to either cell type. Initial focusing had been performed using the DNA-staining channel, here visualising live cells stained with SiR-Hoechst, followed by rapid fine-focus adjustment in the GFP fluorescence channel used to visualise NPC and NB proteins. Next, three equally sized rectangular areas comprising an NE segment and adjacent cytoplasmic (C) and nuclear (N) regions were selected (2), and mean fluorescence intensities in these ROIs were quantified (3) using ImageJ. The rectangle dimensions were chosen such that the labelled NE occupied approximately one third of the NE rectangle, allowing background-corrected NE signal yields to be calculated as: NE signal = NE rectangle – ($^1/_3$ × C rectangle + $^1/_3$ × N rectangle). The individual NE signal yields already formed two clearly separated intensity distributions. Cells within the small region of overlap were left unassigned, and equal fractions of outliers were excluded from both sides of each distribution to avoid upward or downward bias of the respective mean values. For the co-cultured population of sfGFP-TPR and sfGFP-TPRΔ1–58 cells shown here, the resulting mean NE signal intensity ratio from three experiments, each with 100 analysed nuclei, was 1:0.46. **(G)** Automated image-based selection and quantification of NE-associated fluorescence signals in co-cultured cell populations. **(G1)** User interface of the Napari plugin *Nuclei Analysis*, showing the input panels for single-image analysis and batch analysis. **(G2)** Exemplary processing steps for automated analysis of a live cell image of a co-cultured population of sfGFP-TPR- and sfGFP-TPR Δ1–58-expressing cells. Four representative nuclei are boxed throughout the image series, including one nucleus of each cell type that passed or failed the selection criteria, respectively. The original LCI image is shown at the upper left. Automated U-Net-based detection of NE masks is shown at the upper right, including several incomplete NE rings. After shape-based filtering, shown at the lower left, only nuclei with closed NE rings were retained for analysis. For these nuclei, the inner nucleoplasmic and outer cytoplasmic rim masks used for local background correction are depicted in green and red, respectively. Following quantification and cluster assignment, shown at the lower right, NE masks are depicted in blue for the lower-intensity sfGFP-TPR Δ1–58 population and in orange for the higher-intensity sfGFP-TPR population. **(G3)** Magnified views of the four nuclei boxed in G2, illustrating examples that failed or passed the automated selection procedure. For the two excluded nuclei shown on the left, magenta arrowheads mark discontinuities in the predicted NE rings; black panels indicate that these nuclei were excluded from subsequent analysis. The two nuclei shown on the right fulfilled the selection criteria and were retained throughout NE-mask generation, background correction, and cluster assignment. **(G4)** Control plot output generated for each individually analysed image. The upper row shows, from left to right, (1) the original grayscale LCI image, (2) the NE masks detected by U-Net, (3) the subset of NE masks retained for intensity quantification after shape-based filtering for closed rings (genus = 1), and (4) the inner nucleoplasmic and outer cytoplasmic rim masks generated by expanding the respective NE masks on either side. In the third image, mask colours indicate assignment of the individual nuclei to either of the two cell populations, while the numbers identify those nuclei for which individual intensity values were exported to the accompanying data file, thereby allowing each measurement to be traced back to its respective nucleus. The plots in the lower row show, from left to right, the NE signal intensities before background subtraction, the corresponding intensities after subtraction of cytoplasmic and nucleoplasmic background signals, the background-corrected NE intensities plotted against the mean nucleoplasmic rim intensities used as an additional parameter for population assignment, and the background-corrected NE intensities of the two assigned populations displayed as a scatter dot plot. The latter additionally provides the ratio between the mean signal intensities of the two populations. **(G5)** Automated image-based quantification of relative sfGFP signal yields at the NEs of sfGFP-TPR- and sfGFP-TPR Δ1–58-expressing cells retained from a single LCI image analysed as exemplified in G2–G4. The arithmetic mean for the sfGFP-TPR WT cells is set to 100%. **(G6)** Batch analysis applying the same automated selection, quantification, and population-assignment procedure to sets of multiple images, each obtained from five independently performed experiments in which co-cultured populations of HeLa P2 WT cells, expressing either sfGFP-TPR or sfGFP-TPR Δ1–58, were analysed by LCI. **(H)** Characterisation of TPR-specific antibodies by immunoblotting and RNAi experiments. **(H1)** IB of total cell proteins from HeLa P2 cells using peptide antibodies targeting defined *Hs*TPR epitopes, with the respective target regions given in parentheses. Immunolabelling was performed on the representative Ponceau S-stained membrane shown here and on replicate membranes of the same kind. **(H2)** Representative IFM of HeLa P2 cells treated with TPR siRNAs to assess antibody target specificity by correlating TPR knockdown (KD) with the corresponding IF signals. Some cells that had remained non-transfected within the TPR siRNA-treated populations are shown as reference. Cells were permeabilised prior to fixation and then double-labelled with epitope-specific rabbit antibodies targeting different regions of TPR's NBD, as formerly defined (Hase *et al.*, 2001), and with a mouse monoclonal TPR antibody that labels both the T1 and T2 subpopulations of TPR. Bar, 10 μm.

Supplemental Figure S3 (1/3)

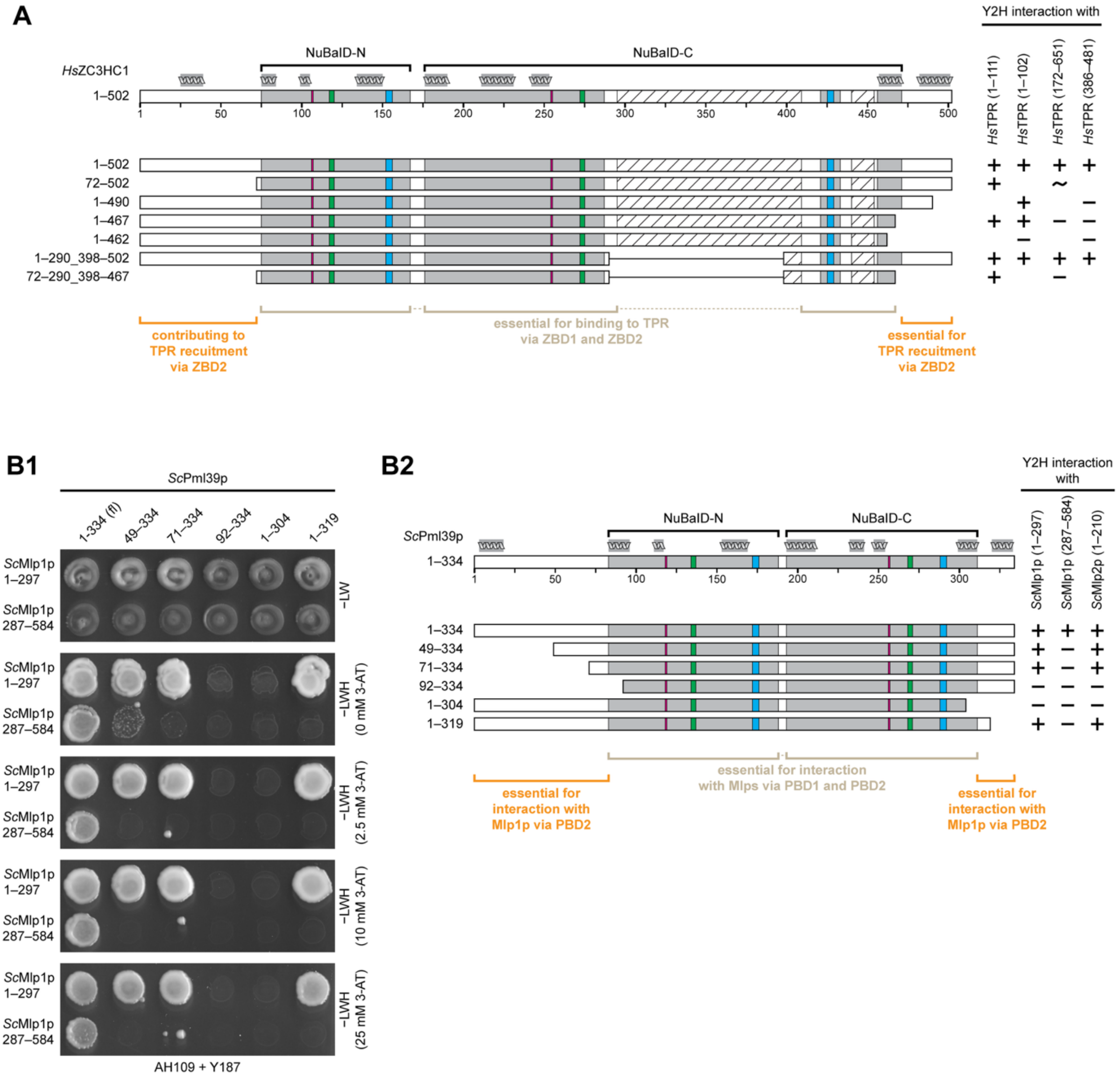

### Supplemental Figure S3 (2/3)

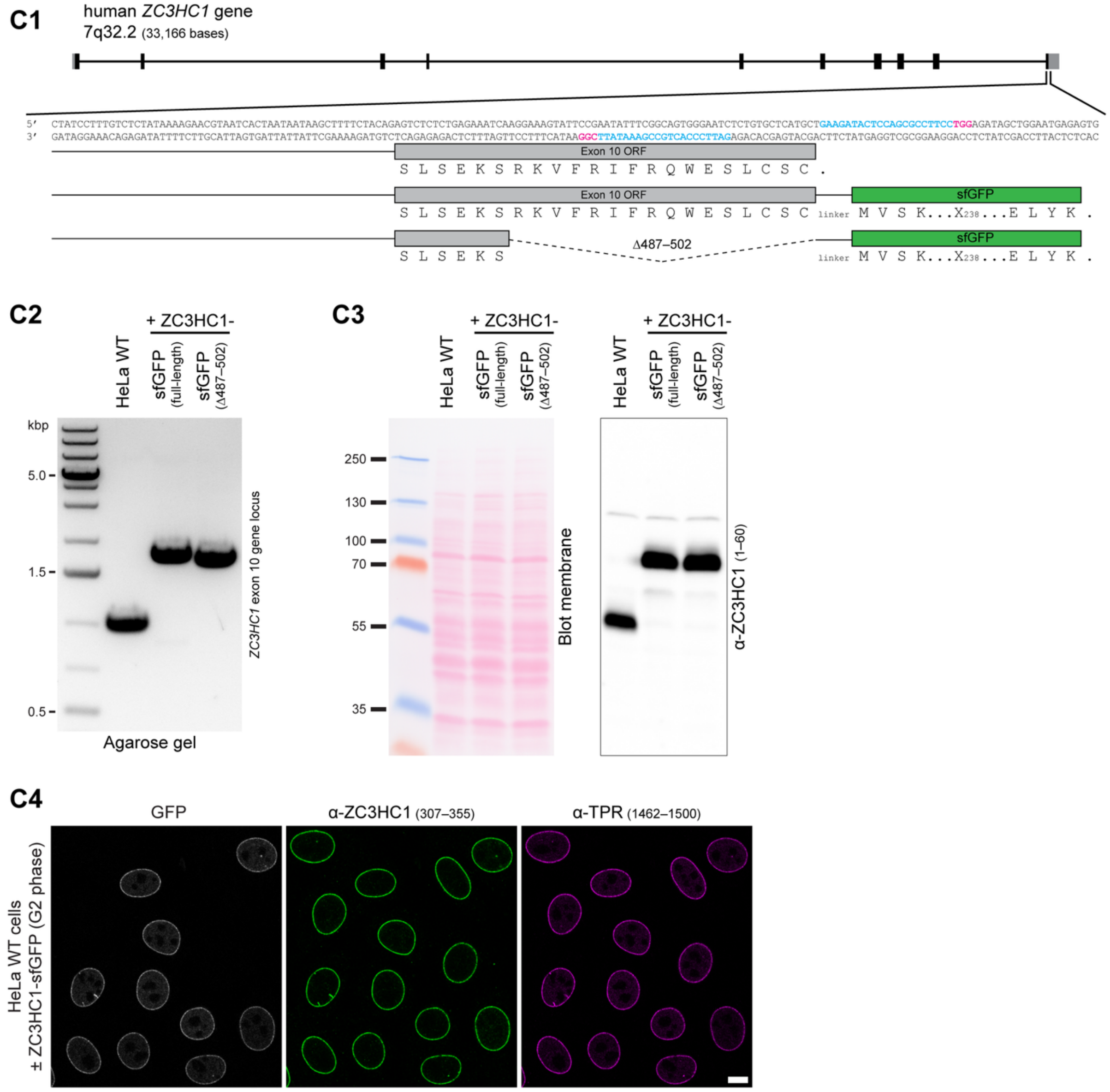

Supplemental Figure S3 (3/3)

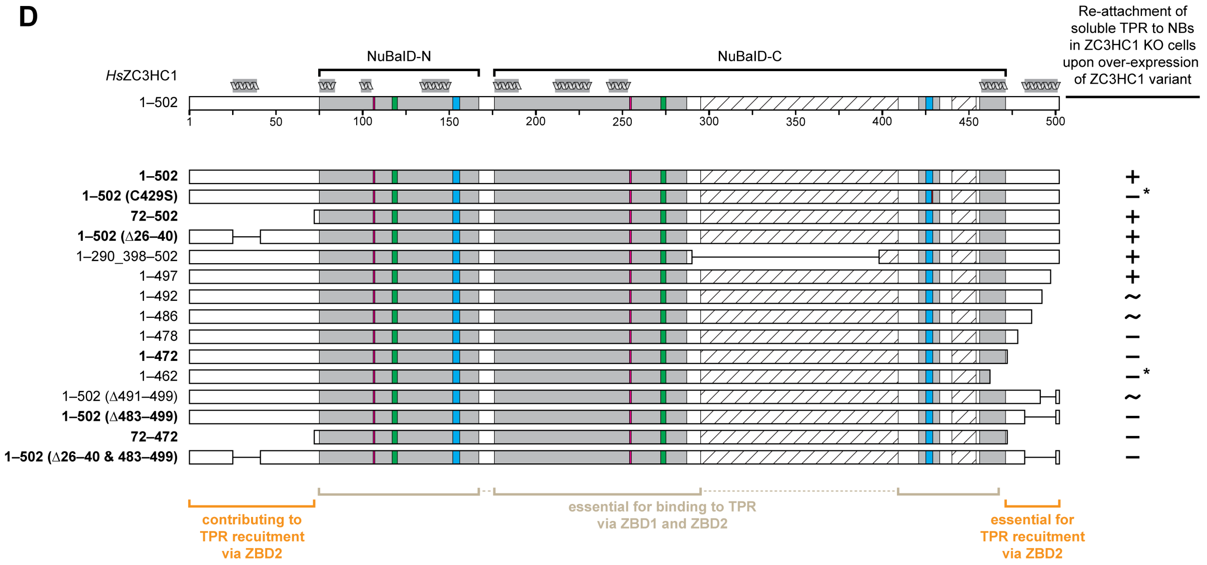


**Supplemental Figure S3. Evolutionarily conserved NuBaID-flanking elements are required for the engagement with TPR's ZBD2. (A)** Summary Y2H interaction data for full-length ZC3HC1 or deletion mutants thereof and TPR segments containing either ZBD1 (TPR 1–111 and 1–102) or ZBD2 (TPR 172–651 and 386–481) corresponding to the data presented in Figure 3A. Schematics of *Hs*ZC3HC1 and its deletion mutants are drawn to scale, with the two BLDs (NuBaID-N and NuBaID-C) highlighted in grey. Each BLD's C-X(2)-C tetrapeptide and H-X(3)-C pentapeptide, with the putative zinc-coordinating residues, are highlighted as green and blue boxes, respectively. Magenta boxes mark the positions of evolutionarily conserved G-W dipeptides. Hatched boxes represent loop-like insertions within the second BLD predicted to be largely disordered. For simplicity, other potentially unstructured regions beyond the outer BLD boundaries are not highlighted. The relative positions of nine α-helices that are part of the ZC3HC1 structure are schematically depicted above the uppermost ZC3HC1 scheme and are also drawn to scale (for further details regarding the structural characteristics of ZC3HC1, see Gunkel *et al.*, 2023). The capability of the ZC3HC1 polypeptides and TPR segments to engage in a Y2H interaction is indicated (+, colony growth even at elevated concentrations of 3-AT; ~, notably attenuated colony growth; −, no colony growth). **(B)** Y2H experiments analysing interactions of full-length *Sc*Pml39p and mutants thereof with segments of *Sc*Mlp1p and *Sc*Mlp2p. Mlp segments were fused at their NT to GAL4-AD while the NT of Pml39p was fused to GAL4-BD. **(B1)** Representative colony growth on −LW medium and after replica-plating onto −LWH medium without or with different concentrations of 3-AT, visualising Y2H interactions. Deletion of either the N- or C-terminal region flanking the NuBaID of Pml39p did not noticeably affect binding to PBD1 of Mlp1p but abolished binding to PBD2. **(B2)** Summary of Y2H interaction data for full-length Pml39p or deletion mutants thereof and Mlp segments containing either PBD1 (Mlp1p 1–297 and Mlp2p 1–210) or PBD2 (Mlp1p 287–584). Schematics of Pml39p and its deletion mutants are drawn to scale and according to the ZC3HC1 scheme in A. A hatched box is absent because the second BLD of Pml39p lacks the unstructured loop common to homologs in other phyla. Y2H interactions are indicated as in A. **(C)** Characterisation of HeLa P2 cell lines expressing C-terminally sfGFP-tagged full-length and truncated ZC3HC1. **(C1)** Schematic depiction of part of the human *ZC3HC1* gene comprising exon 10, its natural stop codon, the site of CRISPR/Cas9n-mediated insertion of the sfGFP ORF, and the segment deleted in the mutant. The pair of single guide RNA (sgRNA) sequences used for site-specific knock-in are shown in blue lettering, except for the respective protospacer-adjacent motifs (PAMs), highlighted in magenta. In addition, the deduced aa sequences of the non-tagged WT progenitor and the tagged progeny cell lines at the transitions between ZC3HC1 and sfGFP are provided for comparison. The linker sequence between ZC3HC1 and sfGFP, indicated by three dashes (-), reads GSGGSAGSAAGSGEFTGS for the full-length protein and GSAGSAS for the truncation mutant. **(C2)** Genotyping by PCR amplification. Note that three *ZC3HC1* alleles exist in the HeLa P2 cell line (Gunkel *et al.*, 2021), as in HeLa Kyoto (Supplemental Figure S4F3; Landry *et al.*, 2013), in the HeLa progenitor cell line CCL-2, and in HeLa S3 (Adey *et al.*, 2013). PCR products derive from the original untagged progenitor HeLa P2 line, the homozygous line in which all three *ZC3HC1* alleles encode sfGFP-tagged full-length ZC3HC1, and the mutant line in which all three alleles encode sfGFP-tagged ZC3HC1 Δ487–502. Separate donor templates encoding either full-length or Δ487–502 ZC3HC1, each followed by the sfGFP ORF, had been used to generate the respective cell lines; the larger PCR products thus document on-target integration of the corresponding donor construct into all three *ZC3HC1* alleles of each cell line. Note that the minor size difference between the PCR products representing the GFP-tagged full-length and Δ487–502 cell lines is due to the mutant being 81 nt shorter than the WT version, because of 48 deleted ZC3HC1 nucleotides and a linker between ZC3HC1 and sfGFP that is 33 nt shorter than in the WT version. **(C3)** IB of total cell extracts from the three cell lines described in C2. Immunolabelling was performed on the Ponceau S-stained membrane shown here, using an *Hs*ZC3HC1 antibody targeting aa 1–60. **(C4)** Triple-FM of co-cultured populations of HeLa P2 WT cells with non-tagged ZC3HC1 and HeLa P2 cells homozygously expressing ZC3HC1-sfGFP. Cells were additionally immunolabelled with ZC3HC1 and TPR antibodies, revealing no apparent differences in the NE-associated signal intensities for either ZC3HC1 or TPR. For corresponding IFM of co-cultured populations of HeLa P2 WT cells with non-tagged ZC3HC1 and HeLa P2 cells homozygously expressing ZC3HC1-sfGFP Δ487–502, see Figure 3B1. Bar, 10 µm. **(D)** Summary of experiments in which WT ZC3HC1 or deletion mutants thereof were ectopically expressed in a HeLa P2 ZC3HC1 KO cell line containing a notable nucleoplasmic pool of TPR. Schematics of ZC3HC1 and its deletion mutants are as in A. The capability of each ZC3HC1 version to bind to the NE and to recruit the nucleoplasmic TPR pool back to the NE is indicated separately (+, binding or recruitment; ~, uncertain or only traces of binding or recruitment; −, no binding or recruitment). Corresponding live-cell images for representative ZC3HC1 variants are shown in Figure 3E.

#### Supplemental Figure S4 (1/15)

**A1**

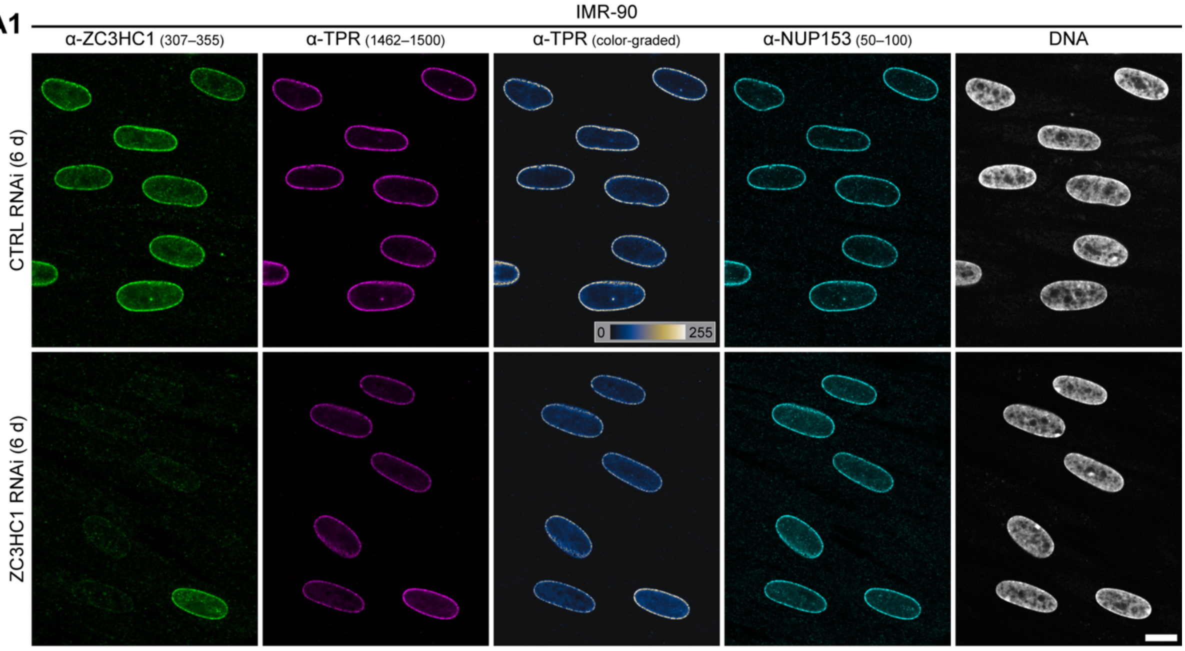


**A2**

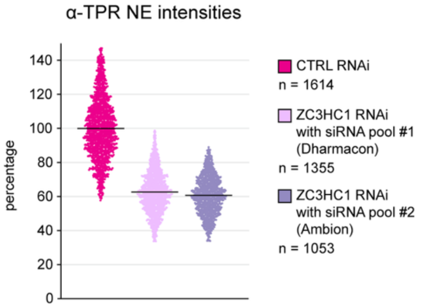

### Supplemental Figure S4 (2/15)

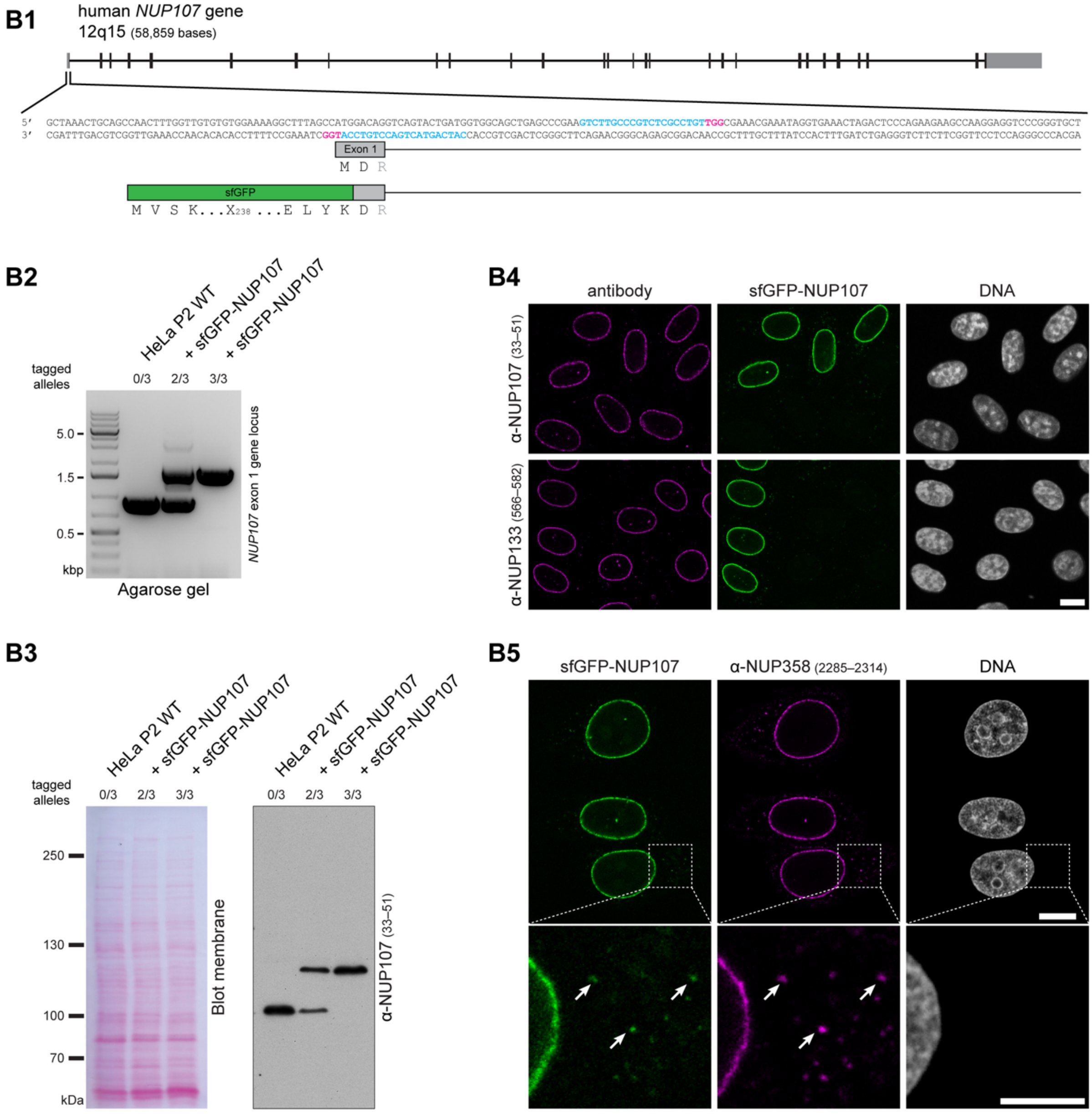

Supplemental Figure S4 (3/15)

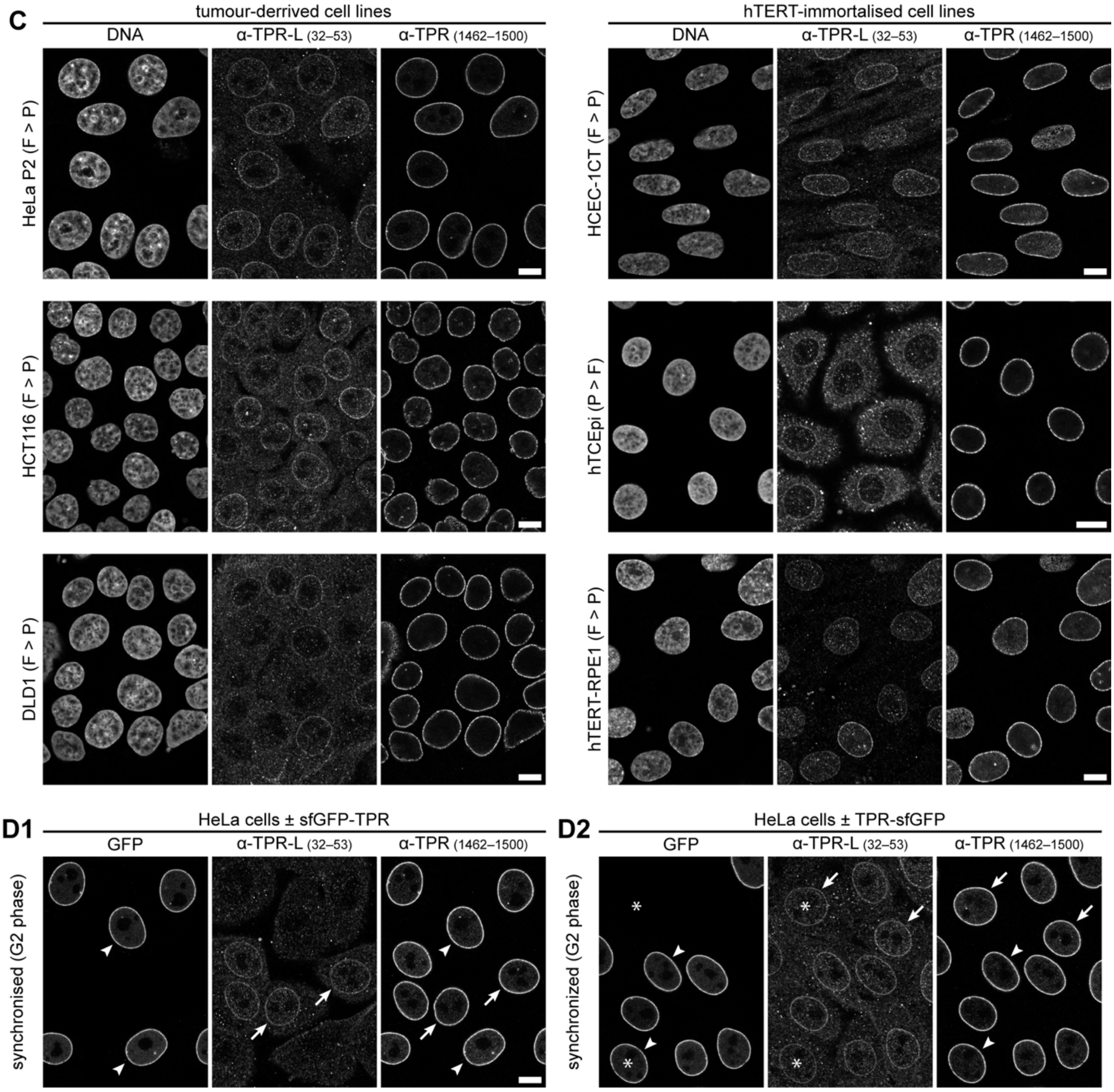

Supplemental Figure S4 (4/15)

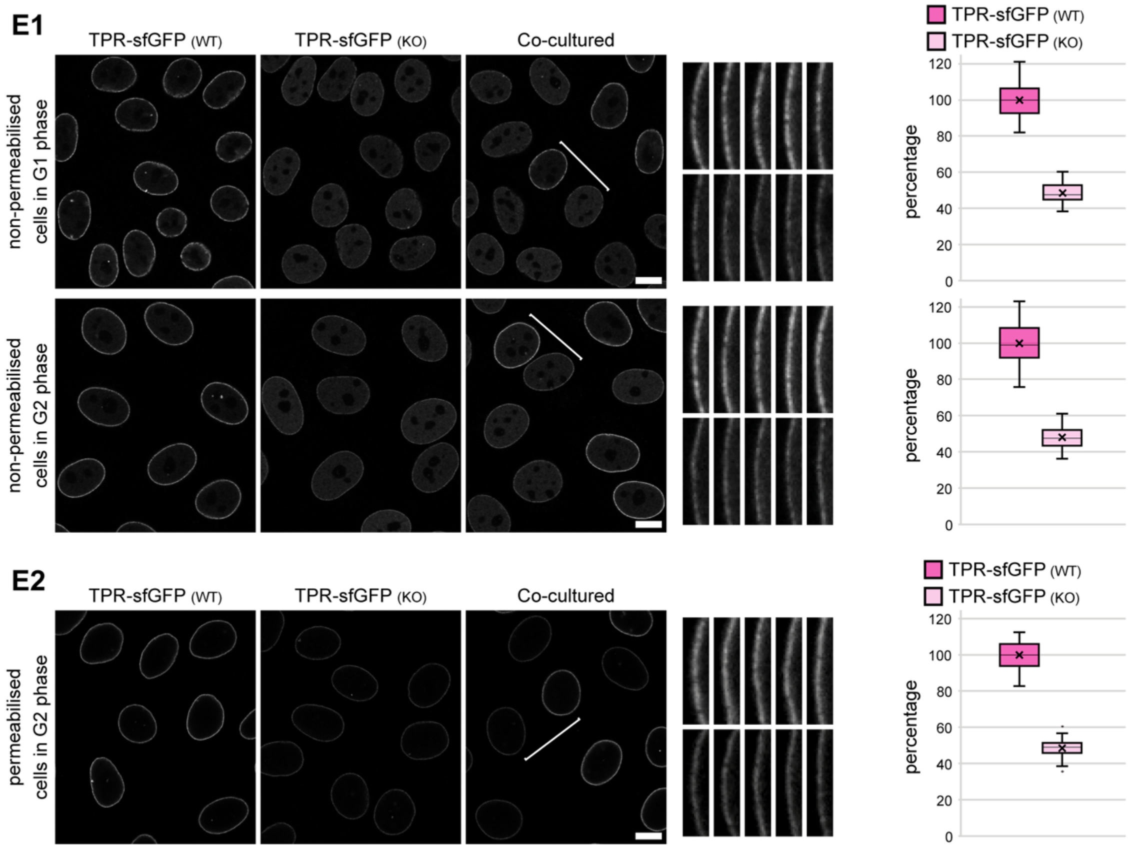

Supplemental Figure S4 (5/15)

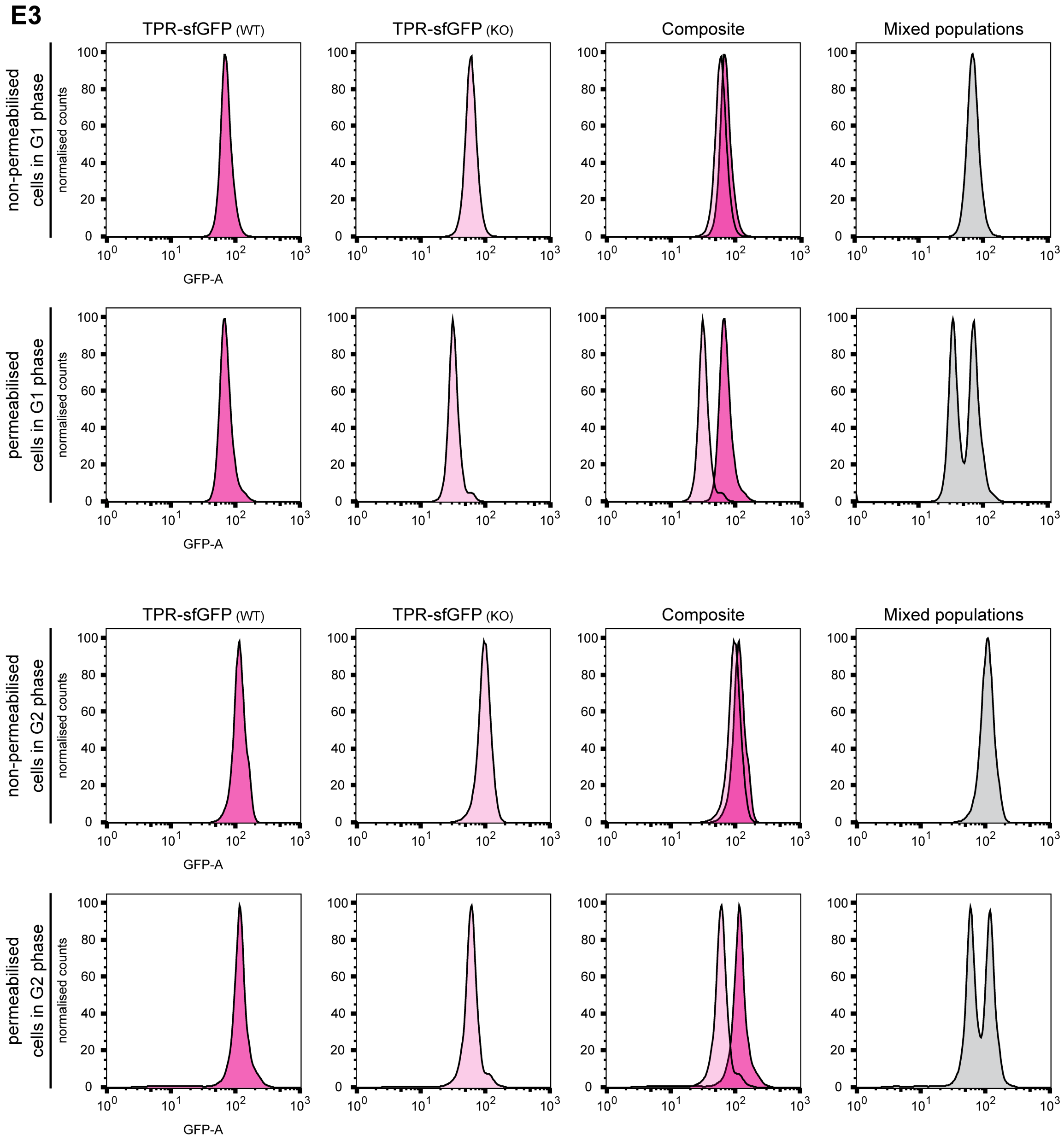

#### Supplemental Figure S4 (6/15)

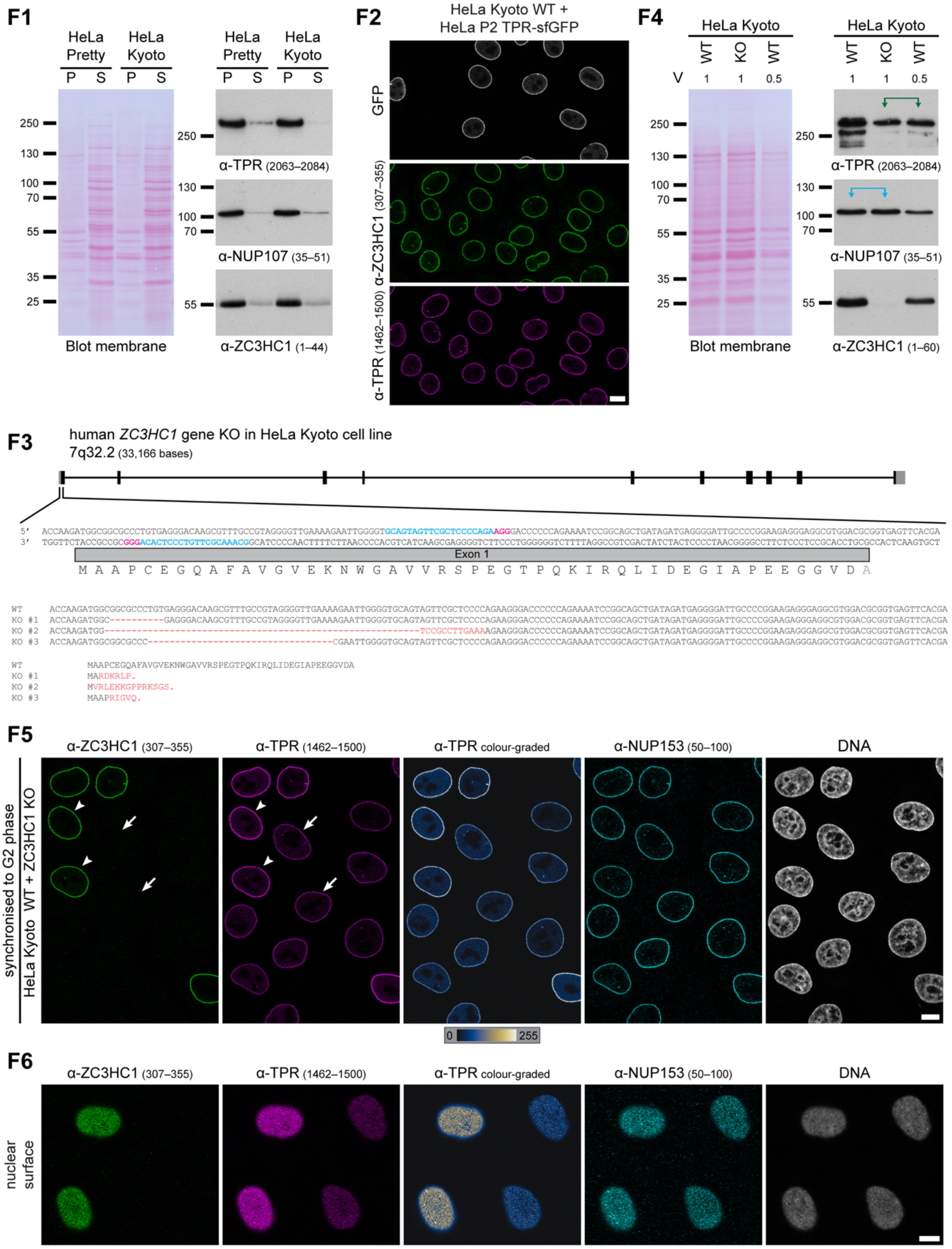

Supplemental Figure S4 (7/15)

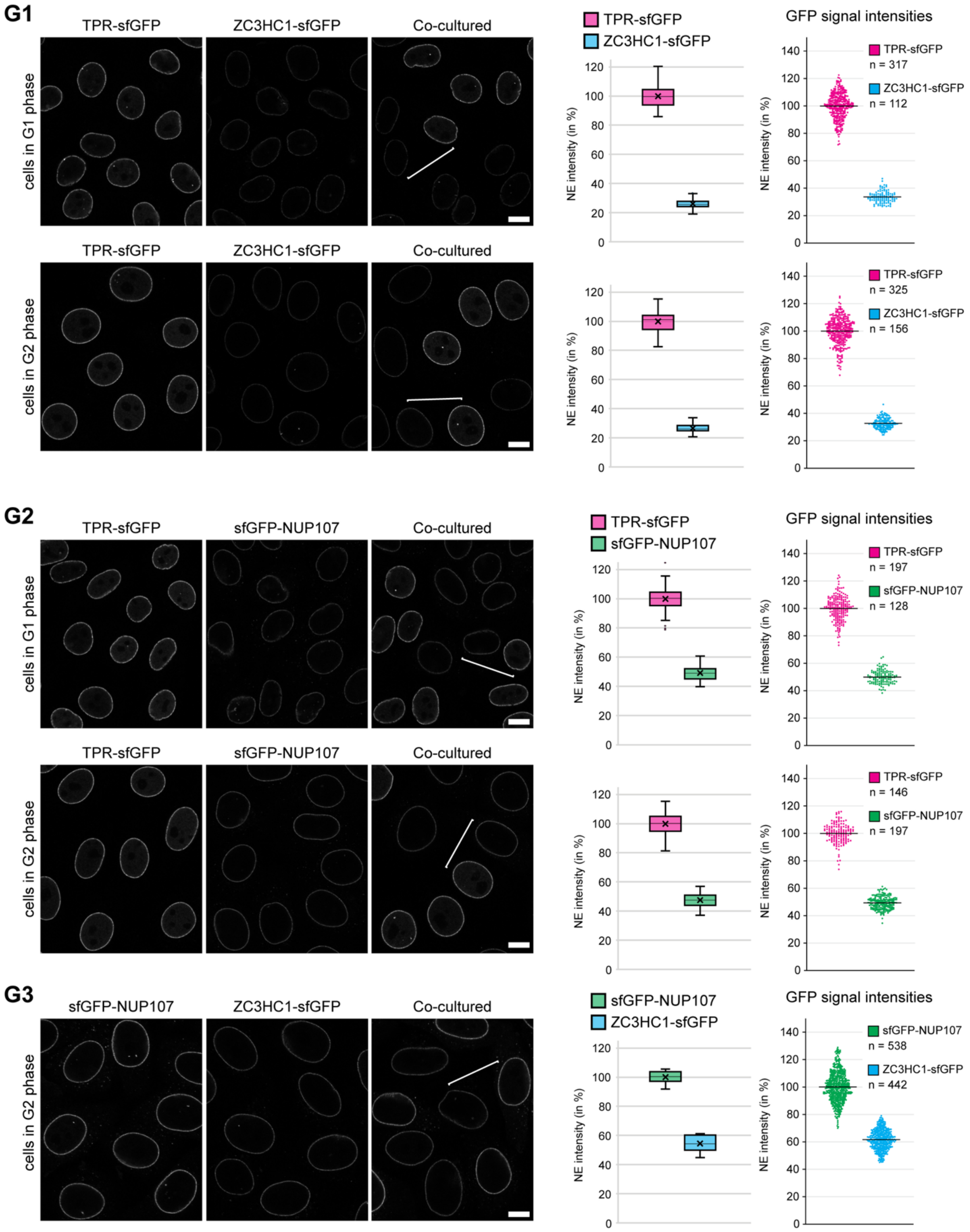

Supplemental Figure S4 (8/15)

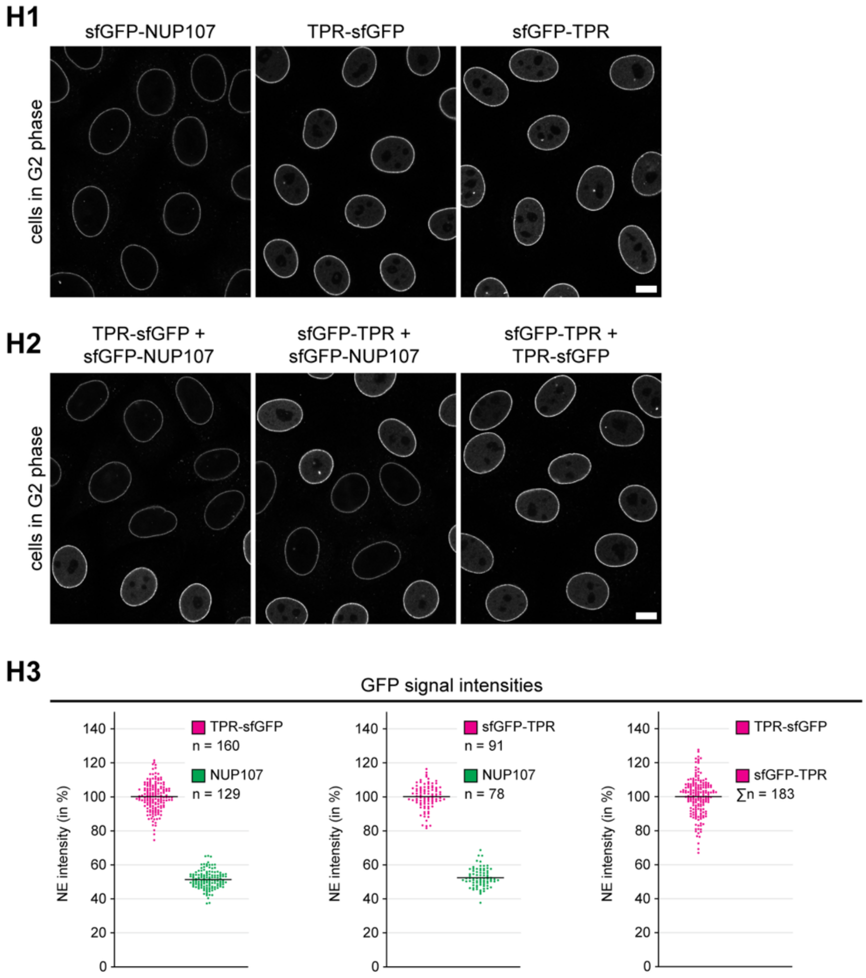

### Supplemental Figure S4 (9/15)

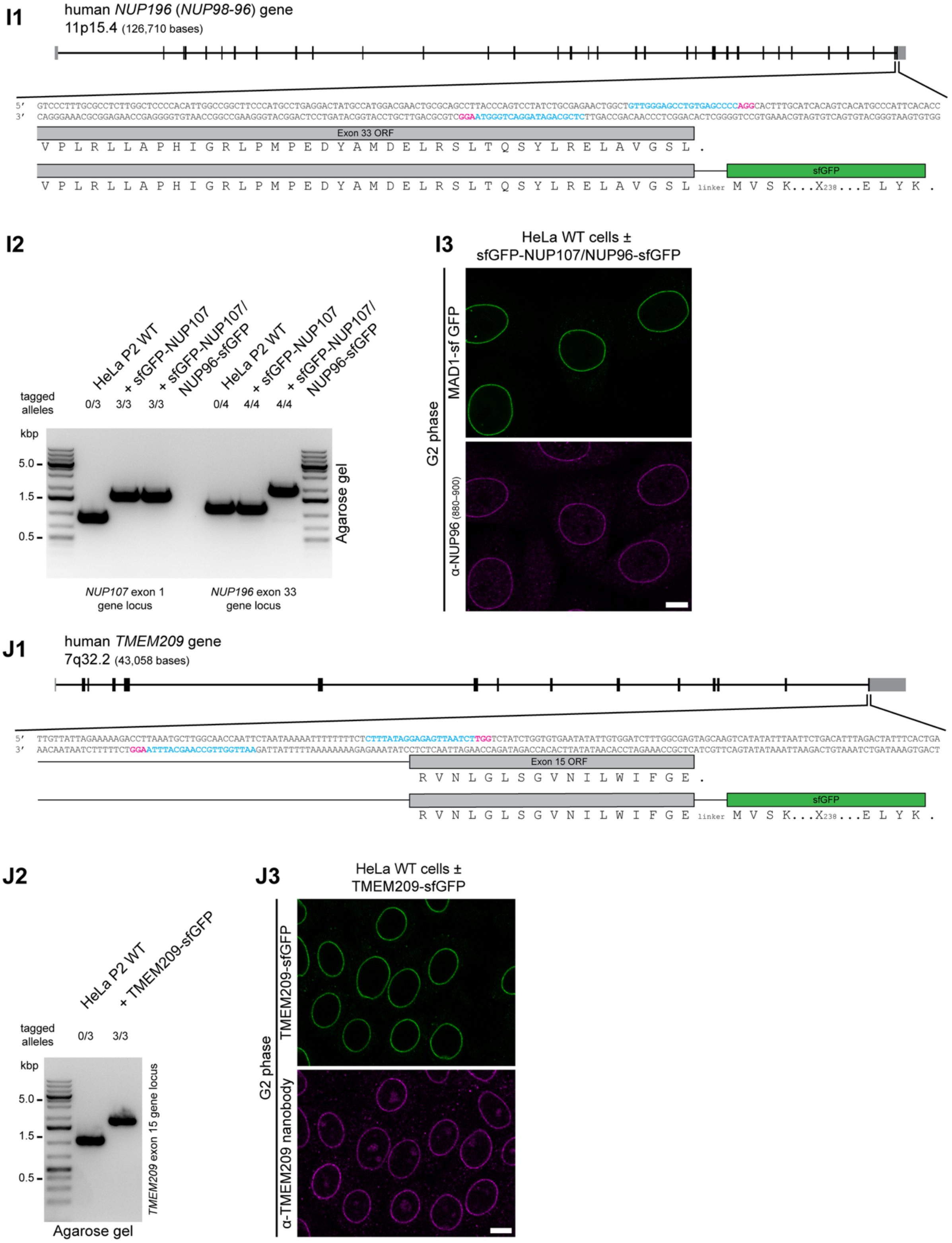

Supplemental Figure S4 (10/15)

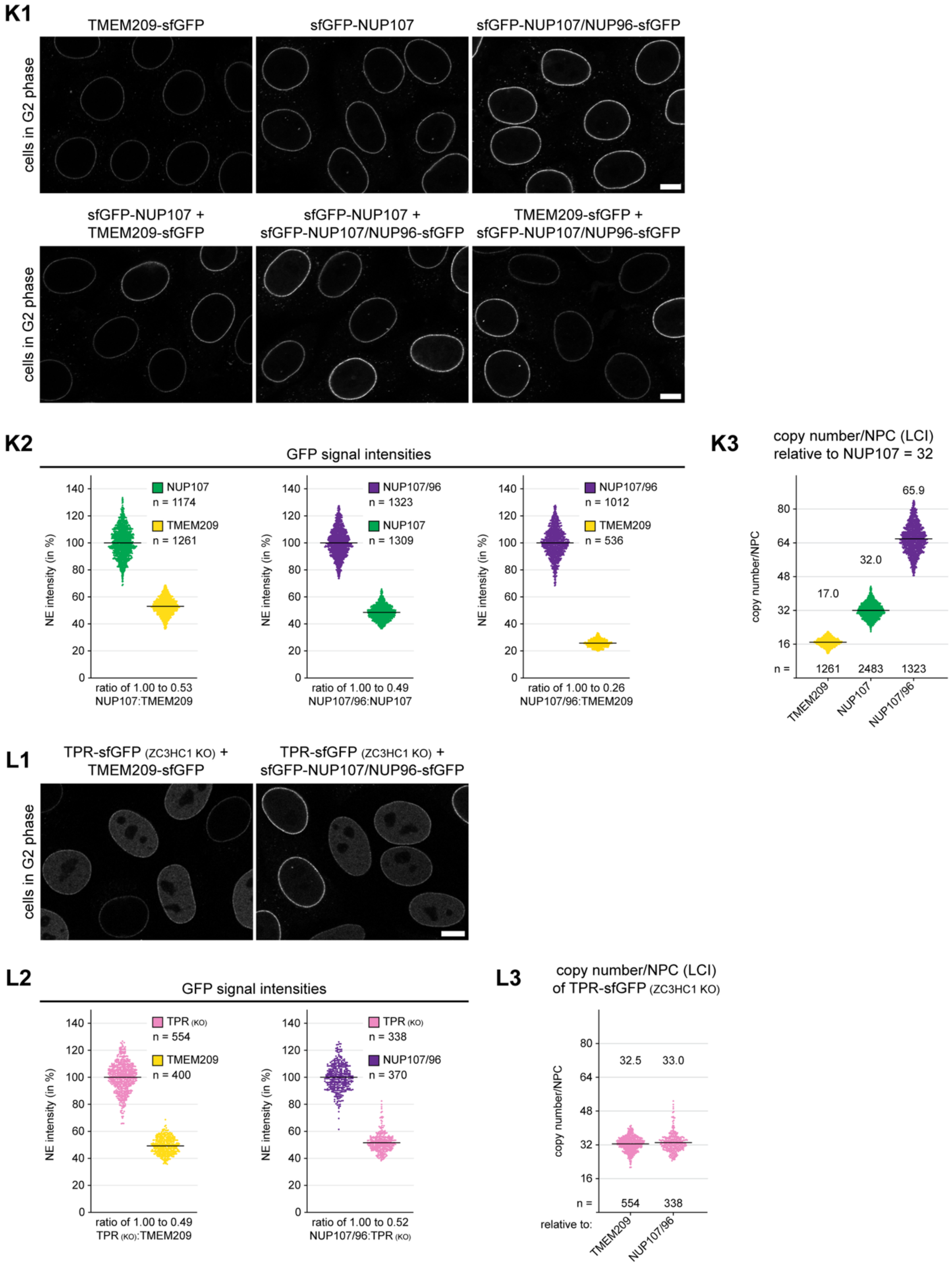

#### Supplemental Figure S4 (11/15)

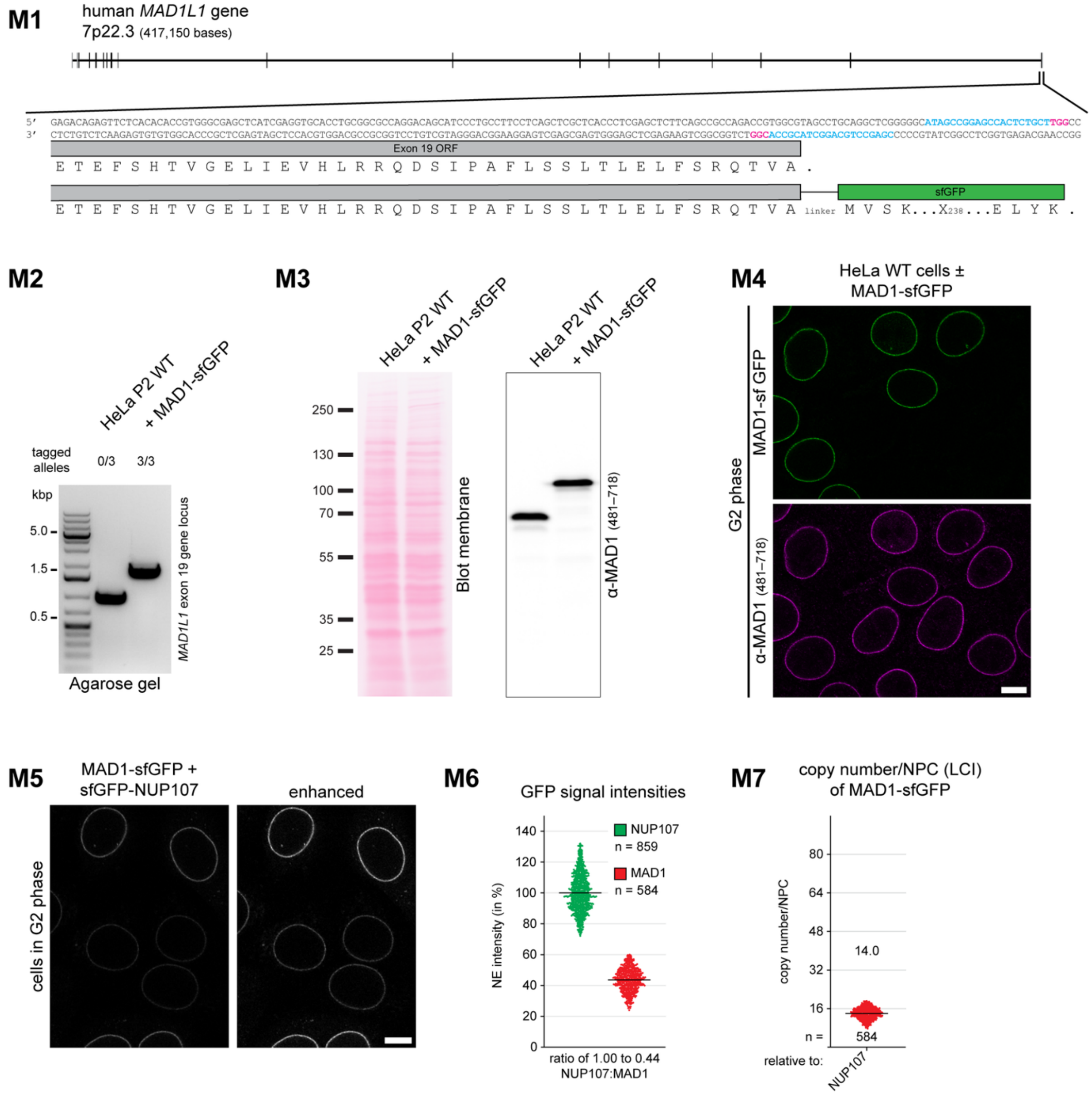

Supplemental Figure S4 (12/15)

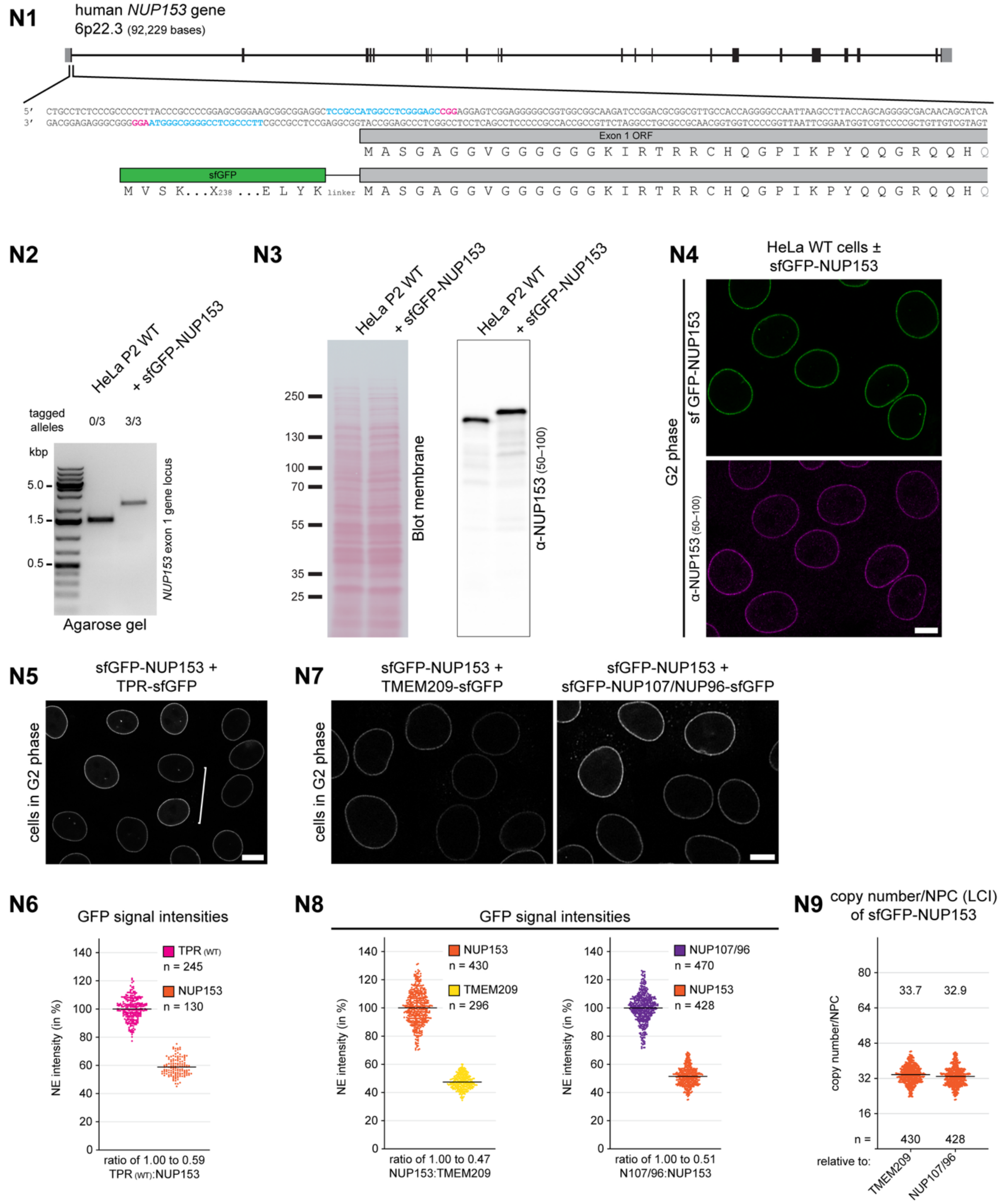

#### Supplemental Figure S4 (13/15)

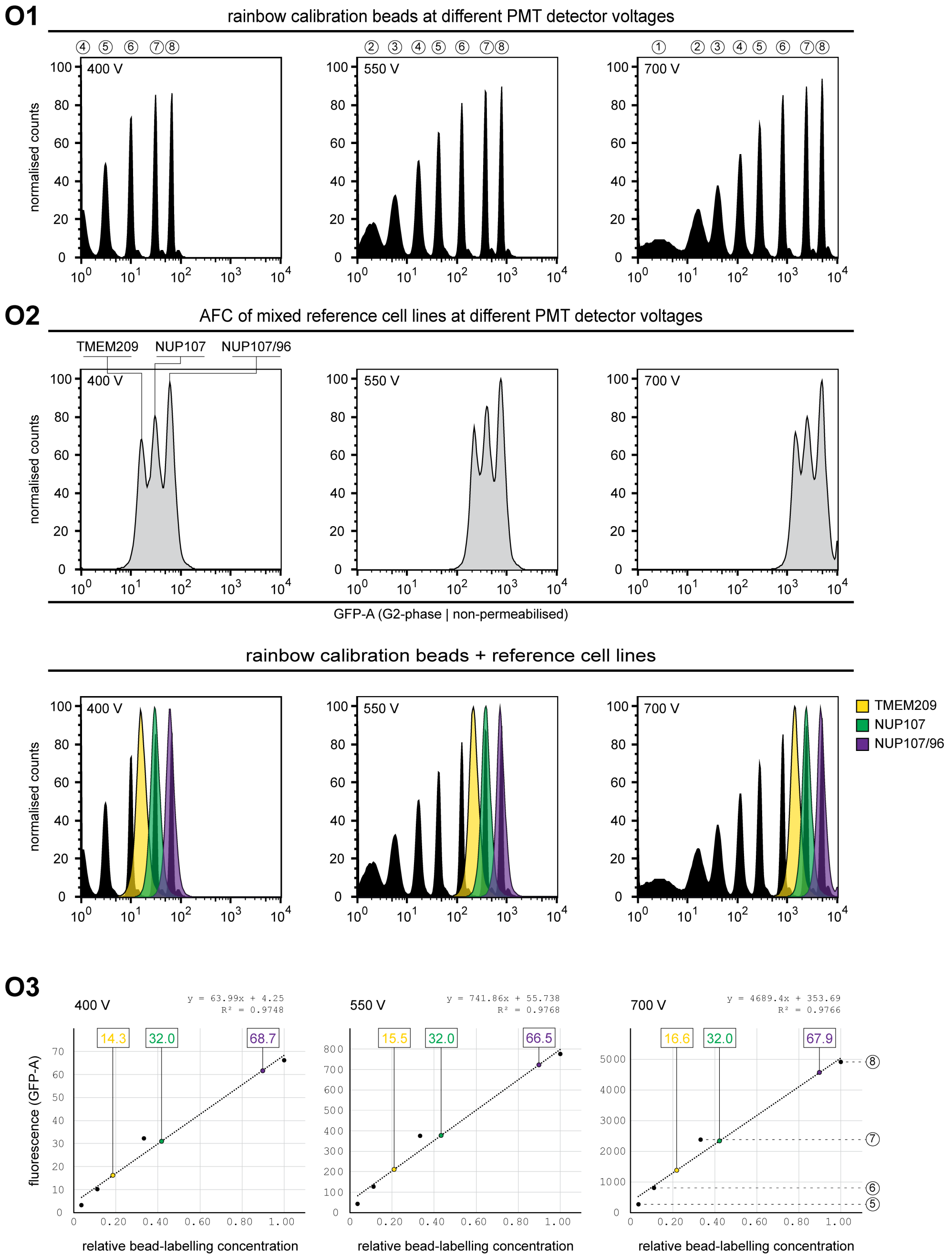

Supplemental Figure S4 (14/15)

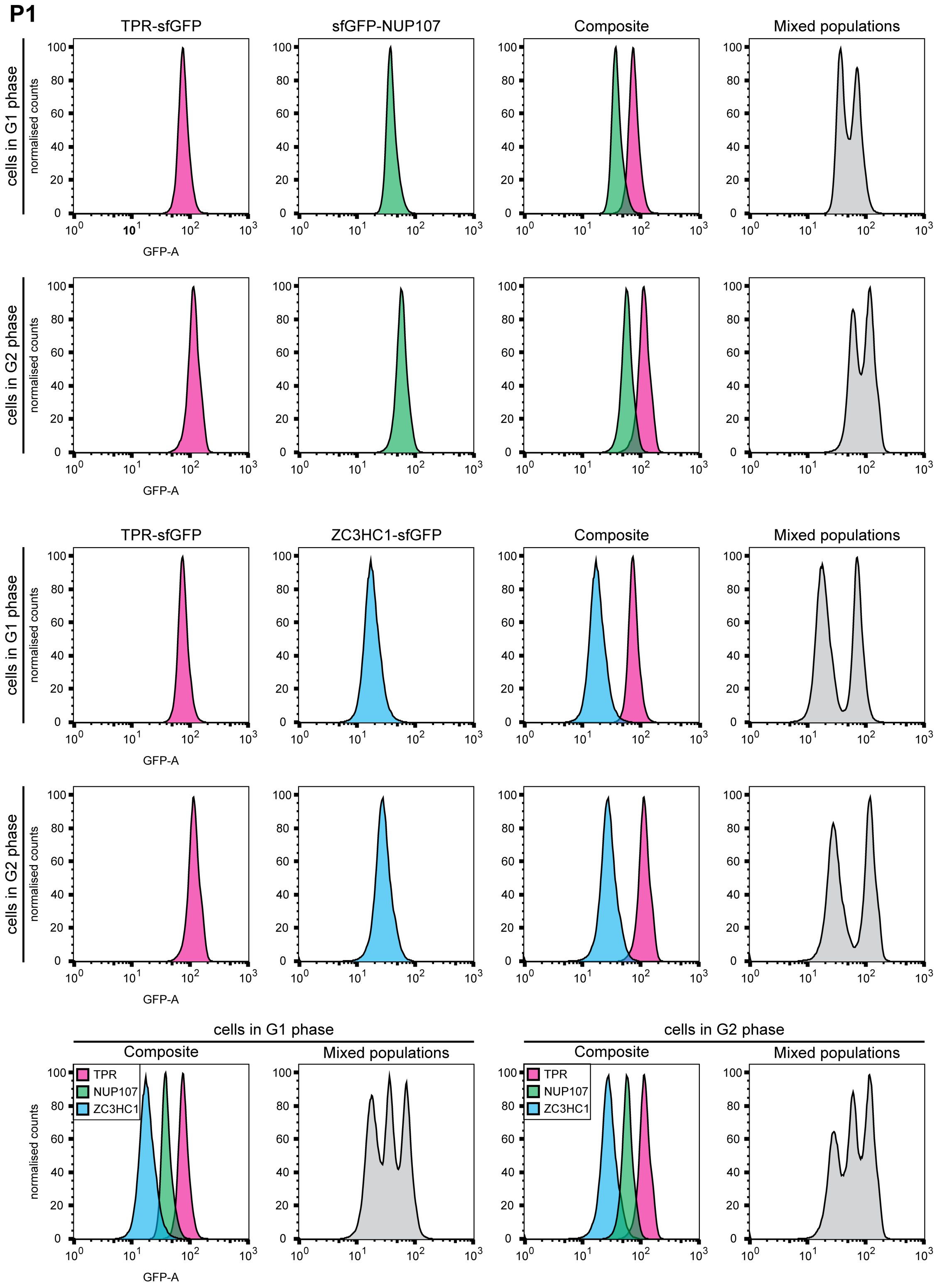

Supplemental Figure S4 (15/15)

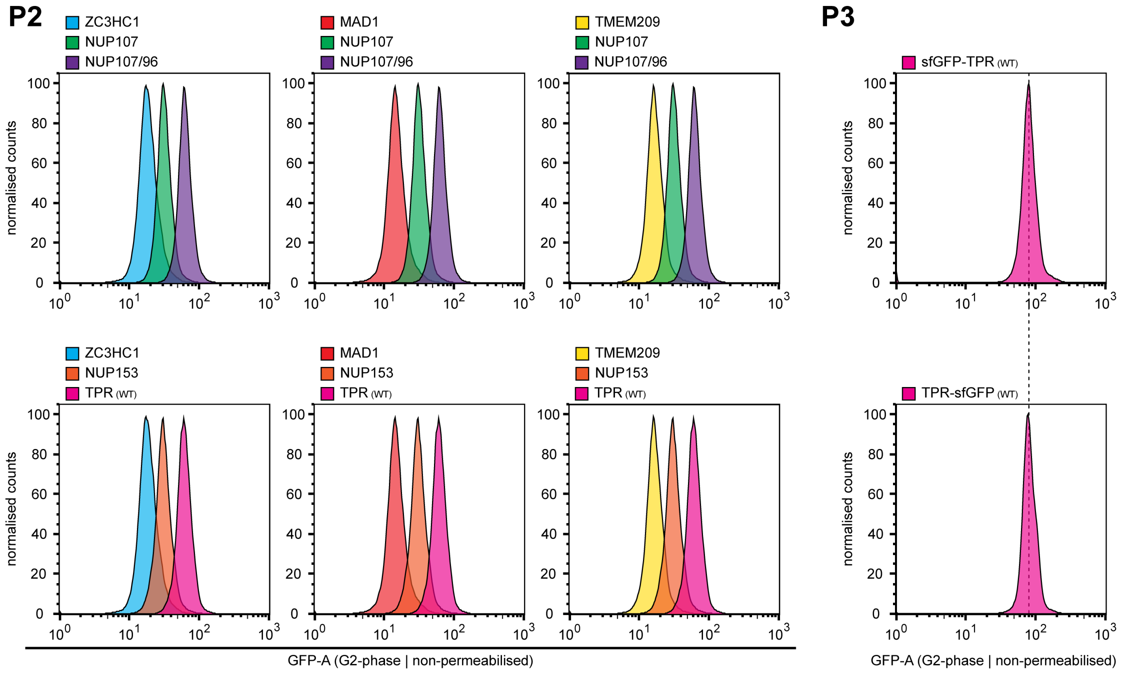


**Supplemental Figure S4. Extended fluorescence-based analyses of relative and absolute amounts of TPR, ZC3HC1, and other NPC- and NB-associated proteins, and characterisation of the GFP-tagged cell lines used for copy-number determination. (A)** RNAi-mediated depletion of ZC3HC1 in IMR-90 cells results in a substantial reduction of the NE-associated amounts of TPR. **(A1)** Representative IFM of IMR-90 cells treated with control siRNAs (CTRL) or siRNAs targeting ZC3HC1. Cells were triple-labelled with antibodies against ZC3HC1, TPR and NUP153. The micrograph for TPR is also shown colour-graded next to an LUT. Bar, 10 µm. **(A2)** Automated image-based quantification of NE-associated TPR immunofluorescence in IMR-90 cells treated with control siRNAs or two different sets of siRNAs targeting ZC3HC1. **(B)** Characterisation of a HeLa P2 cell line homozygously expressing sfGFP-NUP107. **(B1)** Schematic depiction of the human *NUP107* gene, the nucleotide sequence of exon 1 and its flanking regions, and the site of CRISPR/Cas9n-mediated insertion of the sfGFP ORF. The pair of single guide RNA (sgRNA) sequences used for site-specific knock-in are shown in blue lettering, except for the respective protospacer-adjacent motifs (PAMs), highlighted in magenta. In addition, the amino acids encoded by exon 1 of the non-tagged WT progenitor and some of those encoded by the sfGFP ORF are provided, as is the aa sequence of the sfGFP-NUP107 transition. **(B2)** Genotyping by PCR amplification. Note that the HeLa P2 cell line likely has three *NUP107* alleles, as in HeLa Kyoto (Landry *et al.*, 2013) and in the HeLa progenitor cell line CCL-2 (Adey *et al.*, 2013), while HeLa S3 may have three or four *NUP107* alleles (Adey *et al.*, 2013). PCR products derive from the original untagged progenitor HeLa P2 line (0/3), a heterozygous line in which two of the three *NUP107* alleles had been sfGFP-tagged (2/3), and a homozygous line in which all three *NUP107* alleles had been tagged (3/3). **(B3)** IB of total cell extracts from the three cell lines described in B2. Immunolabelling was performed on the Ponceau S-stained membrane shown here, using an epitope-specific *Hs*NUP107 antibody targeting aa 33–51. **(B4)** Double-FM of co-cultured populations of HeLa P2 WT cells with non-tagged NUP107 and HeLa P2 cells homozygously expressing sfGFP-NUP107. Immunolabelling with antibodies against NUP107 and NUP133, a direct binding partner of NUP107 within the Y-complex (e.g., Belgareh *et al.*, 2001; Berke *et al.*, 2004), revealed essentially indistinguishable NE signal intensities for NUP107 and NUP133 in both cell lines. Bar, 10 µm. **(B5)** IFM of HeLa P2 cells homozygously expressing sfGFP-NUP107. Immuno-labelling was performed with antibodies against NUP358, an NPC protein located at the NPC's cytoplasmic side (Wu *et al.*, 1995; Yokoyama *et al.*, 1995) and also at pore complexes of AL in different species and cell types (e.g., Cordes *et al.*, 1996, 1997a, 1997b), including tumour cell lines such as HeLa (e.g., Cordes *et al.*, 1997b; Lin *et al.*, 2026). In HeLa P2 cells, however, AL are only sporadically observed but can be identified by the co-localisation of sfGFP-NUP107 with numerous other NPC proteins, including the AL marker protein NUP358. In the magnified insets, some of these cytoplasmic co-localisations are marked by arrows. Bars, 10 µm (overview) and 5 µm (magnified ROI). **(C)** Expression of *Hs*TPR-L occurs in a wide range of human cell lines. Double-IFM of three tumour and three hTERT-immortalised cell lines with rabbit peptide antibodies against aa 32–53 of *Hs*TPR-L and a mAb targeting aa 1462–1500 of canonical *Hs*TPR, an epitope accessible in all NPC-associated subpopulations of *Hs*TPR. Cells were either fixed prior to permeabilisation (F > P) or permeabilised prior to fixation (P > F). While the NEs within a given cell population appear uniformly labelled with the pan-TPR antibody, the amounts of NE-associated TPR-L appear to differ notably between individual cells in some cell lines, such as HCT116 and hTERT-RPE1, and to a lesser extent in others. Bars, 10 µm. **(D)** TPR-L does not confound copy-number determinations based on sfGFP-tagged canonical TPR. **(D1)** N-terminal sfGFP-tagging of TPR at the canonical start codon abolishes detectable expression of TPR-L. Double-IFM of co-cultured populations of G2-synchronised HeLa P2 WT cells with non-tagged TPR and HeLa P2 cells homozygously expressing sfGFP-TPR. Cells were immunolabelled with the TPR mAb against aa 1462–1500 that recognises all TPR polypeptides and with antibodies specifically targeting TPR-L, revealing that TPR-L is no longer detectable in sfGFP-TPR-expressing cells. Two nuclei with sfGFP-TPR are marked by arrowheads, while two cells with non-tagged TPR are marked by arrows. Bar, 10 µm. **(D2)** Corresponding analysis of co-cultured populations of HeLa P2 cells with non-tagged TPR and cells with C-terminally tagged TPR-sfGFP. C-terminal tagging results in NE association of the sfGFP-tagged versions of both TPR-L and canonical TPR, with the mean amount ratio between TPR-L and canonical TPR remaining similar to that in HeLa P2 cells expressing non-tagged TPR. Arrows and arrowheads mark representative cells with non-tagged TPR and TPR-sfGFP, respectively. Asterisks mark one cell of each type, representing examples in which the NE-associated signal intensities for TPR-L differ to a similar extent from those of neighbouring cells with and without TPR-sfGFP. This further indicates that C-terminal sfGFP tagging does not noticeably alter the degree to which TPR-L polypeptides associate with the NB. Bar, 10 µm. **(E)** ZC3HC1 KO in HeLa P2 cells results in a substantial reduction of the NE-associated amounts of TPR, as visualised by fluorescence microscopy and AFC of cell cycle-synchronised HeLa WT and KO cells expressing TPR-sfGFP. **(E1)** LCI and quantification of TPR-sfGFP signal yields at the NEs of WT and ZC3HC1 KO cells, applying the manual quantification procedure described previously (Gunkel *et al.*, 2021; Supplemental Figure S2F). Cells were grown as single-type cell populations in separate wells, shown as reference, and as co-cultured populations in the same well. Specimens were imaged in parallel using identical microscope settings. Brackets mark exemplary pairs of WT and KO nuclei of the kind used for signal quantifications. Bars, 10 µm. Each quantification was based on 50 randomly chosen pairs of NE segments from neighbouring WT and KO cells. (**E2**) Corresponding analysis after detergent extraction to remove soluble TPR while retaining NB-associated TPR. **(E3)** AFC of HeLa WT cells and ZC3HC1 KO cells expressing TPR-sfGFP. Cells were analysed either as single-type cell populations, shown as magenta and light-magenta distribution plots together with the corresponding composite, or as a co-cultured population, shown in grey. The single-parameter histograms display the relative fluorescence of ~50,000 cells per cell line, with the respective peak values set to 100%. In addition, ~100,000 cells were analysed for the co-cultured populations. Cells were cell-cycle-synchronised and harvested in G1 or G2 and analysed either intact or after brief detergent permeabilisation to remove soluble TPR while retaining NB-associated TPR. Fluorescence intensities represent total amounts of emitted light (GFP-A), as holds for the AFC data shown in Figure 4. In intact cells, the fluorescence peak of the ZC3HC1 KO population reached ~85–88% of the WT value, resulting in only a minor offset between the peaks and preventing their separation in co-cultured populations. After detergent extraction, however, the residual NE-associated fluorescence peak of the KO cells amounted to only ~47–49% of that of the equally permeabilised WT cells, allowing both populations to be resolved. Note that the G2-synchronised co-cultured population plot is also shown in Figure 4I. **(F)** The relative amounts of NE-associated TPR, ZC3HC1, and NUP107 are similar in HeLa P2 and HeLa Kyoto cells. (**F1**) IB of the soluble proteins (S) released during extraction of HeLa P2 and HeLa Kyoto cells with TX-100 in NB-s buffer (Gunkel and Cordes, 2022), and of the corresponding insoluble pellet fractions (P), which include the lamina-NPC-NB (LNN)-enriched material (for further details, see Gunkel *et al.*, 2021). Together, the S and P fractions represent the entire protein content of a cell. Immunolabelling was performed on the Ponceau S-stained membrane shown here, revealing similar amount relationships between TPR, NUP107, and ZC3HC1 in both cell lines. **(F2)** Double-IFM of co-cultured populations of HeLa Kyoto WT cells with non-tagged TPR and HeLa P2 cells homozygously expressing TPR-sfGFP, revealing similar NE signal intensities for TPR and ZC3HC1 in both cell lines. **(F3)** Schematic depiction of part of the human *ZC3HC1* gene and the nucleotide and deduced aa sequences corresponding to exon 1, which was the target of CRISPR/Cas9n-conferred disruption in HeLa Kyoto cells. The sgRNA sequences are shown in blue lettering, with the respective PAMs highlighted in magenta. The genomic sequences of WT *ZC3HC1* and the disrupted alleles of the resulting HeLa Kyoto ZC3HC1 KO line, are shown for comparison, together with the corresponding aa sequences. The red-coloured sequences denote nucleotides and amino acids that are not part of the genuine ZC3HC1 sequence but result from rearrangements, insertions and frameshifts caused by CRISPR/Cas9n-editing. **(F4)** IB of LNN material obtained from HeLa Kyoto WT and ZC3HC1 KO cells harvested at ~90% confluency. For direct comparison, equal proportions of the adjusted LNN materials from WT and KO cells were loaded (1× V), alongside half this amount of the WT material (0.5× V). Immunolabelling was on the upper and lower parts of the membranes shown here. NUP107 signal intensities in the 1× V lanes, marked by an arrow-tipped bracket in blue, were essentially identical for the WT and KO cells. By contrast, TPR signal intensity in the KO amounted to at most about half of that in the WT and was similar to that in the WT 0.5× V lane, as indicated by the dark-green bracket. **(F5)** Triple-IFM of co-cultured HeLa Kyoto WT and ZC3HC1 KO cells. Arrowheads mark NEs of representative WT cells and arrows those of ZC3HC1 KO cells. NE-associated TPR signal intensities are approximately twofold higher in WT than in KO cells. Bar, 10 µm. **(F6)** Triple-IFM of neighbouring HeLa Kyoto WT and ZC3HC1 KO nuclei from a co-cultured population, imaged at their polar regions. While the NE-associated signal intensity for TPR is notably lower in the KO cell, NPC densities appear essentially indistinguishable between WT and KO cells. Bar, 10 µm. **(G)** GFP signal intensities at the NEs of cells expressing sfGFP-NUP107 are lower than those in cells with TPR-sfGFP but exceed those in cells with ZC3HC1-sfGFP. **(G1–G3)** LCI and quantification of GFP signal yields at the NEs of cells expressing **(G1)** TPR-sfGFP or ZC3HC1-sfGFP, **(G2)** TPR-sfGFP or sfGFP-NUP107, and **(G3)** sfGFP-NUP107 or ZC3HC1-sfGFP. Cells were grown as single-type populations in separate wells, shown as reference, and as co-cultured populations in the same well. Specimens were imaged in parallel using identical microscope settings. Brackets mark exemplary pairs of nuclei used for manual quantification. Box plots show the results of manual quantifications performed as described previously (Gunkel *et al.*, 2021) and in Supplemental Figure S2F, based on 50 randomly chosen pairs of neighbouring NE segments. Scatter plots show the results of automated quantifications as described in Supplemental Figure S2G. The numbers of nuclei included in the automated analyses are indicated in the plots. Bars, 10 µm. **(H)** NE-associated GFP signal intensities in cells expressing either sfGFP-TPR or TPR-sfGFP are essentially indistinguishable, indicating that NPC- and NB-association of TPR is largely unaffected by whether TPR is sfGFP-tagged at its N- or C-terminus. **(H1, H2)** LCI of cells grown as single-type populations (H1) or as co-cultured populations (H2). **(H3)** Scatter plots show the results of automated quantifications as described in Supplemental Figure S2G. The numbers of nuclei included in the automated analyses are indicated in the plots. Note that the automated NE detection software is not capable of distinguishing the two co-cultured cell lines expressing sfGFP-TPR and TPR-sfGFP cells from the LCI micrographs. Bar, 10 µm. **(I)** Characterisation of a HeLa P2 cell line homozygously expressing both sfGFP-NUP107 and NUP96-sfGFP. **(I1)** Schematic depiction of the human *NUP196* gene, part of the nucleotide sequence of the ORF of exon 33, terminated by the stop codon of *NUP196*, and the site of CRISPR/Cas9n-mediated insertion of the sfGFP ORF. The pair of sgRNA sequences used for site-specific knock-in are shown in blue lettering, with the respective PAMs highlighted in magenta. In addition, the amino acids encoded by the depicted part of exon 33 and some of those encoded by the sfGFP ORF are shown. The linker sequence between NUP96 and sfGFP, indicated by three dashes (-), reads GSAGSAAGSGTGS. NUP196 is the precursor of NUP98 and NUP96, which are generated by autoproteolytic cleavage (Fontoura *et al.*, 1999). CRISPR/Cas9n-mediated insertion of the sfGFP ORF into the *NUP196* alleles was performed in the HeLa P2 cell line already homozygously expressing sfGFP-NUP107, characterised in Supplemental Figure S4B. **(I2)** Genotyping by PCR amplification. Note that four *NUP196* alleles likely exist in the HeLa P2 cell line, as in HeLa Kyoto (Landry *et al.*, 2013), in the HeLa progenitor cell line CCL-2, and in HeLa S3 (Adey *et al.*, 2013). PCR products derive from the original untagged progenitor line (0/3 and 0/4) for the sfGFP-NUP107 cell line (see Supplemental Figure S4B), which was the progenitor for the cell line in which all *NUP96* alleles had been additionally sfGFP-tagged (3/3 and 4/4). **(I3)** Double-FM of co-cultured populations of HeLa P2 WT cells with non-tagged NUP107 and NUP96 and HeLa P2 cells homozygously expressing both sfGFP-NUP107 and NUP196-sfGFP. Immunolabelling with antibodies against NUP96 revealed essentially indistinguishable NE signal intensities for both NUPs in the two cell lines. Bar, 10 µm. **(J)** Characterisation of a HeLa P2 cell line homozygously expressing TMEM209-sfGFP. **(J1)** Schematic depiction of the human *TMEM209* gene, part of the nucleotide sequence of the ORF of exon 15, terminated by the stop codon of *TMEM209*, and the site of CRISPR/Cas9n-mediated insertion of the sfGFP ORF. The sgRNA sequences are shown in blue lettering, with the PAMs highlighted in magenta. In addition, the amino acids encoded by the depicted part of exon 15 and some of those encoded by the sfGFP ORF are shown. The linker sequence between

TMEM209 and sfGFP, indicated by three dashes (-), reads GSAGSAAGSGTG. **(J2)** Genotyping by PCR amplification. Note that three *TMEM209* alleles exist in the HeLa P2 cell line, as in HeLa Kyoto (Landry *et al.*, 2013), in the HeLa progenitor cell line CCL-2 and in HeLa S3 (Adey *et al.*, 2013). PCR products derive from the original untagged progenitor HeLa P2 line (0/3) and a homozygous line in which all *TMEM209* alleles had been tagged (3/3). **(J3)** Double-FM of co-cultured populations of HeLa P2 WT cells with non-tagged TMEM209 and HeLa P2 cells homozygously expressing TMEM209-sfGFP. Cells had been co-cultured and synchronized to G2-phase. After removal of the culture medium, the cells were permeabilised for 5 min with 60 µg/mL digitonin in PBS, followed by washings with PBS and blocking for 30 min in PBS with 1% BSA. Immunolabelling with a dye-coupled nanobody specific for *Hs*TMEM209 (clone BM10D02; Gräser, 2024), whose target-specificity was verified on a HeLa P2 TMEM209 KO cell line (our unpublished data), was for 60 min at RT. Following brief washes in PBS, cells were imaged immediately, revealing essentially indistinguishable NE signal intensities for TMEM209 in both cell lines. Bar, 10 µm. **(K)** Comparative LCI and quantitative analyses of the three reference cell lines reveals an approximately linear relationship between their copy numbers and NE-associated sfGFP signal intensities. **(K1)** LCI and automated quantification of GFP signal yields at the NEs of cells expressing TMEM209-sfGFP, sfGFP-NUP107 or both sfGFP-NUP107 and NUP96-sfGFP (sfGFP-NUP107/NUP96-sfGFP). Cells were grown as single-type populations in separate wells and as co-cultured populations in the same well. Specimens were imaged in parallel using identical microscope settings. Bars, 10 µm. (**K2**) Scatter plots show automated NE signal quantifications as described in Supplemental Figure S2G. **(K3)** Conversion of the mean NE-associated sfGFP signal intensities into copy-number equivalents. The mean value for sfGFP-NUP107 was assigned a copy number of 32 per NPC, and the corresponding mean fluorescence values of TMEM209-sfGFP and sfGFP-NUP107/NUP96-sfGFP were converted relative to this reference, yielding an approximate copy-number relationship of 16:32:64 for TMEM209:NUP107:NUP107/NUP96. **(L)** The NPC-associated amount of TPR in ZC3HC1 KO cells is approximately half the combined copy number of NUP107 and NUP96 and approximately twice that of TMEM209. **(L1)** LCI of ZC3HC1 KO cells expressing TPR-sfGFP, compared with WT cells expressing sfGFP-NUP107/NUP96-sfGFP or TMEM209-sfGFP. Cells were grown and imaged as described for K1. Bar, 10 µm. (**L2**) Scatter plots show automated NE signal quantifications as described in Supplemental Figure S2G. **(L3)** Conversion of the mean NE-associated sfGFP signal intensities into copy-number equivalents. The mean value was assigned either a copy number of 64 per NPC for sfGFP-NUP107/NUP96-sfGFP or a copy number of 16 per NPC for TMEM209-sfGFP, and the corresponding mean fluorescence values of TPR-sfGFP in ZC3HC1 KO cells were converted relative to these two reference values. **(M)** Characterisation of a HeLa P2 cell line homozygously expressing MAD1-sfGFP. **(M1)** Schematic depiction of the human *MAD1L1* gene, part of the nucleotide sequence of the ORF of exon 19, terminated by the stop codon of *MAD1L1*, and the site of CRISPR/Cas9n-mediated insertion of the sfGFP ORF. The sgRNA sequences are shown in blue lettering, with the PAMs highlighted in magenta. In addition, the amino acids encoded by the depicted part of exon 19 and some of those encoded by the sfGFP ORF are shown. The linker sequence between MAD1 and sfGFP, indicated by three dashes (-), reads GSAGSAAGSGTGS. **(M2)** Genotyping by PCR amplification. Note that three *MAD1L1* alleles exist in the HeLa P2 cell line, as in HeLa Kyoto (Landry *et al.*, 2013), in the HeLa progenitor cell line CCL-2, and in HeLa S3 (Adey *et al.*, 2013).. PCR products derive from the original untagged progenitor line (0/3) and a homozygous line in which all *MAD1L1* alleles had been tagged (3/3). **(M3)** IB of total cell extracts from cell lines described in M2. Immunolabelling was performed on the Ponceau S-stained membrane shown here, using an epitope-specific *Hs*MAD1 antibody targeting aa 481–718. **(M4)** Double-FM of co-cultured populations of HeLa P2 WT cells with non-tagged MAD1 and HeLa P2 cells homozygously expressing MAD1-sfGFP. Immunolabelling revealed essentially indistinguishable NE signal intensities for MAD1 in both cell lines. Bar, 10 µm. **(M5)** LCI of cells expressing MAD1-sfGFP or sfGFP-NUP107. Cells were grown and imaged as described for K1. Bar, 10 µm. (**M6**) Scatter plots show automated NE signal quantifications as described in Supplemental Figure S2G. **(M7)** Conversion of the mean NE-associated sfGFP signal intensities into copy-number equivalents. The mean value was assigned a copy number of 32 per NPC for sfGFP-NUP107 and the corresponding mean fluorescence values of MAD1-sfGFP cells were converted relative to this reference. **(N)** Characterisation of a HeLa P2 cell line homozygously expressing sfGFP-NUP153. **(N1)** Schematic depiction of the human *NUP153* gene, part of the nucleotide sequence of the ORF of exon 1, and the site of CRISPR/Cas9n-mediated insertion of the sfGFP ORF. The sgRNA sequences are shown in blue lettering, with the PAMs highlighted in magenta. In addition, some of the amino acids encoded by exon 1 and some of those encoded by the sfGFP ORF are shown. The linker sequence between sfGFP and NUP153, indicated by three dashes (-), reads GSAGSAAGSG. **(N2)** Genotyping by PCR amplification. Note that three *NUP153* alleles likely exist in the HeLa P2 cell line, as in HeLa Kyoto (Landry *et al.*, 2013), in the HeLa progenitor cell line CCL-2 and in HeLa S3 (Adey *et al.*, 2013). PCR products derive from the original untagged progenitor line (0/3) and a homozygous line in which all *NUP153* alleles had been tagged (3/3). **(N3)** IB of total cell extracts from untagged HeLa P2 cells and the cell lines described in N2. Immunolabelling was performed on the Ponceau S-stained membrane shown here, using an epitope-specific *Hs*NUP153 antibody targeting aa 50–100. **(N4)** Double-FM of co-cultured populations of HeLa P2 WT cells with non-tagged NUP153 and HeLa P2 cells homozygously expressing sfGFP-NUP153. Immunolabelling revealed essentially indistinguishable NE signal intensities for NUP153 in both cell lines. Bar, 10 µm. **(N5)** LCI of cells expressing sfGFP-NUP153 or TPR-sfGFP. Cells were grown and imaged as described for Figure B–G. **(N6)** Automated quantification of GFP signal yields at the NEs of cells imaged for N5. Scatter plots show automated NE signal quantifications as described in Supplemental Figure S2G. Bar, 10 µm. **(N7)** LCI of cells expressing sfGFP-NUP153, compared with cells expressing TMEM209-sfGFP or sfGFP-NUP107/NUP96-sfGFP. Cells were grown and imaged as described for K1. Bar, 10 µm. (**N8**) Scatter plots show automated NE signal quantifications as described in Supplemental Figure S2G. **(N9)** Conversion of the mean NE-associated sfGFP signal intensities into copy-number equivalents. The mean value was assigned either a copy number of 64 per NPC for sfGFP-NUP107/NUP96-sfGFP or a copy number of 16 per NPC for TMEM209-sfGFP, and the corresponding mean fluorescence values of sfGFP-NUP153 cells were converted relative to these two reference values. **(O)** AFC settings applied in the current study provided a linear detection range spanning both fluorescent calibration beads and the fluorescence signals of the different reference cell lines. **(O1)** AFC of ProLine Rainbow Beads (BioRad) comprising eight bead populations (1–8). Seven populations (2–8) had been prepared with serially diluted amounts of the same fluorescent dye stock (1×, 1/3, 1/9, 1/27, 1/81, 1/243, and 1/729), while the dimmest population (1) had been left undyed. AFC was performed with three different settings, applying PMT voltages of 400 V, 550 V and 700 V, respectively. **(O2)** AFC of HeLa P2 reference cell lines homozygously expressing either TMEM209-sfGFP, sfGFP-NUP107, or sfGFP-NUP107/NUP96-sfGFP, here analysed as a co-cultured population of all three cell lines (upper panel) or with beads and cell lines separately analysed on the same day using identical AFC settings (lower panel). Fluorescence intensities represent total amounts of emitted light (GFP-A). **(O3)** Combined representation of the AFC data shown in O1 and O2. The y-axis indicates measured fluorescence values, while the x-axis indicates the relative bead-labelling concentration (1/dilution factor). For clarity, only the four brightest bead populations (5–8) are shown here (black dots), together with the corresponding linear regression and its equation.

This equation was then applied to the measured fluorescence values of the three reference cell lines to derive corresponding bead-equivalent values on the x-axis, i.e. the relative bead-labelling concentrations that, according to the regression, would yield the measured GFP fluorescence values. The resulting coordinates are shown on the regression line as coloured dots for TMEM209-sfGFP (yellow), sfGFP-NUP107 (green) and sfGFP-NUP107/NUP96-sfGFP (purple). Next, the measured fluorescence value of sfGFP-NUP107 was assigned the established copy number of 32 polypeptides per NPC. Relative to this reference number, the measured fluorescence values of TMEM209-sfGFP and sfGFP-NUP107/NUP96-sfGFP were converted into corresponding sfGFP copy-number equivalents per NPC (boxed numbers in colour). **(P)** Compilation of AFC distribution plots for cell lines analysed either as single-type cell populations, shown as coloured distributions together with the corresponding composite data, or as co-cultured populations, shown in grey. G1- and G2-synchronised cells were analysed as described in E3. Fluorescence intensities again represent total amounts of emitted light (GFP-A). Each group of experiments presented here as P1, P2, and P3 was conducted on a single day under identical conditions, while the conditions differed slightly between P1, P2, and P3.

#### Supplemental Figure S5 (1/6)

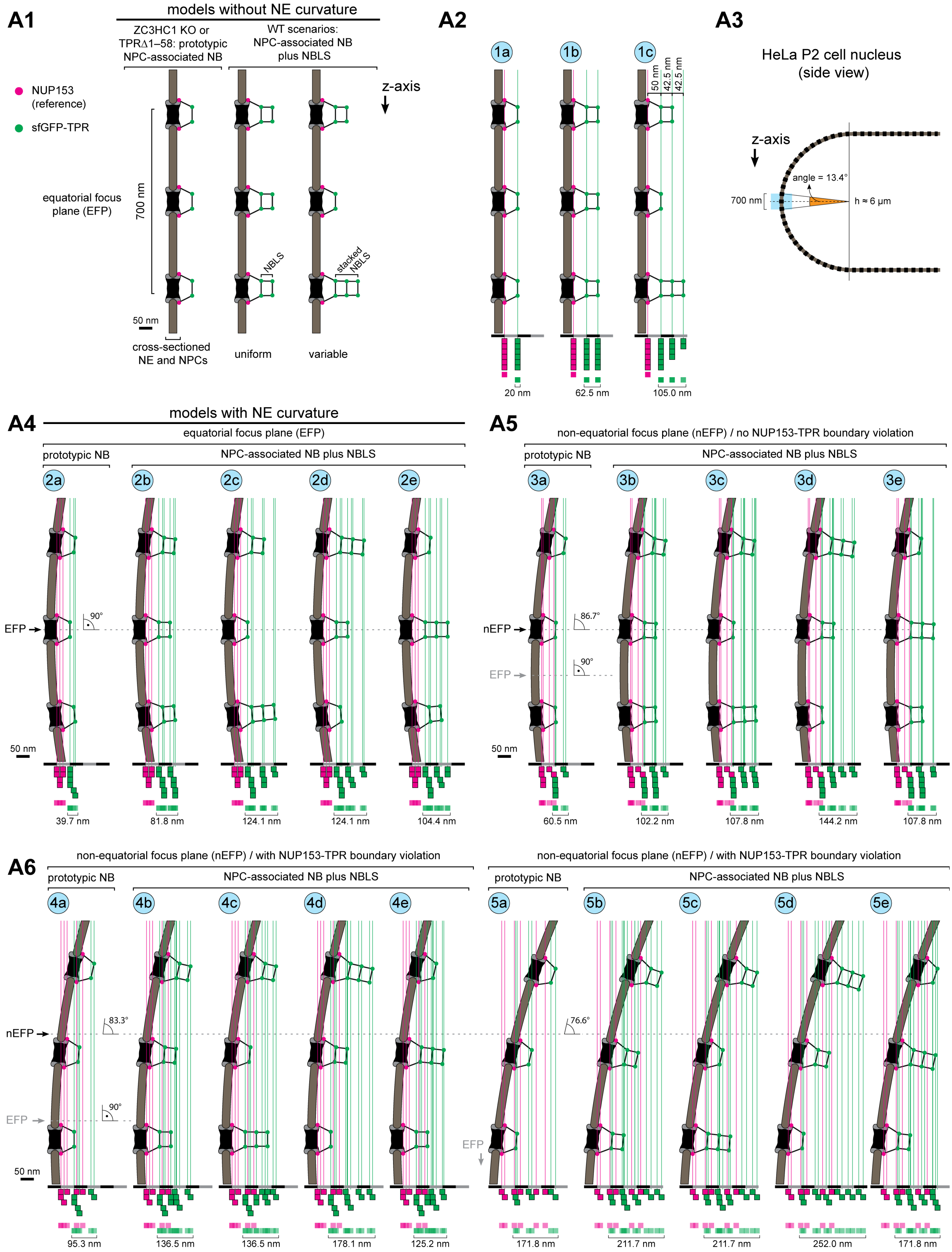

Supplemental Figure S5 (2/6)

**A7**

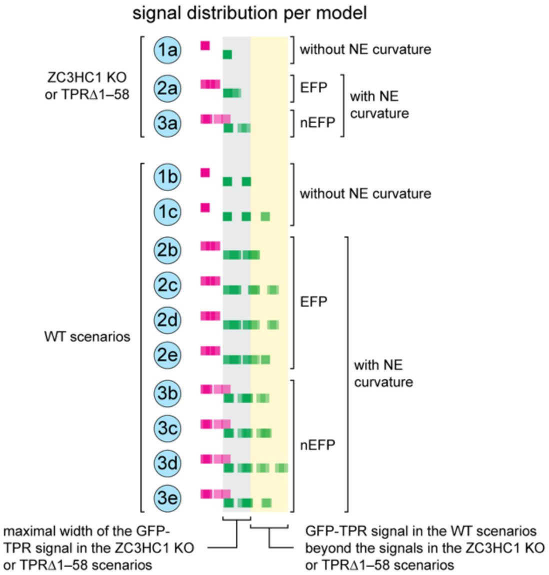


**A8**

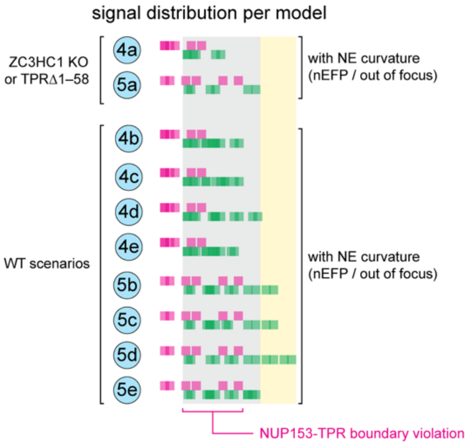


**B1**

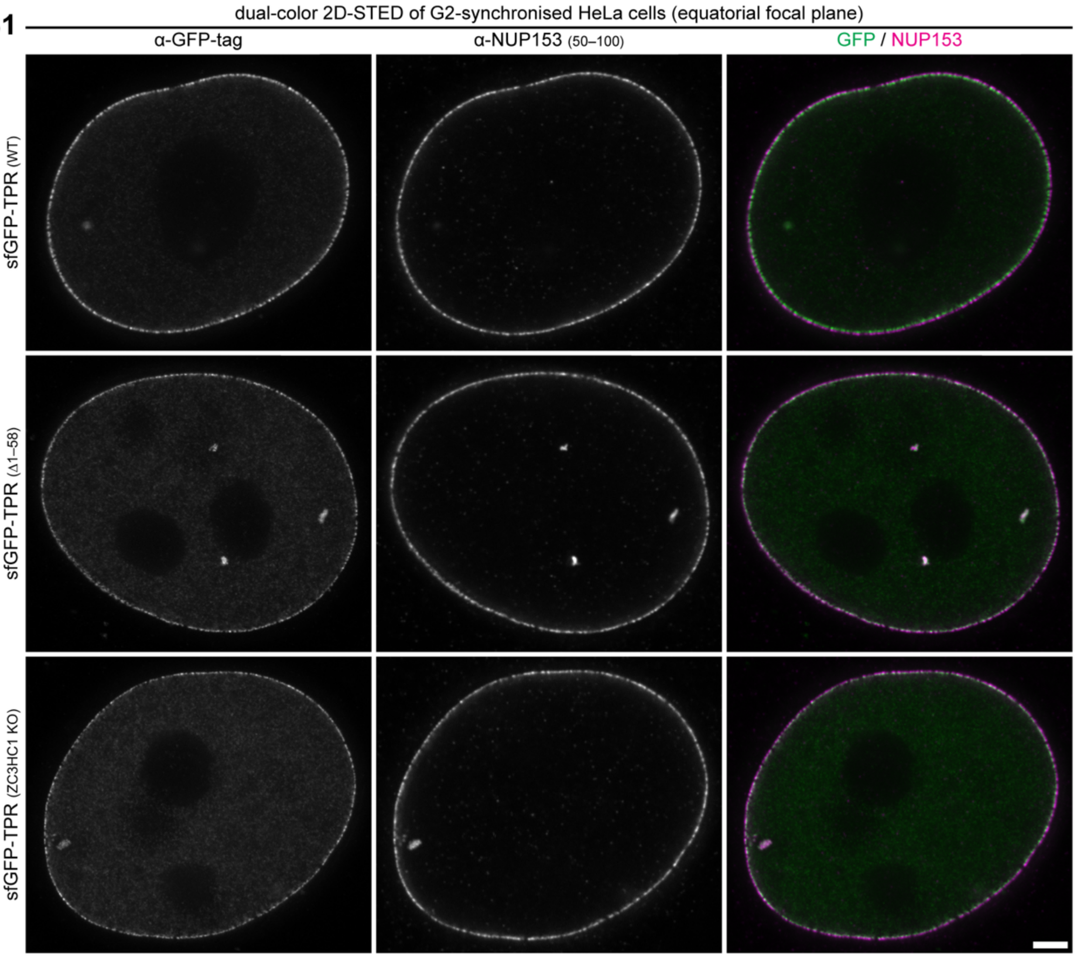

Supplemental Figure S5 (3/6)

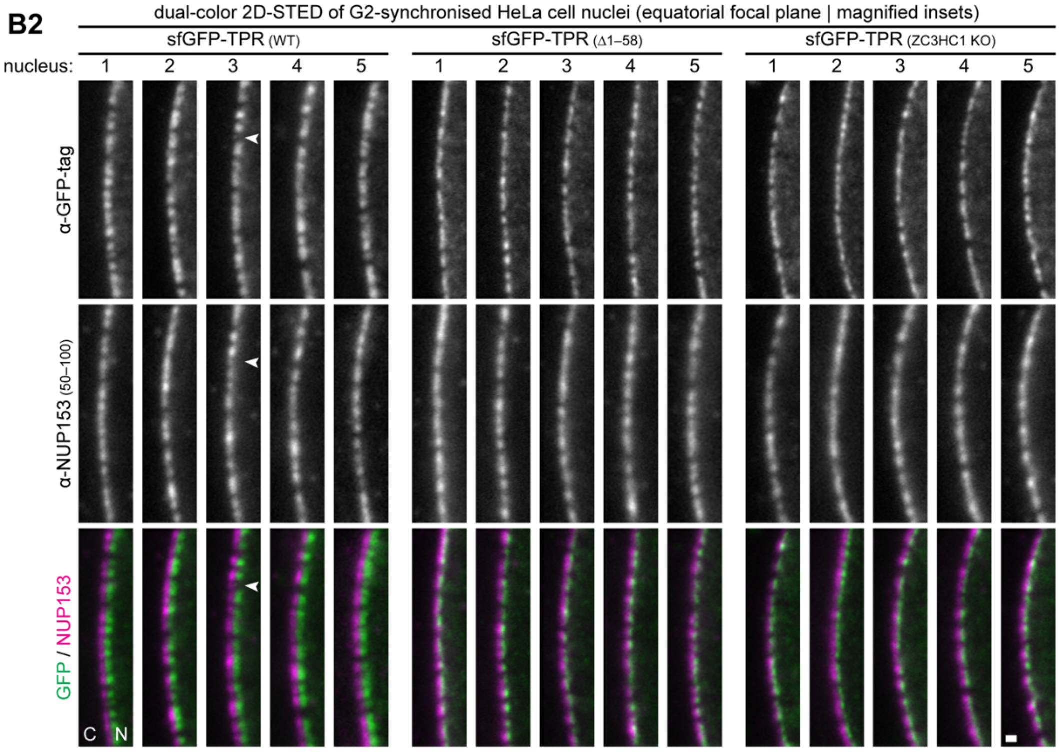

Supplemental Figure S5 (4/6)

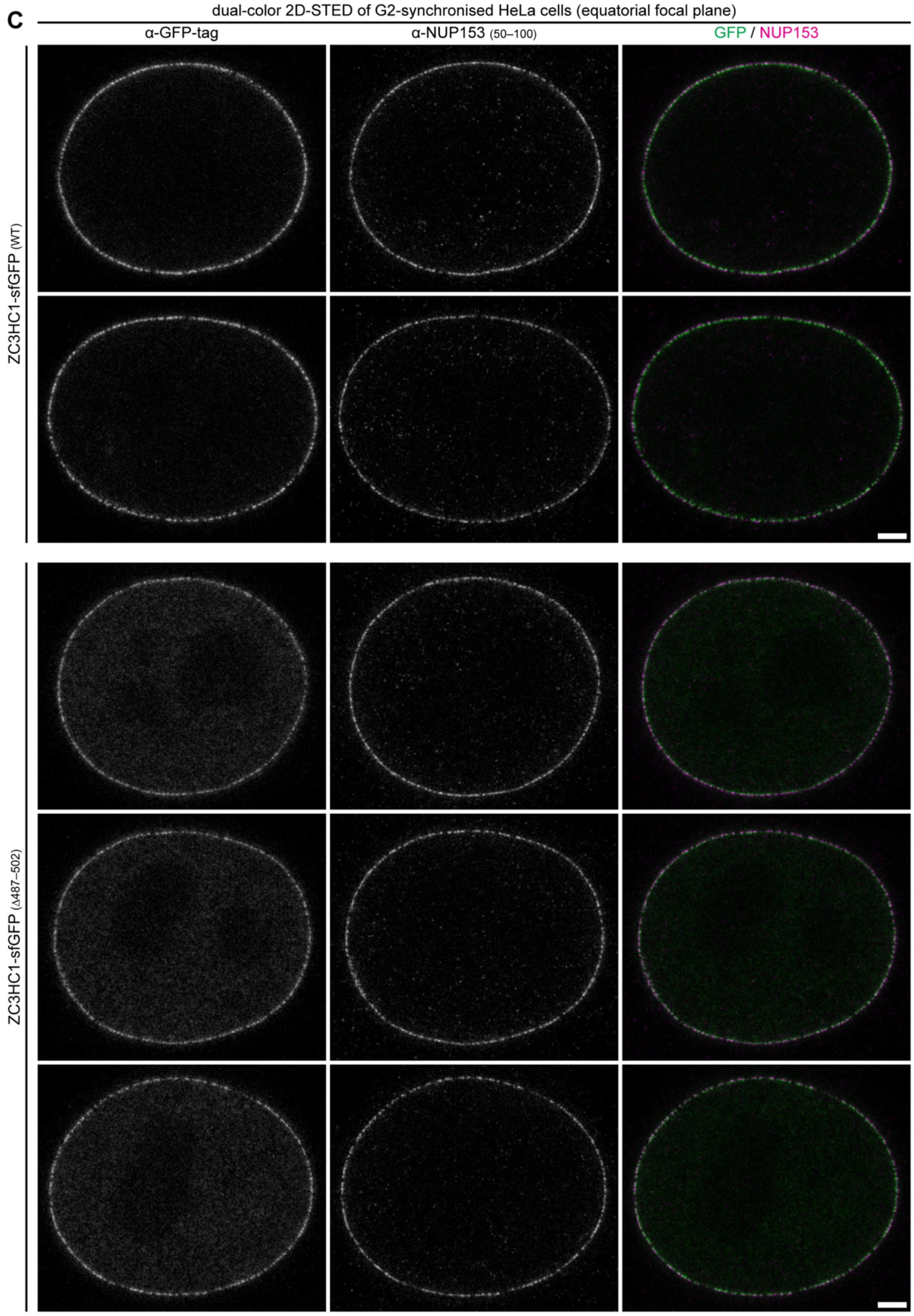

#### Supplemental Figure S5 (5/6)

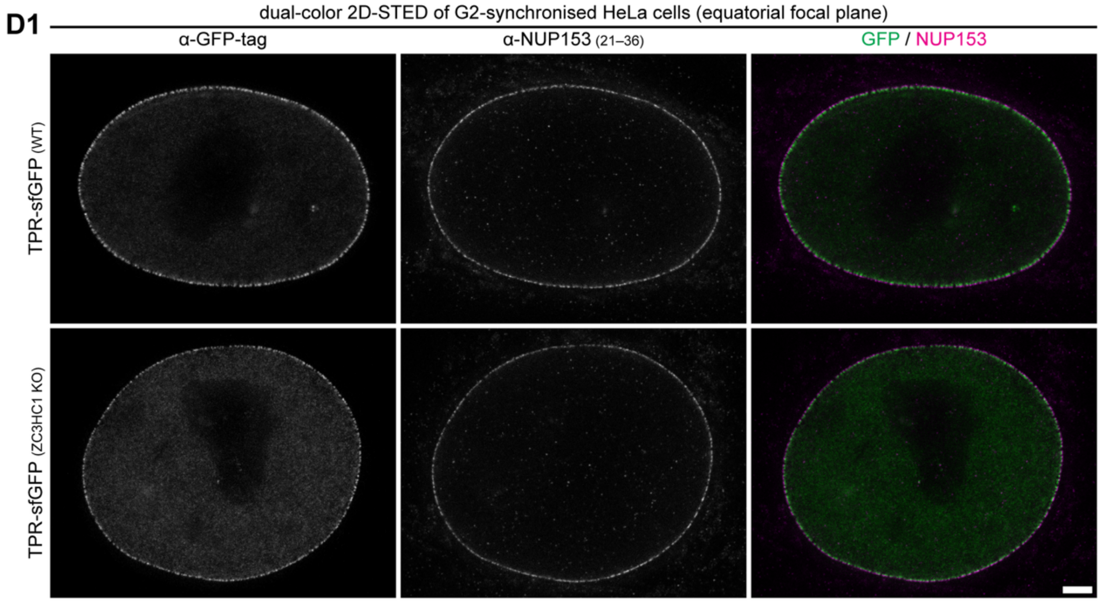


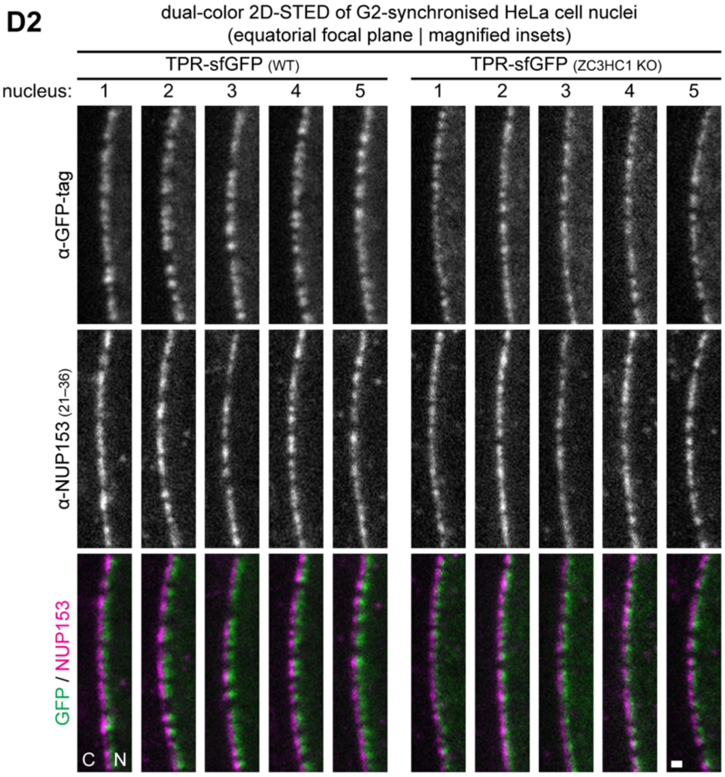

Supplemental Figure S5 (6/6)

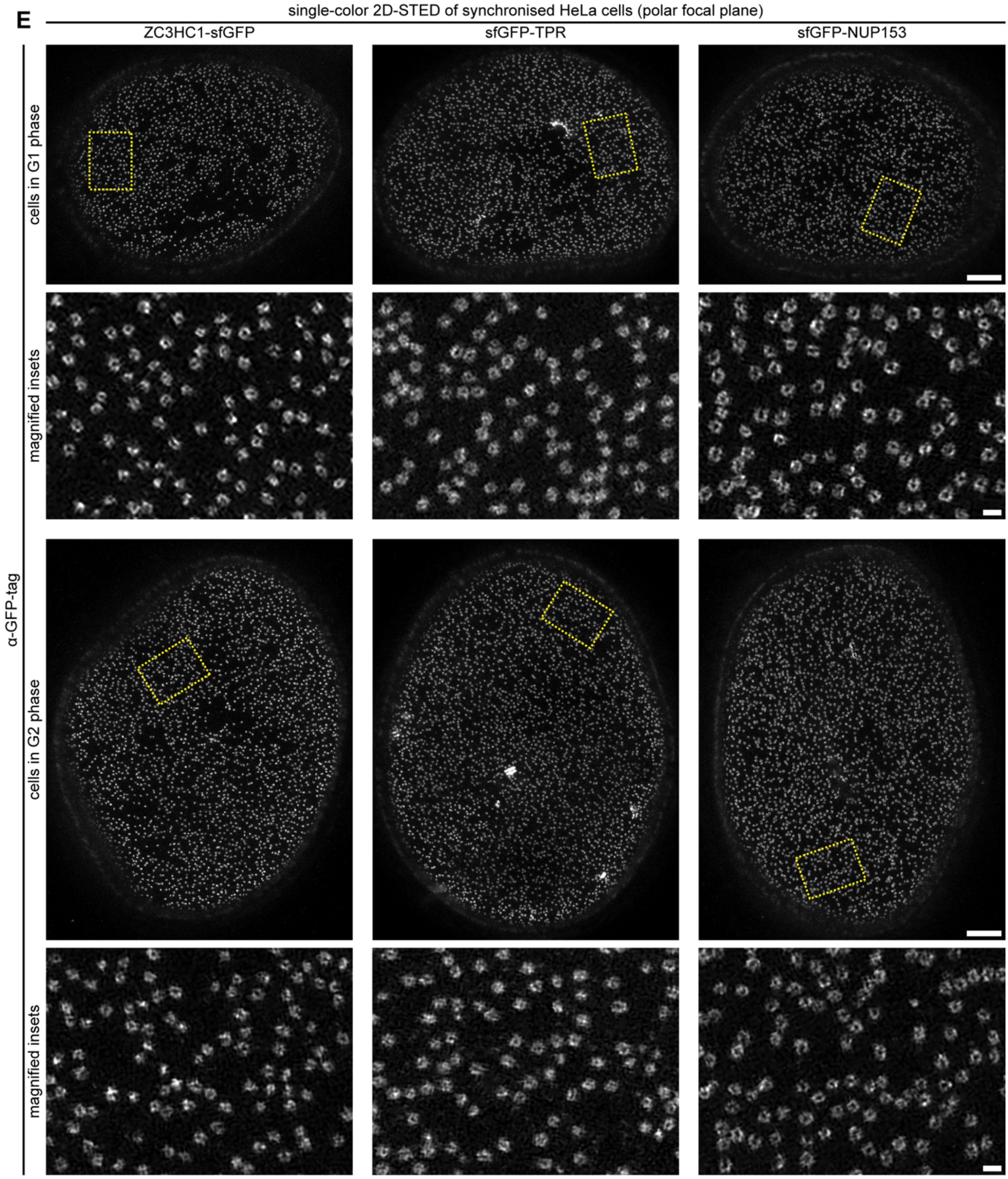


**Supplemental Figure S5. Geometrical modelling of 2D-STED signal distributions and extended STED imaging support the interpretation of how TPR and ZC3HC1 are positioned at NPC-associated NBs and NBLSs. (A)** Geometrical considerations for interpreting 2D-**STED** signal distributions at NPC-associated NBs and NBLSs. **(A1)** Schematic cross-sectional representations of NPCs embedded in an idealised planar NE without curvature. In ZC3HC1 KO cells, or in cells expressing only the N-terminally truncated TPRΔ1–58, NPCs are depicted as possessing only the prototypic NPC-anchored NB. For WT cells, two conceivable scenarios are shown: one in which all NPCs carry an identical NB:NBLS ensemble, and another in which individual NPCs differ in the number of NBLSs appended to the NB proper. The positions of the sfGFP tags at TPR's N-termini are indicated in green, and that of the N-terminal NUP153 region used as the NR reference in magenta. The dimensions of the NPCs, NBs, and NBLSs are drawn approximately to scale relative to each other, based on corresponding structural measurements obtained for *Xenopus* oocyte NBs and NBLSs (Scheer *et al.*, 1988; Ris and Malecki, 1993; Gunkel *et al.*, 2021). Also indicated to scale is a range of ~700 nm along the optical z axis, corresponding approximately to the axial resolution of conventional 2D-STED microscopy. Based on a mean NPC density of ~11.6 NPCs/$\mu m^2$ determined for HeLa P2 cells (our unpublished data), and assuming an idealised regular NPC distribution across the NE, a range of ~700 nm would encompass approximately three successive NPCs along the z axis within the planar NE depicted here. **(A2)** The same models as in A1, now illustrating how the individual fluorescence signals originating from the depicted NUP153 and sfGFP-TPR positions would combine in the resulting 2D-STED image in the absence of NE curvature. Magenta and green lines drawn parallel to the optical z axis connect corresponding NUP153 and sfGFP-TPR positions within the three NPCs depicted over the ~700-nm axial range. Because the respective positions are aligned along the z axis and occur at the same distance from the NE, their fluorescence signals would overlap at the same position along a line profile drawn perpendicular to the NE. To illustrate this signal superposition, each magenta or green position shown in the NPC:NB:NBLS models is represented below by one square of the corresponding colour, with squares representing signals at the same position stacked vertically. Thus, the six NUP153 positions depicted in each model give rise to a single stack of six magenta squares. Likewise, the six TPR T1 positions in model 1a give rise to one stack of six green squares. In the uniform WT scenario shown in 1b, the six T1 and six T2 positions produce two corresponding stacks of six green squares, whereas in the variable WT scenario shown in 1c, the different numbers of NBLSs appended to the three NPCs result in successive stacks of six, four, and two green squares for the T1, T2, and T3 layers, respectively. For simplicity, each TPR layer depicted in the structural models contains the same number of sfGFP positions; this assumption serves only to illustrate the superposition of the respective fluorescence signals and does not imply equal TPR copy numbers in the NB and individual NBLSs. The horizontal width of each square is set to 20 nm, schematically representing the width assigned to an individual fluorescence signal at the lateral resolution considered here. Based on previous TEM, SEM, iTEM, and iSEM analyses of *Xenopus* oocyte NBs and NBLSs (Scheer *et al.*, 1988; Ris and Malecki, 1993; Gunkel *et al.*, 2021), the TPR T1 signal is positioned ~50 nm from the NUP153 reference line, while the centres of successive TPR signal layers are separated by ~42.5 nm. Consequently, the modelled TPR signal extends over 20 nm when only the T1 layer is present, over 62.5 nm when T1 and T2 are present, and over 105 nm when T1, T2, and T3 are present, measured from the outer edge of the first to the outer edge of the most distal TPR signal. (**A3**) Schematic side view of part of a representative HeLa P2 cell nucleus, illustrating the NE curvature that has to be considered when NPCs and their nuclear appendices are imaged in side view by conventional 2D-STED microscopy. Systematic profilometry and analysis of confocal z-stacks from numerous HeLa P2 nuclei between mid-G1 and mid-G2 (our unpublished data) revealed roundish to ellipsoidal nuclear outlines when viewed from the pole, whereas side views – as schematically depicted here for one side of the nucleus – approximated a stadium- or capsule-shaped profile. In the idealised representation shown here, the curved NE connecting the upper and lower nuclear surfaces is approximated by a semicircle whose diameter corresponds to the mean nuclear height (*h*) of ~6 µm determined for FA-fixed HeLa P2 cells. When a 700-nm interval along the optical z axis is centred on the equatorial focal plane, the corresponding segment of a semicircular NE with a diameter of 6 µm spans an angle of ~13.4°. **(A4)** The same structural scenarios as in A1 and A2, now modelled for the curved NE geometry introduced in A3 and with the equatorial focal plane (EFP) passing through the middle of the three depicted NPCs. Model 2a represents the situation in ZC3HC1 KO cells or in cells expressing TPRΔ1–58 in the ZC3HC1 WT background, in which only the NPC-anchored NB exists. Model 2b represents a WT scenario in which all three NPCs carry identical NB:NBLS ensembles, whereas models 2c–2e represent different WT scenarios that vary in the number of NBLSs appended to the three NBs and also differ with regard to which of the three NBs are equipped with NBLSs. As in A2, magenta and green lines connect the corresponding NUP153 and sfGFP-TPR positions, and the fluorescence signals originating from each of these positions are represented below the structural models by squares of the corresponding colour. Owing to the NE's curvature, the corresponding NUP153 and TPR positions at the three NPCs no longer lie on common lines parallel to the optical z axis. Instead, their fluorescence signals become slightly displaced relative to one another in the resulting image, rather than being superimposed as in the curvature-free models in A2. The lowermost row additionally shows the resulting overall NUP153 and sfGFP-TPR signal distributions after summation of the individual signal contributions. Numbers indicate the overall width of the modelled sfGFP-TPR signal distributions. **(A5)** The corresponding five scenarios, now modelled for a non-equatorial focal plane (nEFP) passing through the middle NPC, which is tilted by 3.3° from the perpendicular orientation at the EFP (86.7° instead of 90°). The EFP itself now lies between the second and third NPC. This displacement of the focal plane further increases the positional spread of the NUP153 and sfGFP-TPR signals compared with the corresponding models in A4. Nevertheless, the resulting NUP153 and TPR signal distributions either remain separated or overlap only marginally, to an extent represented by less than the width of a single square. Such situations were classified as showing no NUP153-

TPR boundary violation, because the NUP153 and TPR signals would still appear as clearly distinguishable distributions, with only minor signal overlap in the corresponding STED images. **(A06)** Two sets of the same structural scenarios as in A4 and A5, now modelled for more strongly displaced non-equatorial focal planes. With increasing displacement from the equator, the effects of NE curvature become more pronounced, causing the individual NUP153 and sfGFP-TPR signals to become progressively more displaced relative to one another in the resulting image. In these examples, the resulting summed signal distributions overlap substantially, such that part of the NUP153 signal is positioned farther from the NPC than part of the TPR signal, while part of the TPR signal is positioned closer to the NPC than part of the NUP153 signal. We refer to this loss of the normally expected spatial ordering of the two signal distributions as an NUP153-TPR boundary violation. NE regions displaying such obvious boundary violations were excluded from the line-profile analyses shown in Figure 5B and 5D. **(A7)** Summary of the modelled NUP153 and sfGFP-TPR signal distributions for all scenarios in A2, A4, and A5 that do not exhibit an NUP153–TPR boundary violation. Only the resulting summed signal distributions are shown. **(A8)** Corresponding summary of the modelled signal distributions for the scenarios in A6 that exhibit an NUP153–TPR boundary violation. **(B-D)** Two-colour STED microscopy of HeLa P2 cells. Cells were labelled with anti-GFP-tag nanobodies and anti-NUP153 antibodies. Images of representative nuclei were acquired at the equatorial focal plane, with image acquisition settings identical for those cell lines that were directly compared. **(B)** STED images of cells expressing full-length sfGFP-TPR or sfGFP-TPRΔ1–58 in the HeLa P2 WT background, and sfGFP-TPR in the ZC3HC1 KO background. **(B1)** STED images of the nuclei shown in Figure 5A, here presenting the individual channels next to the composite images. Bar, 2 µm. **(B2)** Higher-magnification views of representative NE regions from five different nuclei per cell line. The nuclear (N) and cytoplasmic (C) sides of the NE are indicated. Bar, 200 nm. **(C)** STED images of representative nuclei of HeLa P2 cells expressing full-length ZC3HC1-sfGFP or ZC3HC1Δ487–502-sfGFP. Two of the five nuclei shown here are identical to those shown in Figure 5C. Bars, 2 µm. **(D)** STED images of cells expressing full-length TPR-sfGFP in a HeLa P2 WT and ZC3HC1 KO background. **(D1)** STED images entire nuclei. Bar, 2 µm. **(D2)** Higher-magnification STED images of representative NE regions. Bar, 200 nm. **(E)** Single-colour STED microscopy of cell-cycle-synchronised HeLa P2 cells expressing ZC3HC1-sfGFP, sfGFP-TPR, or sfGFP-NUP153, labelled only with anti-GFP-tag nanobodies. Images were acquired at the nuclear pole. For each cell line, one representative nucleus in G1 phase and one in G2 phase are shown. Rectangles mark selected regions of the nuclear surface that are shown at higher magnification in the corresponding lower rows, revealing ring-like signal distributions for sfGFP-NUP153, sfGFP-TPR, and ZC3HC1-sfGFP. Bars, 2 µm (overview images) and 200 nm (magnified insets).

Supplemental Figure S6 (1/3)

A

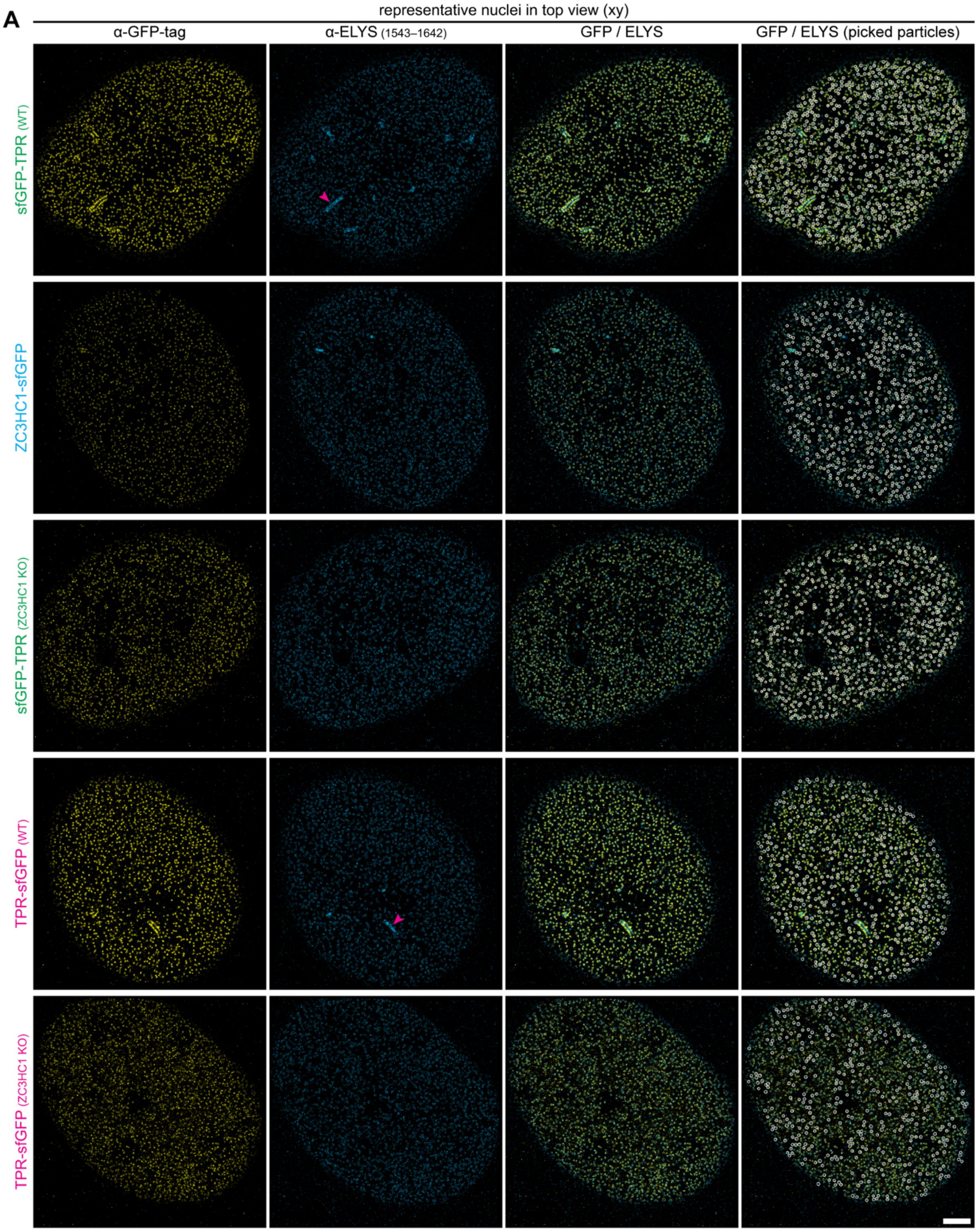

Supplemental Figure S6 (2/3)

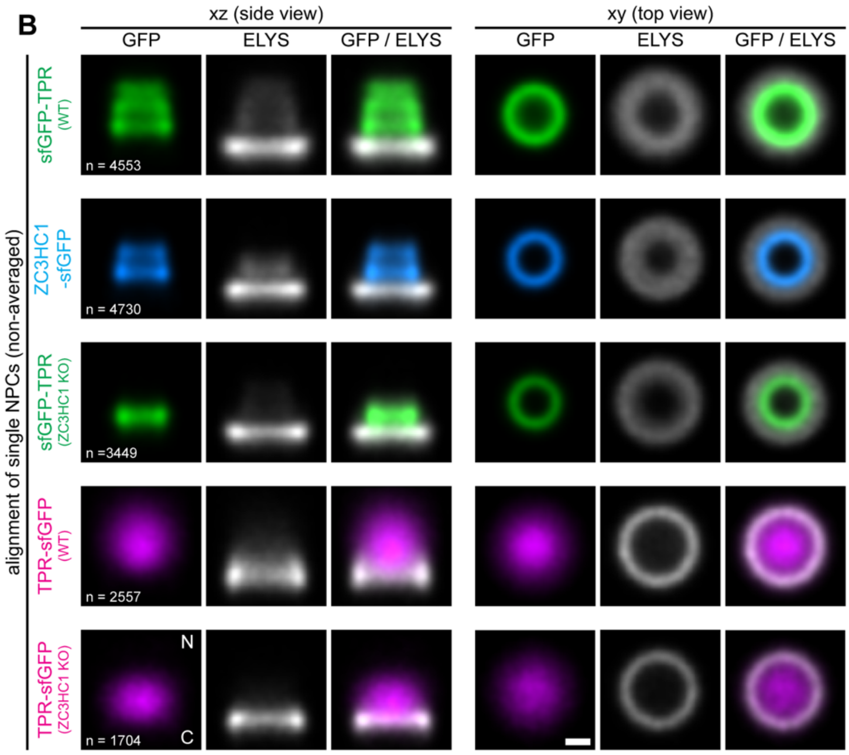


**Supplemental Figure S6. 4Pi-STORM analysis of automatically picked particles distinguishes the stratified NB:NBLS architecture formed by ZC3HC1 and TPR's scaffold-forming domain from the disordered distribution of TPR's CTD. (A)** Overview 4Pi-STORM images of polar regions of representative nuclei from HeLa P2 cells expressing sfGFP-TPR, ZC3HC1-sfGFP, or TPR-sfGFP in the WT background, and sfGFP-TPR or TPR-sfGFP in ZC3HC1 KO cells. The immunolabelled sfGFP-tags are shown in yellow and the immunolabelled ELYS in blue. Particles identified by the automated particle-picking procedure are encircled in the corresponding images on the right. Magenta arrowheads mark local invaginations of the NE sporadically observed in some HeLa P2 nuclei. Bar, 2 µm. **(B)** Composite views of COM-aligned stacks of NPC:NB- and NPC:NB:NBLS-containing particles. The same automatically picked particles and computationally aligned particle stacks as shown in Figure 6B are presented here, but with the ELYS channel additionally displayed separately, revealing some crosstalk from ELYS into GFP, while crosstalk from GFP into ELYS is virtually imperceptible across the different sfGFP-tagged cell lines. Bars, 50 nm.

#### Supplemental Figure S7 (1/5)

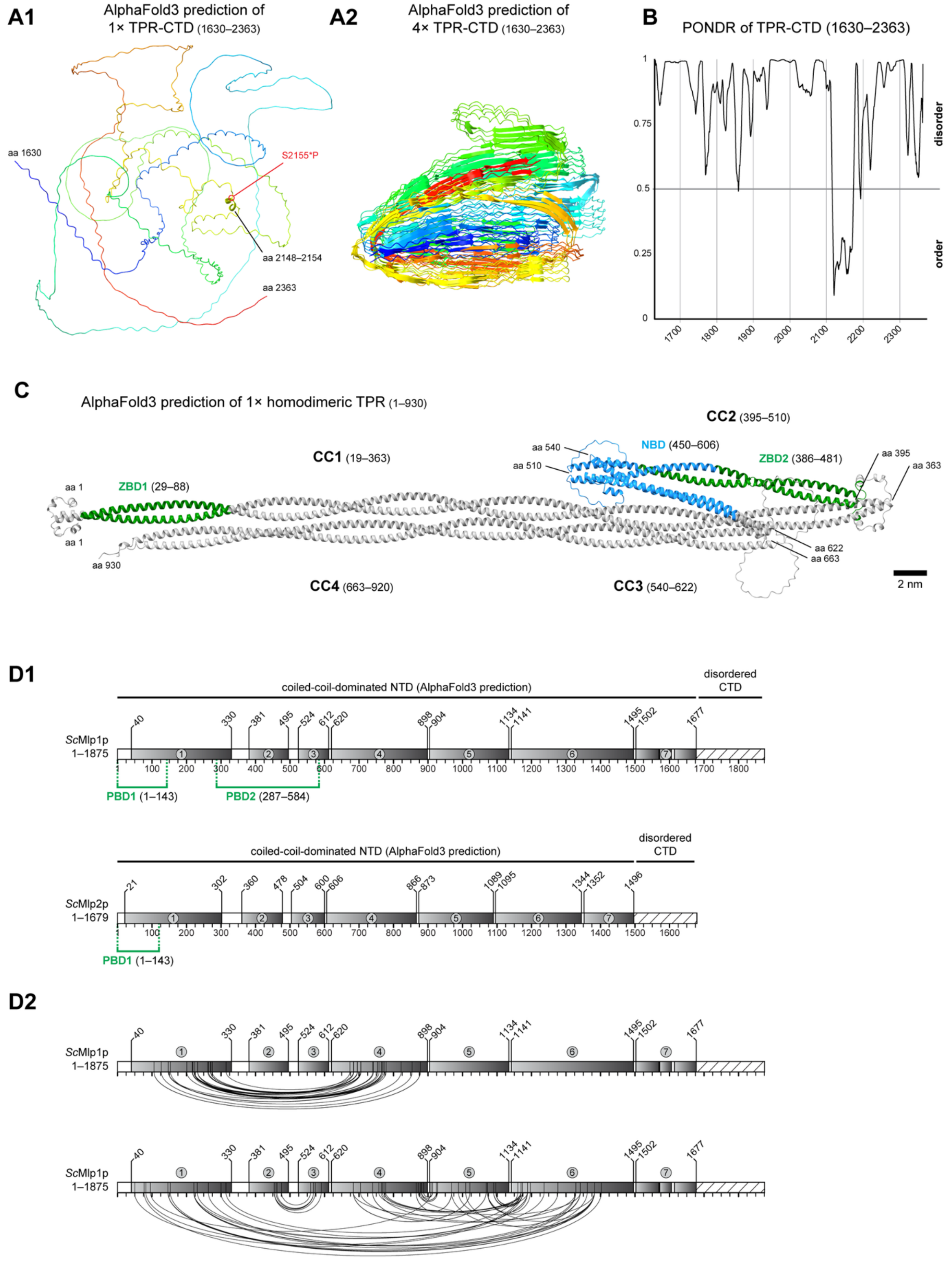

Supplemental Figure S7 (2/5)

**D3**

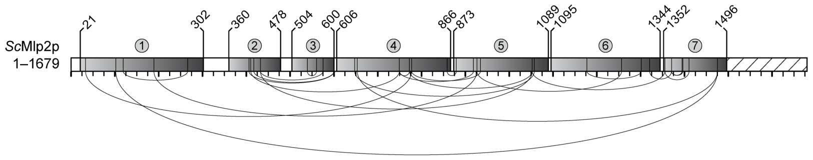


**D4**

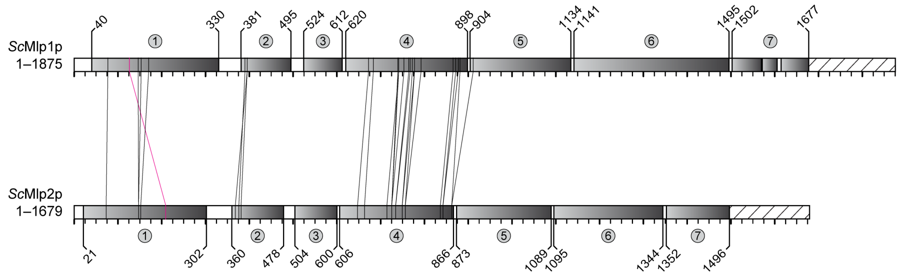


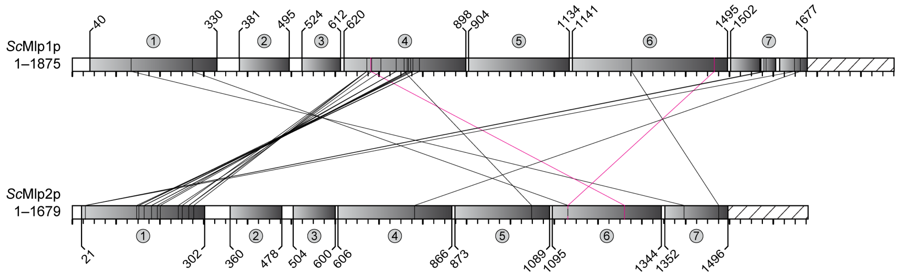


**E1**

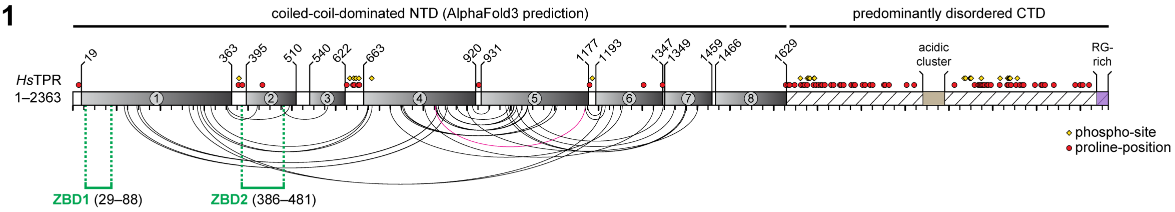


**E2**

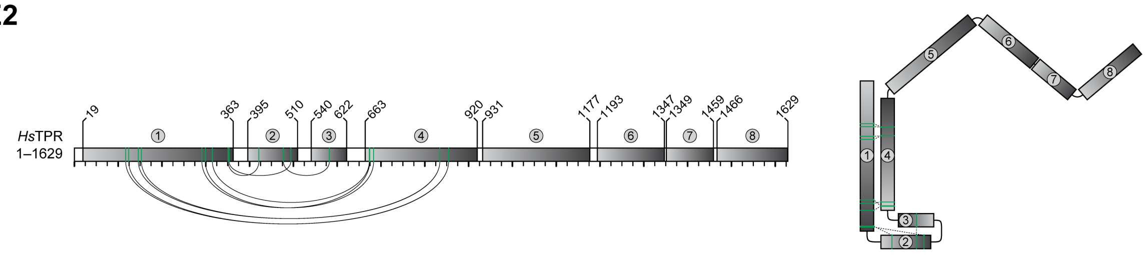

Supplemental Figure S7 (3/5)

**E3**

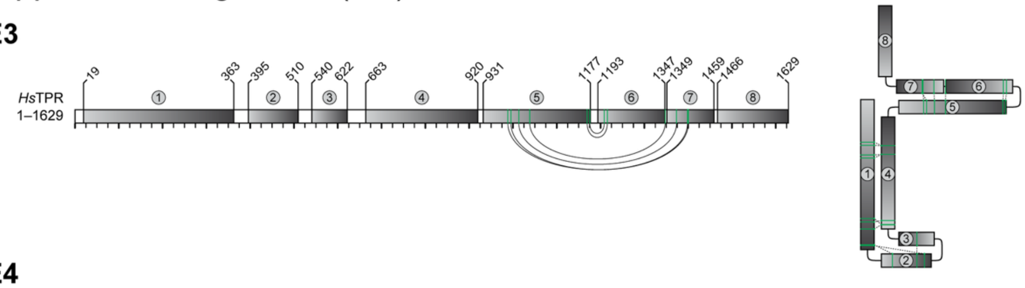


**E4**

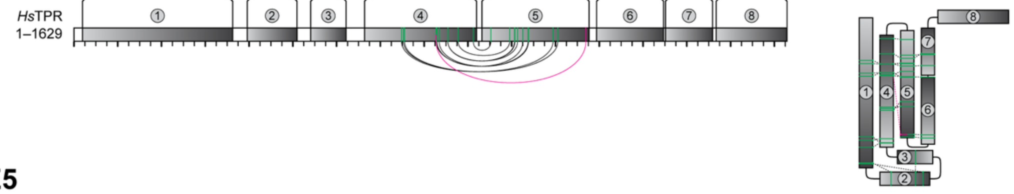


**E5**

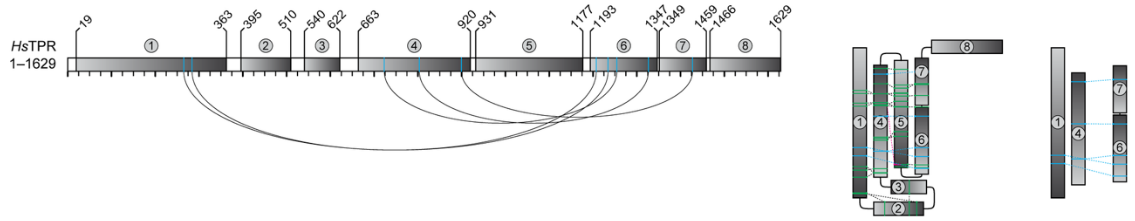


**E6**

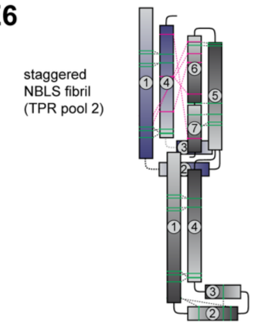


**E7**

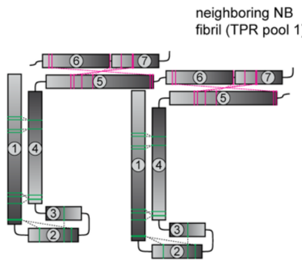


**E8**

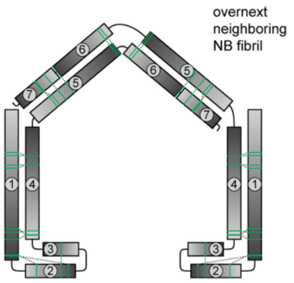


**E9**

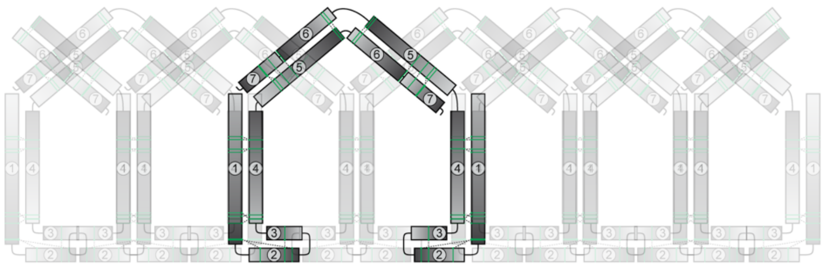

**E10**

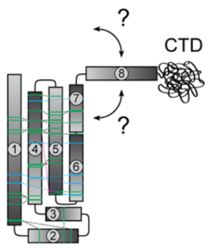

#### Supplemental Figure S7 (4/5)

**F**

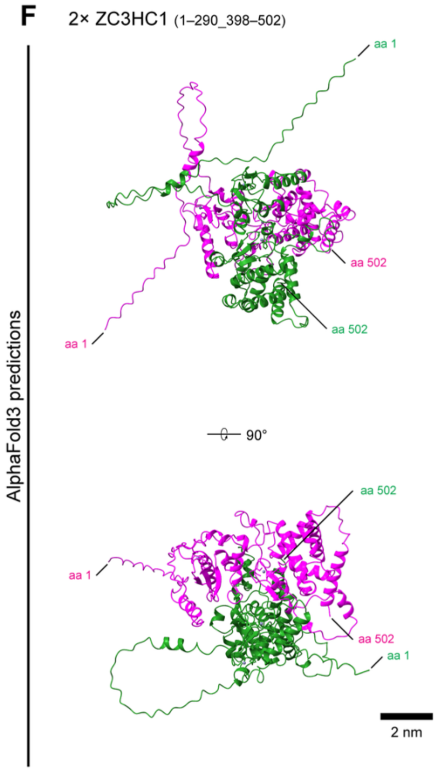


**G**

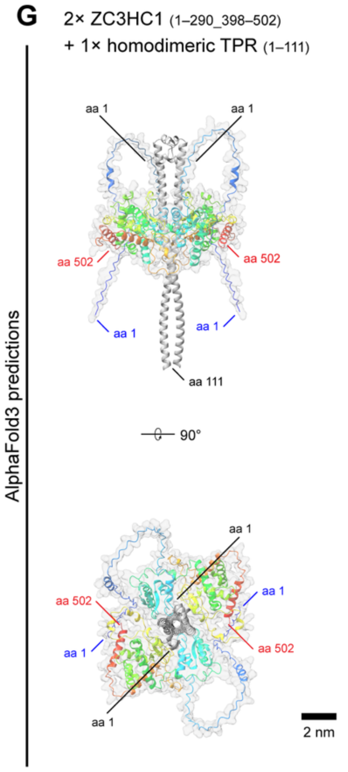


**H**

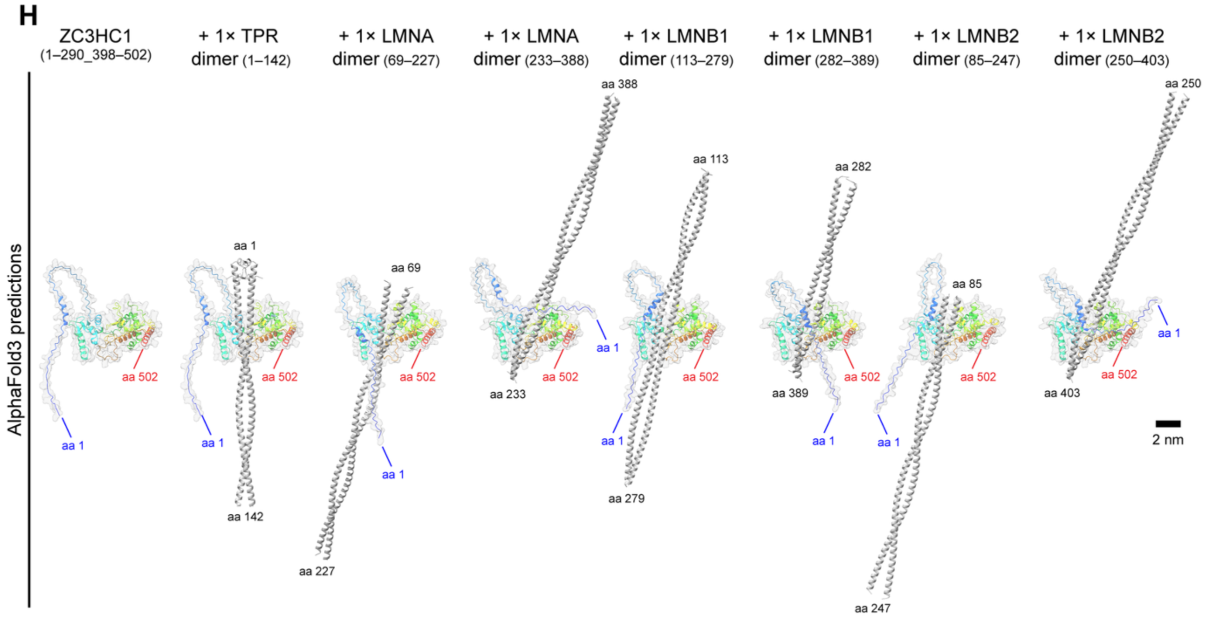

Supplemental Figure S7 (5/5)

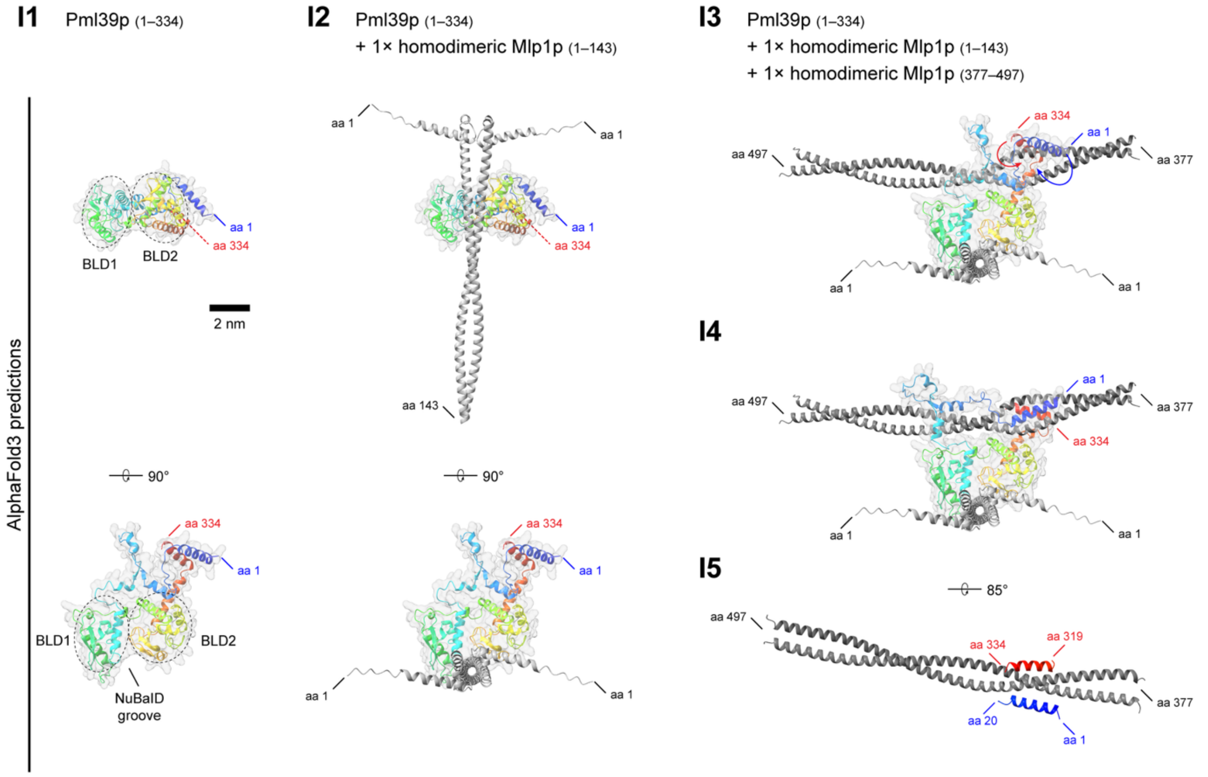


**Supplemental Figure S7. Extended analyses of TPR/Mlp structural organisation and predicted modes of interaction with ZC3HC1/Pml39p. (A)** AlphaFold3 predictions of TPR's CTD. **(A1)** Prediction for a single TPR CTD (aa 1630–2363), illustrating its predominantly disordered conformation and a short α-helical segment comprising aa 2148–2154. The adjacent phosphosite S2155 is indicated. **(A2)** Prediction generated with four TPR CTDs modelled together, in which substantial portions of the individual CTDs, predicted to be largely disordered when modelled in isolation, are arranged in β-sheet structures. **(B)** PONDR profile of TPR's CTD (aa 1630–2363). Values above 0.5 indicate predicted disorder, whereas values below 0.5 indicate predicted order. A prominent ordered segment between aa 2116 and 2173 encompasses the short α-helical region at aa 2148–2154 also predicted by AlphaFold3. **(C)** AlphaFold3 prediction of a homodimeric TPR fragment comprising aa 1–930, with CC1–4 and the positions of ZBD1, ZBD2, and NBD indicated. The prediction illustrates folding of CC4 onto CC1 and the resulting antiparallel arrangement of these two CC segments. CC2 and

CC3 are shown here in their original AlphaFold3-predicted positions. This arrangement differs from the model in Figure 7B, where these two CCs were repositioned via the flexible spacer regions between the CC segments to illustrate an alternative geometry that may better accommodate exposure of the NBD for NR binding. Bar, 2 nm. **(D)** Published XL-MS data for *S. cerevisiae* Mlp1p and Mlp2p mapped onto their predicted CC organisation. **(D1)** Schematics of Mlp1p and Mlp2p showing AlphaFold3-predicted CC segments and the positions of PBD1 and PBD2. **(D2)** Intramolecular cross-links reported for Mlp1p. Lines connect the amino acid residues identified as cross-linked in the respective XL-MS datasets. For clarity, cross-links between CTD residues are not shown. **(D3)** Corresponding intramolecular cross-links reported for Mlp2p. **(D4)** Intermolecular cross-links detected between Mlp1p and Mlp2p. Grey lines indicate cross-links that can be explained by either a parallel, non-staggered alignment of unfolded Mlp1p and Mlp2p homodimers, as shown above, or by an alignment of multiply folded Mlp1p and Mlp2p homodimers, as shown below. Magenta lines indicate cross-links that cannot be explained by either arrangement. **(E)** Published XL-MS data for human TPR and the extent to which proposed successive folding steps of its coiled-coil-dominated NTD are compatible with these data. **(E1)** Intramolecular TPR cross-links compiled from available XL-MS datasets and mapped onto the linear TPR scheme shown in Figure 7A. Lines connect the amino acid residues identified as cross-linked in the respective datasets. The cross-links shown here are subsequently considered in different subsets in E2–10, either in the context of successive folding steps of a single TPR homodimer or of possible arrangements involving two or more TPR homodimers. The folding arrangements shown in E2–5 and E10 are compatible with the cross-links indicated by grey lines. The single cross-link shown in magenta cannot be plausibly explained by any of the folding arrangements considered here. For clarity, cross-links between residues within the CTD are not shown. **(E2)** Cross-links between CC1 and CC4 and between CC2 and CC3, consistent with folding of CC4 onto CC1 and of CC3 onto CC2, resulting in antiparallel alignment of the respective CC pairs. The corresponding folded arrangement is shown schematically on the right. **(E3)** Cross-links between CC5 and CC6/7, consistent with folding of CC6/7 onto CC5. **(E4)** Cross-links between CC4 and CC5, consistent with folding of the CC5–7 segment onto CC1/4. The single cross-link incompatible with this arrangement is highlighted in magenta. **(E5)** Cross-links between CC1 and CC6 and between CC4 and CC6/7, consistent with the final folding step shown here, in which the CC5–7 segment folds onto the CC1/4 bundle. These cross-links are additionally shown in the simplified representation on the far right, in which CC2, CC3, CC5, and CC8 are omitted for clarity. **(E6–E9)** Hypothetical arrangements involving two or more TPR homodimers, assessed for compatibility with the available XL-MS data. For clarity, CC8 is omitted in these Figures. **(E6)** Hypothetical staggered alignment of TPR homodimers belonging to the NPC-anchored and ZC3HC1-appended TPR subpopulations T1 and T2. Numerous reported cross-links connect residues in CC1/4 with residues in CC6/7 and are incompatible with the arrangement shown here, in which CC1/4 of the TPR T2 homodimer would align with CC6/7 of the TPR T1 homodimer such that CC4 of T2 runs antiparallel to CC6/7 of T1. By contrast, within an individual folded TPR homodimer, CC4 and CC6/7 would be aligned in parallel, as shown in Figure 7C2 and 7C3 and Supplemental Figure S7E3–S7E5. For clarity, only CC1–4 are shown for the TPR T2 homodimer at the upper left of the Figure. **(E7)** Hypothetical staggered alignment of branched-off CC5–7 segments of neighbouring NB fibrils of the TPR T1 subpopulation. In this arrangement, CC5 of one NB-fibril-forming TPR homodimer would align in parallel with CC6/7 of a TPR homodimer belonging to a neighbouring NB fibril. Such an arrangement would be incompatible with the numerous reported cross-links between CC5 and CC6/7. Within an individual folded TPR homodimer, by contrast, CC5 and CC6/7 would be aligned antiparallel, as shown in Figure 7C2 and 7C3 and Supplemental Figure S7E3–S7E5. For clarity, only parts of the three TPR homodimers are shown: CC6/7 for the homodimer at the upper left, CC1–7 for the homodimer on the left, and CC1–5 for the homodimer on the right. **(E8)** Hypothetical antiparallel alignment of the branched-off CC5–7 segments of next-but-one neighbouring fibrils. Such an arrangement would be compatible with the reported CC5:CC6/7 cross-links, provided that no further intramolecular folding involving CC5–7 occurred beyond the folding of CC1:CC4. The reason for considering next-but-one rather than directly neighbouring fibrils is twofold: first, zig-zag-like arrangements between the distal parts of next-but-one NB fibrils have been described in *Xenopus* oocytes (Gunkel *et al.*, 2021), and second, the lengths of the entire CC5–7 segments would allow the shallow angle of 110° between CC5 and CC6, as shown here, only for such an arrangement. The directly neighbouring fibril, which would be positioned between the two TPR homodimers depicted here, is not shown. **(E9)** The arrangement shown in E8, now integrated into an array of NB fibrils. Directly neighbouring fibrils, omitted in E8, are indicated by shaded TPR homodimers. Repetition of such antiparallel interactions between the CC5–7 segments of next-but-one neighbouring fibrils in a criss-crossed, interleaved manner could, in principle, generate a TR of zig-zag-like appearance (Gunkel *et al.*, 2021). **(E10)** Model in which CC8 does not participate in the formation of a folded NB fibril or TR but instead remains flexible and projects the CTD away from the NB-fibril-forming NTD bundle. CC1–7 are arranged as in the folding models shown above, whereas CC8 is depicted as potentially acting as a jib that could position the CTD further towards the longitudinal mid-axis of the NB/NBLS. While no cross-links from CC1–7 with CC8 have so far been reported, the proposed positioning of CC8 nonetheless remains speculative, as indicated by the question marks next to arrows illustrating possible movements of CC8. The CTD is shown schematically. **(F)** AlphaFold3 prediction generated with two ZC3HC1 polypeptides, illustrating a potential ZC3HC1 homodimer. Two orientations are shown, with one of the ZC3HC1 polypeptides coloured in green and the other in magenta. For clarity, the largely unstructured loop within BLD2, most of which lies between aa 290 and

398 and which is dispensable for NB association of ZC3HC1 and subsequent recruitment of TPR T2 polypeptides (Gunkel *et al.*, 2023), is omitted. Bar, 2 nm. **(G)** AlphaFold3 prediction generated with two ZC3HC1 polypeptides and one homodimeric TPR fragment comprising aa 1–111. The predicted ZC3HC1 dimer embraces the ZBD1-containing TPR coiled coil, with the NuBaID grooves of both ZC3HC1 polypeptides contributing to the interaction. As in F, the largely unstructured BLD2 loop of ZC3HC1 is omitted for clarity. Two orientations are shown. Bar, 2 nm. **(H)** AlphaFold3 predictions illustrating promiscuity of the NuBaID groove/TBE1 in silico. ZC3HC1 was modelled either alone or together with a homodimeric TPR fragment comprising aa 1–142 or with different CC-forming fragments of LMNA, LMNB1, or LMNB2**.** In these predictions, AlphaFold3 places several unrelated CCs into the TBE1 groove in a manner similar to TPR's ZBD1, although there is no evidence for such promiscuity in vivo. As in F and G, the largely unstructured BLD2 loop is omitted for clarity. Bar, 2 nm. **(I)** AlphaFold3 predictions of the interaction between *S. cerevisiae* Pml39p and Mlp1p. **(I1)** Prediction of Pml39p, illustrating its two BLD modules and the NuBaID groove formed between them, corresponding to the Mlp-binding element MBE1. Two orientations are shown. Bar, 2 nm. **(I2)** Prediction of Pml39p together with a homodimeric Mlp1p fragment comprising aa 1–143, illustrating accommodation of the PBD1-containing CC within the MBE1 groove. Two orientations are shown. **(I3)** Representation containing one Pml39p polypeptide, a homodimeric Mlp1p fragment comprising aa 1–143 and containing PBD1, and a second homodimeric Mlp1p fragment comprising aa 377–497 and containing PBD2. Pml39p is shown in the same conformation as in the lower representation in I2, with its near-N-terminal and near-C-terminal α-helices not yet contacting PBD2. Blue and red arrows illustrate how these two α-helices could approach the PBD2-containing CC from opposite sides and clasp it. **(I4)** Corresponding AlphaFold3 prediction with the same components and viewed from the same direction as in I3, now showing the near-N-terminal and near-C-terminal α-helices of Pml39p

positioned on opposite sides of the PBD2-containing CC and clasping it. **(I5)** For clarity, only the PBD2-containing Mlp1p CC and the two NuBaID-flanking α-helices of Pml39p are shown, with the near-N-terminal α-helix in blue and the near-C-terminal α-helix in red.

Supplemental Figure S8 (1/4)

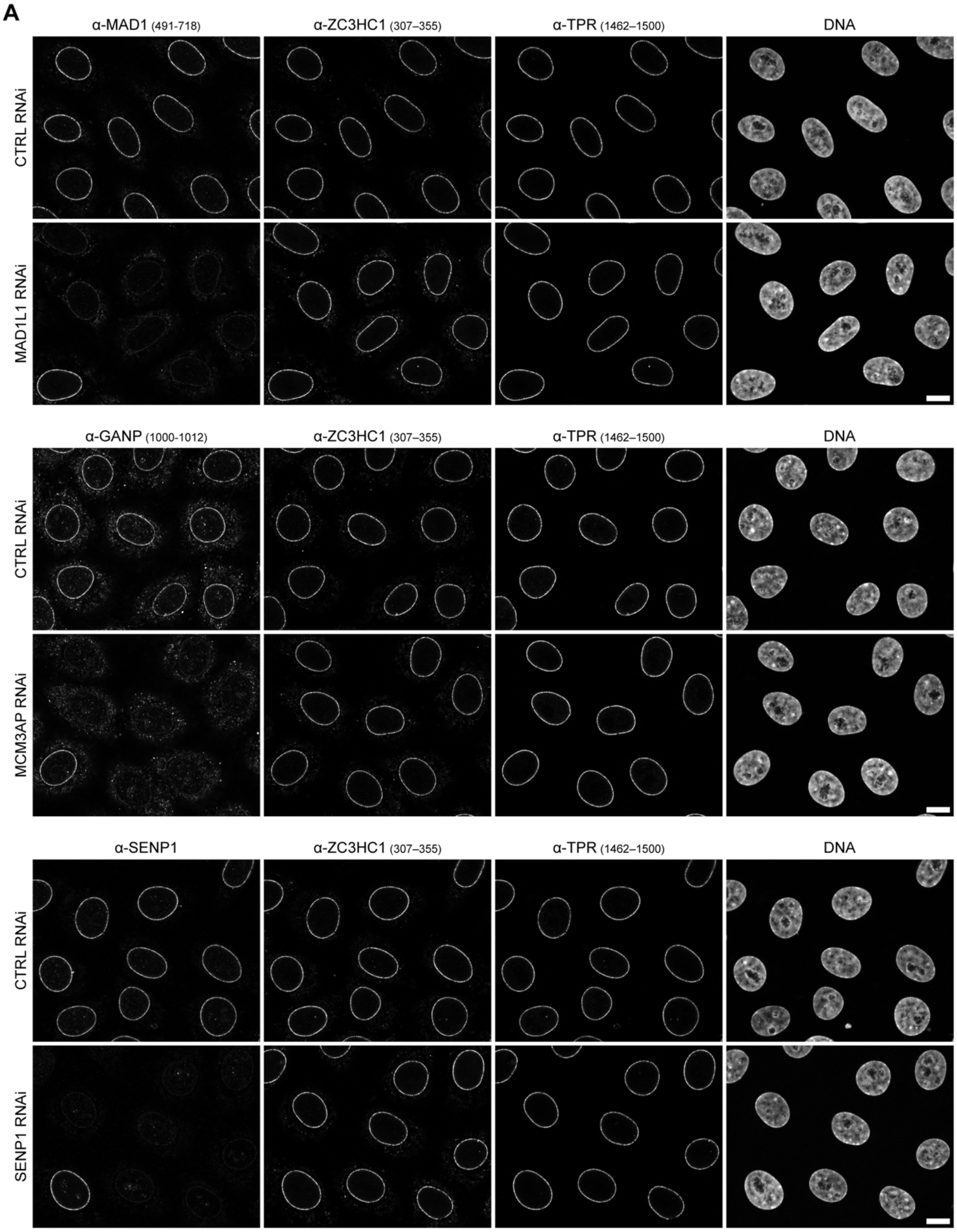

#### Supplemental Figure S8 (2/4)

**B1**

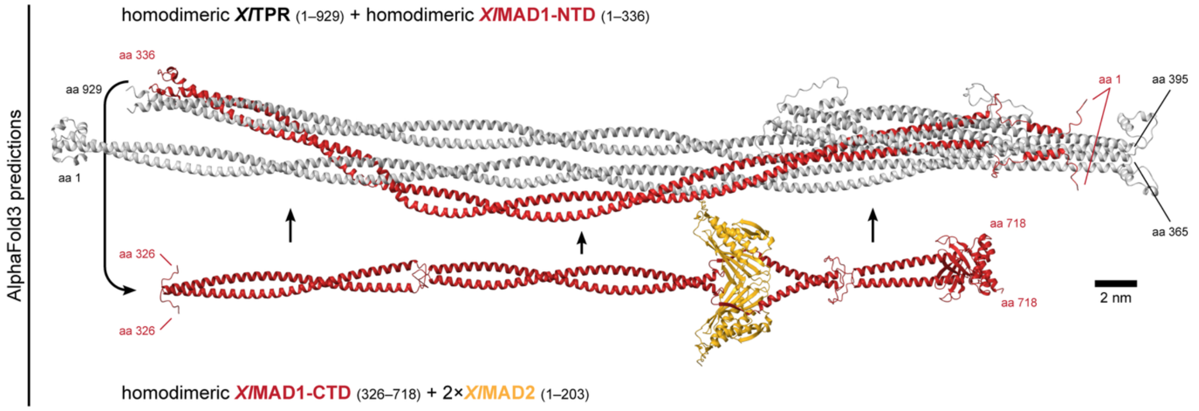


**B2**

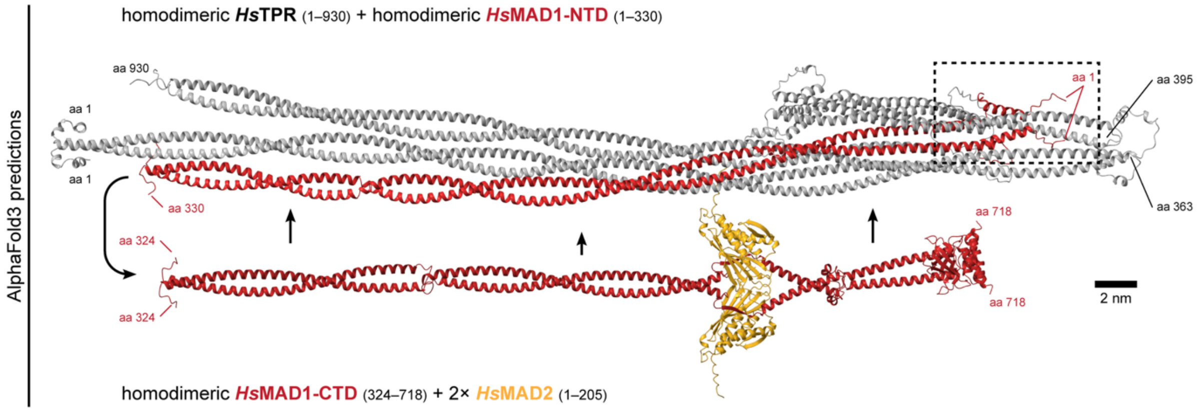


**B3**

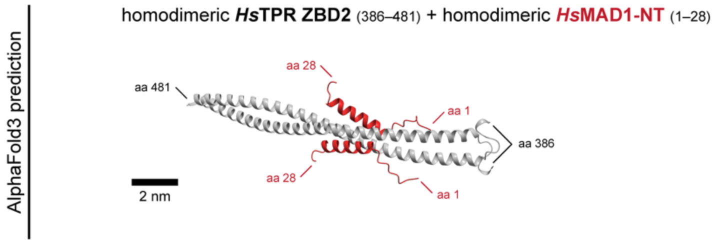

### Supplemental Figure S8 (3/4)

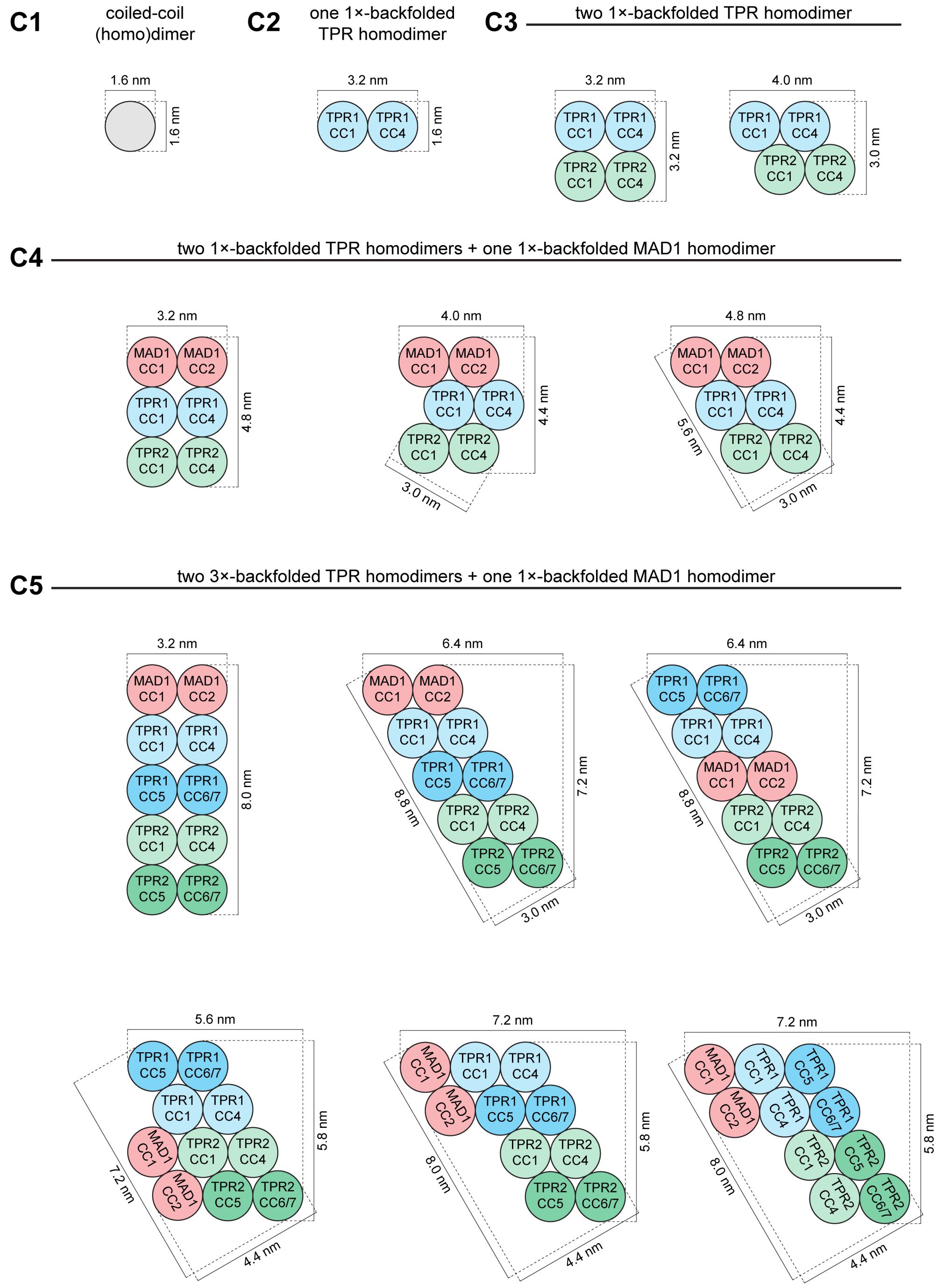

Supplemental Figure S8 (4/4)

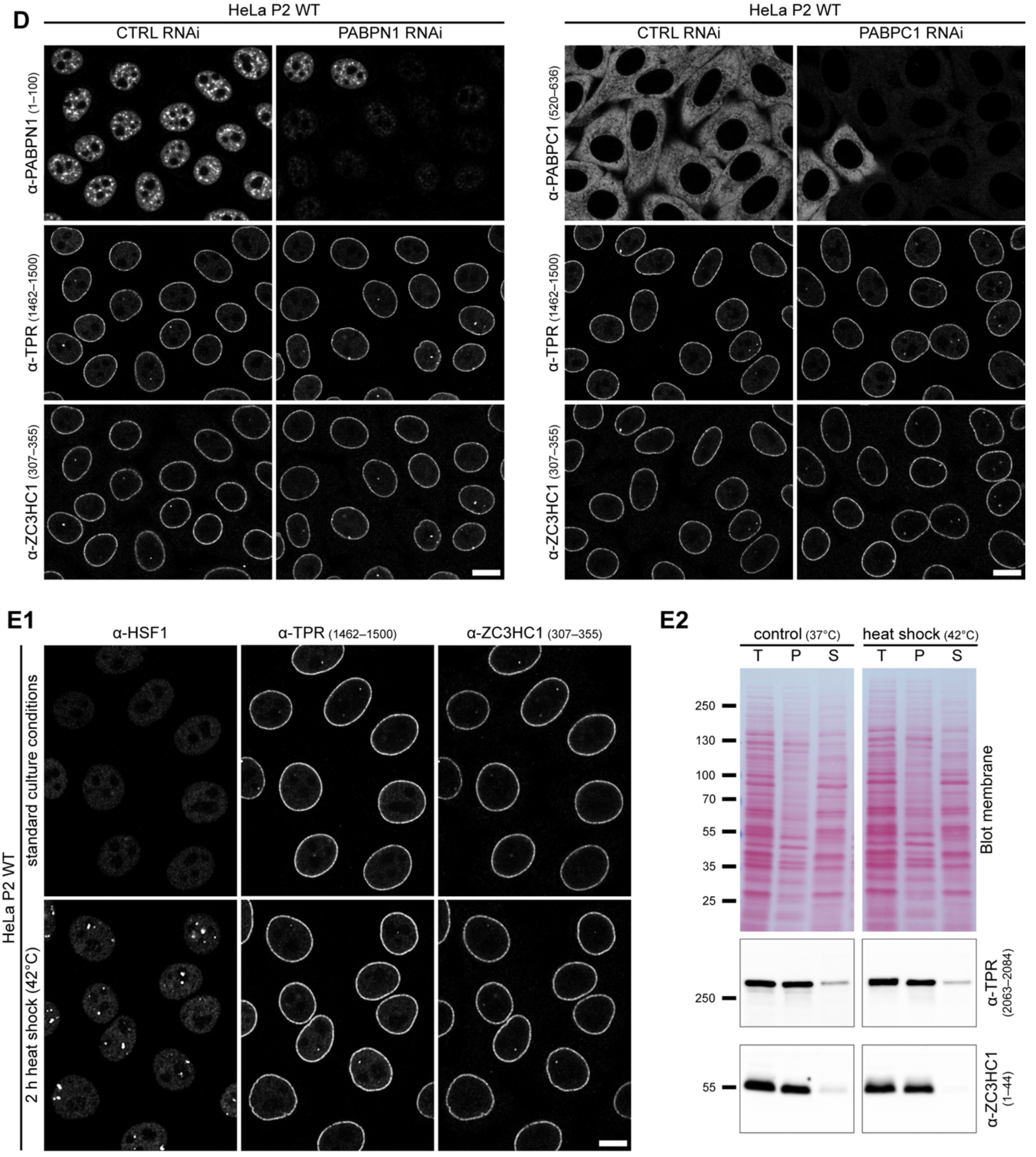


**Supplemental Figure S8. Predicted TPR:MAD1 arrangements and preserved NPC association of TPR and ZC3HC1 upon depletion of other NB-associated proteins or poly(A)-binding proteins and after heat shock**. **(A)** Representative IFM of HeLa P2 cells treated with control (CTRL) siRNAs or siRNAs targeting MAD1L1, MCM3AP/GANP, or SENP1 to assess whether depletion of these NB-associated proteins affects NE association of TPR and ZC3HC1. Cells were triple-labelled with antibodies against the respective RNAi target, ZC3HC1, and TPR. While the respective target signals are strongly reduced upon RNAi, NE association of TPR and ZC3HC1 remains largely unaffected. Some cells that had remained non-transfected within the respective siRNA-treated populations are shown as reference. Bar, 10 µm. **(B)** AlphaFold3 predictions of interactions between TPR and MAD1. **(B1)** Prediction of the homodimeric *Xenopus laevis* TPR segment aa 1–929, comprising CC1–4, together with the first of the two homodimeric CC segments of *Xl*MAD1 (MAD1-NTD, aa 1–336). The second homodimeric MAD1 CC segment (MAD1-CTD, aa 326–718), which binds two MAD2 polypeptides, was modelled separately together with two copies of *Xl*MAD2 (aa 1–203). The known folding of the MAD1-CTD onto the MAD1-NTD, resulting in antiparallel alignment of the two MAD1 CCs, is indicated by arrows. The two predictions are shown together to illustrate how the folded MAD1 homodimer could extend along the CC1:CC4 region of TPR. Bar, 2 nm. **(B2)** Corresponding AlphaFold3 predictions for the human proteins, with the TPR:MAD1 arrangement predicted for homodimeric TPR (aa 1–930) together with the homodimeric NTD of *Hs*MAD1 (aa 1–330), and the homodimeric MAD1-CTD (aa 324–718) modelled separately together with two *Hs*MAD2 polypeptides (aa 1–205). **(B3)** AlphaFold3 prediction of a homodimeric human TPR fragment comprising ZBD2 (aa 386–481) together with the two N-terminal α-helices of the *Hs*MAD1 homodimer, located between aa 1–28. The prediction places the two MAD1 α-helices on opposite sides of the ZBD2-containing region of TPR CC2, thereby potentially shielding ZBD2 and contributing to preventing ZC3HC1 binding to TPR homodimers of the T1 subpopulation. **(C)** Schematic cross-sectional models of conceivable TPR:MAD1 arrangements within NB fibrils and their estimated dimensions. **(C1–C4)** Geometrical considerations for arrangements in which the TPR homodimers are folded only once, with CC4 aligned alongside CC1. **(C1)** Cross-section of a prototypic homodimeric CC, for which a diameter of 1.6 nm was assumed. **(C2)** Corresponding cross-section of a once-folded TPR homodimer, in which CC1 and CC4 form two aligned homodimeric CCs, resulting in estimated cross-sectional dimensions of 3.2 x 1.6 nm. **(C3)** Two principally conceivable arrangements of two such once-folded TPR homodimers. **(C4)** Conceivable arrangements formed by two once-folded TPR homodimers together with one once-folded MAD1 homodimer. MAD1 is shown either positioned between the two TPR bundles or alongside an arrangement in which the two TPR bundles contact each other directly. The approximate cross-sectional dimensions of the different arrangements are indicated. **(C5)** Conceivable cross-sectional arrangements comprising two severalfold-folded TPR homodimers, in which CC5 and CC6/7 are additionally aligned with the CC1:CC4 region, together with one once-folded MAD1 homodimer. Different possible packings are shown together with their approximate cross-sectional dimensions. (**D)** Representative IFM of HeLa P2 cells treated with control siRNAs (CTRL) or siRNAs targeting PABPN1 or PABPC1 to assess whether depletion of either protein affects NE association of TPR and ZC3HC1. Some cells that had remained non-transfected within the respective siRNA-treated populations are shown as reference. Cells were triple-labelled with antibodies against the respective RNAi target, ZC3HC1 and TPR. While the respective target signals are strongly reduced upon RNAi, NE association of TPR and ZC3HC1 remains largely unaffected. Bars, 10 µm. **(E)** Effects of heat shock on NE association of TPR and ZC3HC1 in HeLa P2 cells. **(E1)** Representative IFM of HeLa P2 cells maintained under standard culture conditions or exposed to heat shock at 42°C for 2 h. Cells were triple-labelled with antibodies against HSF1, TPR and ZC3HC1. While heat shock induced the characteristic redistribution of HSF1, NE association of TPR and ZC3HC1 remained largely unaffected. Bar, 10 µm. **(E2)** IB of cell fractions obtained from parallel cultures of HeLa P2 cells subjected to the same standard and heat-shock conditions as in E1. Cells were extracted with TX-100 in NB-s buffer and subsequently centrifuged at 20,000 × g, resulting in a supernatant (S) and an LNN-enriched pellet (P) fraction. Total cell proteins (T) were loaded for comparison. The amounts loaded in each lane corresponded to approximately equal numbers of cells. Immunolabellings for TPR and ZC3HC1 were performed on the Ponceau S-stained membranes shown above. Under both conditions, TPR and ZC3HC1 remained largely associated with the pellet fraction, with no conspicuous amounts of either protein having been released into the soluble pool upon heat shock.

### Supplemental Tables

#### Supplemental Table S1: Yeast strains.

| Description | Genotype | Source |
|---|---|---|
| AH109 | MATa, trp1-901, leu2-3, 112, ura3-52, his3-200, gal4Δ, gal80Δ, LYS2::GAL1UAS-GAL1$_{TATA}$-HIS3, GAL2$_{UAS}$-GAL2$_{TATA}$-ADE2, URA3::MEL1$_{UAS}$-MEL1$_{TATA}$-lacZ | Clontech |
| CG-1945 | MATa, ura3-52, his3-200, ade2-101, lys2-801, trp1-901, leu2-3,112, gal4-542, gal80-538, cyhr2, LYS2::GAL1$_{UAS}$-GAL1$_{TATA}$-HIS3, URA3::GAL4$_{17\text{-mers}(x3)}$-CYC1$_{TATA}$-lacZ | Clontech |
| Y187 | MATα, ura3-52, his3-200, ade2-101, trp1-901, leu2-3,112, gal4Δ, met–, gal80Δ, URA3::GAL1$_{UAS}$-GAL1$_{TATA}$-lacZ | Clontech |

#### Supplemental Table S2: Expression vectors.

| Description | Backbone (resistance) | Source |
|---|---|---|
| GAL4-BD (empty bait vector) | pGBT9 (Amp) | Clontech, Mountain View, CA, USA |
| GAL4-BD-*Hs*ZC3HC1(1–502) | pGBT9 (Amp) | (Gunkel *et al.*, 2023) |
| GAL4-BD-*Hs*ZC3HC1(72–502) | pGBT9 (Amp) | This study |
| GAL4-BD-*Hs*ZC3HC1(1–490) | pGBT9 (Amp) | This study |
| GAL4-BD-*Hs*ZC3HC1(1–467) | pGBT9 (Amp) | This study |
| GAL4-BD-*Hs*ZC3HC1(1–462) | pGBT9 (Amp) | This study |
| GAL4-BD-*Hs*ZC3HC1(1–290_398–502) | pGBT9 (Amp) | This study |
| GAL4-BD-*Hs*ZC3HC1(72–290_398–467) | pGBT9 (Amp) | This study |
| GAL4-AD (empty prey vector) | pGAD424 (Amp) | Clontech, Mountain View, CA, USA |
| GAL4-AD-*Hs*TPR-L(1–76) | pGAD424 (Amp) | This study |
| GAL4-AD-*Hs*TPR(1–60) | pGAD424 (Amp) | This study |
| GAL4-AD-*Hs*TPR(1–74) | pGAD424 (Amp) | This study |
| GAL4-AD-*Hs*TPR(1–88) | pGAD424 (Amp) | This study |
| GAL4-AD-*Hs*TPR(1–102) | pGAD424 (Amp) | (Gunkel *et al.*, 2023) |
| GAL4-AD-*Hs*TPR(1–111) = TPR-L(77–187) | pGAD424 (Amp) | This study |
| GAL4-AD-*Hs*TPR(1–175) | pGAD424 (Amp) | This study |
| GAL4-AD-*Hs*TPR(11–109) | pGAD424 (Amp) | This study |
| GAL4-AD-*Hs*TPR(20–111) | pGAD424 (Amp) | This study |
| GAL4-AD-*Hs*TPR(29–175) | pGAD424 (Amp) | This study |
| GAL4-AD-*Hs*TPR(43–175) | pGAD424 (Amp) | This study |
| GAL4-AD-*Hs*TPR(54–175) | pGAD424 (Amp) | This study |
| GAL4-AD-*Hs*TPR(110–342) | pGAD424 (Amp) | This study |
| GAL4-AD-*Hs*TPR(110–377) | pGAD424 (Amp) | This study |
| GAL4-AD-*Hs*TPR(172–651) | pGAD424 (Amp) | This study |
| GAL4-AD-*Hs*TPR(233–342) | pGAD424 (Amp) | This study |
| GAL4-AD-*Hs*TPR(233–499) | pGAD424 (Amp) | This study |
| GAL4-AD-*Hs*TPR(275–450) | pGAD424 (Amp) | This study |
| GAL4-AD-*Hs*TPR(275–481) | pGAD424 (Amp) | This study |
| GAL4-AD-*Hs*TPR(275–539) | pGAD424 (Amp) | This study |
| GAL4-AD-*Hs*TPR(347–499) | pGAD424 (Amp) | This study |
| GAL4-AD-*Hs*TPR(347–543) | pGAD424 (Amp) | This study |
| GAL4-AD-*Hs*TPR(361–539) | pGAD424 (Amp) | This study |
| GAL4-AD-*Hs*TPR(386–481) | pGAD424 (Amp) | This study |
| GAL4-AD-*Hs*TPR(386–539) | pGAD424 (Amp) | (Gunkel *et al.*, 2023) |
| GAL4-AD-*Hs*TPR(411–539) | pGAD424 (Amp) | This study |
| GAL4-AD-*Hs*TPR(432–539) | pGAD424 (Amp) | This study |
| GAL4-AD-*Hs*TPR(450–543) | pGAD424 (Amp) | This study |
| GAL4-AD-*Hs*TPR(608–940) | pGAD424 (Amp) | This study |
| GAL4-AD-*Hs*TPR(926–1178) | pGAD424 (Amp) | This study |
| GAL4-AD-*Hs*TPR(1129–1632) | pGAD424 (Amp) | This study |
| GAL4-AD-*Hs*TPR(1177–1632) | pGAD424 (Amp) | This study |
| GAL4-AD-*Hs*TPR(1618–1917) | pGAD424 (Amp) | This study |
| GAL4-AD-*Hs*TPR(1894–2138) | pGAD424 (Amp) | This study |
| GAL4-AD-*Hs*TPR(2110–2363) | pGAD424 (Amp) | This study |
| GAL4-BD-*Sc*Pml39p(1–334) | pGBT9 (Amp) | (Gunkel *et al.*, 2023) |
| GAL4-BD-*Sc*Pml39p(49–334) | pGBT9 (Amp) | This study |
| GAL4-BD-*Sc*Pml39p(71–334) | pGBT9 (Amp) | This study |
| GAL4-BD-*Sc*Pml39p(92–334) | pGBT9 (Amp) | This study |
| GAL4-BD-*Sc*Pml39p(1–319) | pGBT9 (Amp) | This study |
| GAL4-BD-*Sc*Pml39p(1–304) | pGBT9 (Amp) | This study |
| GAL4-AD-*Sc*Mlp1p(1–297) | pGAD424 (Amp) | (Gunkel *et al.*, 2023) |
| GAL4-AD-*Sc*Mlp1p(287–584) | pGAD424 (Amp) | (Gunkel *et al.*, 2023) |

| | | |
|---|---|---|
| GAL4-AD-*Sc*Mlp1p(1–210) | pGAD424 (Amp) | (Gunkel *et al.*, 2023) |
| EGFP-SV40-N1 | pEGFP-N1 (Kan) | |
| *Hs*TPR(1–651)-EGFP-SV40NLS | pEGFP-N1 (Kan) | This study |
| *Hs*TPR(1–334)-EGFP-SV40NLS | pEGFP-N1 (Kan) | This study |
| *Hs*TPR(1–398)-EGFP-SV40NLS | pEGFP-N1 (Kan) | This study |
| *Hs*TPR(172–651)-EGFP-SV40NLS | pEGFP-N1 (Kan) | This study |
| *Hs*TPR(172–539)-EGFP-SV40NLS | pEGFP-N1 (Kan) | This study |
| *Hs*TPR(172–481)-EGFP-SV40NLS | pEGFP-N1 (Kan) | This study |
| *Hs*TPR(172–453)-EGFP-SV40NLS | pEGFP-N1 (Kan) | This study |
| *Hs*TPR(276–651)-EGFP-SV40NLS | pEGFP-N1 (Kan) | This study |
| *Hs*TPR(275–539)-EGFP-SV40NLS | pEGFP-N1 (Kan) | This study |
| *Hs*TPR(276–481)-EGFP-SV40NLS | pEGFP-N1 (Kan) | This study |
| *Hs*TPR(450–651)-EGFP-SV40NLS | pEGFP-N1 (Kan) | This study |
| *Hs*TPR(479–651)-EGFP-SV40NLS | pEGFP-N1 (Kan) | This study |
| *Hs*TPR(432–651)-EGFP-SV40NLS | pEGFP-N1 (Kan) | This study |
| Myc-*Hs*TPR(1–479)-SV40NLS | pRc/CMV (Amp) | (Hase *et al.*, 2001) |
| Myc-*Hs*TPR(276–651)-SV40NLS | pRc/CMV (Amp) | (Hase *et al.*, 2001) |
| Myc-*Hs*TPR(276–519)-SV40NLS | pRc/CMV (Amp) | (Hase *et al.*, 2001) |
| Myc-*Hs*TPR(357–519)-EGFP-SV40NLS | pRc/CMV (Amp) | (Hase *et al.*, 2001) |
| Myc-*Hs*TPR(436–606)-EGFP-SV40NLS | pRc/CMV (Amp) | (Hase *et al.*, 2001) |
| h*Sp*Cas9n(BB)-2A-Puro | PX462 V2.0 (Amp) | Addgene plasmid #62987, gift from Feng Zhang (Ran *et al.*, 2013a, 2013b) |
| sgRNA NT1 (*Hs*TPR) / hSpCas9n(BB)-2A-Puro | PX462 V2.0 (Amp) | (Gunkel and Cordes, 2022) |
| sgRNA NT2 (*Hs*TPR) / hSpCas9n(BB)-2A-Puro | PX462 V2.0 (Amp) | (Gunkel and Cordes, 2022) |
| sgRNA NT3 (*Hs*TPR) / hSpCas9n(BB)-2A-Puro | PX462 V2.0 (Amp) | This study |
| sgRNA NT4 (*Hs*TPR) / hSpCas9n(BB)-2A-Puro | PX462 V2.0 (Amp) | This study |
| sgRNA CT1 (*Hs*TPR) / hSpCas9n(BB)-2A-Puro | PX462 V2.0 (Amp) | (Gunkel and Cordes, 2022) |
| sgRNA CT2 (*Hs*TPR) / hSpCas9n(BB)-2A-Puro | PX462 V2.0 (Amp) | (Gunkel and Cordes, 2022) |
| sgRNA NT1 (*Hs*NUP107) / hSpCas9n(BB)-2A-Puro | PX462 V2.0 (Amp) | This study |
| sgRNA NT2 (*Hs*NUP107) / hSpCas9n(BB)-2A-Puro | PX462 V2.0 (Amp) | This study |
| sgRNA NT1 (*Hs*NUP153) / hSpCas9n(BB)-2A-Puro | PX462 V2.0 (Amp) | This study |
| sgRNA NT2 (*Hs*NUP153) / hSpCas9n(BB)-2A-Puro | PX462 V2.0 (Amp) | This study |
| sgRNA CT1 /(*Hs*NUP96) hSpCas9n(BB)-2A-Puro | PX462 V2.0 (Amp) | This study |
| sgRNA CT2 (*Hs*NUP96) / hSpCas9n(BB)-2A-Puro | PX462 V2.0 (Amp) | This study |
| sgRNA CT1 (*Hs*MAD1) / hSpCas9n(BB)-2A-Puro | PX462 V2.0 (Amp) | This study |
| sgRNA CT2 (*Hs*MAD1) / hSpCas9n(BB)-2A-Puro | PX462 V2.0 (Amp) | This study |
| sgRNA CT1 (*Hs*TMEM209) / hSpCas9n(BB)-2A-Puro | PX462 V2.0 (Amp) | This study |
| sgRNA CT2 (*Hs*TMEM209) / hSpCas9n(BB)-2A-Puro | PX462 V2.0 (Amp) | This study |
| sgRNA NT1 (*Hs*ZC3HC1) / hSpCas9n(BB)-2A-Puro | PX462 V2.0 (Amp) | (Gunkel *et al.*, 2021) |
| sgRNA NT2 (*Hs*ZC3HC1) / hSpCas9n(BB)-2A-Puro | PX462 V2.0 (Amp) | (Gunkel *et al.*, 2021) |
| sgRNA CT1 (*Hs*ZC3HC1) / hSpCas9n(BB)-2A-Puro | PX462 V2.0 (Amp) | (Gunkel and Cordes, 2022) |
| sgRNA CT2 (*Hs*ZC3HC1) / hSpCas9n(BB)-2A-Puro | PX462 V2.0 (Amp) | (Gunkel and Cordes, 2022) |
| H14-ZZ-*Bd*SUMO-*Hs*TPR(354–452)-Cys | pQE80N (Kan) | This study |
| Integration vector TPR-sfGFP | pQE80N (Kan) | (Gunkel and Cordes, 2022) |
| Integration vector sfGFP-TPR | pQE80N (Kan) | (Gunkel and Cordes, 2022) |
| Integration vector sfGFP-TPR (Δ1–43) | pQE80N (Kan) | This study |
| Integration vector sfGFP-TPR (Δ1–47) | pQE80N (Kan) | This study |
| Integration vector sfGFP-TPR (Δ1–50) | pQE80N (Kan) | This study |
| Integration vector sfGFP-TPR (Δ1–58) | pQE80N (Kan) | This study |
| Integration vector sfGFP-NUP107 | pQE80N (Kan) | This study |
| Integration vector sfGFP-NUP153 | pUC57 (Amp) | This study |
| Integration vector NUP96-sfGFP | pUC57 (Amp) | (Weber *et al.*, 2023) |
| Integration vector MAD1-sfGFP | pUC57 (Amp) | This study |
| Integration vector TMEM209-sfGFP | pUC57 (Amp) | This study |
| Integration vector ZC3HC1-sfGFP | pQE80N (Kan) | (Gunkel and Cordes, 2022) |
| Integration vector ZC3HC1-sfGFP (Δ487–502) | pQE80N (Kan) | (Gunkel and Cordes, 2022) |
| Dox-inducible mCherry-*HsZC3H*C1(1–502) | pTetOne (Amp) | (Gunkel and Cordes, 2022) |
| Dox-inducible mCherry-*HsZC3H*C1(1–502 \| C429S) | pTetOne (Amp) | (Gunkel and Cordes, 2022) |
| Dox-inducible mCherry-*HsZC3H*C1(72–502) | pTetOne (Amp) | This study |
| Dox-inducible mCherry-*HsZC3H*C1(1–502 Δ26–40) | pTetOne (Amp) | This study |
| Dox-inducible mCherry-*HsZC3H*C1(1–290_398–502) | pTetOne (Amp) | This study |
| Dox-inducible mCherry-*HsZC3H*C1(1–497) | pTetOne (Amp) | This study |
| Dox-inducible mCherry-*HsZC3H*C1(1–492) | pTetOne (Amp) | This study |
| Dox-inducible mCherry-*HsZC3H*C1(1–486) | pTetOne (Amp) | This study |
| Dox-inducible mCherry-*HsZC3H*C1(1–478) | pTetOne (Amp) | This study |
| Dox-inducible mCherry-*HsZC3H*C1(1–472) | pTetOne (Amp) | This study |
| Dox-inducible mCherry-*HsZC3H*C1(1–462) | pTetOne (Amp) | This study |
| Dox-inducible mCherry-*HsZC3H*C1(1–502 Δ491–499) | pTetOne (Amp) | This study |
| Dox-inducible mCherry-*HsZC3H*C1(1–502 Δ483–499) | pTetOne (Amp) | This study |
| Dox-inducible mCherry-*HsZC3H*C1(72–472) | pTetOne (Amp) | This study |
| Dox-inducible mCherry-*HsZC3H*C1(1–502 Δ26–40 & Δ483–499) | pTetOne (Amp) | This study |

| | | |
|---|---|---|
| mCherry-*HsZC3H*C1(1–502) | pEYFP-C1 (Kan) | (Gunkel and Cordes, 2022) |
| mCherry-*HsZC3H*C1(1–502 \| C429S) | pEYFP-C1 (Kan) | (Gunkel and Cordes, 2022) |
| mCherry-*HsZC3H*C1(72–502) | pEYFP-C1 (Kan) | This study |
| mCherry-*HsZC3H*C1(1–502 Δ26–40) | pEYFP-C1 (Kan) | This study |
| mCherry-*HsZC3H*C1(1–290 398–502) | pEYFP-C1 (Kan) | This study |
| mCherry-*HsZC3H*C1(1–497) | pEYFP-C1 (Kan) | This study |
| mCherry-*HsZC3H*C1(1–492) | pEYFP-C1 (Kan) | This study |
| mCherry-*HsZC3H*C1(1–486) | pEYFP-C1 (Kan) | This study |
| mCherry-*HsZC3H*C1(1–478) | pEYFP-C1 (Kan) | This study |
| mCherry-*HsZC3H*C1(1–472) | pEYFP-C1 (Kan) | This study |
| mCherry-*HsZC3H*C1(1–462) | pEYFP-C1 (Kan) | This study |
| mCherry-*HsZC3H*C1(1–502 Δ491–499) | pEYFP-C1 (Kan) | This study |
| mCherry-*HsZC3H*C1(1–502 Δ483–499) | pEYFP-C1 (Kan) | This study |
| mCherry-*HsZC3H*C1(72–472) | pEYFP-C1 (Kan) | This study |
| mCherry-*HsZC3H*C1(1–502 Δ26–40 & Δ483–499) | pEYFP-C1 (Kan) | This study |

#### Supplemental Table S3: Cell lines.

| **Cell line** | **Source** |
|---|---|
| HeLa P2 WT | (Gunkel *et al.*, 2021) |
| HeLa P2 ZC3HC1 KO | (Gunkel *et al.*, 2021) |
| HeLa P2 TPR-sfGFP (ZC3HC1 KO) | (Gunkel and Cordes, 2022) |
| HeLa P2 sfGFP-TPR (ZC3HC1 KO) | (Gunkel and Cordes, 2022) |
| HeLa P2 TPR-sfGFP | (Gunkel and Cordes, 2022) |
| HeLa P2 sfGFP-TPR | this study |
| HeLa P2 sfGFP-TPR (Δ1–43) | this study |
| HeLa P2 sfGFP-TPR (Δ1–47) | this study |
| HeLa P2 sfGFP-TPR (Δ1–50) | this study |
| HeLa P2 sfGFP-TPR (Δ1–58) | this study |
| HeLa P2 ZC3HC1-sfGFP | this study |
| HeLa P2 ZC3HC1-sfGFP (Δ487–502) | this study |
| HeLa P2 sfGFP-NUP107 | this study |
| HeLa P2 sfGFP-NUP107; NUP96-sfGFP | this study |
| HeLa P2 sfGFP-NUP153 | this study |
| HeLa P2 MAD1-sfGFP | this study |
| HeLa P2 TMEM209-sfGFP | this study |
| IMR-90 (RRID: CVCL_0347) | (Nichols *et al.*, 1977) |
| IMR-90 ZC3HC1 KO (enriched population) | this study |
| HeLa Kyoto (RRID: CVCL_1922) | Shuh Narumiya (Kyoto University, Japan)<br>Kindly provided by Jan Ellenberg |
| HeLa Kyoto ZC3HC1 KO | this study |

#### Supplemental Table S4: sgRNA sequences.

| **Description** | **Vector backbone** | **gRNA sequence (sense)** | **Purpose** |
|---|---|---|---|
| *Hs*TPR sgRNA NT1 | PX462 V2.0 | GGGGCGGCATGAGAAATTTA | sfGFP integration (NT, Exon 1) |
| *Hs*TPR sgRNA NT2 | PX462 V2.0 | gTCTCGGATTGCTGATCAGCA | sfGFP integration (NT, Exon 1) |
| *Hs*TPR sgRNA NT3 | PX462 V2.0 | gCCCGAGAGTGTCAAAGCTTG | sfGFP integration (NT, Exon 2) |
| *Hs*TPR sgRNA NT4 | PX462 V2.0 | gCACAAGTCTCTCCTGACTGT | sfGFP integration (NT, Exon 2) |
| *Hs*TPR sgRNA CT1 | PX462 V2.0 | gCAGAGGAAATATTAATTAAA | sfGFP integration (CT, Exon 51) |
| *Hs*TPR sgRNA CT2 | PX462 V2.0 | gCTCCTCTCCCTCCCATTGCA | sfGFP integration (CT, Exon 51) |
| *Hs*NUP107 sgRNA NT1 | PX462 V2.0 | GTCTTGCCCGTCTCGCCTGT | sfGFP integration (NT, Exon 1) |
| *Hs*NUP107 sgRNA NT2 | PX462 V2.0 | gCATCAGTACTGACCTGTCCA | sfGFP integration (NT, Exon 1) |
| *Hs*NUP153 sgRNA NT1 | PX462 V2.0 | gTCCGCCATGGCCTCGGGAGC | sfGFP integration (NT, Exon 1) |
| *Hs*NUP153 sgRNA NT2 | PX462 V2.0 | gTTCCCGCTCCGGGGCGGGTA | sfGFP integration (NT, Exon 1) |
| *Hs*NUP196 sgRNA CT1 | PX462 V2.0 | GTTGGGAGCCTGTGAGCCCC | sfGFP integration (CT, Exon 33) |
| *Hs*NUP196 sgRNA CT2 | PX462 V2.0 | gCTCGCAGATAGGACTGGGTA | sfGFP integration (CT, Exon 33) |
| *Hs*MAD1L1 sgRNA CT1 | PX462 V2.0 | gATAGCCGGAGCCACTCTGCT | sfGFP integration (CT, Exon 19) |
| *Hs*MAD1L1 sgRNA CT2 | PX462 V2.0 | gCGAGCCTGCAGGCTACGCCA | sfGFP integration (CT, Exon 19) |
| *Hs*TMEM209 sgRNA CT1 | PX462 V2.0 | gCTTTATAGGAGAGTTAATCT | sfGFP integration (CT, Exon 15) |
| *Hs*TMEM209 sgRNA CT2 | PX462 V2.0 | gAATTGGTTGCCAAGCATTTA | sfGFP integration (CT, Exon 15) |
| *Hs*ZC3HC1 sgRNA NT1 | PX462 V2.0 | GCAGTAGTTCGCTCCCCAGA | knock-out (Exon 1) |
| *Hs*ZC3HC1 sgRNA NT2 | PX462 V2.0 | GCAAACGCTTGTCCCTCACA | knock-out (Exon 1) |
| *Hs*ZC3HC1 sgRNA CT1 | PX462 V2.0 | GAAGATACTCCAGCGCCTTCC | sfGFP integration (CT, Exon 10) |
| *Hs*ZC3HC1 sgRNA CT2 | PX462 V2.0 | GATTCCCACTGCCGAAATATT | sfGFP integration (CT, Exon 10) |

#### Supplemental Table S5: Primers for genotyping.

| Target | Forward primer | Reverse primer |
|---|---|---|
| *Hs*TPR-CT | CCATTTACCCTTTTTCGGTTTGTTAAATGTTCC | CATTGATGTGCCTAGTAGAACAGCAAGAG |
| *Hs*TMEM209-CT | AGACCTTAAATGCTTGGCAACCAATTC | TCCTGATGTCCTCACACTCTTAAC |
| *Hs*NUP196-CT | GTTGGTTCTGGCTGCATTTTTTACTTCC | GGTCACAAGATCCAGAATGGCTAGGG |
| *Hs*NUP107-NT | GCCAGCAACAGAAAAGACTATATCAACAGG | GAGTCTAGTTTCACCTATTTCGTTTCGCC |
| *Hs*NUP153-NT | CCGCCTTAGCAGGGGTAATAG | CCAGGAGTTCATACAGTGGCT |
| *Hs*MAD1L1-CT | CAGAGTTCTCACACACCGTGG | GCACCTCTGAATCATCGGCTC |
| *Hs*ZC3HC1-CT | GTCCTAGACATATCCAGATCATTGTGGG | CTCAGTGACGAAAAACATGGCATATCAGAG |
| *Hs*TPR-NT (for1-rev1) | TGGCAGACAATTACCTCGGACGGTC | CAAGGACGGAAAGAGCATCTCACTTAC |
| *Hs*TPR-NT (for1-rev2) | TGGCAGACAATTACCTCGGACGGTC | GTCTCTGAAGATTTCGATACTCCG |

#### Supplemental Table S6: siRNAs.

| Target | Sense strand (5'-3') | Source (order number) |
|---|---|---|
| Silencer Select negative control #1 | proprietary information | Ambion (4390843) |
| Silencer Select negative control #2 | proprietary information | Ambion (4390846) |
| ON-TARGETplus Non-targeting pool | proprietary information | Dharmacon (D-001810-10) |
| *Hs*PABPN1-1 | GGCCUUAGAUGAGUCCCUA | Ambion (s15626) |
| *Hs*PABPN1-2 | AGUCAACCGUGUUACCAUA | Ambion (s15626) |
| *Hs*PABPN1-3 | CGUAUAUAGAGUUCUCAGA | Ambion (s15626) |
| *Hs*PABPC1-1 | AUCCAUUGAUAAUAAAGCA | Ambion (s25564) |
| *Hs*PABPC1-2 | CCUAAAUGAUCGCAAAGUA | Ambion (s25565) |
| *Hs*PABPC1-3 | UAGGCAACAUAUUCAUUAA | Ambion (s25566) |
| *Hs*TPR-1 | CAAUAUGAAGGUCGAAUUA | Ambion (s14355) |
| *Hs*TPR-2 | GAAGUUCAUACUAAGCGUA | Ambion (s14354) |
| *Hs*TPR-3 | GAGUCUGCGUUAUCGACAA | Ambion (s14353) |
| *Hs*TPR-4 (ON-TARGETplus Human TPR (7175) siRNA – SMARTpool) | proprietary information | Dharamacon (L-010548-00) |
| *Hs*ZC3HC1-1 | GAACAACCUUCAUUGGAAU | Ambion (s28276) |
| *Hs*ZC3HC1-2 | CAAGGAAAGUAUUCCGAAU | Ambion (s28277) |
| *Hs*ZC3HC1-3 | CACUCGUCUGUGCAAAAUA | Ambion (s28278) |
| *Hs*ZC3HC1-4 (ON-TARGETplus Human ZC3HC1 (51530) siRNA – SMARTpool) | proprietary information | Dharamacon (L-016879-02) |

#### Supplemental Table S7: Antibodies.

| Antibody (target region) | Source |
|---|---|
| Mouse monoclonal anti-*Hs*MAD1 (481–718*), clone BB3-8 | Merck Millipore (Burlington, MA, USA; MABE867)<br>* region mapped (Gunkel and Cordes, 2022) |
| Mouse monoclonal anti-*Hs*TPR, clone 203-37 (1462–1500) | (Cordes *et al.*, 1997b; Hase *et al.*, 2001; Gunkel *et al.*, 2021) |
| Mouse monoclonal anti-*Hs*ZC3HC1, clone B-10 (1–60*) | Santa Cruz Biotechnology (Dallas, TX, USA; sc-365058)<br>* epitope mapped (Gunkel *et al.*, 2021) |
| Rabbit monoclonal anti-Myc-tag, clone 71D10 | Cell Signaling Technology, Danvers, MA, USA (#2278) |
| Rabbit monoclonal anti-*Hs*PABPN1, clone EP3000Y (1–100) | Abcam (Cambridge, United Kingdom; ab75855) |
| Rabbit monoclonal anti-*Hs*SENP1, clone EPR3844 | Abcam (Cambridge, United Kingdom; ab108981) |
| Rabbit polyclonal anti-*Hs*GANP (1000–1012) | (Gunkel and Cordes, 2022) |
| Rabbit polyclonal anti-*Hs*HSF1 | Cell Signaling Technology (#4356) |
| Rabbit polyclonal anti-*Hs*NUP96 (880–900) | (Hase *et al.*, 2001) |
| Rabbit polyclonal anti-*Hs*NUP107 (1–338) | Proteintech (19217-1-AP) |
| Rabbit polyclonal anti-*Hs*NUP153 (50–100) | Abcam, Cambridge, UK (ab84872);<br>Bethyl Laboratories, Montgomery, TX, USA (A301-788A) |
| Rabbit polyclonal anti-*Xl*NUP153 (208–225) | This study |
| Rabbit polyclonal anti-*Hs*PABPC1 (520–636) | Abcam (Cambridge, United Kingdom; ab21060) |
| Rabbit polyclonal anti-*Hs*TPR-L (32–53) | This study |
| Rabbit polyclonal anti-*Hs*TPR (10–27) | This study |
| Rabbit polyclonal anti-*Hs*TPR (327–380) | This study |
| Rabbit polyclonal anti-*Hs*TPR (381–398) | (Gunkel and Cordes, 2022) |
| Rabbit polyclonal anti-*Hs*TPR (496–506) | This study |
| Rabbit polyclonal anti-*Xl*TPR (9–25) | (Gunkel *et al.*, 2021) |
| Rabbit polyclonal anti-*Xl*TPR (363–380) | This study |
| Rabbit polyclonal anti-*Xl*TPR (390–407) | This study |
| Rabbit polyclonal anti-*Xl*TPR (1619–1640) | This study |
| Rabbit polyclonal anti-*Xl*ZC3HC1 (282–392) | (Gunkel *et al.*, 2021) |
| Rabbit polyclonal anti-*Xl*ZC3HC1 (459–477) | (Gunkel *et al.*, 2021) |

| | |
|---|---|
| Guinea pig polyclonal anti-*Hs*NUP107 (33–51) | (Hase *et al.*, 2001) |
| Guinea pig polyclonal anti-*Hs*NUP133 (566–582) | (Krull *et al.*, 2010) |
| Guinea pig polyclonal anti-*Hs*NUP153 (21–36) | (Ferrando-May *et al.*, 2001) |
| Guinea pig polyclonal anti-*Hs*TPR (2063–2084) | (Cordes *et al.*, 1997b) |
| Guinea pig polyclonal anti-*Hs*ZC3HC1 (1–44) | (Gunkel *et al.*, 2021) |
| Guinea pig polyclonal anti-*Hs*ZC3HC1 (307–355) | (Gunkel *et al.*, 2021) |
| Anti-*Hs*TMEM209 nanobody, clone BM10D02, conjugated with Alexa647 | Kindly provided by Dirk Görlich |
| FluoTag-X4 anti-GFP-tag sdAb conjugated with STAR 635P | NanoTag Biotechnologies GmbH (Göttingen, Germany; N0304) |
| F(ab')2-Goat anti-Rabbit IgG (H+L) Cross-Adsorbed Secondary Antibody, conjugated with Alexa594 PLUS | Invitrogen (Carlsbad, CA, USA; A48284) |
| Donkey polyclonal anti-guinea pig IgG (H&L, minimal cross reactions) conjugated with Alexa488, Cy3, Cy5 or HRP | Jackson ImmunoResearch (706-545-148, 706-165-148, 706-175-148 or 706-035-148) |
| Donkey polyclonal anti-mouse IgG (H&L, minimal cross reactions) conjugated with Alexa488, Cy3, Cy5 or HRP | Jackson ImmunoResearch (715-545-150, 715-165-150, 715-175-150 or 715-035-150) |
| Donkey polyclonal anti-rabbit IgG (H&L, minimal cross reactions) conjugated with Alexa488, Cy3, Cy5 or HRP | Jackson ImmunoResearch (711-545-152, 711-165-152, 711-175-152 or 711-035-152) |
